## Supplementary figures and images for "Chemo- and optogenetic activation of hypothalamic Foxb1-expressing neurons and their terminal endings in the rostral-dorsolateral PAG leads to tachypnea, bradycardia, and immobility"

### SupplementalFile_S1

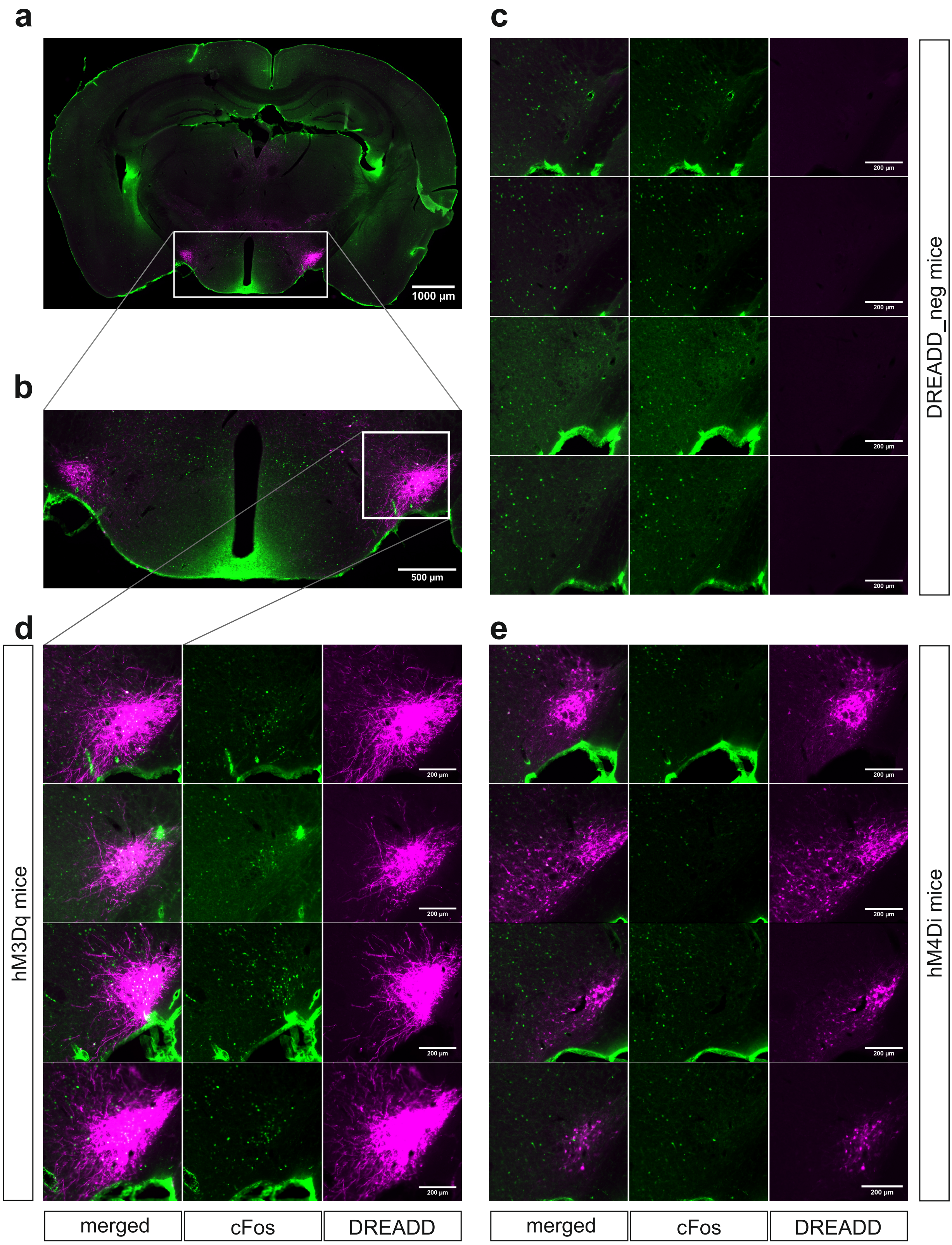
