## SupplementaryFile_S2 for "Chemo- and optogenetic activation of hypothalamic Foxb1-expressing neurons and their terminal endings in the rostral-dorsolateral PAG leads to tachypnea, bradycardia, and immobility"

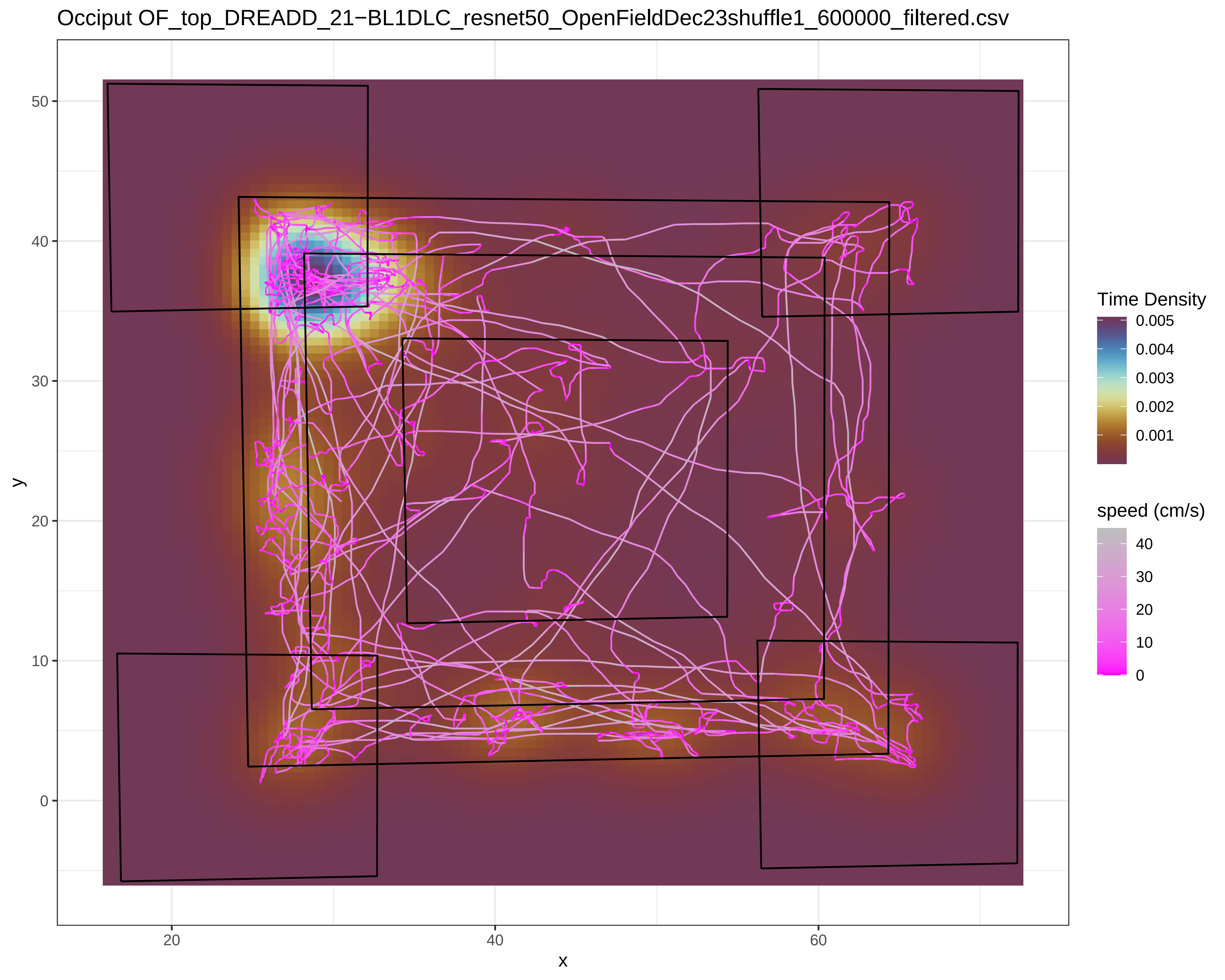

Occiput OF_top_DREADD_21−BL1DLC_resnet50_OpenFieldDec23shuffle1_600000_filtered.csv
y
50
40
30
20
10
0
x
20
40
60
Time Density
0.005
0.004
0.003
0.002
0.001
speed (cm/s)
40
30
20
10
0

Occiput OF_top_DREADD_21−BL2DLC_resnet50_OpenFieldDec23shuffle1_600000_filtered.csv

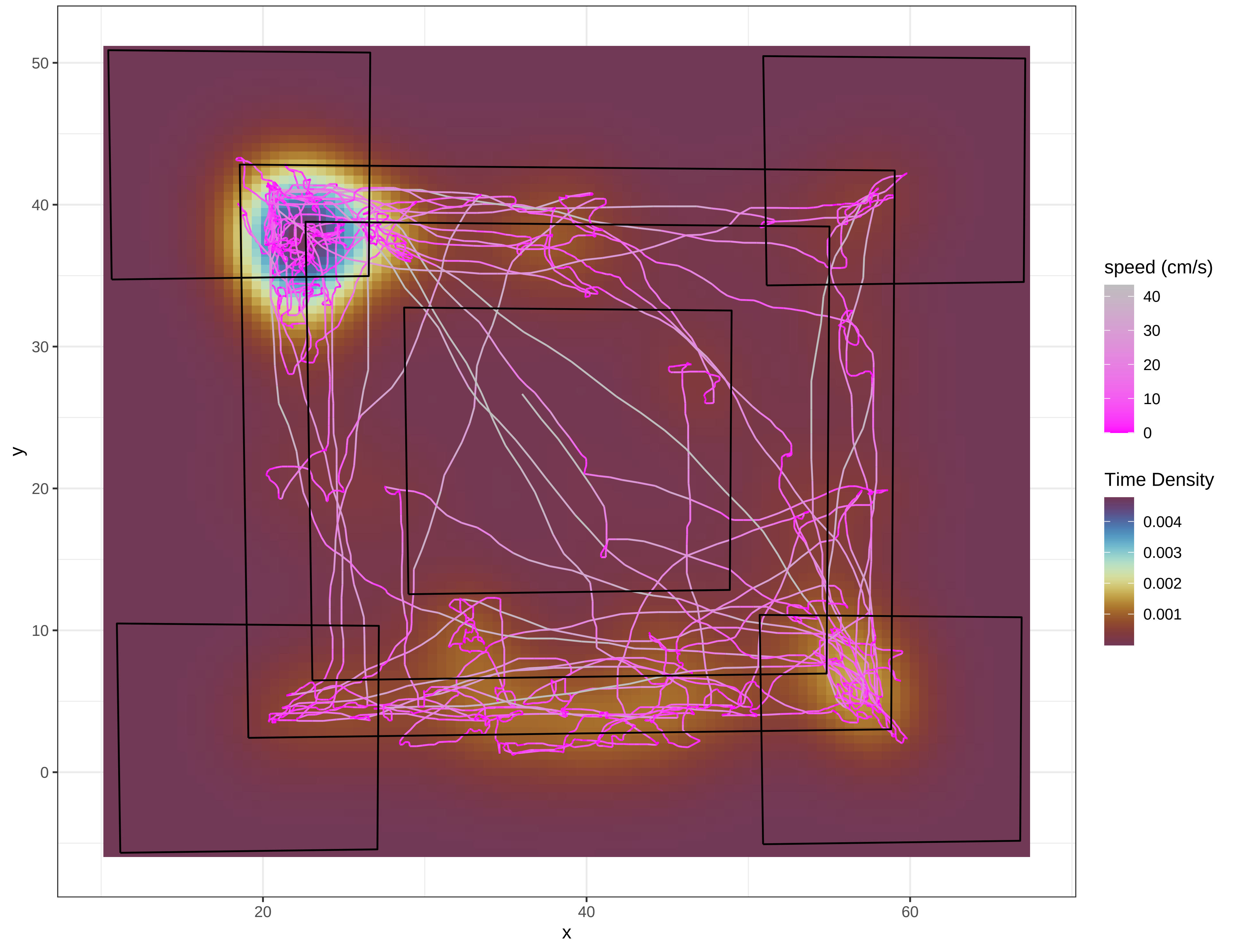

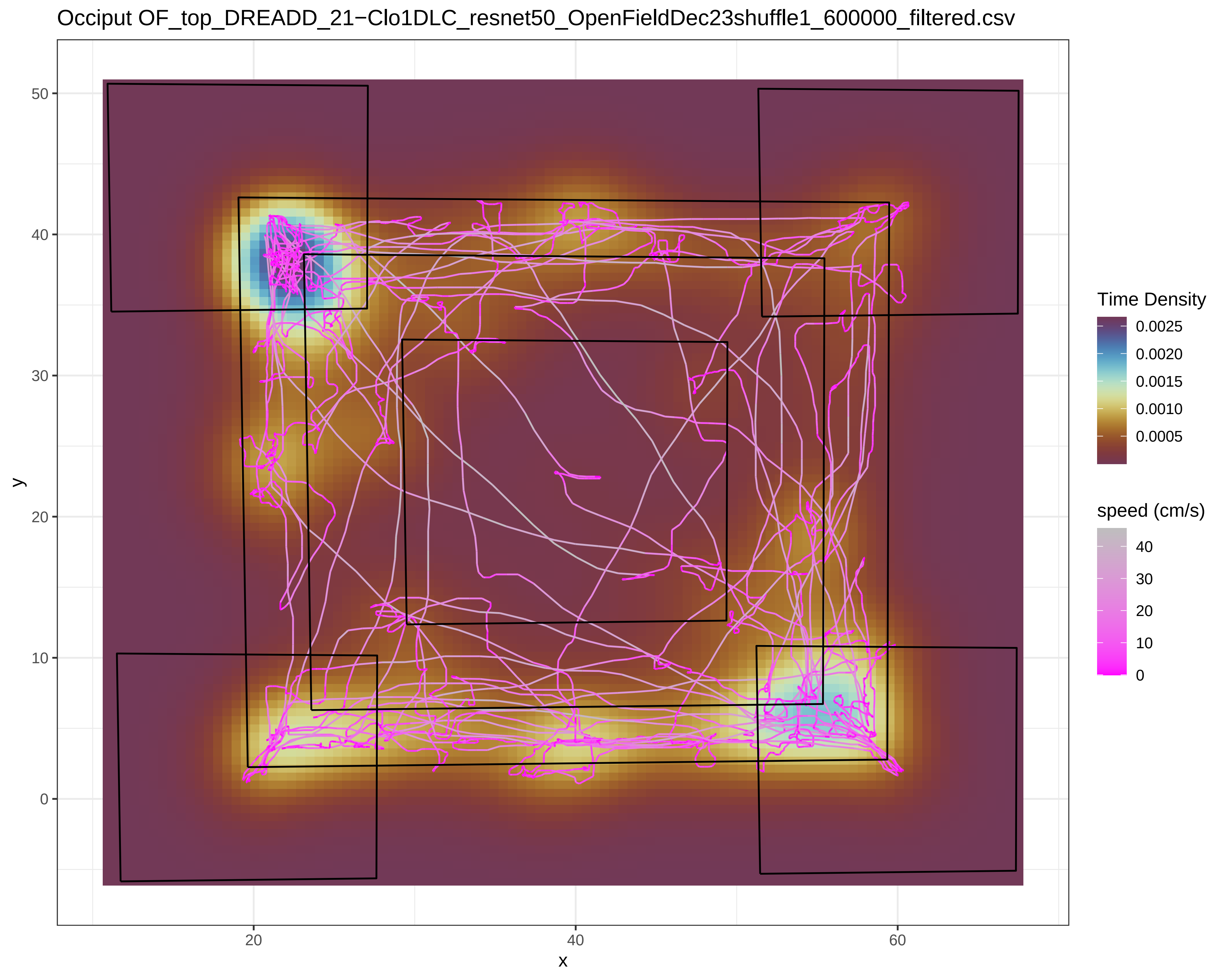

Occiput OF_top_DREADD_21−Clo1DLC_resnet50_OpenFieldDec23shuffle1_600000_filtered.csv
Time Density
0.0025
0.0020
0.0015
0.0010
0.0005
speed (cm/s)
40
30
20
10
0
y
x
50
40
30
20
10
0
20
40
60

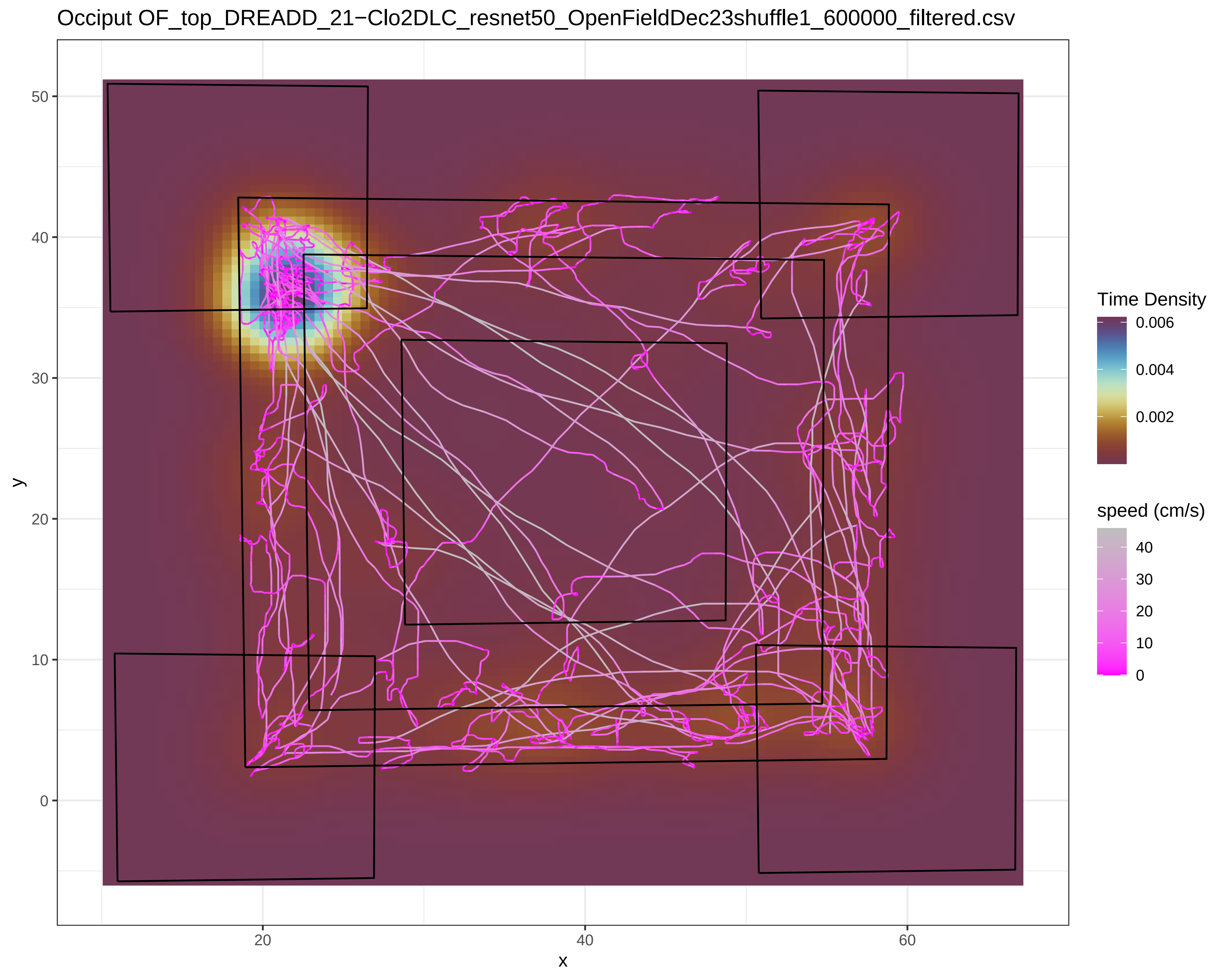

Occiput OF_top_DREADD_21−Clo2DLC_resnet50_OpenFieldDec23shuffle1_600000_filtered.csv
y
50
40
30
20
10
0
x
20
40
60
Time Density
0.006
0.004
0.002
speed (cm/s)
40
30
20
10
0

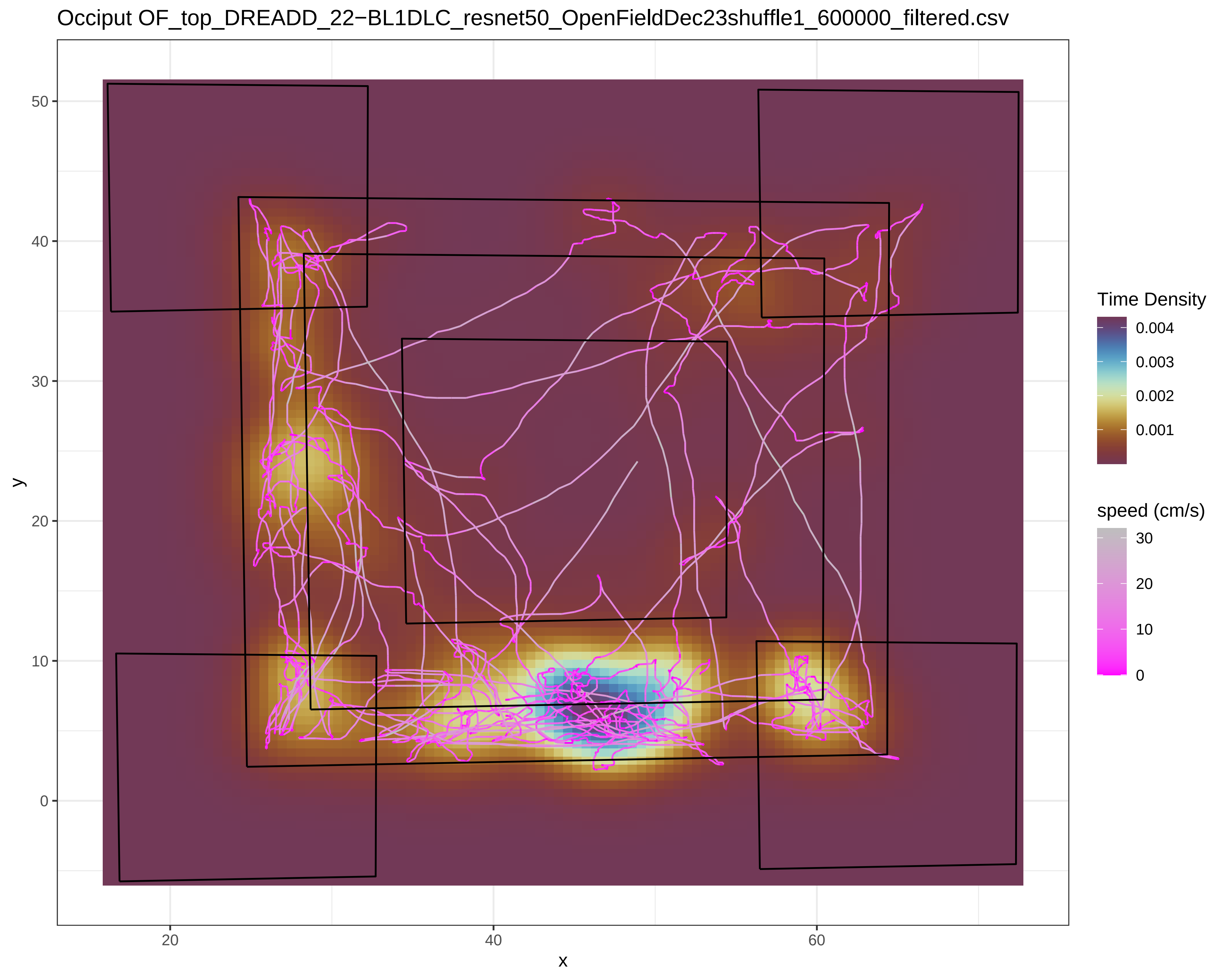
Occiput OF_top_DREADD_22−BL1DLC_resnet50_OpenFieldDec23shuffle1_600000_filtered.csv
50
40
30
20
10
0
y
20
40
60
x
Time Density
0.004
0.003
0.002
0.001
speed (cm/s)
30
20
10
0

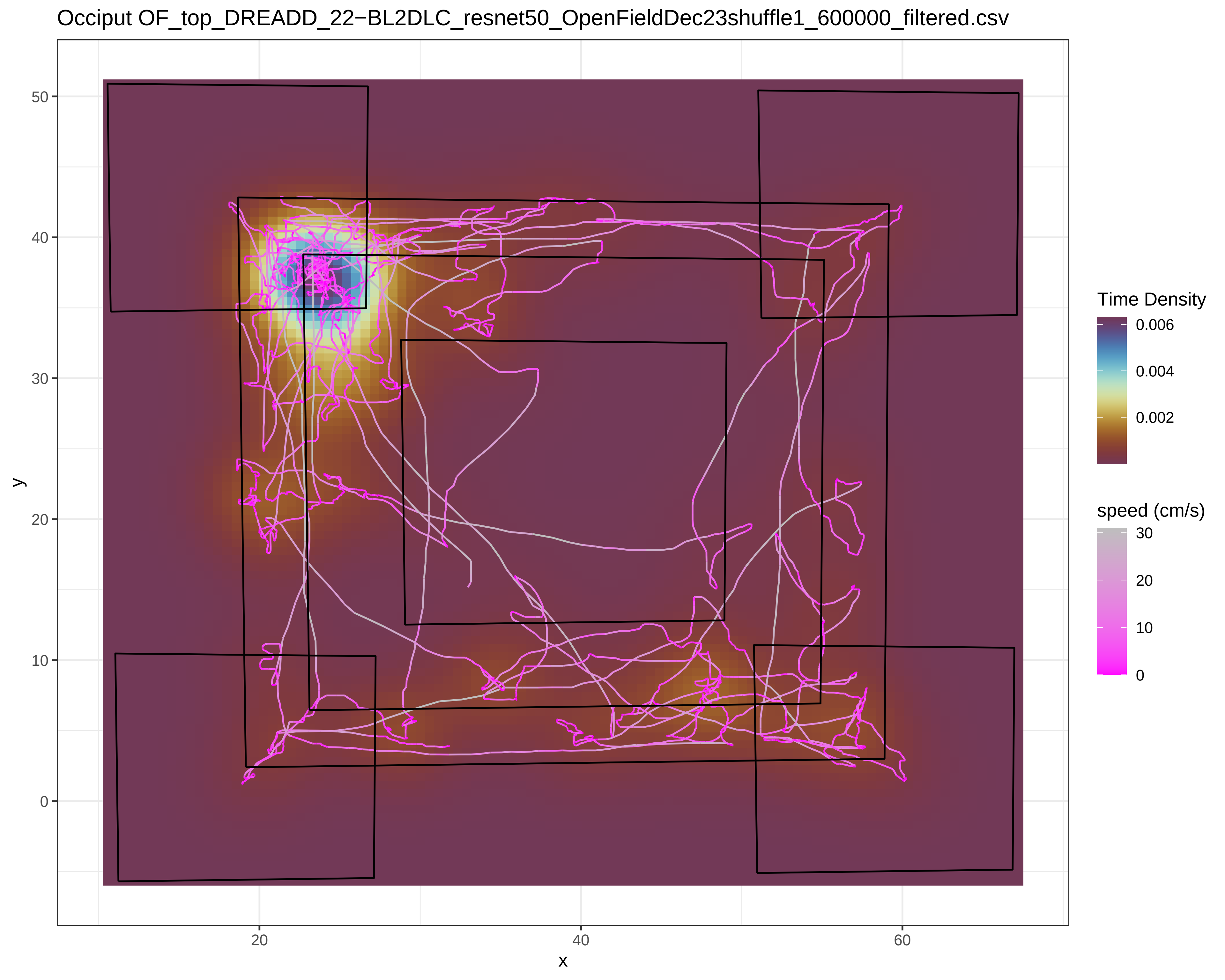
Occiput OF_top_DREADD_22−BL2DLC_resnet50_OpenFieldDec23shuffle1_600000_filtered.csv
y
x
50
40
30
20
10
0
20
40
60
Time Density
0.006
0.004
0.002
speed (cm/s)
30
20
10
0

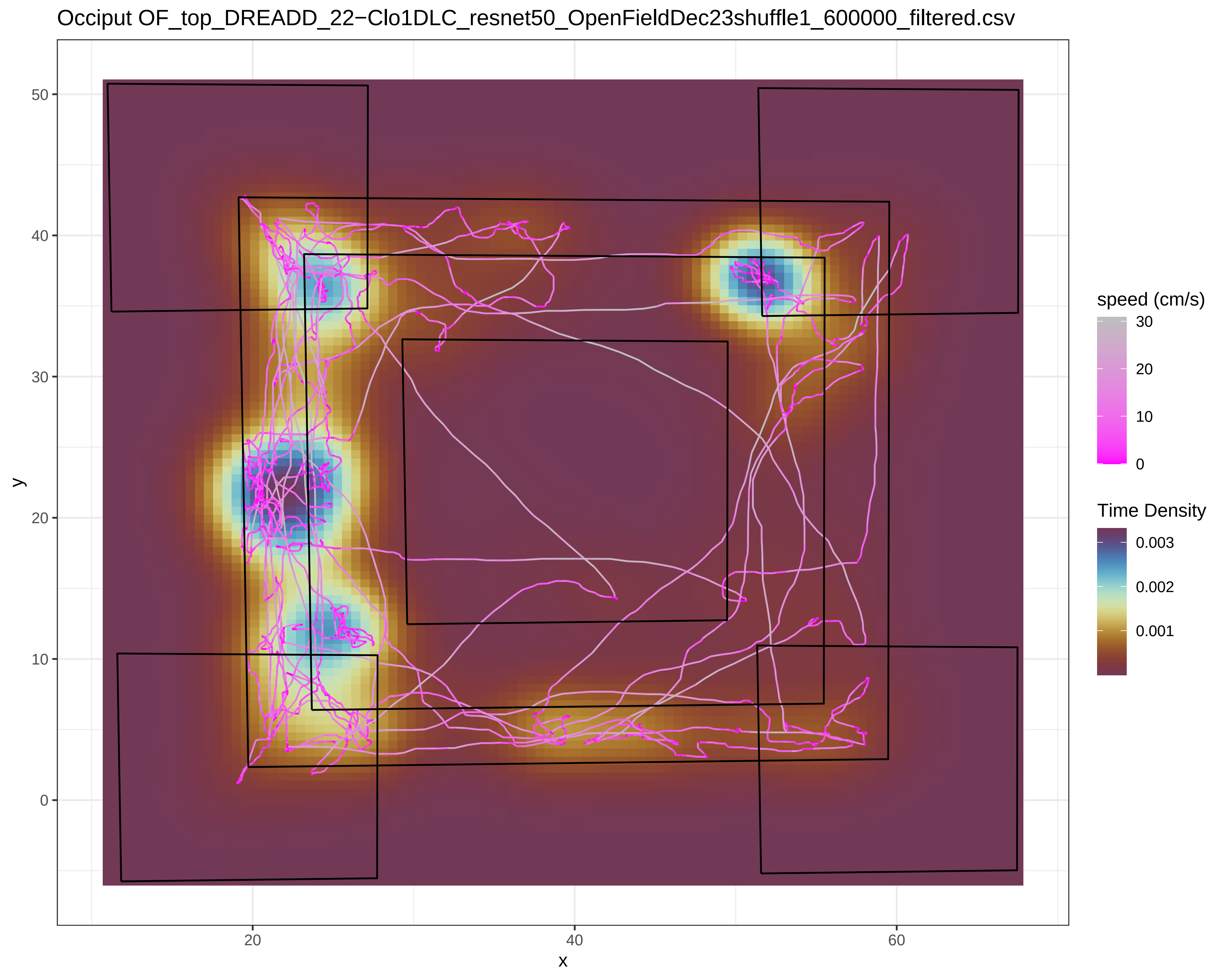
Occiput OF_top_DREADD_22−Clo1DLC_resnet50_OpenFieldDec23shuffle1_600000_filtered.csv
50
40
30
20
10
0
y
20
40
60
x
speed (cm/s)
30
20
10
0
Time Density
0.003
0.002
0.001

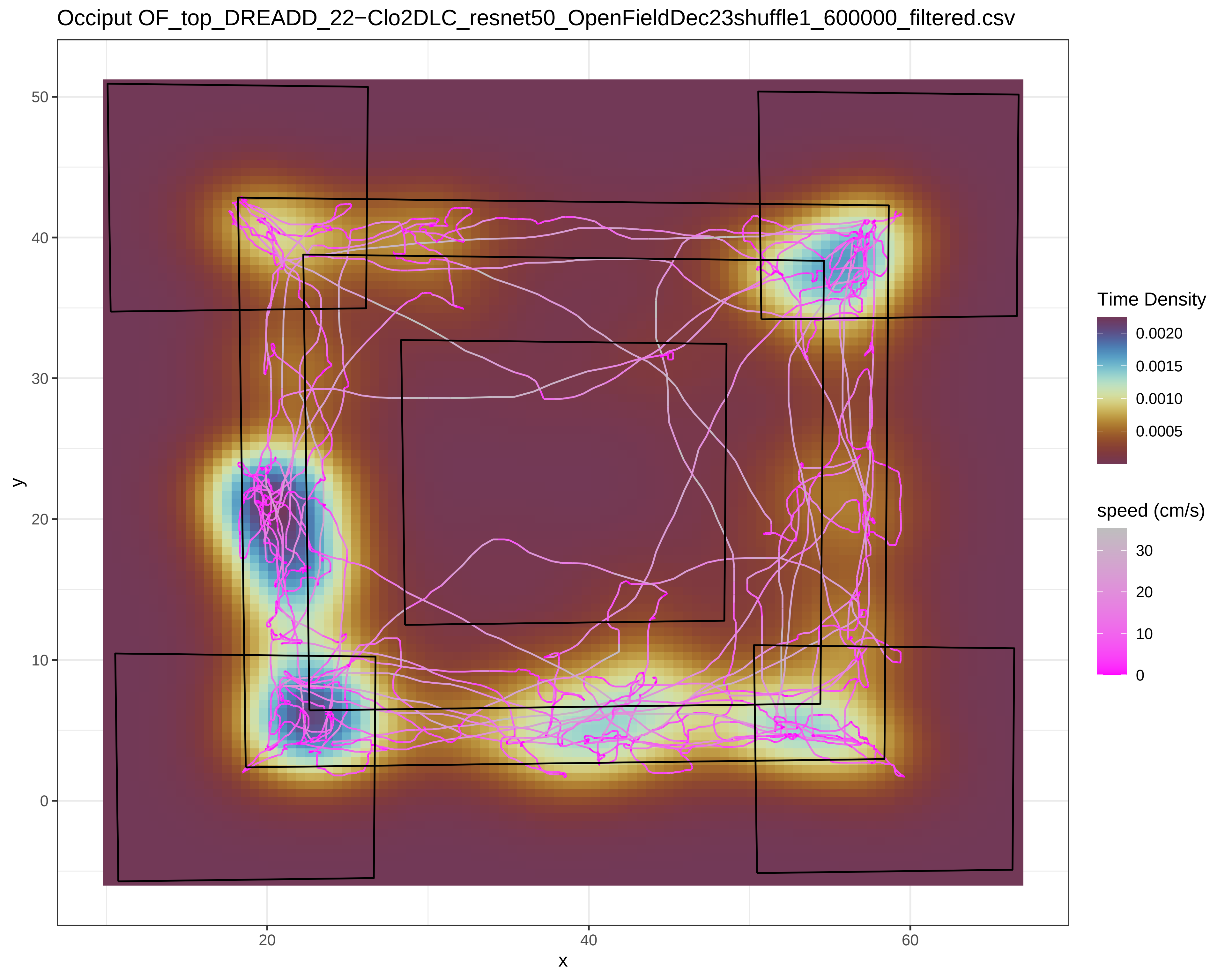

Occiput OF_top_DREADD_22−Clo2DLC_resnet50_OpenFieldDec23shuffle1_600000_filtered.csv
y
50
40
30
20
10
0
x
20
40
60
Time Density
0.0020
0.0015
0.0010
0.0005
speed (cm/s)
30
20
10
0

Occiput OF_top_DREADD_25−BL1DLC_resnet50_OpenFieldDec23shuffle1_600000_filtered.csv

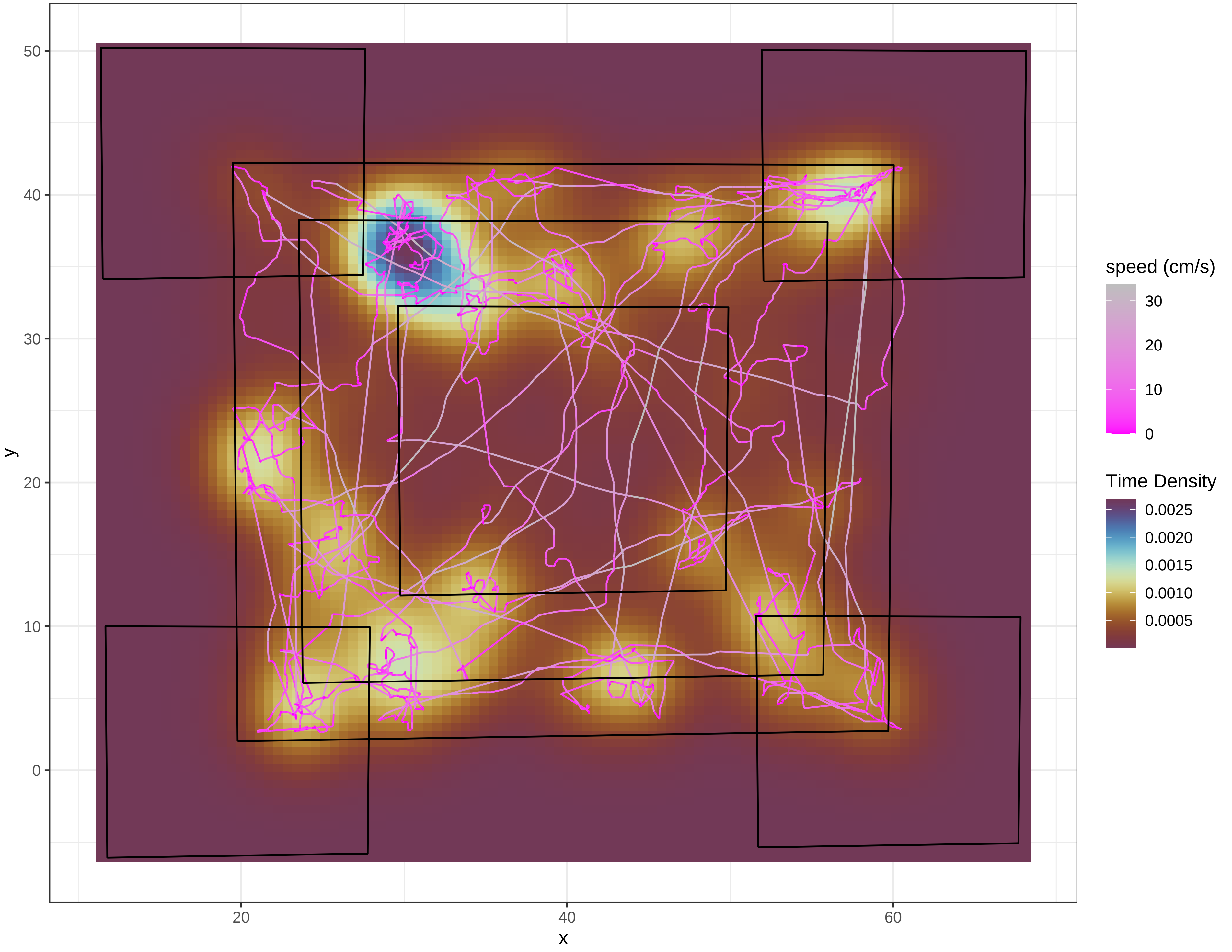

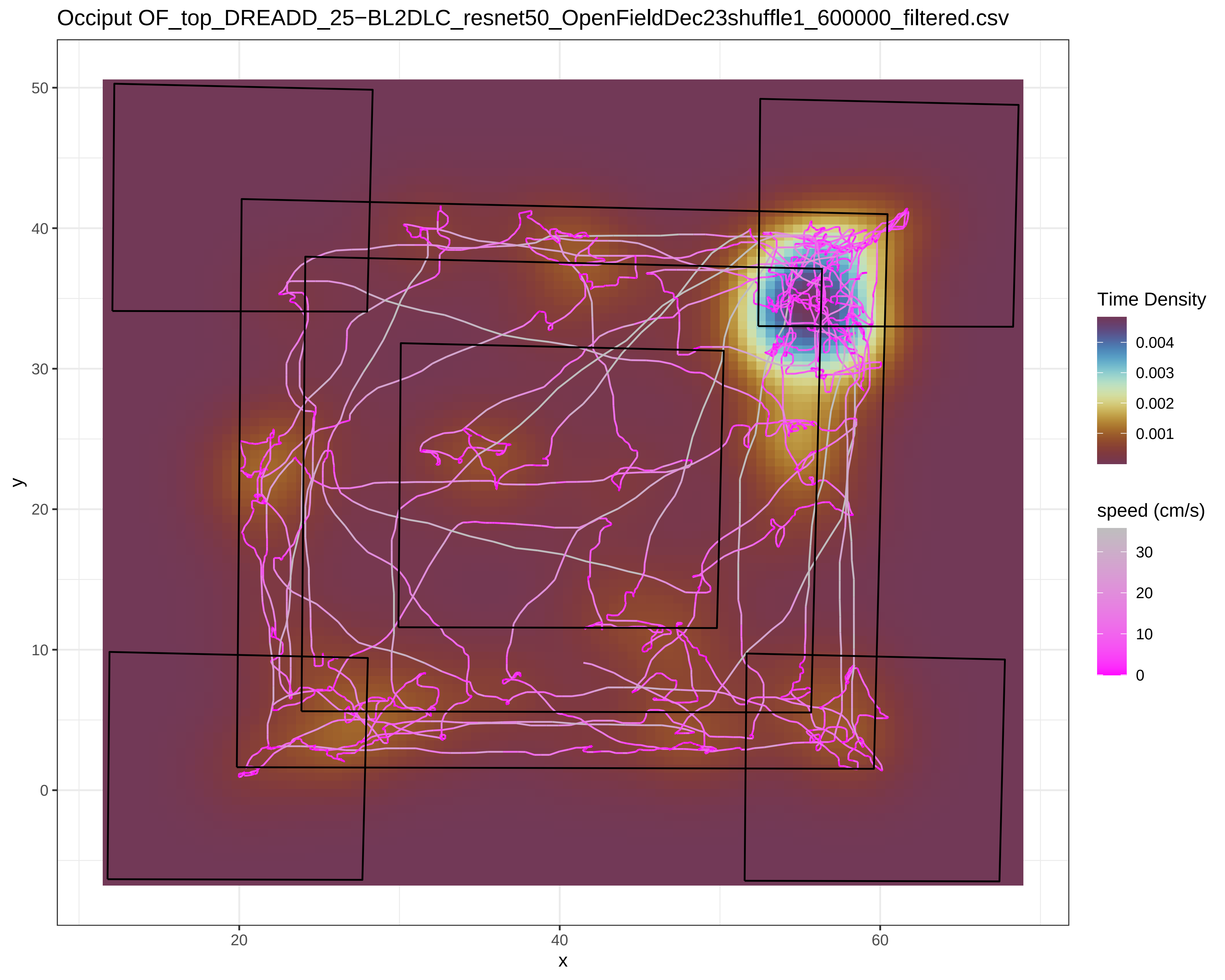

Occiput OF_top_DREADD_25−BL2DLC_resnet50_OpenFieldDec23shuffle1_600000_filtered.csv
y
x
50
40
30
20
10
0
20
40
60
Time Density
0.004
0.003
0.002
0.001
speed (cm/s)
30
20
10
0

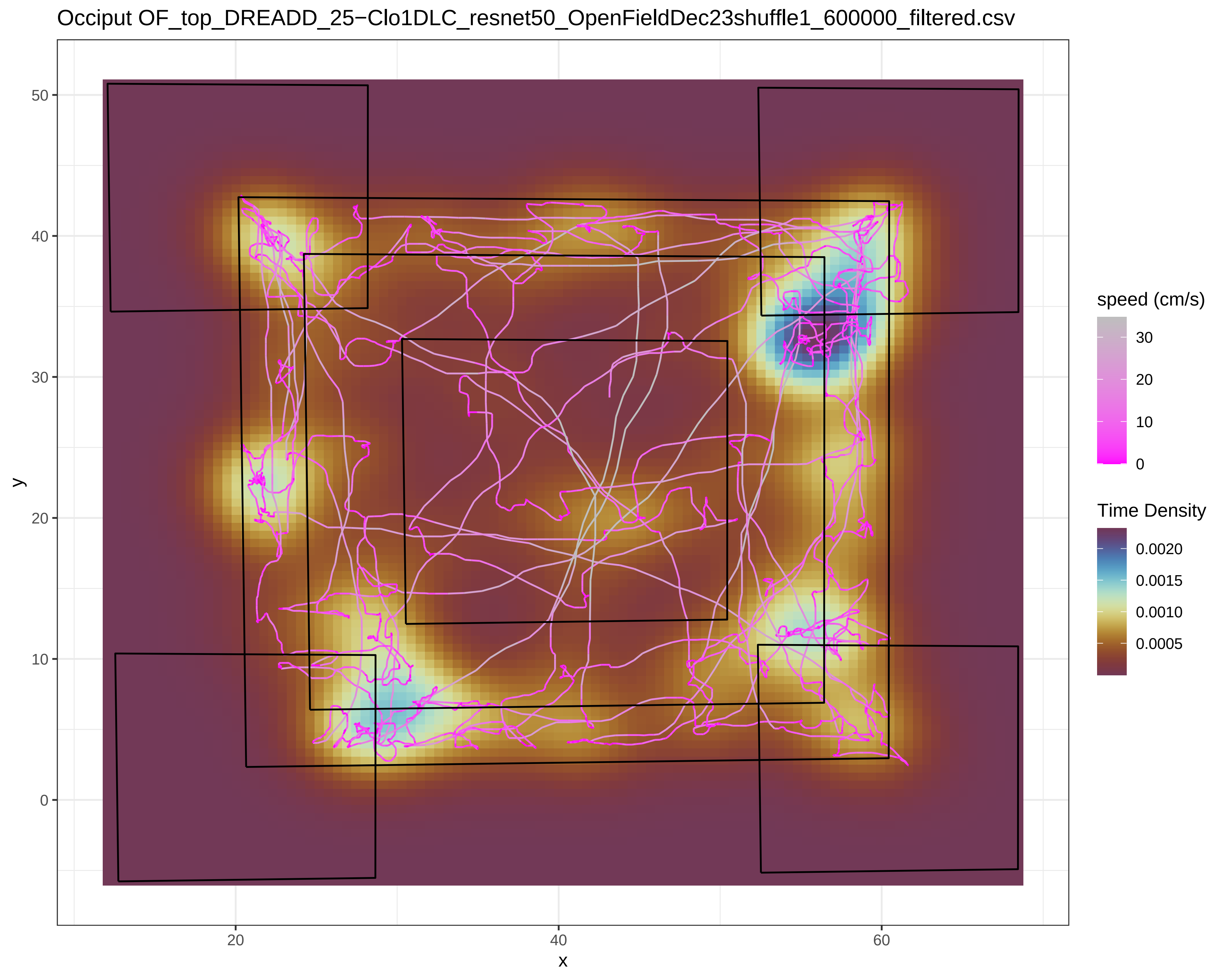

Occiput OF_top_DREADD_25−Clo1DLC_resnet50_OpenFieldDec23shuffle1_600000_filtered.csv
y
x
50
40
30
20
10
0
20
40
60
speed (cm/s)
30
20
10
0
Time Density
0.0020
0.0015
0.0010
0.0005

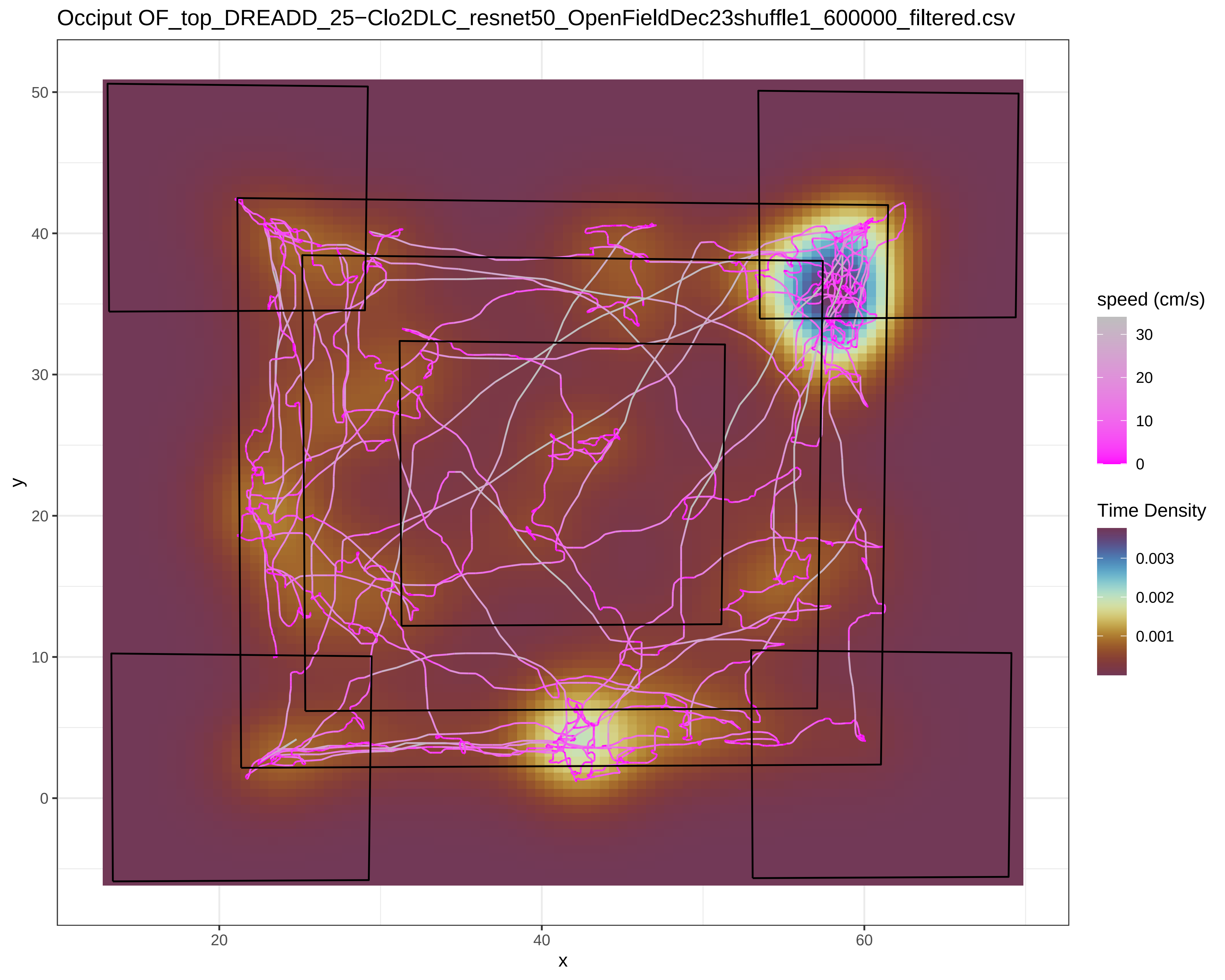

Occiput OF_top_DREADD_25−Clo2DLC_resnet50_OpenFieldDec23shuffle1_600000_filtered.csv
y
x
0
10
20
30
40
50
20
40
60
speed (cm/s)
30
20
10
0
Time Density
0.003
0.002
0.001

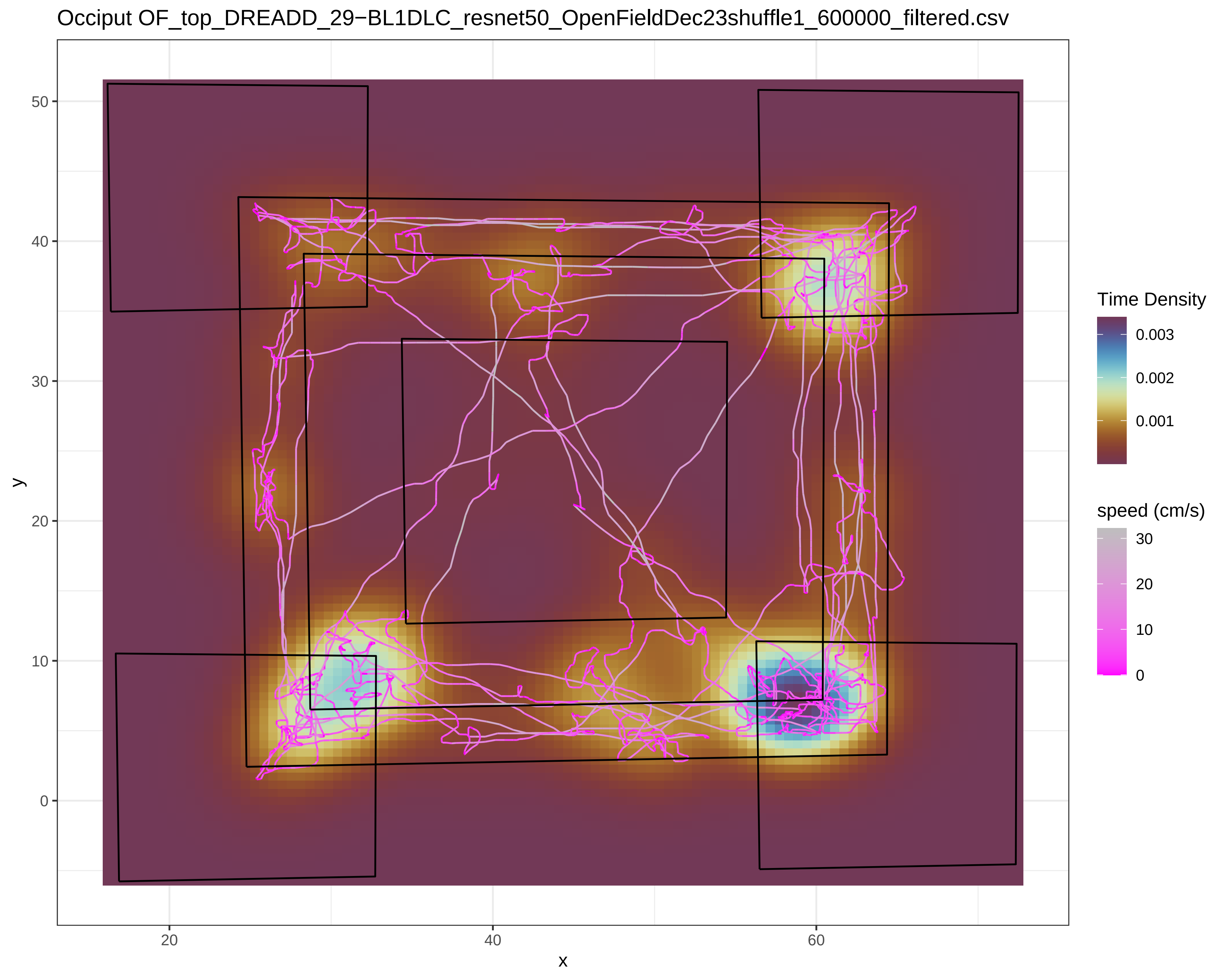

Occiput OF_top_DREADD_29−BL1DLC_resnet50_OpenFieldDec23shuffle1_600000_filtered.csv
y
x
50
40
30
20
10
0
20
40
60
Time Density
0.003
0.002
0.001
speed (cm/s)
30
20
10
0

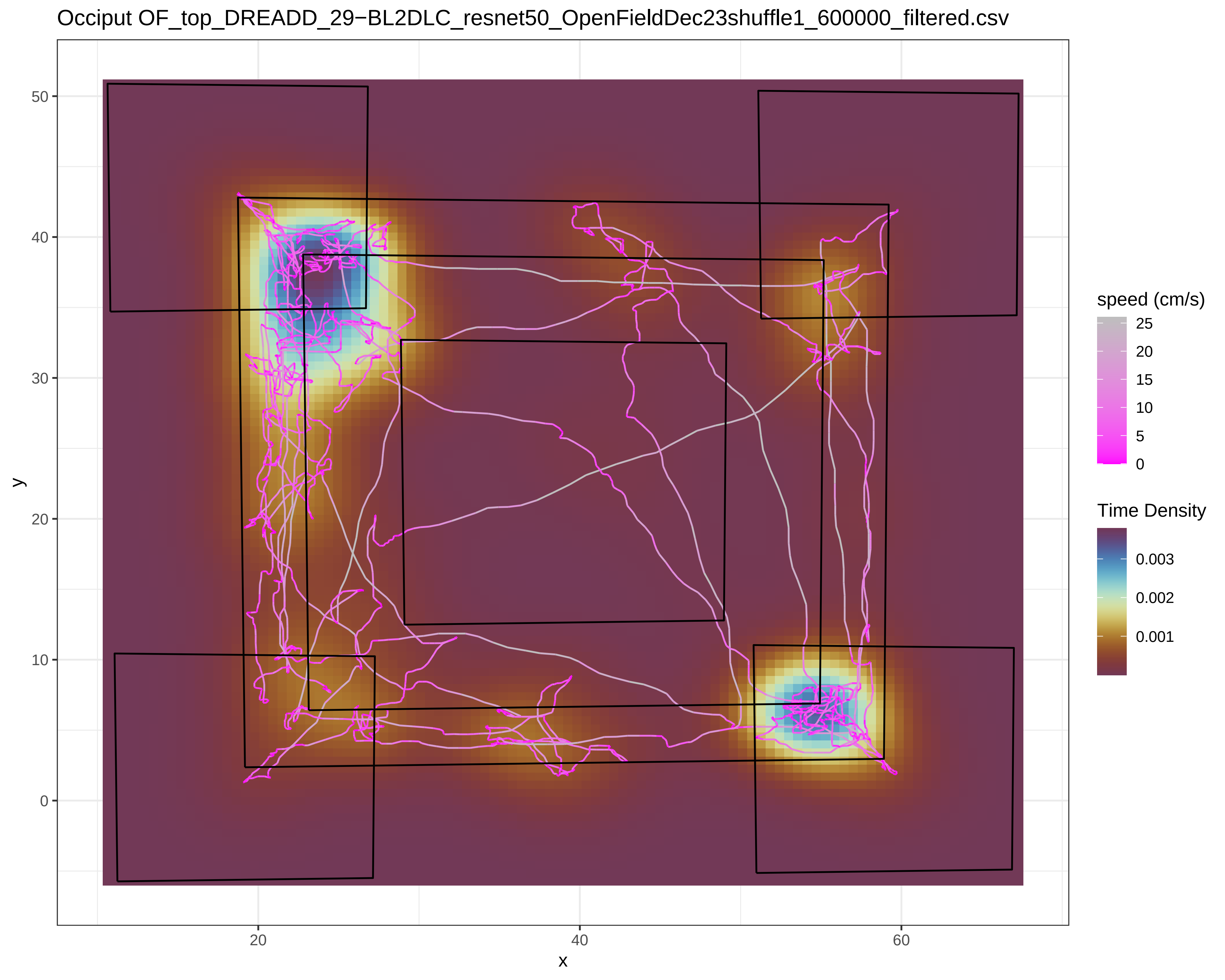
Occiput OF_top_DREADD_29−BL2DLC_resnet50_OpenFieldDec23shuffle1_600000_filtered.csv
y
x
50
40
30
20
10
0
20
40
60
speed (cm/s)
25
20
15
10
5
0
Time Density
0.003
0.002
0.001

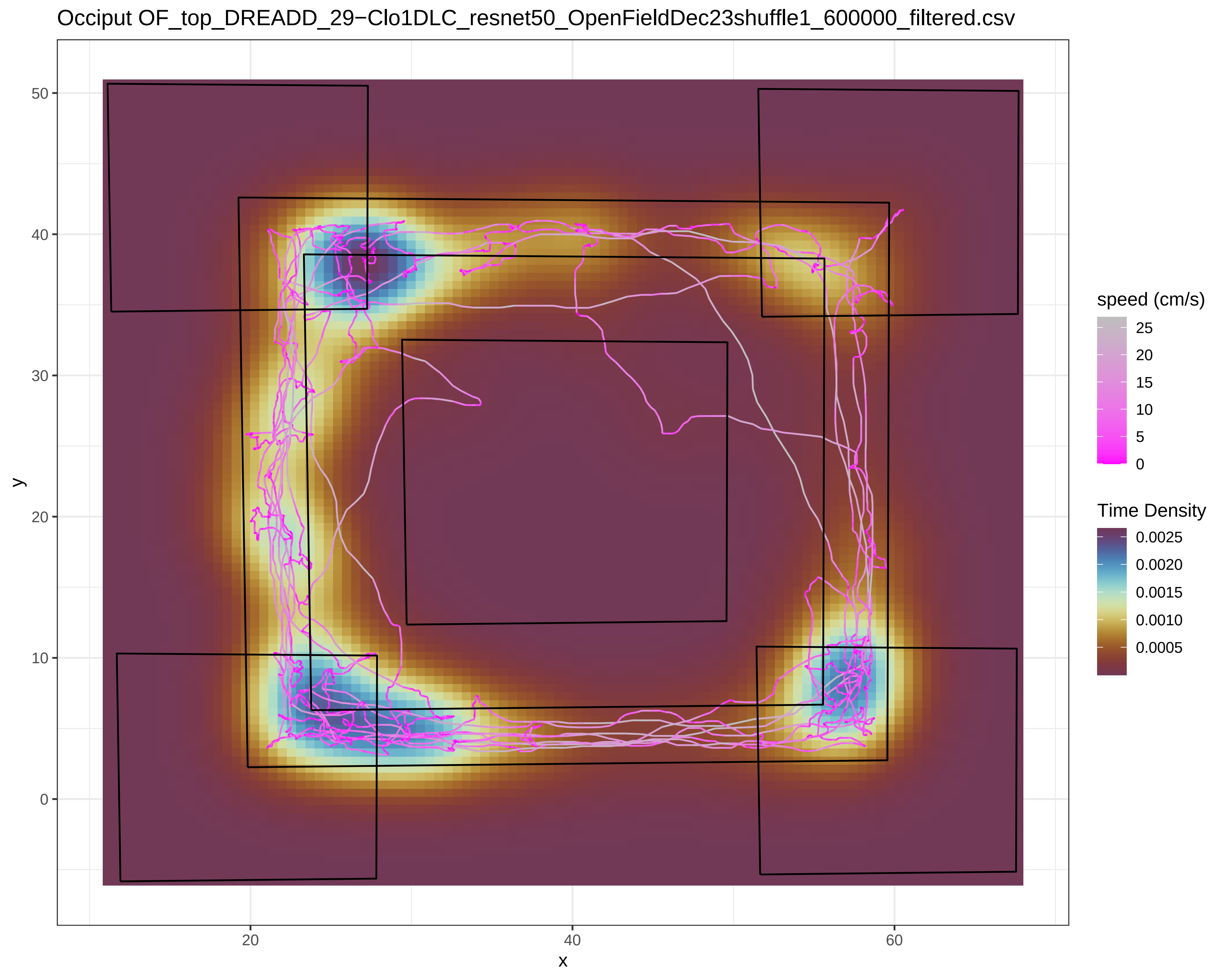

Occiput OF_top_DREADD_29−Clo1DLC_resnet50_OpenFieldDec23shuffle1_600000_filtered.csv
y
x
0
10
20
30
40
50
20
40
60
speed (cm/s)
25
20
15
10
5
0
Time Density
0.0025
0.0020
0.0015
0.0010
0.0005

Occiput OF_top_DREADD_29−Clo2DLC_resnet50_OpenFieldDec23shuffle1_600000_filtered.csv

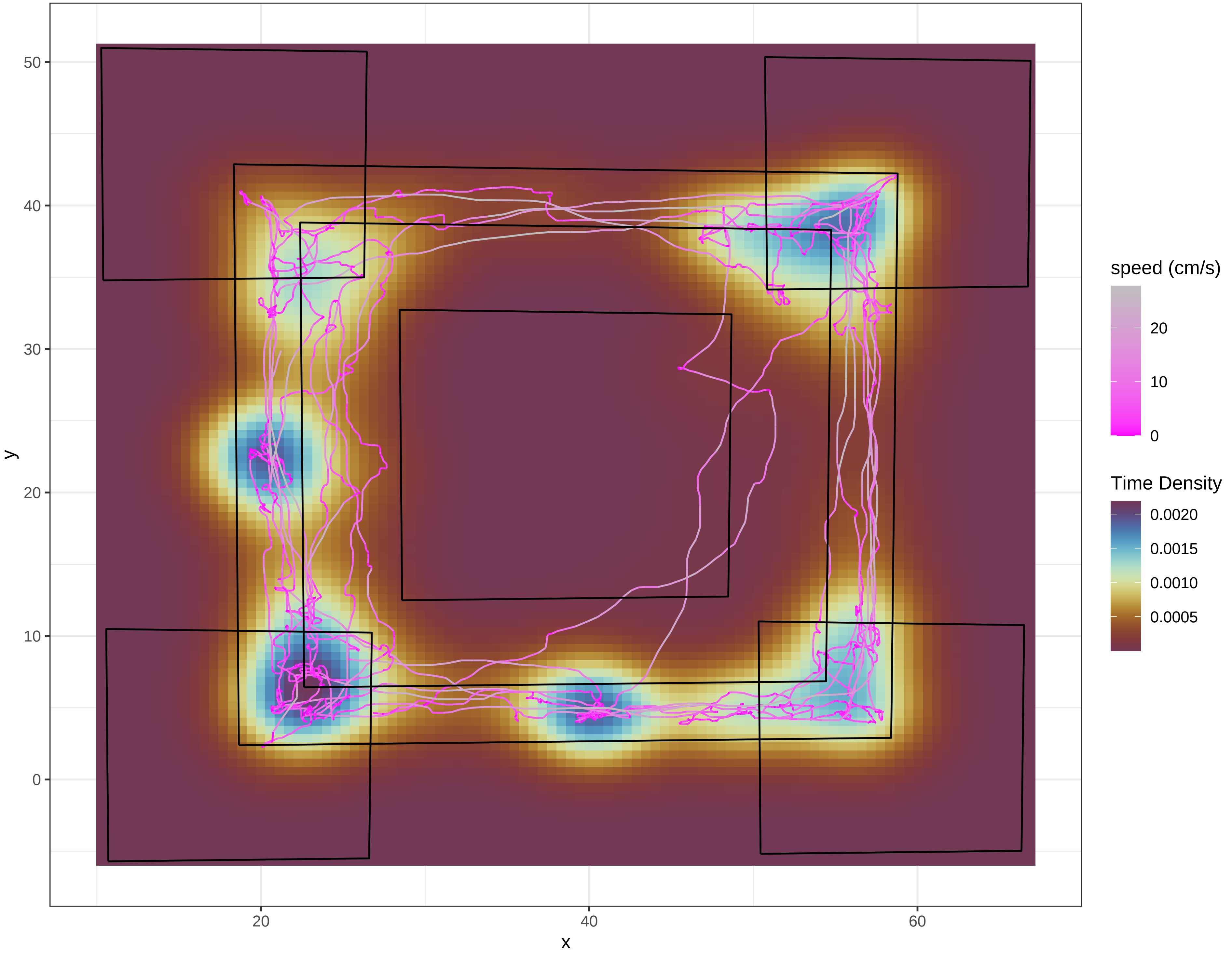

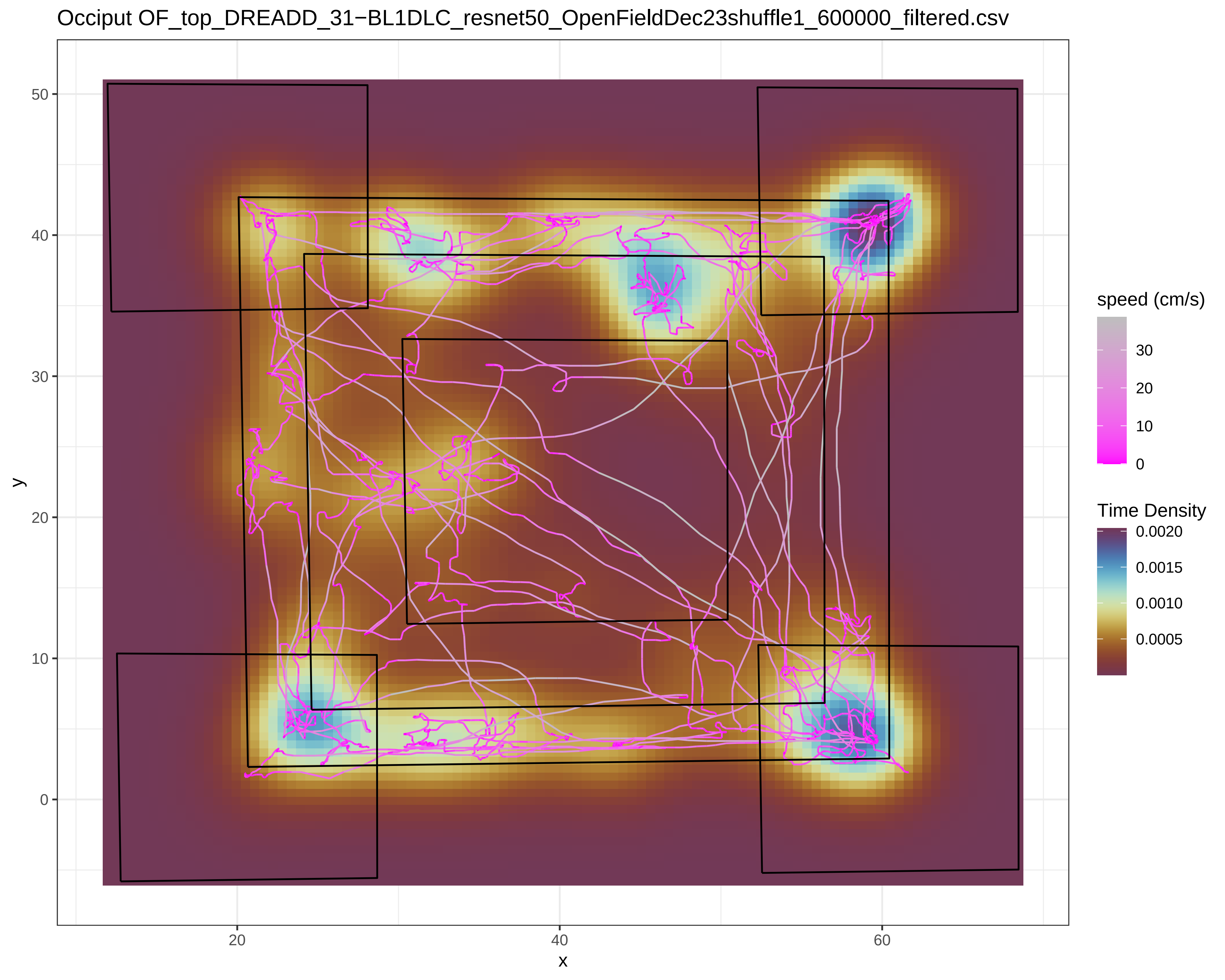
Occiput OF_top_DREADD_31−BL1DLC_resnet50_OpenFieldDec23shuffle1_600000_filtered.csv
y
x
50
40
30
20
10
0
20
40
60
speed (cm/s)
30
20
10
0
Time Density
0.0020
0.0015
0.0010
0.0005

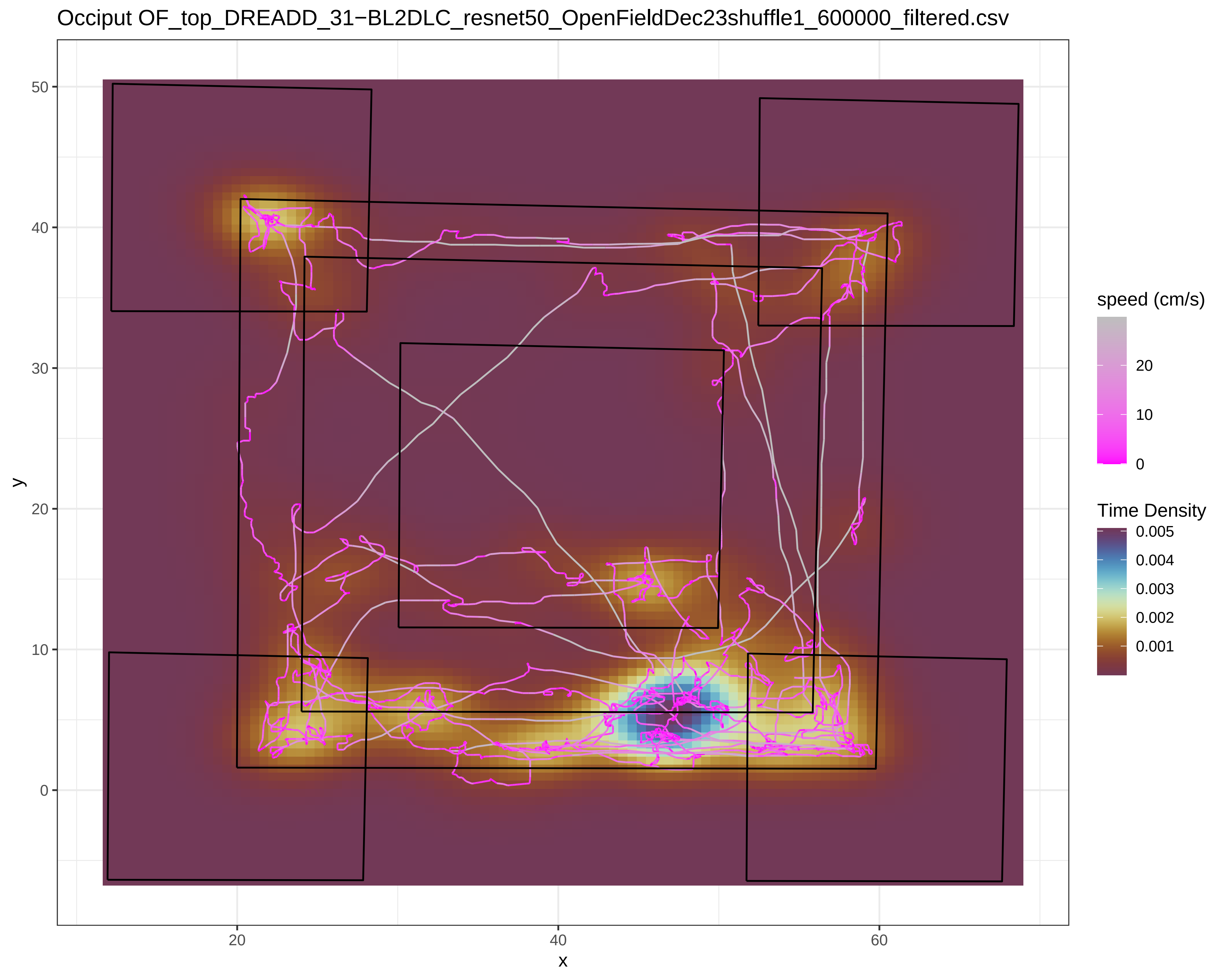
Occiput OF_top_DREADD_31−BL2DLC_resnet50_OpenFieldDec23shuffle1_600000_filtered.csv
y
x
50
40
30
20
10
0
20
40
60
speed (cm/s)
20
10
0
Time Density
0.005
0.004
0.003
0.002
0.001

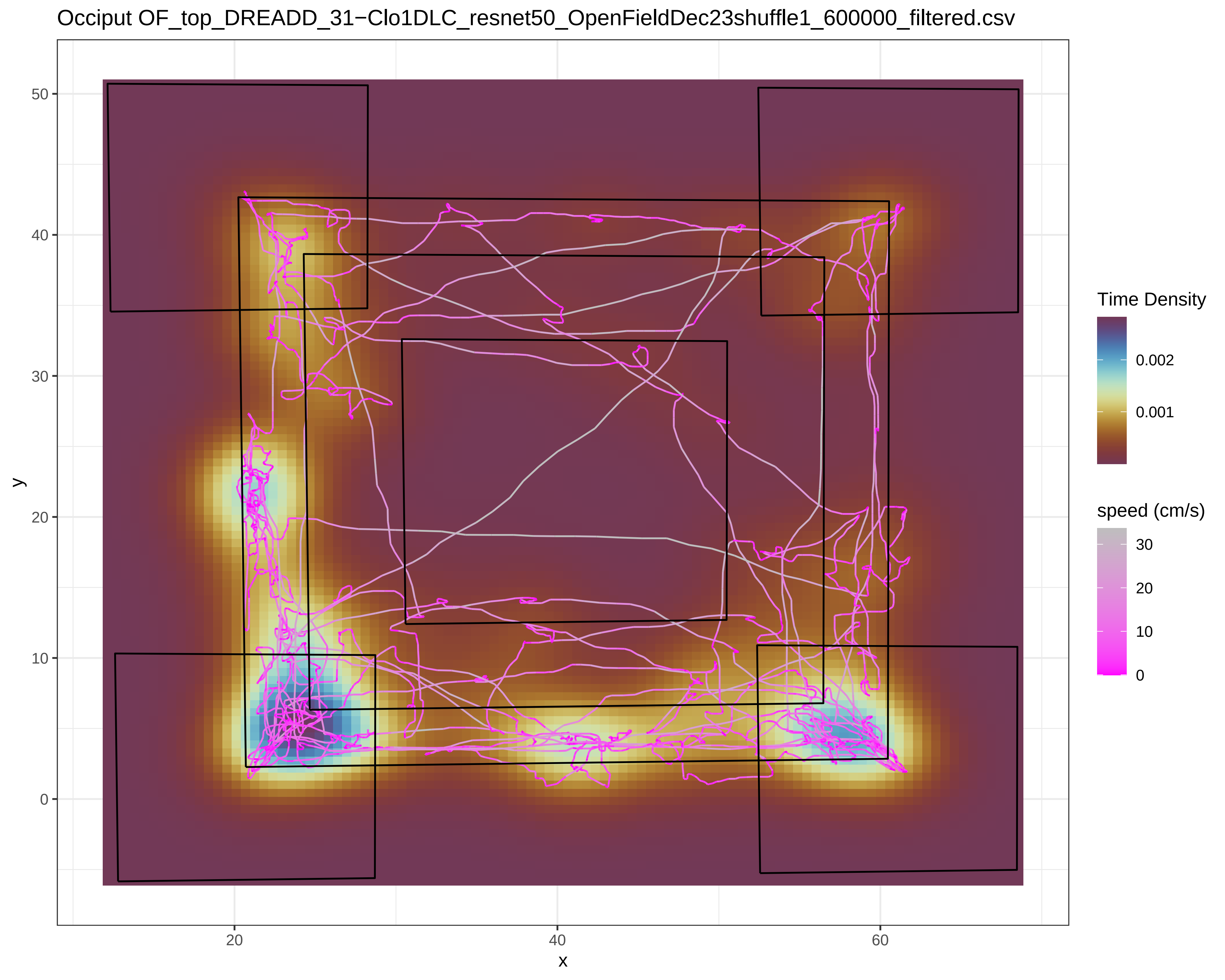

Occiput OF_top_DREADD_31−Clo1DLC_resnet50_OpenFieldDec23shuffle1_600000_filtered.csv
y
x
50
40
30
20
10
0
20
40
60
Time Density
0.002
0.001
speed (cm/s)
30
20
10
0

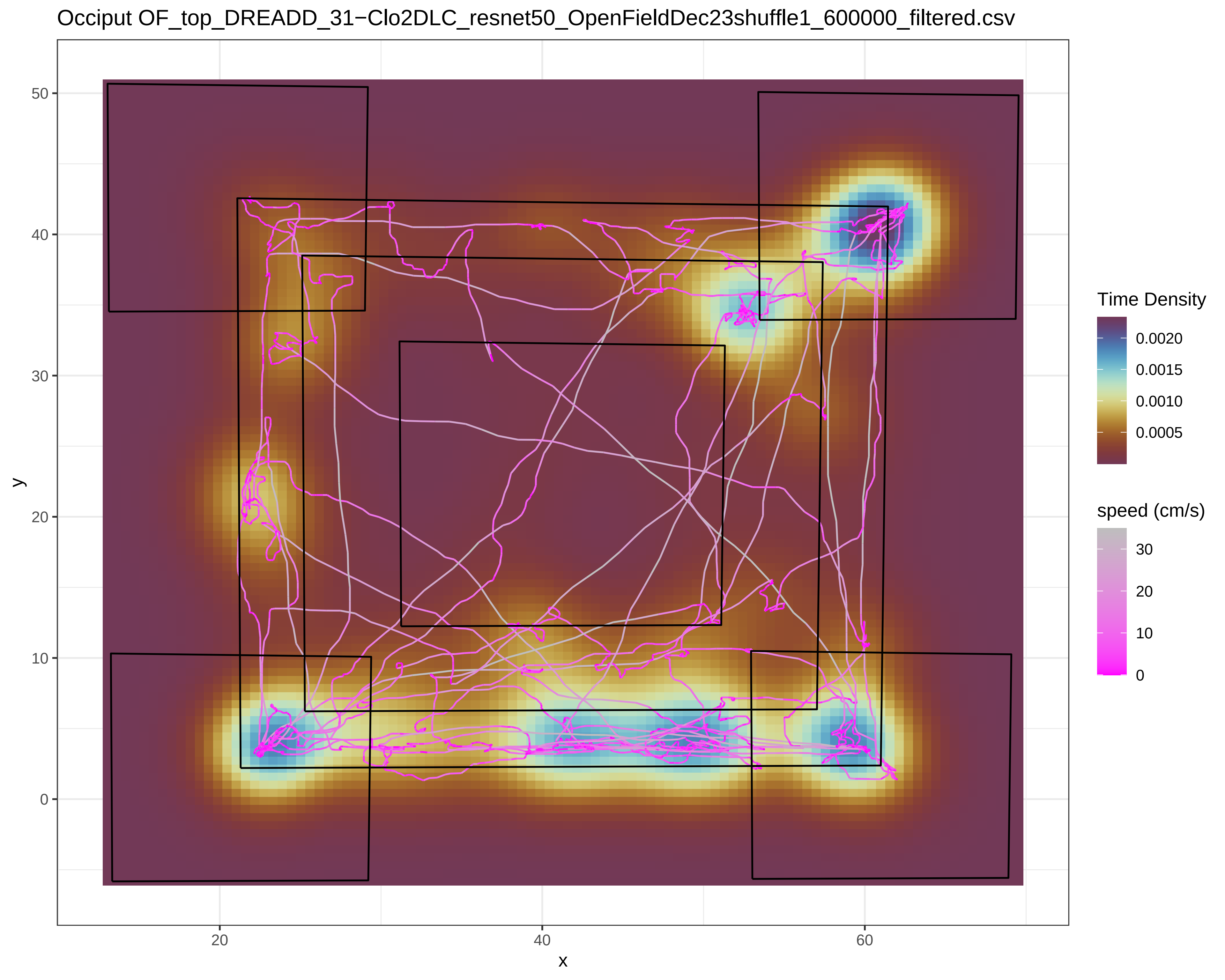

Occiput OF_top_DREADD_31−Clo2DLC_resnet50_OpenFieldDec23shuffle1_600000_filtered.csv
y
x
50
40
30
20
10
0
20
40
60
Time Density
0.0020
0.0015
0.0010
0.0005
speed (cm/s)
30
20
10
0

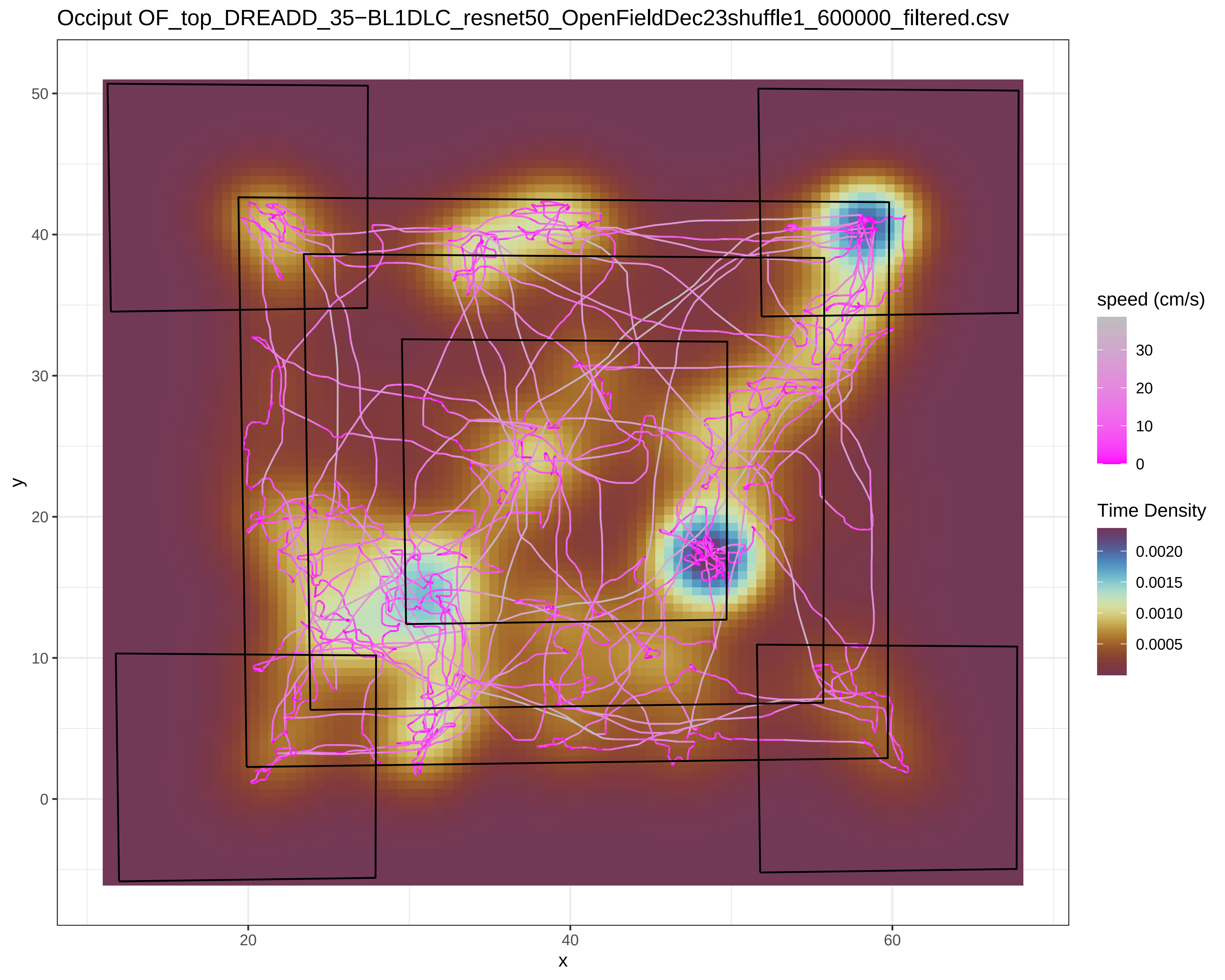
Occiput OF_top_DREADD_35−BL1DLC_resnet50_OpenFieldDec23shuffle1_600000_filtered.csv
speed (cm/s)
30
20
10
0
Time Density
0.0020
0.0015
0.0010
0.0005
y
x
50
40
30
20
10
0
20
40
60

Occiput OF_top_DREADD_35−BL2DLC_resnet50_OpenFieldDec23shuffle1_600000_filtered.csv

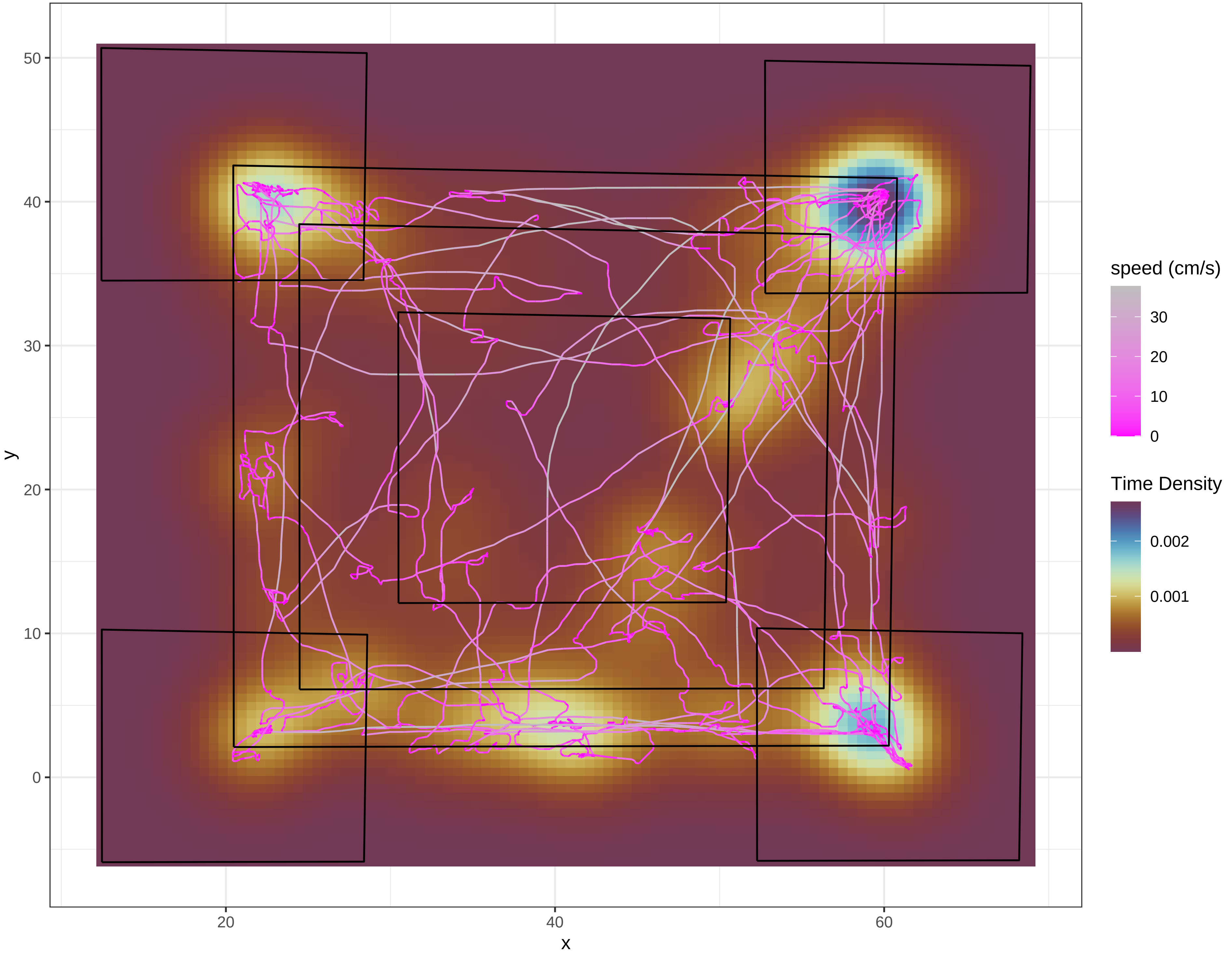

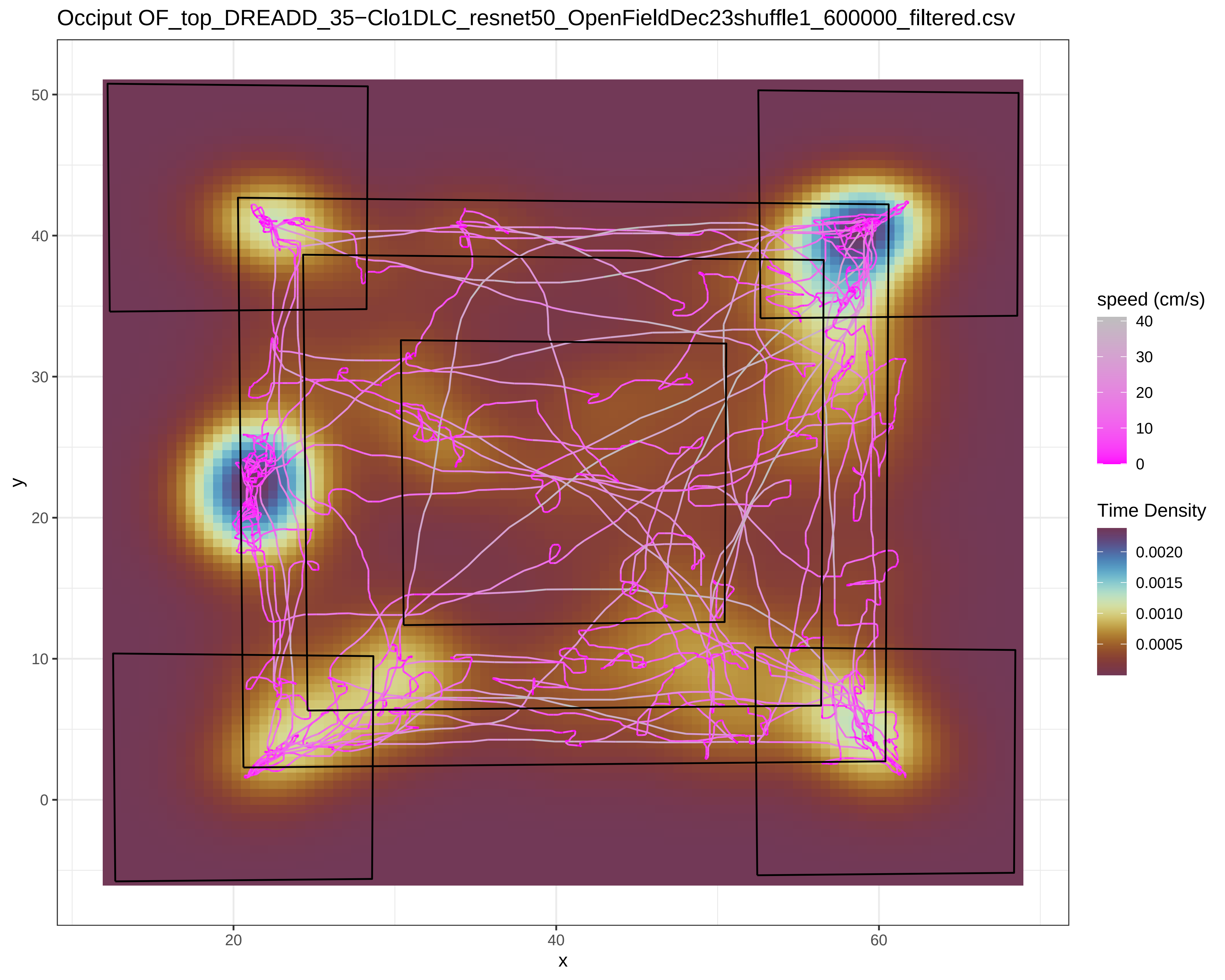
Occiput OF_top_DREADD_35−Clo1DLC_resnet50_OpenFieldDec23shuffle1_600000_filtered.csv
y
50
40
30
20
10
0
x
20
40
60
speed (cm/s)
40
30
20
10
0
Time Density
0.0020
0.0015
0.0010
0.0005

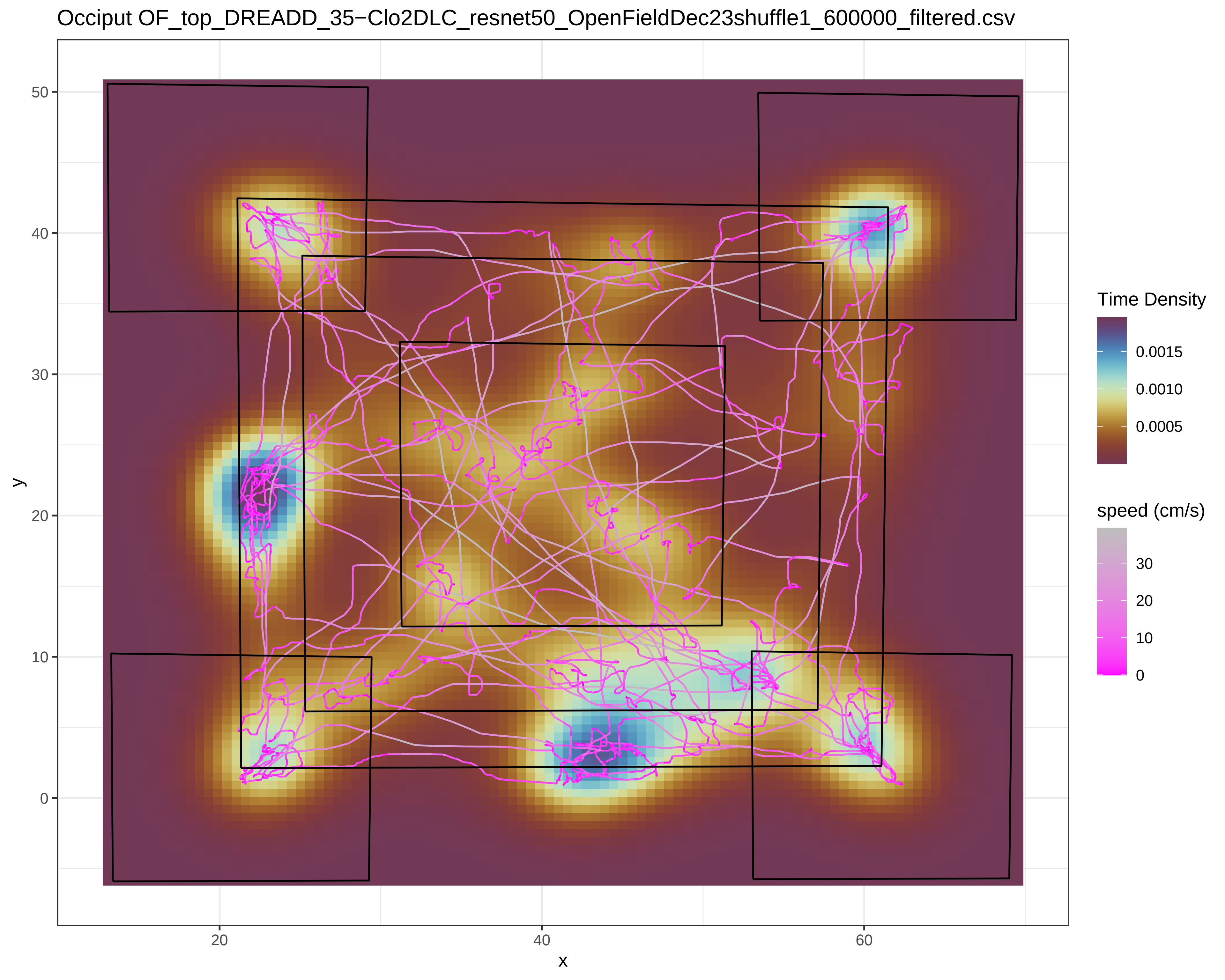
Occiput OF_top_DREADD_35−Clo2DLC_resnet50_OpenFieldDec23shuffle1_600000_filtered.csv
Time Density
0.0015
0.0010
0.0005
speed (cm/s)
30
20
10
0
y
50
40
30
20
10
0
x
20
40
60

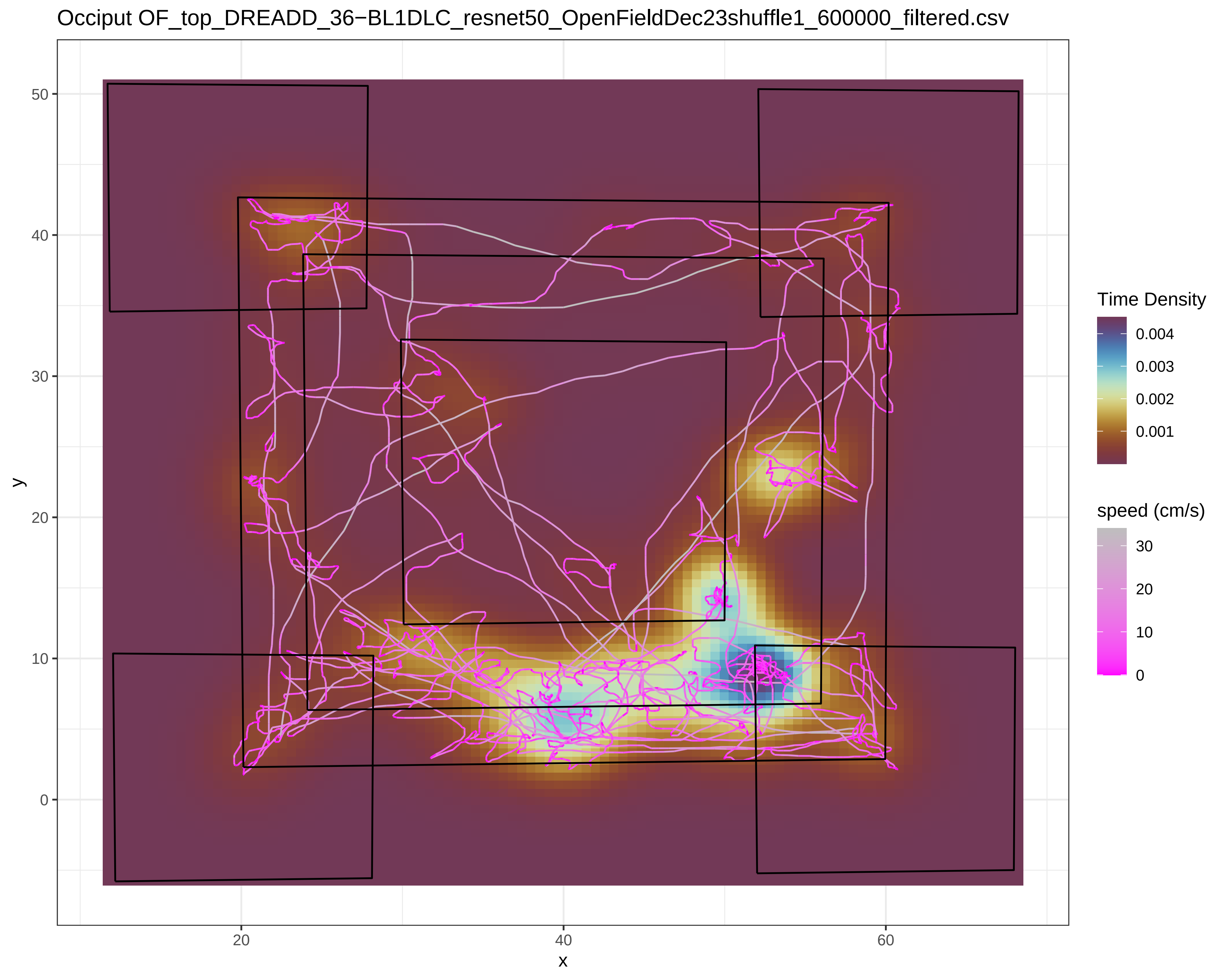
Occiput OF_top_DREADD_36−BL1DLC_resnet50_OpenFieldDec23shuffle1_600000_filtered.csv
y
x
0
10
20
30
40
50
20
40
60
Time Density
0.004
0.003
0.002
0.001
speed (cm/s)
30
20
10
0

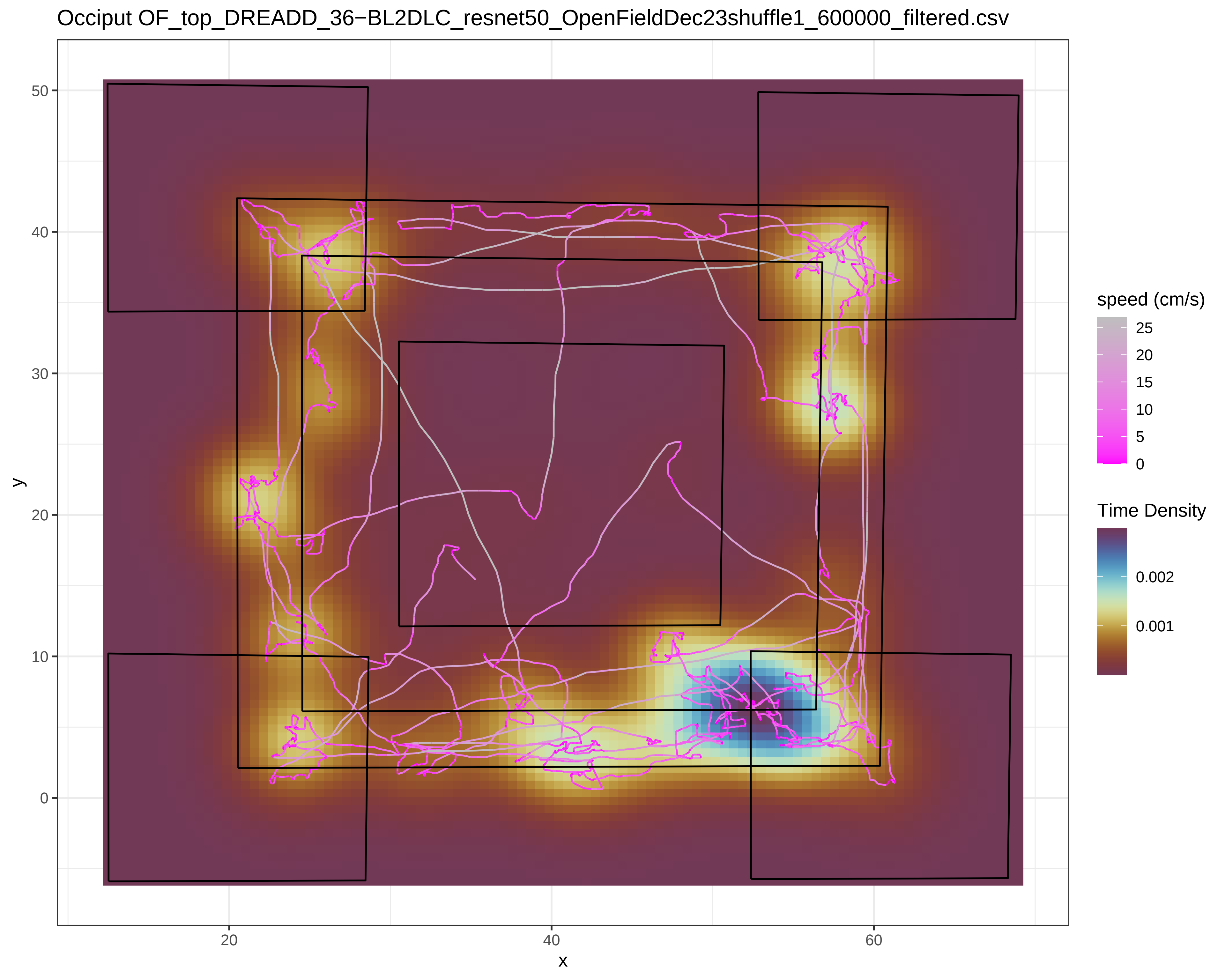
Occiput OF_top_DREADD_36−BL2DLC_resnet50_OpenFieldDec23shuffle1_600000_filtered.csv
y
x
50
40
30
20
10
0
20
40
60
speed (cm/s)
25
20
15
10
5
0
Time Density
0.002
0.001

Occiput OF_top_DREADD_36−Clo1DLC_resnet50_OpenFieldDec23shuffle1_600000_filtered.csv

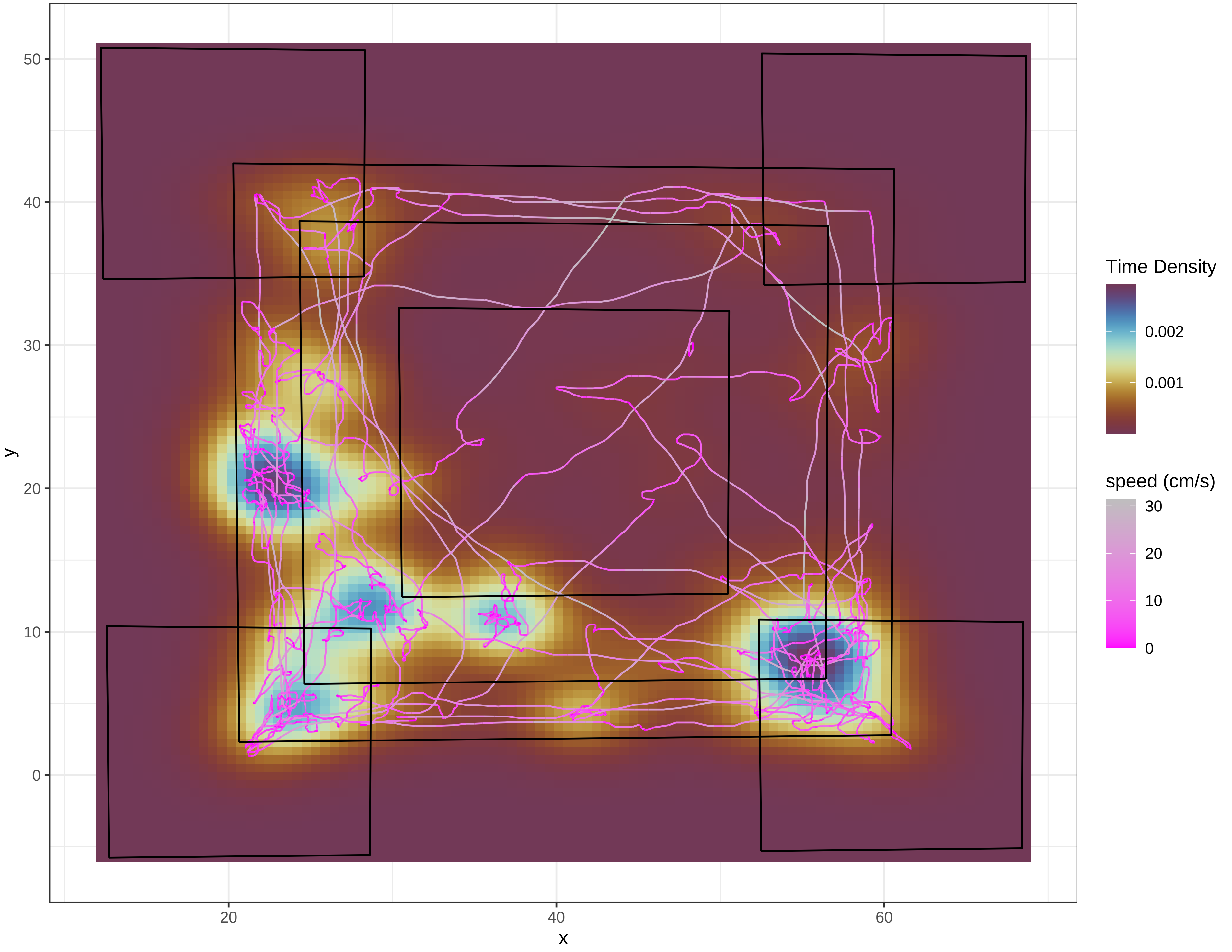

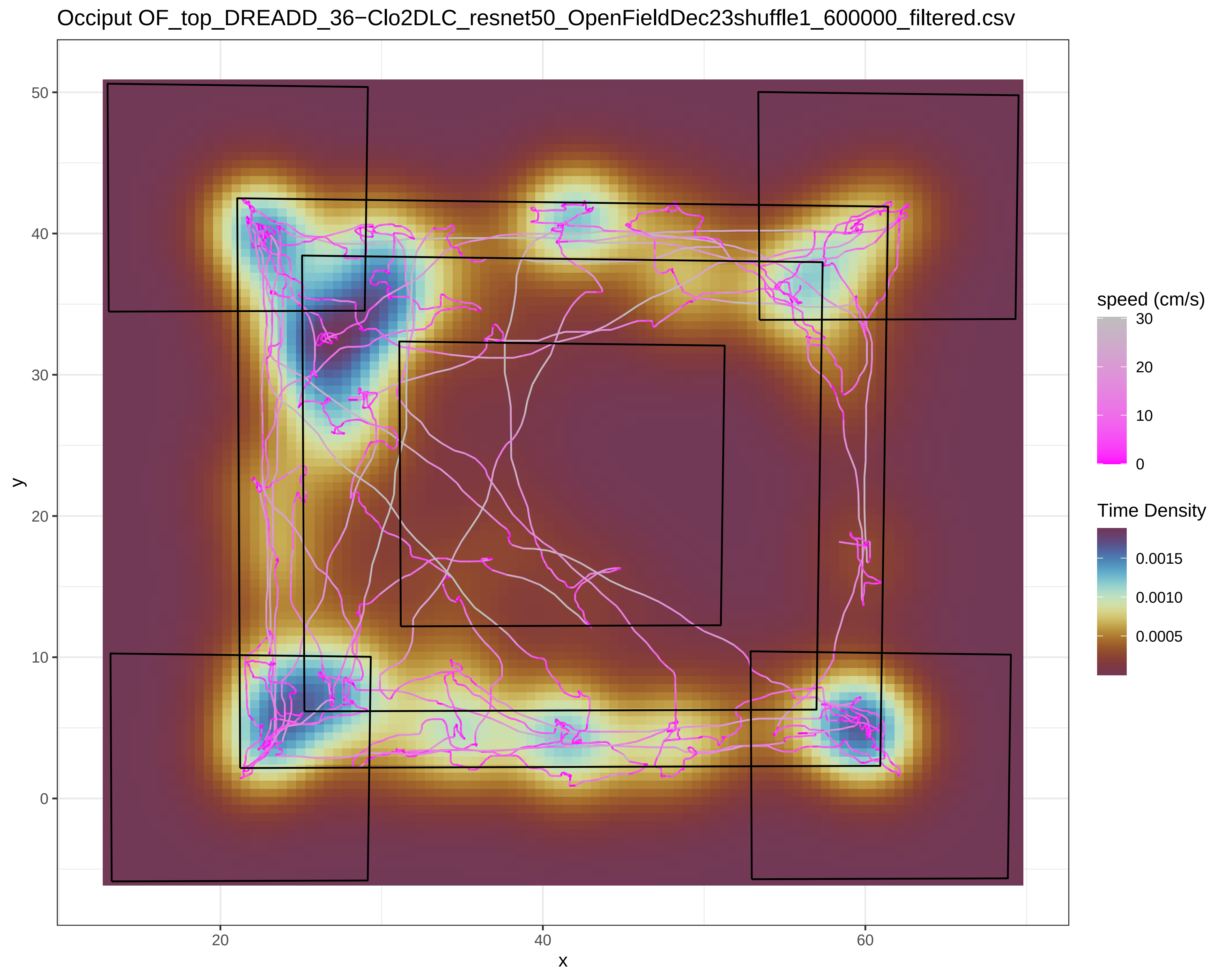

Occiput OF_top_DREADD_36−Clo2DLC_resnet50_OpenFieldDec23shuffle1_600000_filtered.csv
y
x
50
40
30
20
10
0
20
40
60
speed (cm/s)
30
20
10
0
Time Density
0.0015
0.0010
0.0005

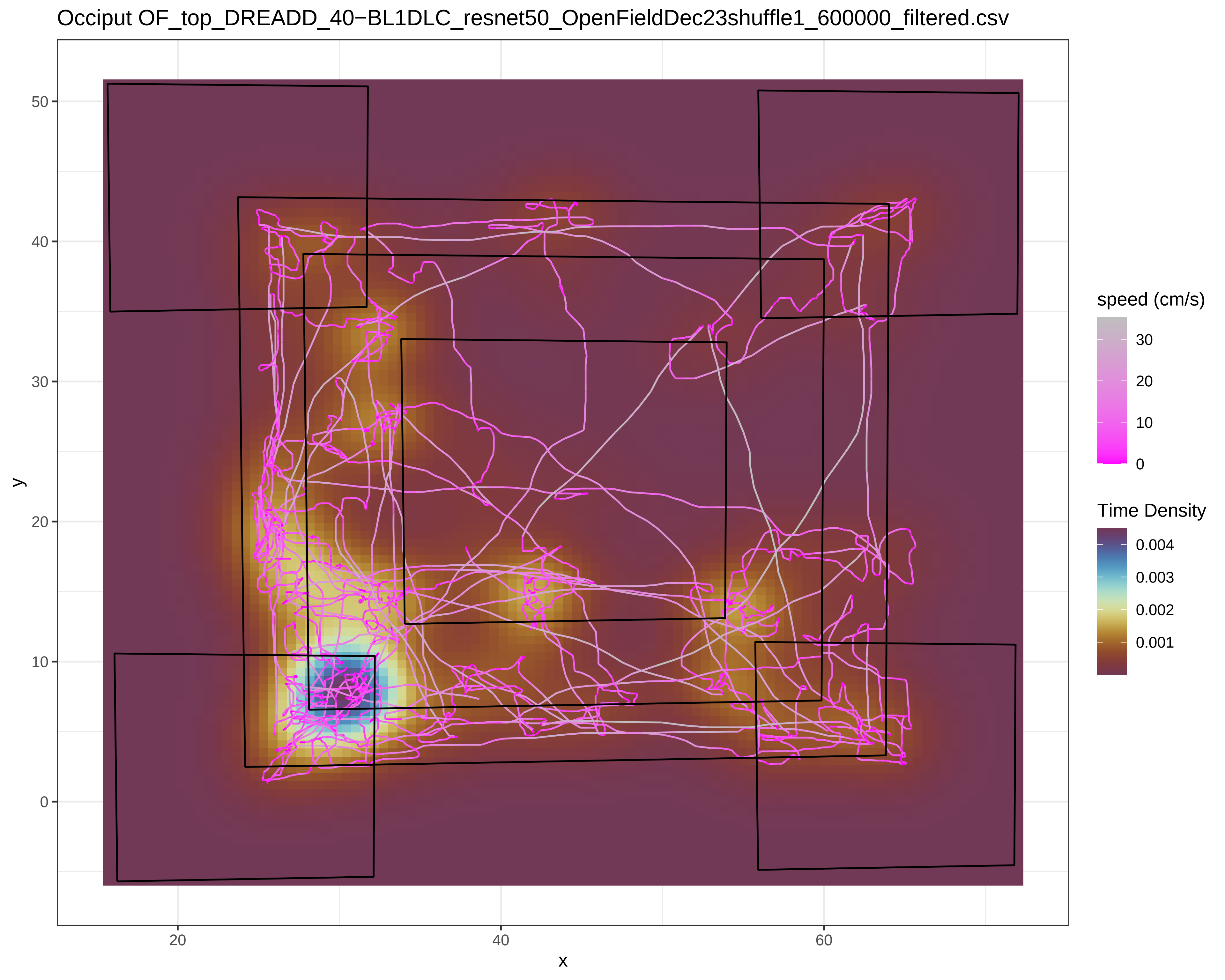
Occiput OF_top_DREADD_40−BL1DLC_resnet50_OpenFieldDec23shuffle1_600000_filtered.csv
y
x
50
40
30
20
10
0
20
40
60
speed (cm/s)
30
20
10
0
Time Density
0.004
0.003
0.002
0.001

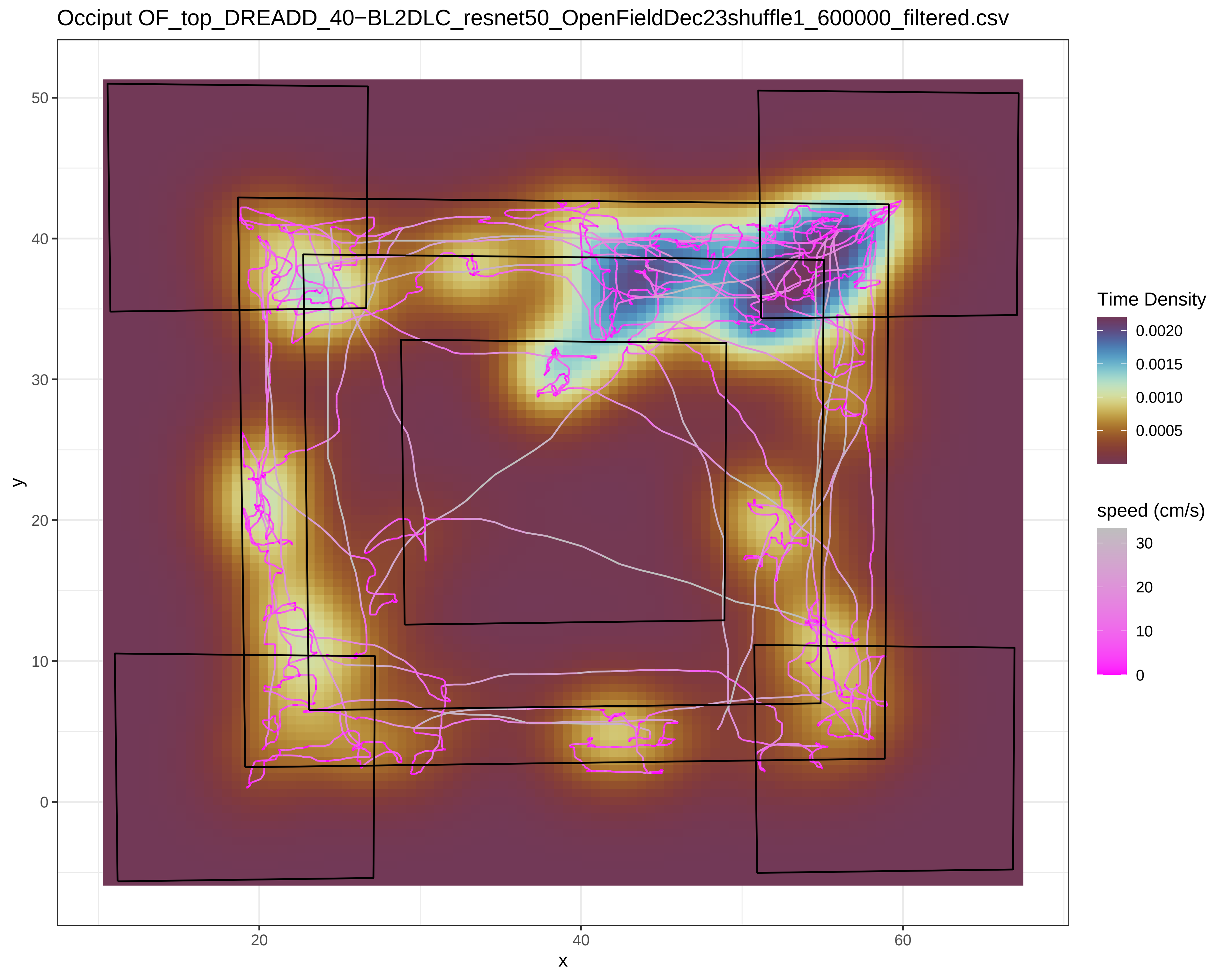
Occiput OF_top_DREADD_40−BL2DLC_resnet50_OpenFieldDec23shuffle1_600000_filtered.csv
Time Density
0.0020
0.0015
0.0010
0.0005
speed (cm/s)
30
20
10
0
y
50
40
30
20
10
0
x
20
40
60

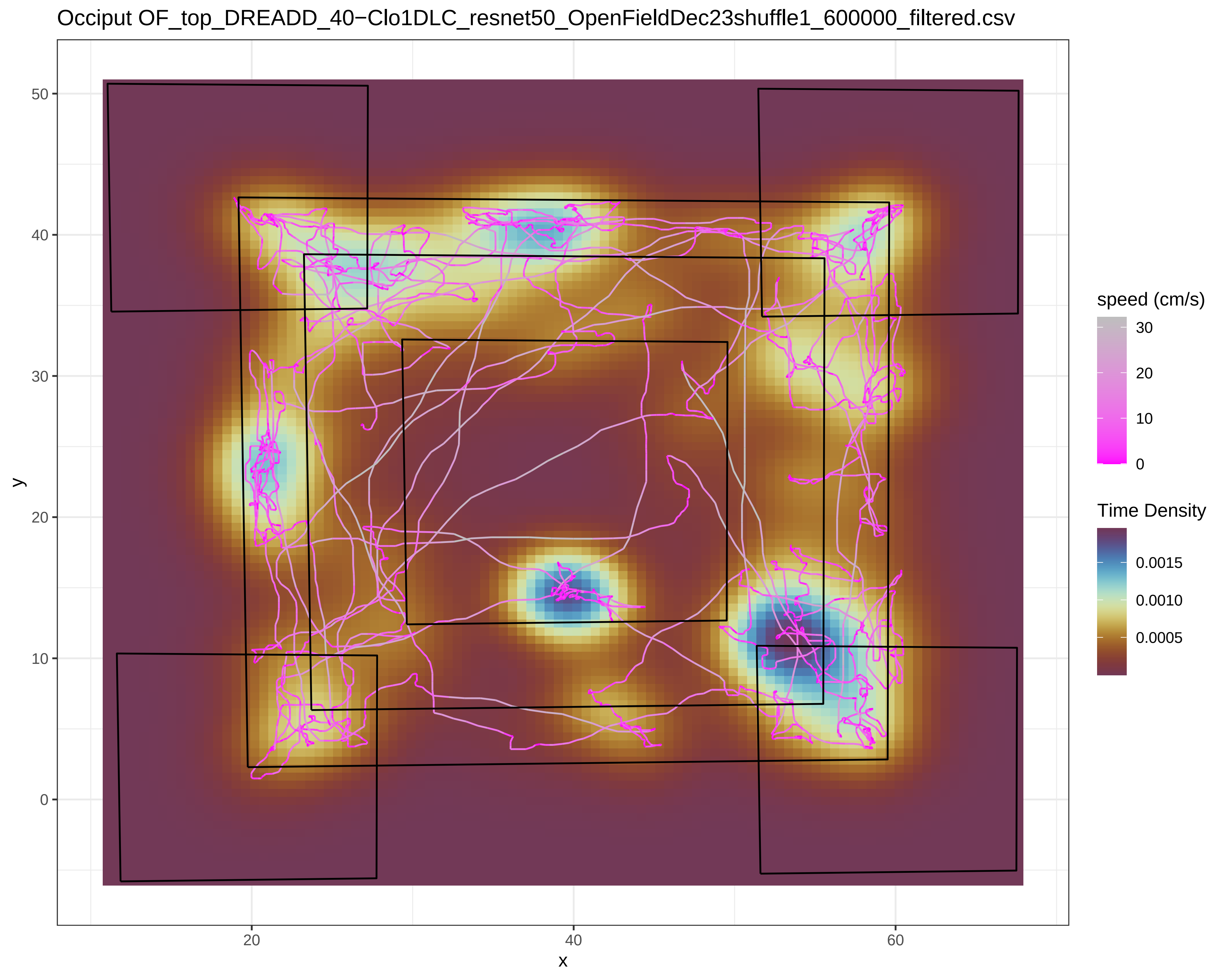
Occiput OF_top_DREADD_40−Clo1DLC_resnet50_OpenFieldDec23shuffle1_600000_filtered.csv
y
50
40
30
20
10
0
x
20
40
60
speed (cm/s)
30
20
10
0
Time Density
0.0015
0.0010
0.0005

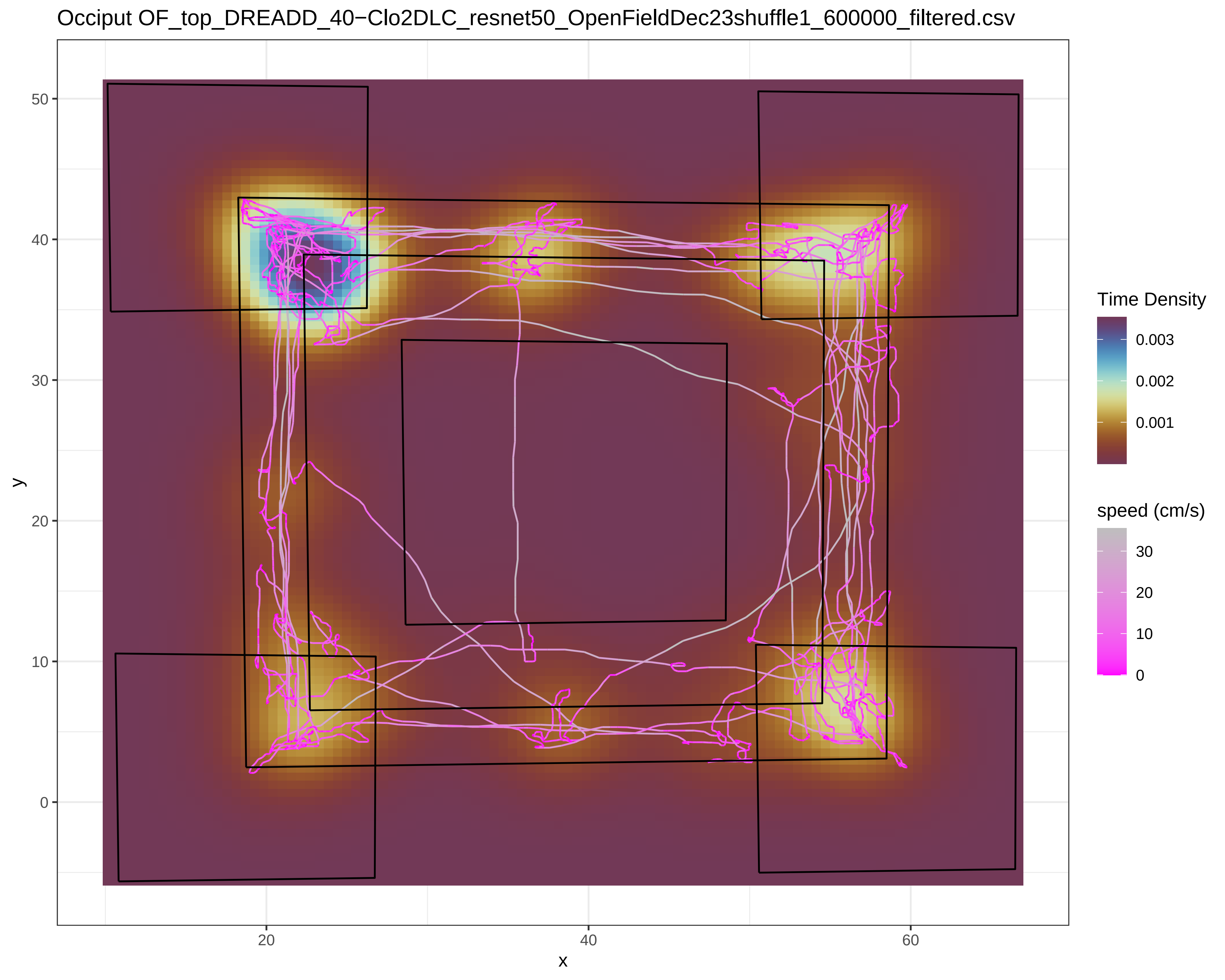
Occiput OF_top_DREADD_40−Clo2DLC_resnet50_OpenFieldDec23shuffle1_600000_filtered.csv
Time Density
0.003
0.002
0.001
speed (cm/s)
30
20
10
0
y
x
50
40
30
20
10
0
20
40
60

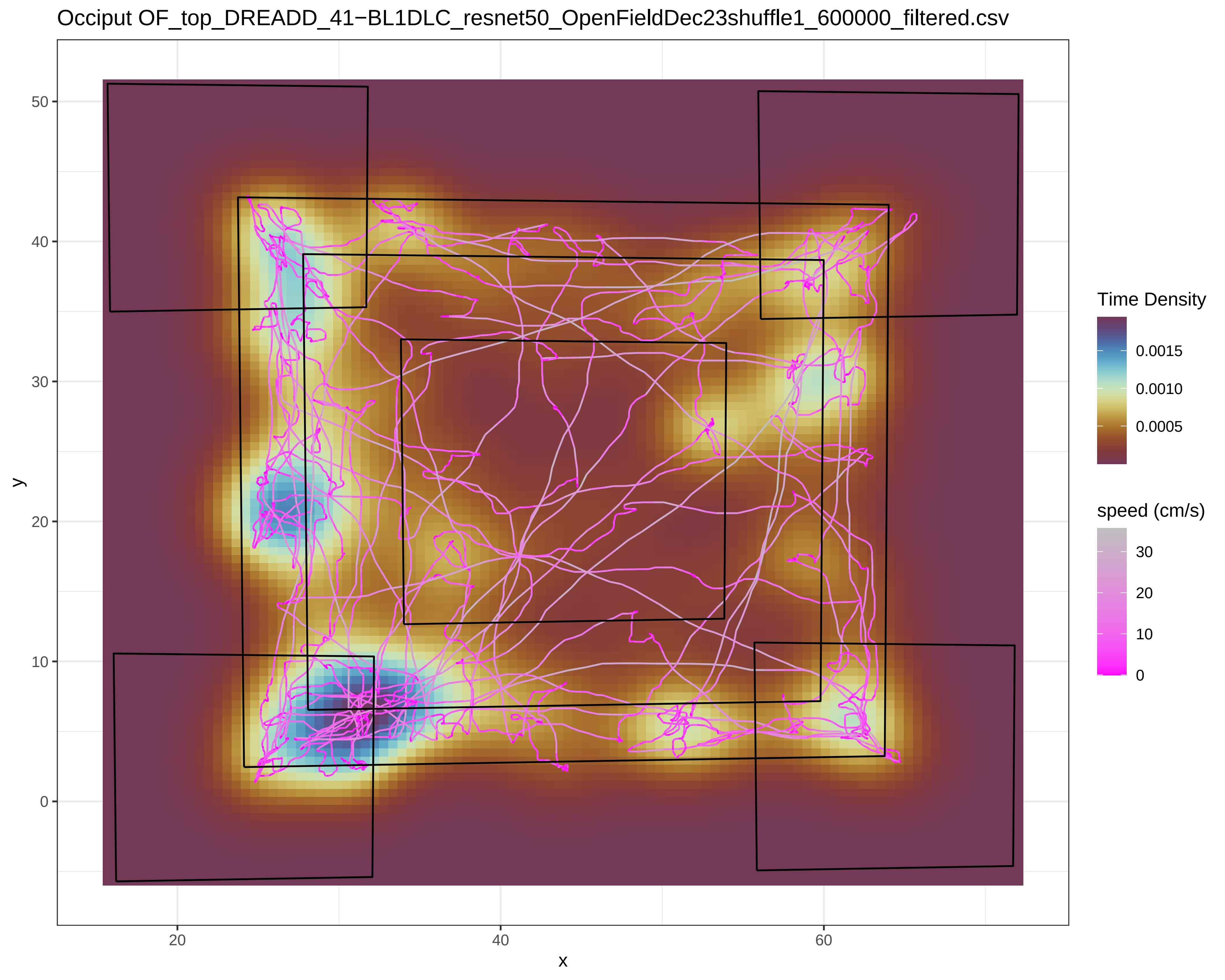
Occiput OF_top_DREADD_41−BL1DLC_resnet50_OpenFieldDec23shuffle1_600000_filtered.csv
y
x
50
40
30
20
10
0
20
40
60
Time Density
0.0015
0.0010
0.0005
speed (cm/s)
30
20
10
0

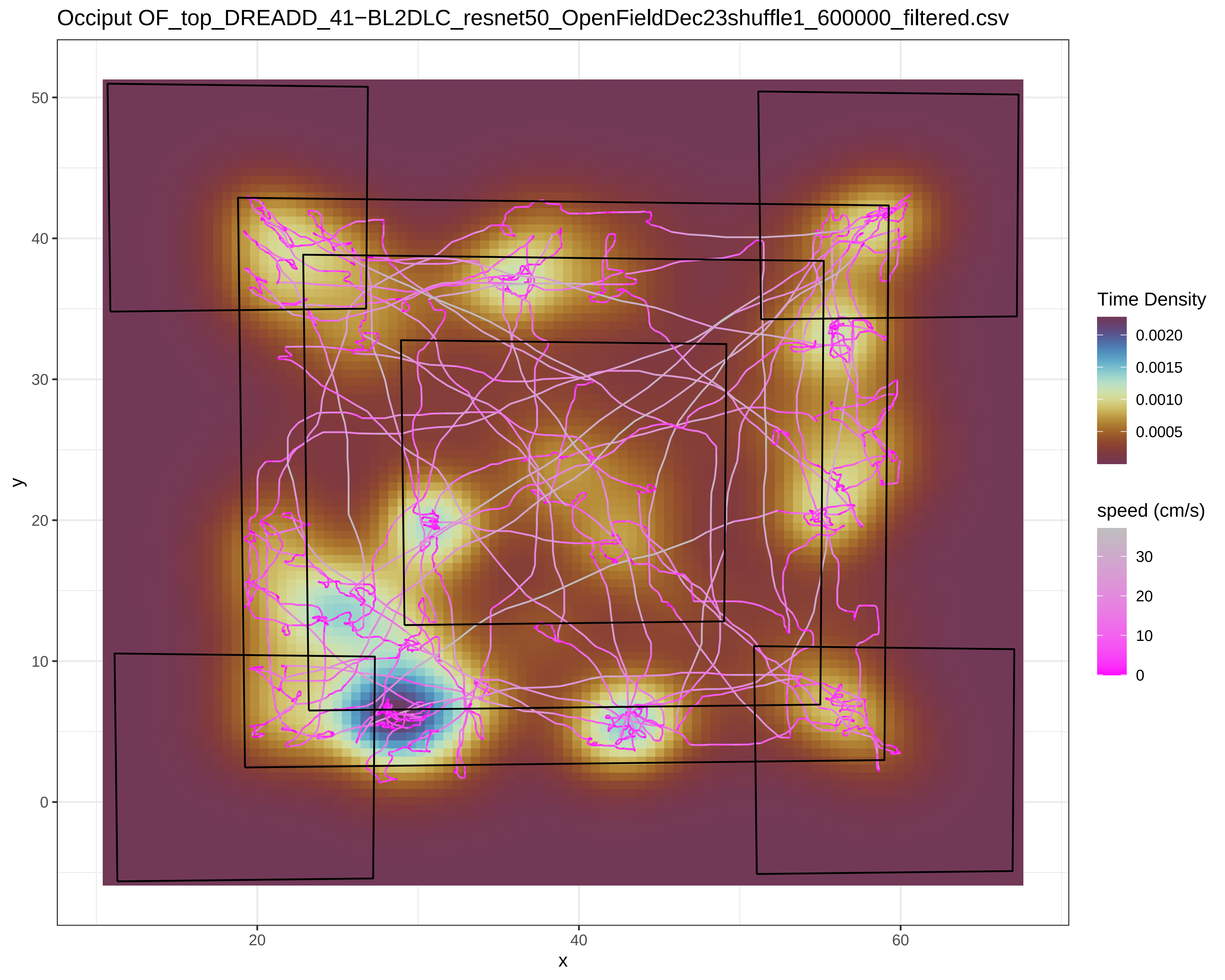

Occiput OF_top_DREADD_41−BL2DLC_resnet50_OpenFieldDec23shuffle1_600000_filtered.csv
y
50
40
30
20
10
0
x
20
40
60
Time Density
0.0020
0.0015
0.0010
0.0005
speed (cm/s)
30
20
10
0

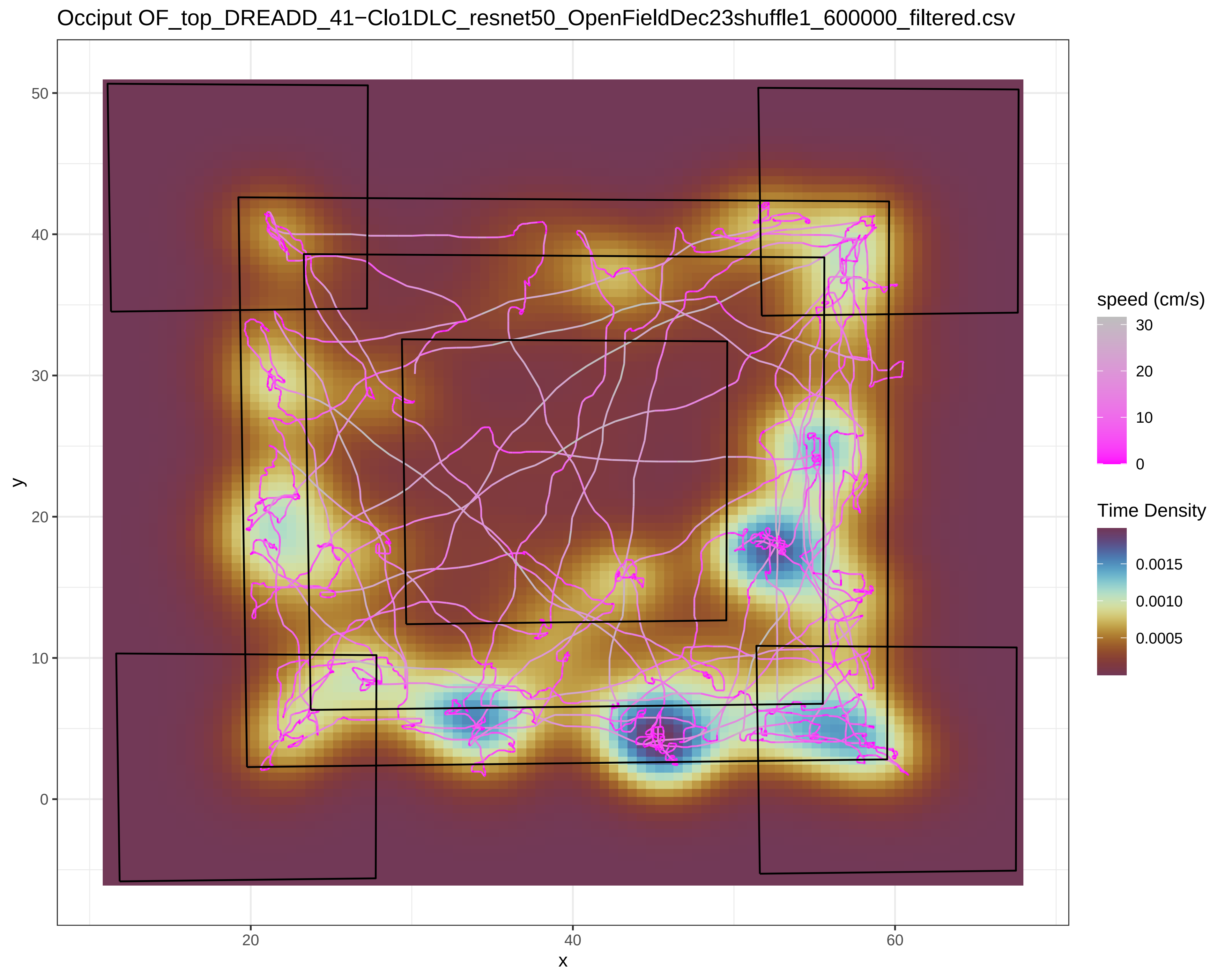

Occiput OF_top_DREADD_41−Clo1DLC_resnet50_OpenFieldDec23shuffle1_600000_filtered.csv
y
x
50
40
30
20
10
0
20
40
60
speed (cm/s)
30
20
10
0
Time Density
0.0015
0.0010
0.0005

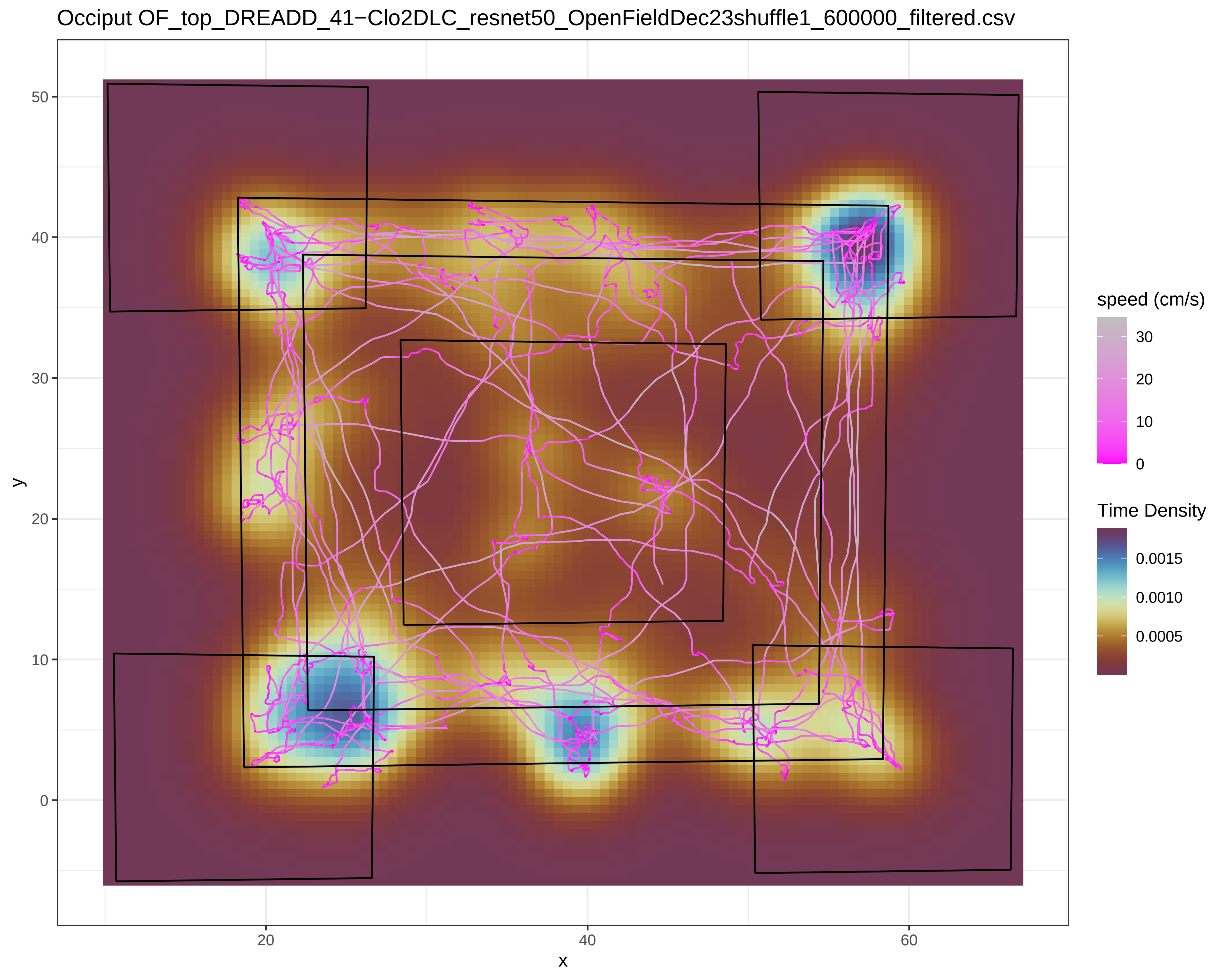
Occiput OF_top_DREADD_41−Clo2DLC_resnet50_OpenFieldDec23shuffle1_600000_filtered.csv
y
x
0
10
20
30
40
50
20
40
60
speed (cm/s)
30
20
10
0
Time Density
0.0015
0.0010
0.0005

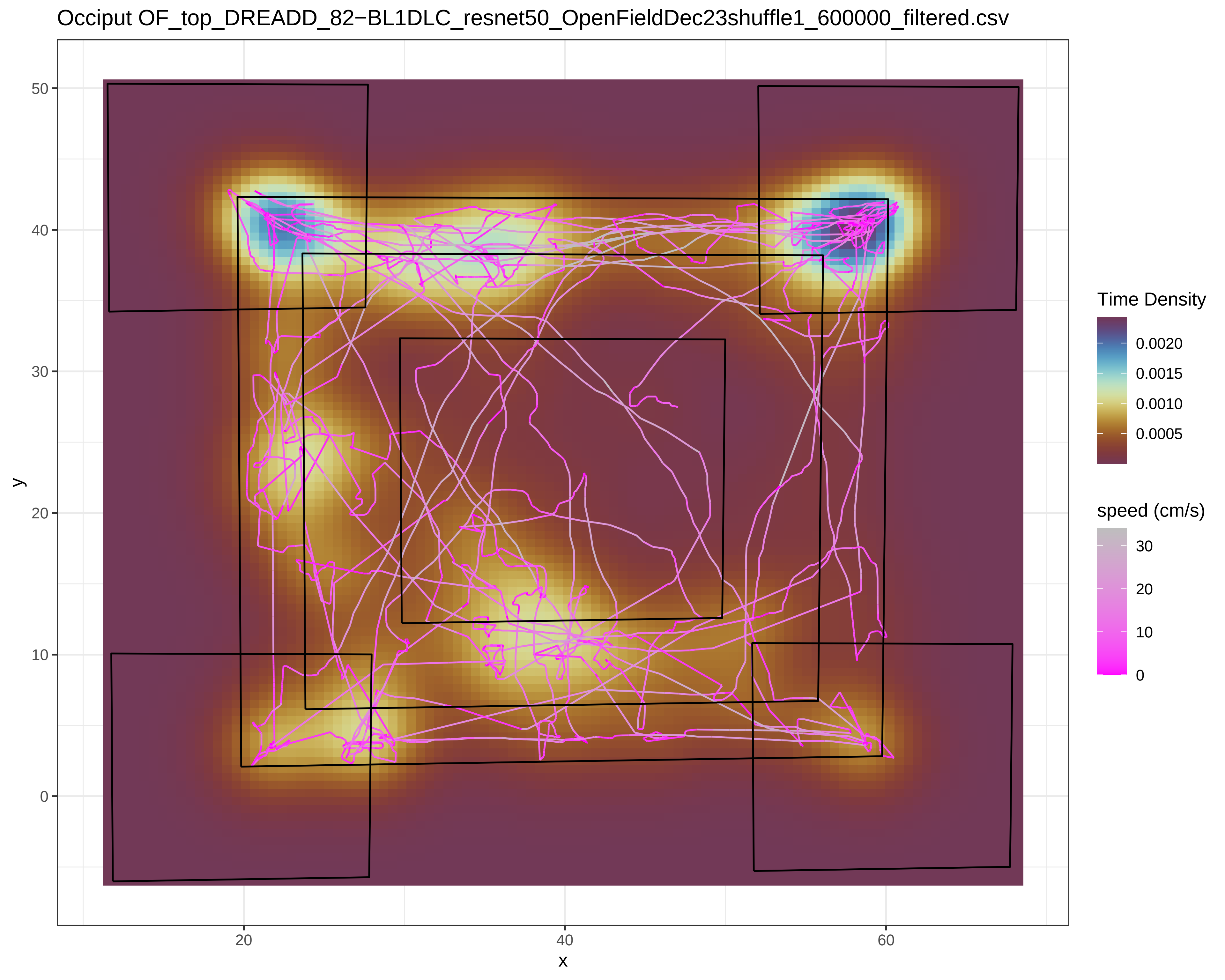
Occiput OF_top_DREADD_82−BL1DLC_resnet50_OpenFieldDec23shuffle1_600000_filtered.csv
y
x
50
40
30
20
10
0
20
40
60
Time Density
0.0020
0.0015
0.0010
0.0005
speed (cm/s)
30
20
10
0

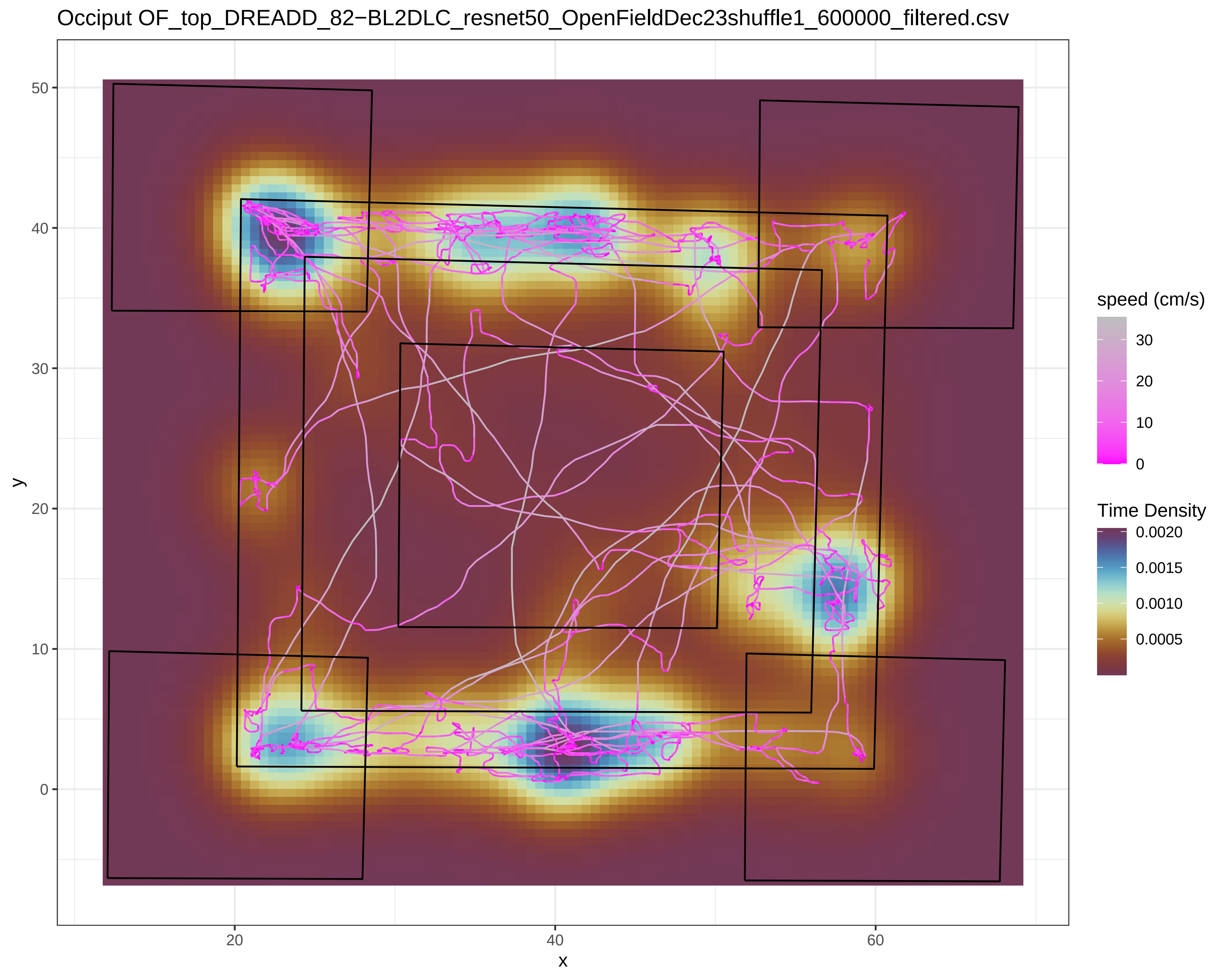

Occiput OF_top_DREADD_82−BL2DLC_resnet50_OpenFieldDec23shuffle1_600000_filtered.csv
y
50
40
30
20
10
0
x
20
40
60
speed (cm/s)
30
20
10
0
Time Density
0.0020
0.0015
0.0010
0.0005

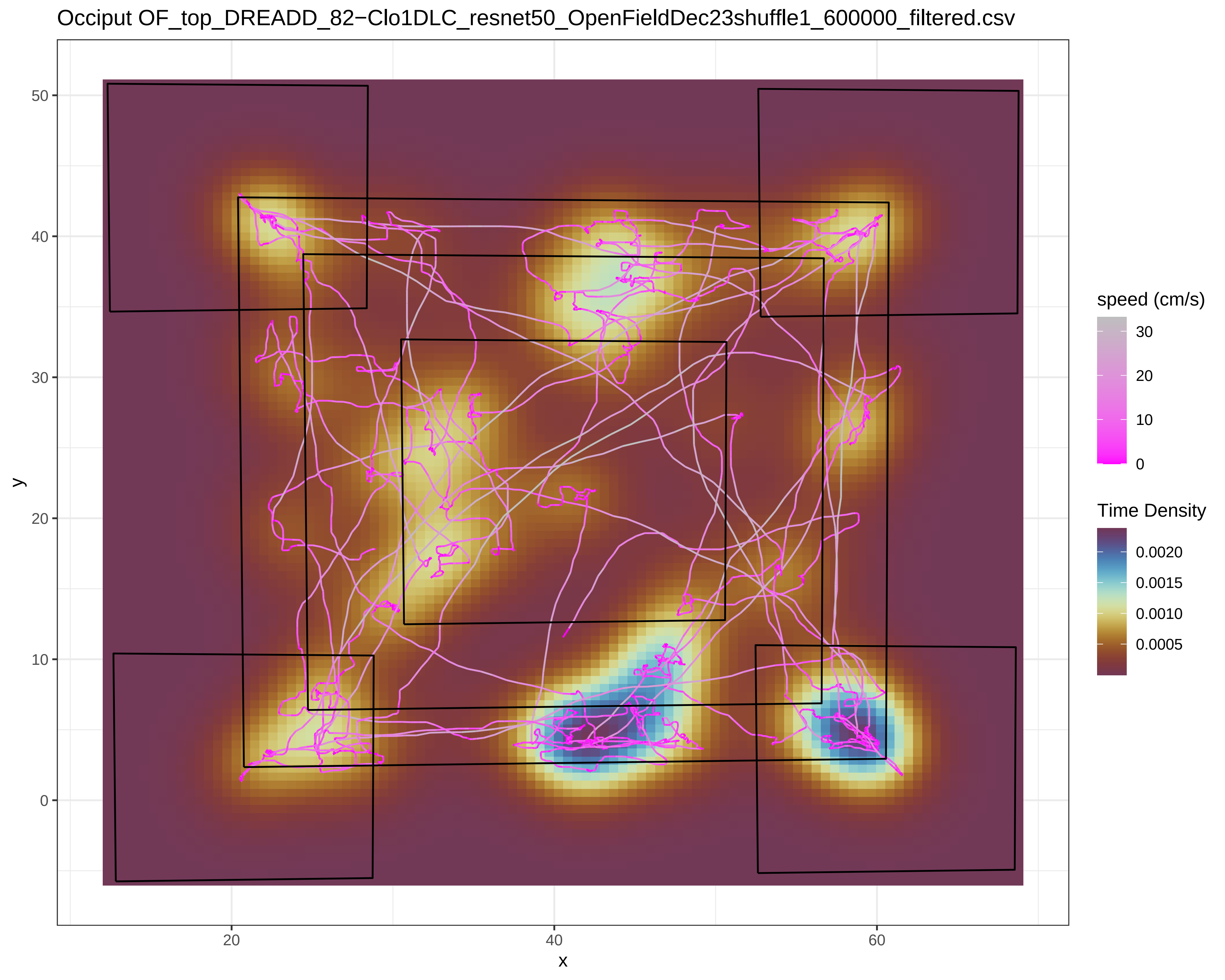
Occiput OF_top_DREADD_82−Clo1DLC_resnet50_OpenFieldDec23shuffle1_600000_filtered.csv
y
x
0
10
20
30
40
50
20
40
60
speed (cm/s)
30
20
10
0
Time Density
0.0020
0.0015
0.0010
0.0005

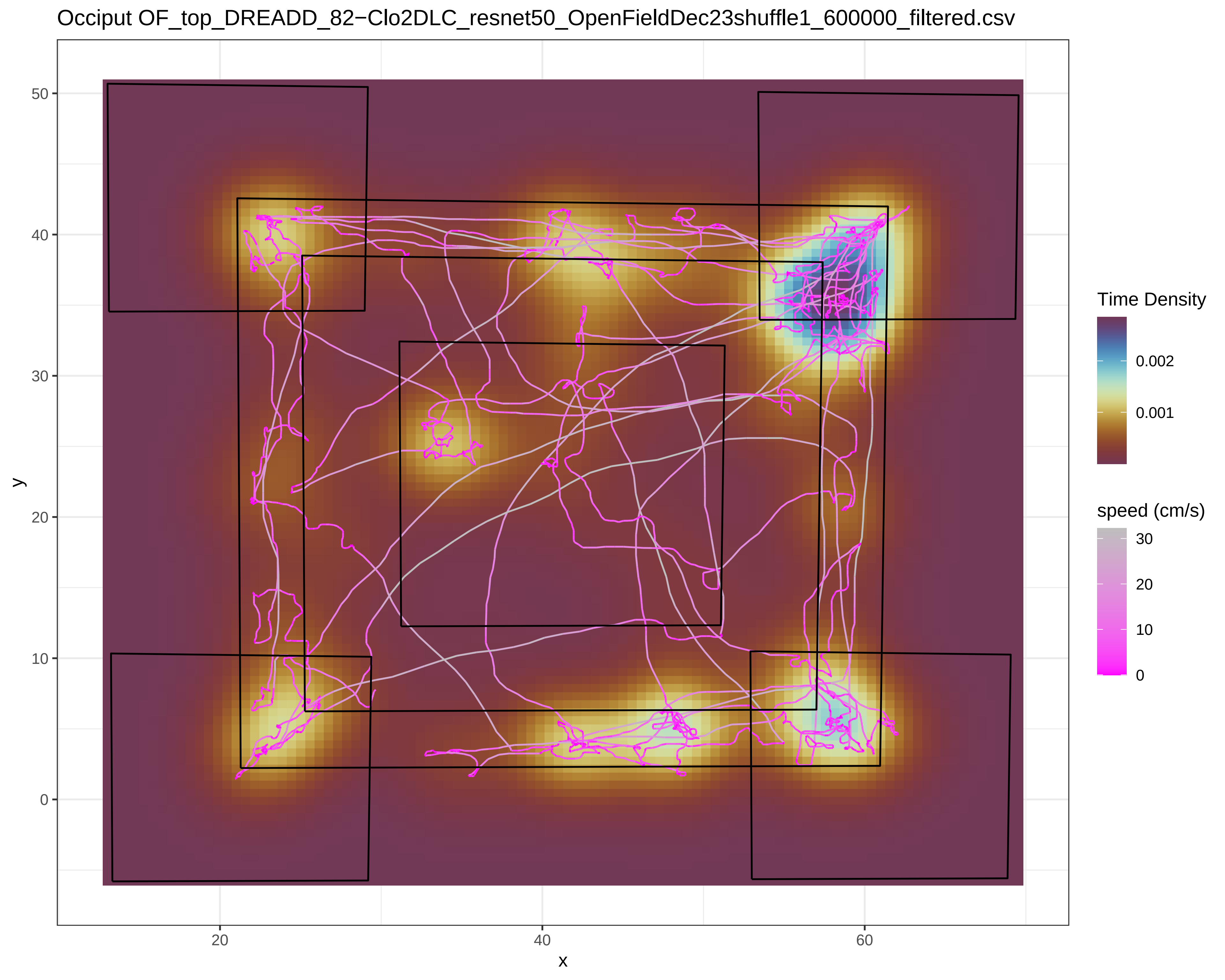
Occiput OF_top_DREADD_82−Clo2DLC_resnet50_OpenFieldDec23shuffle1_600000_filtered.csv
y
50
40
30
20
10
0
x
20
40
60
Time Density
0.002
0.001
speed (cm/s)
30
20
10
0

Occiput OF_top_DREADD_83−BL1DLC_resnet50_OpenFieldDec23shuffle1_600000_filtered.csv

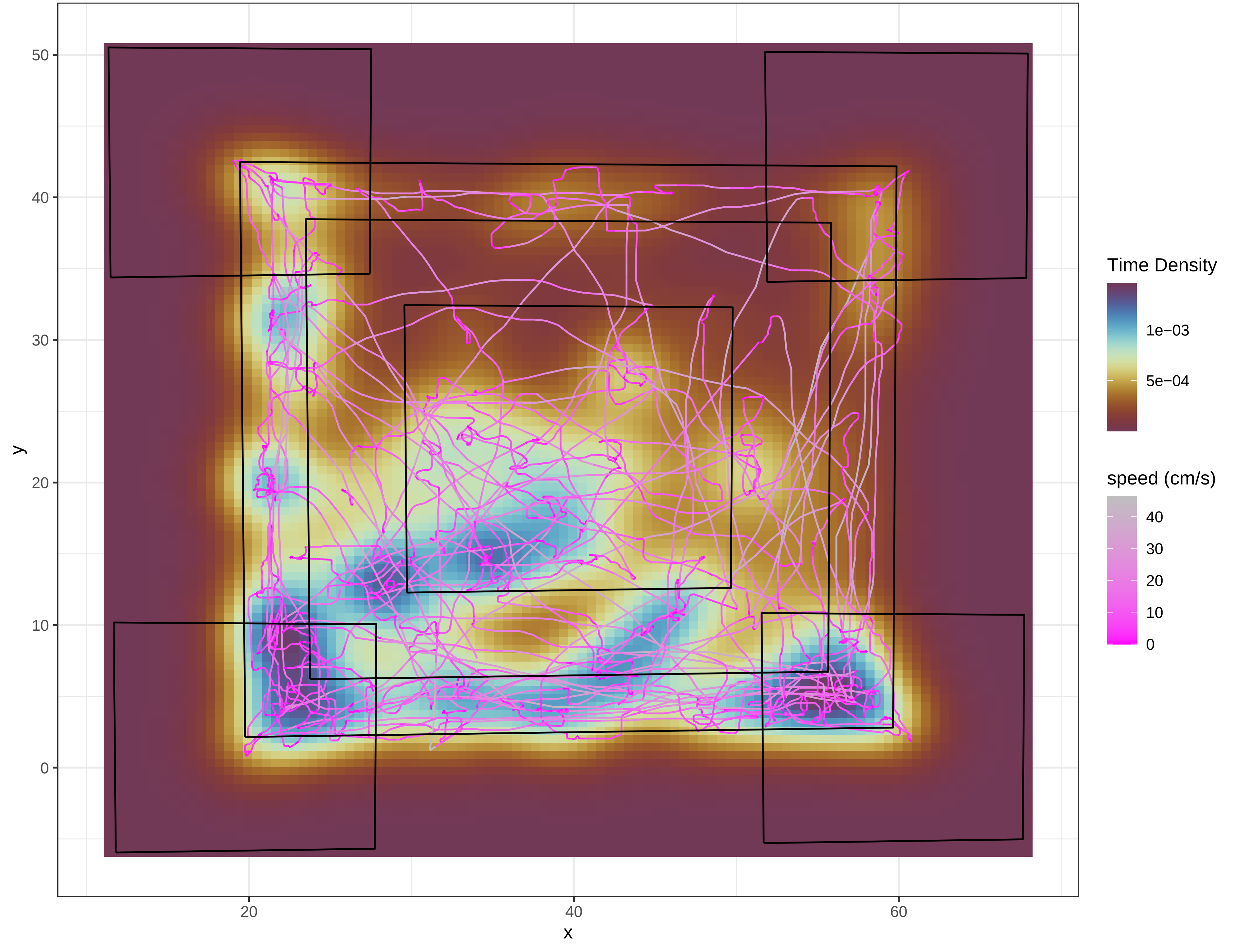

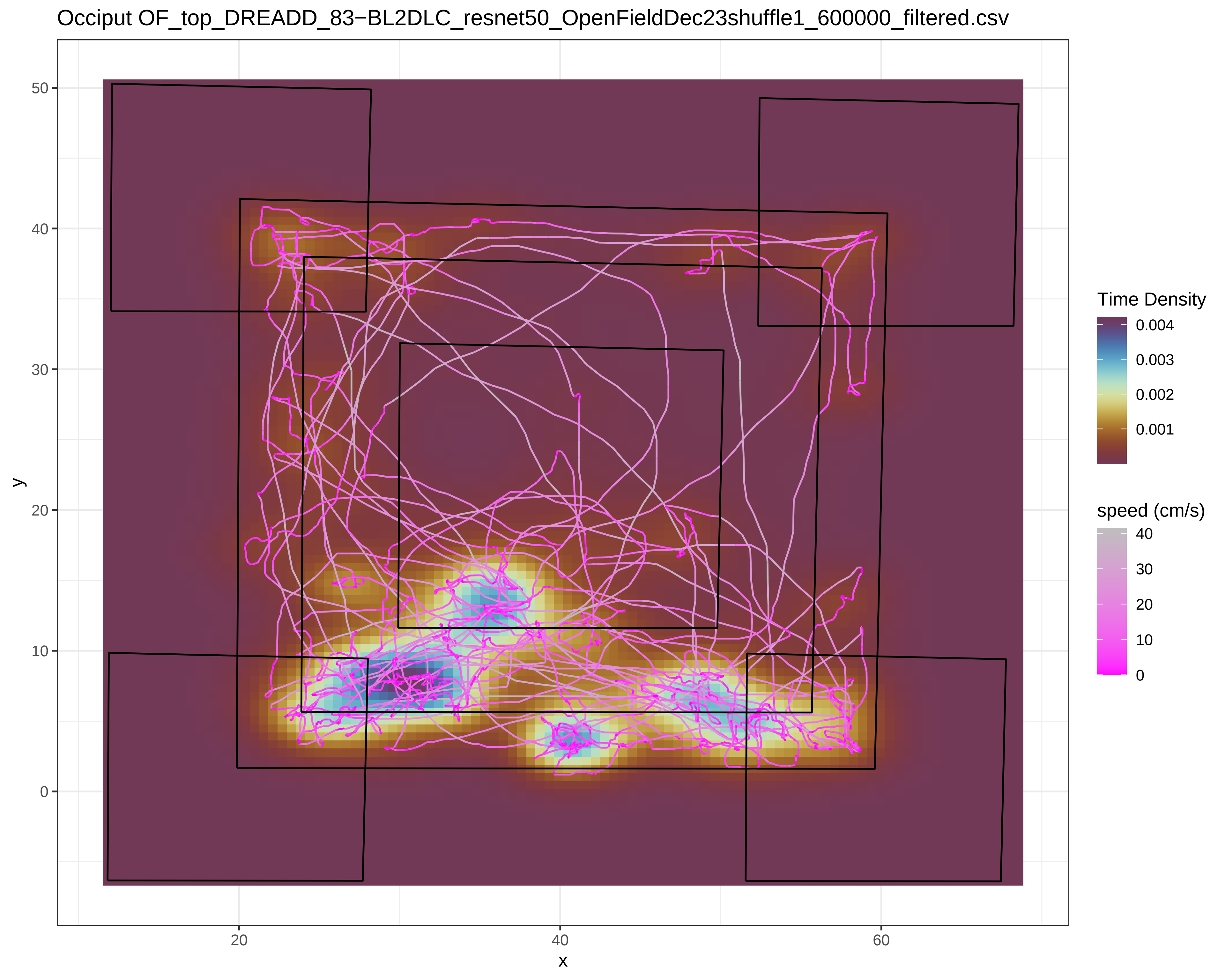

Occiput OF_top_DREADD_83−BL2DLC_resnet50_OpenFieldDec23shuffle1_600000_filtered.csv
y
x
50
40
30
20
10
0
20
40
60
Time Density
0.004
0.003
0.002
0.001
speed (cm/s)
40
30
20
10
0

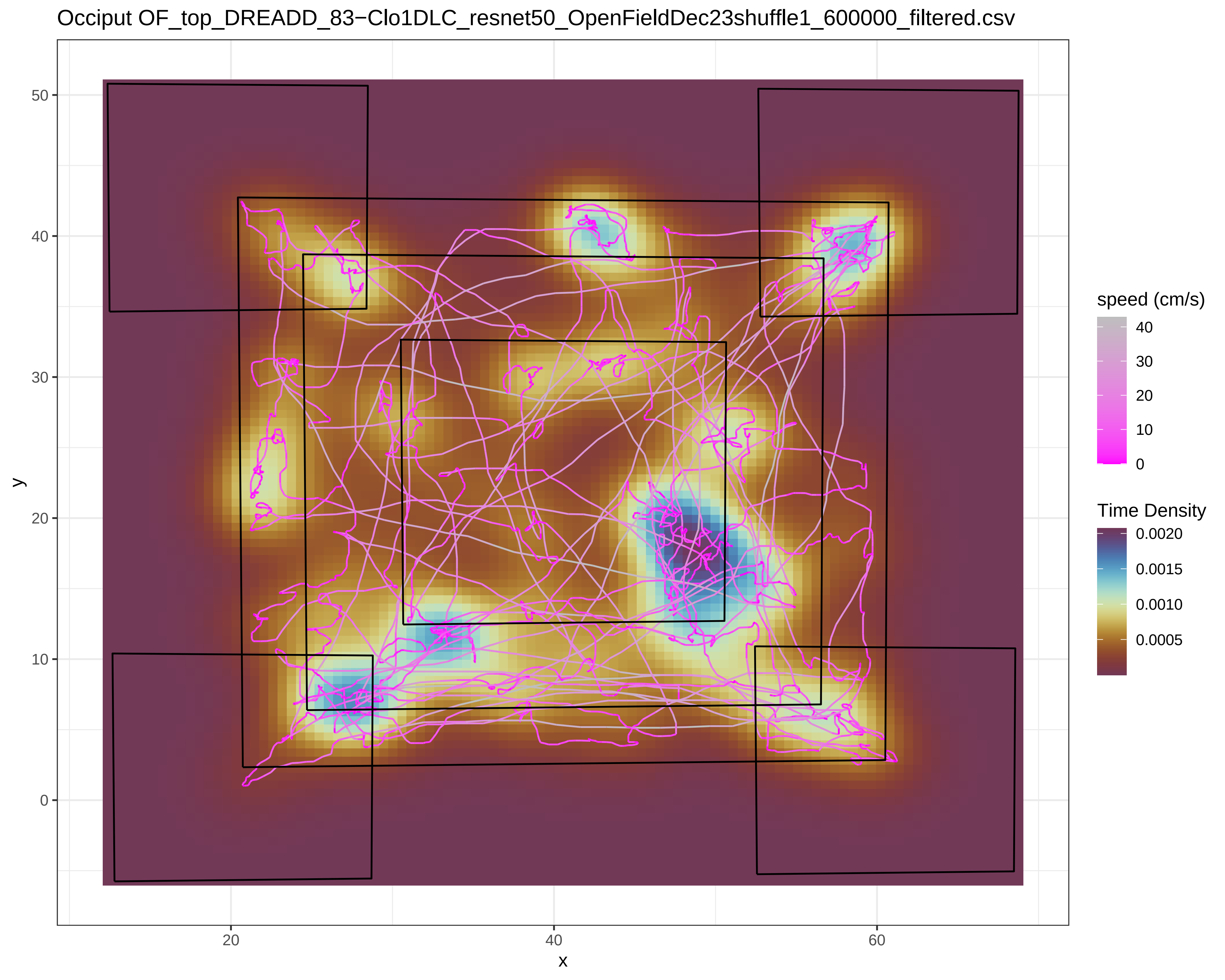

Occiput OF_top_DREADD_83−Clo1DLC_resnet50_OpenFieldDec23shuffle1_600000_filtered.csv
y
x
0
10
20
30
40
50
20
40
60
speed (cm/s)
40
30
20
10
0
Time Density
0.0020
0.0015
0.0010
0.0005

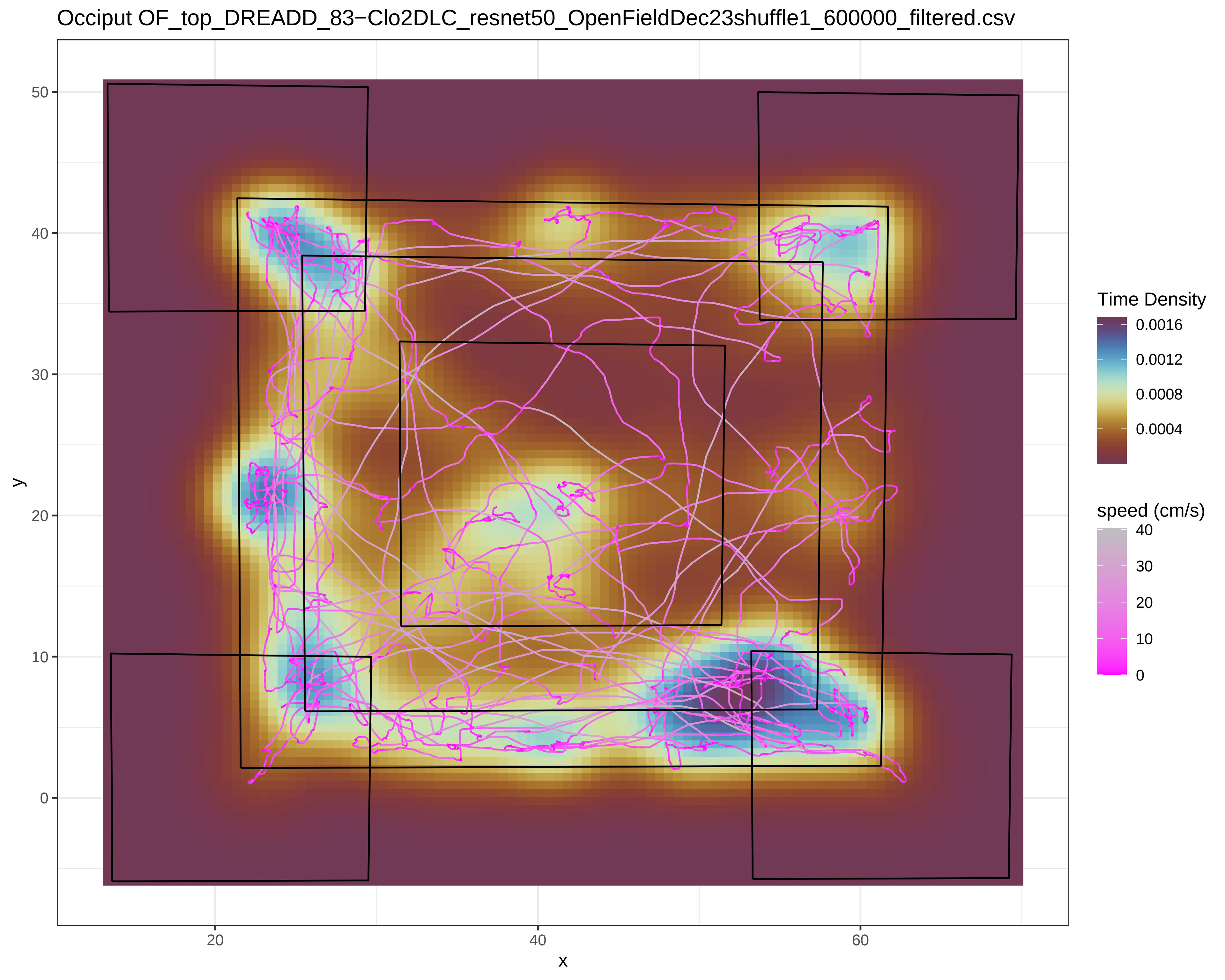
Occiput OF_top_DREADD_83−Clo2DLC_resnet50_OpenFieldDec23shuffle1_600000_filtered.csv
Time Density
0.0016
0.0012
0.0008
0.0004
speed (cm/s)
40
30
20
10
0
y
50
40
30
20
10
0
x
20
40
60

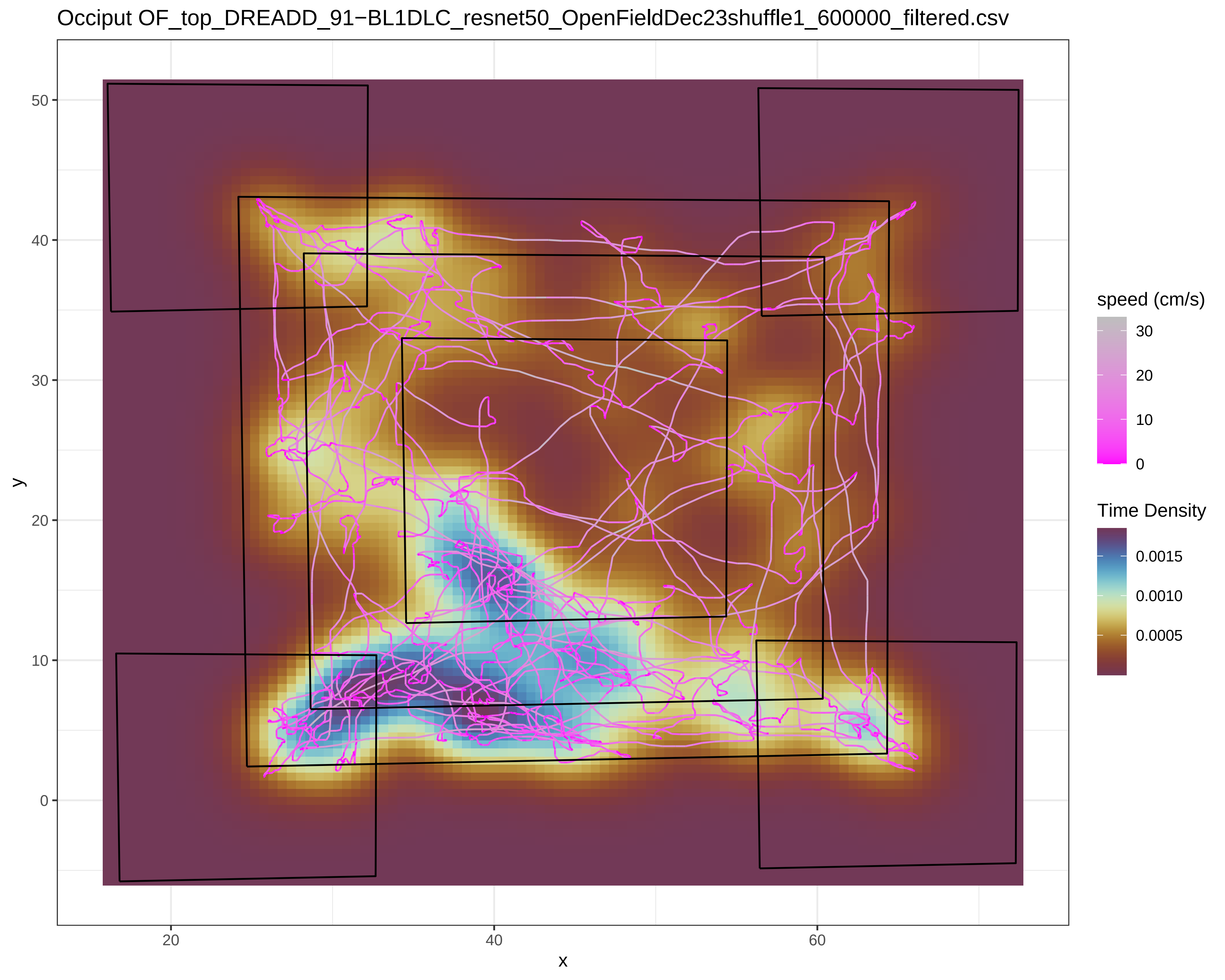
Occiput OF_top_DREADD_91−BL1DLC_resnet50_OpenFieldDec23shuffle1_600000_filtered.csv
y
x
50
40
30
20
10
0
20
40
60
speed (cm/s)
30
20
10
0
Time Density
0.0015
0.0010
0.0005

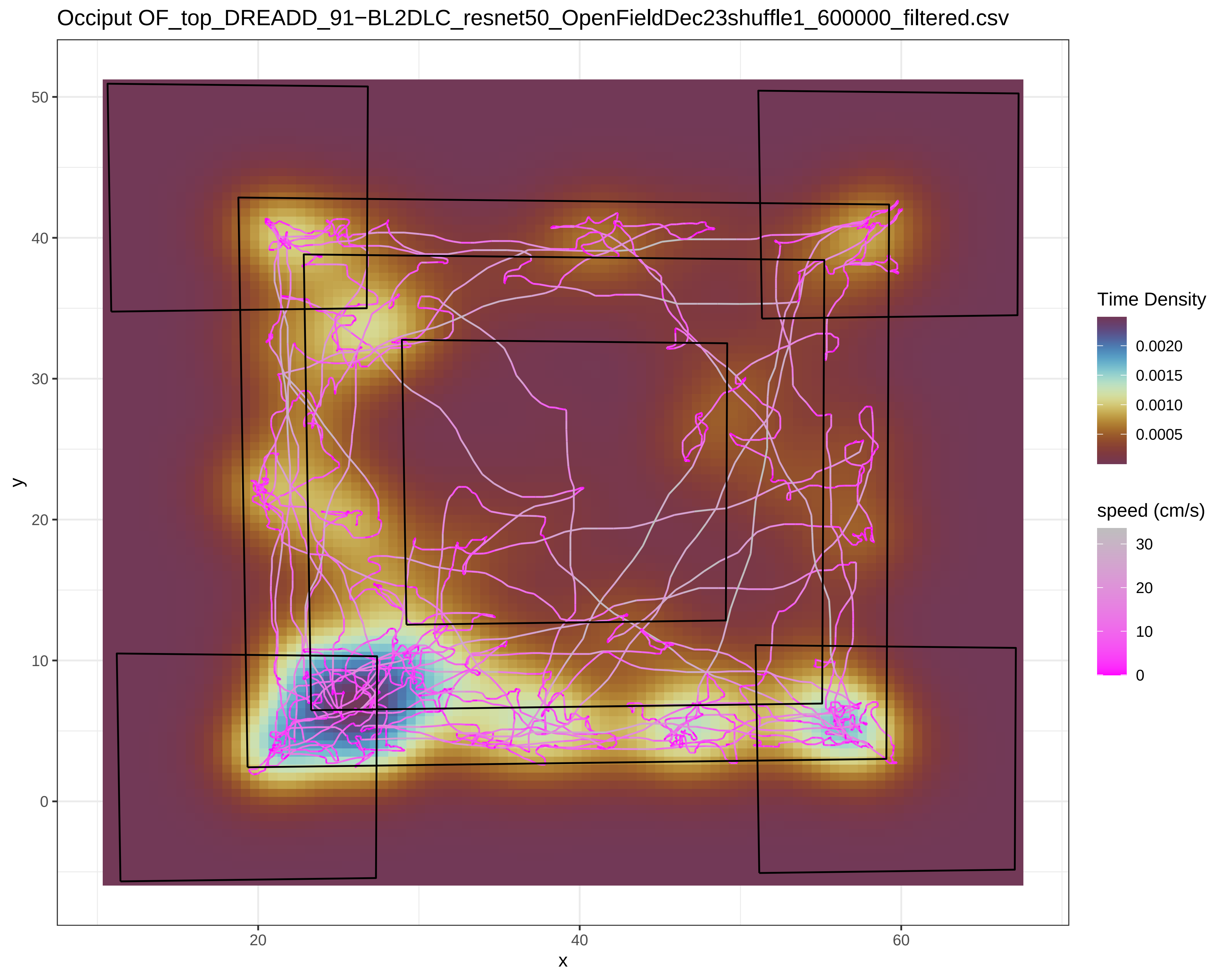
Occiput OF_top_DREADD_91−BL2DLC_resnet50_OpenFieldDec23shuffle1_600000_filtered.csv
y
50
40
30
20
10
0
x
20
40
60
Time Density
0.0020
0.0015
0.0010
0.0005
speed (cm/s)
30
20
10
0

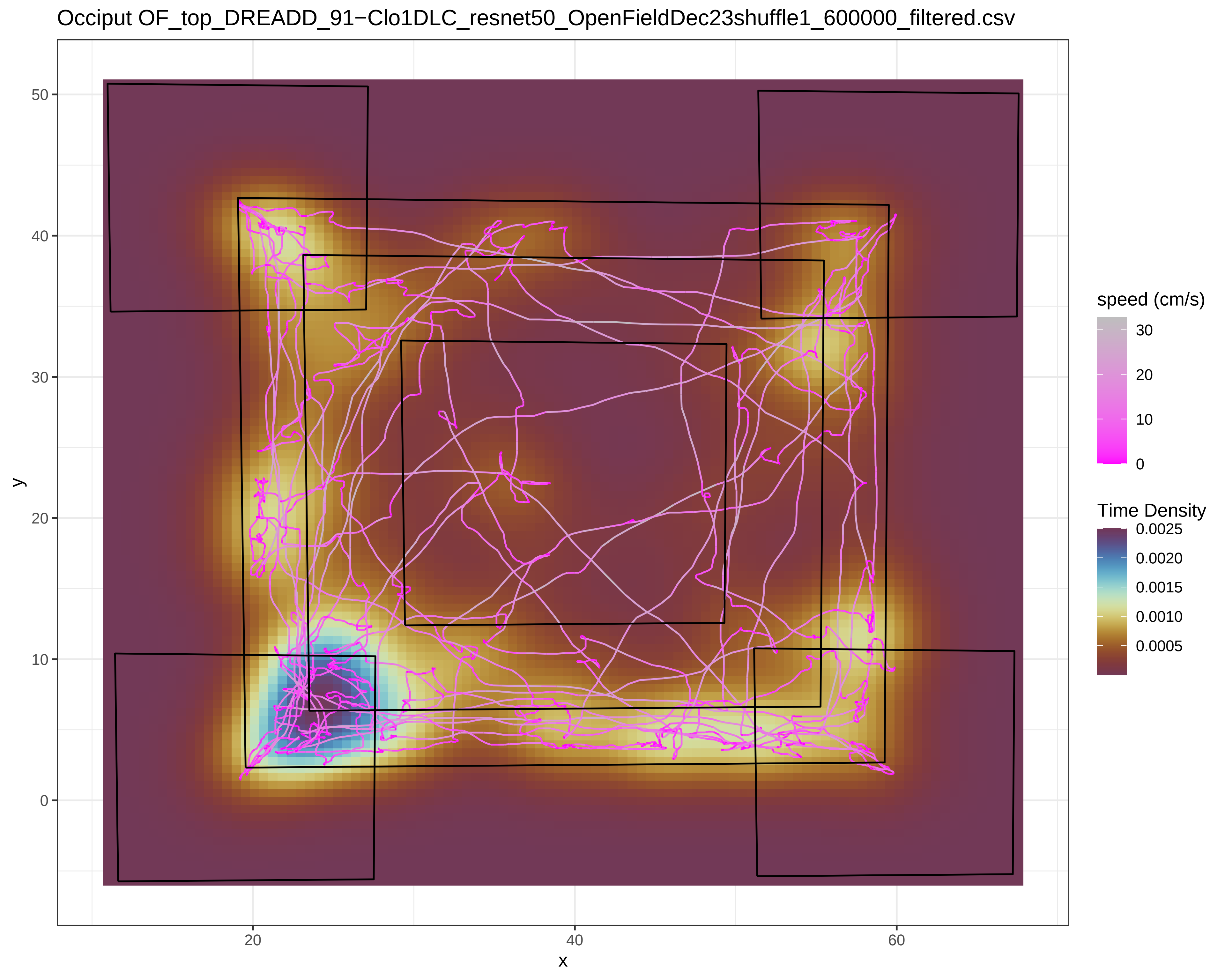
Occiput OF_top_DREADD_91−Clo1DLC_resnet50_OpenFieldDec23shuffle1_600000_filtered.csv
y
x
50
40
30
20
10
0
20
40
60
speed (cm/s)
30
20
10
0
Time Density
0.0025
0.0020
0.0015
0.0010
0.0005

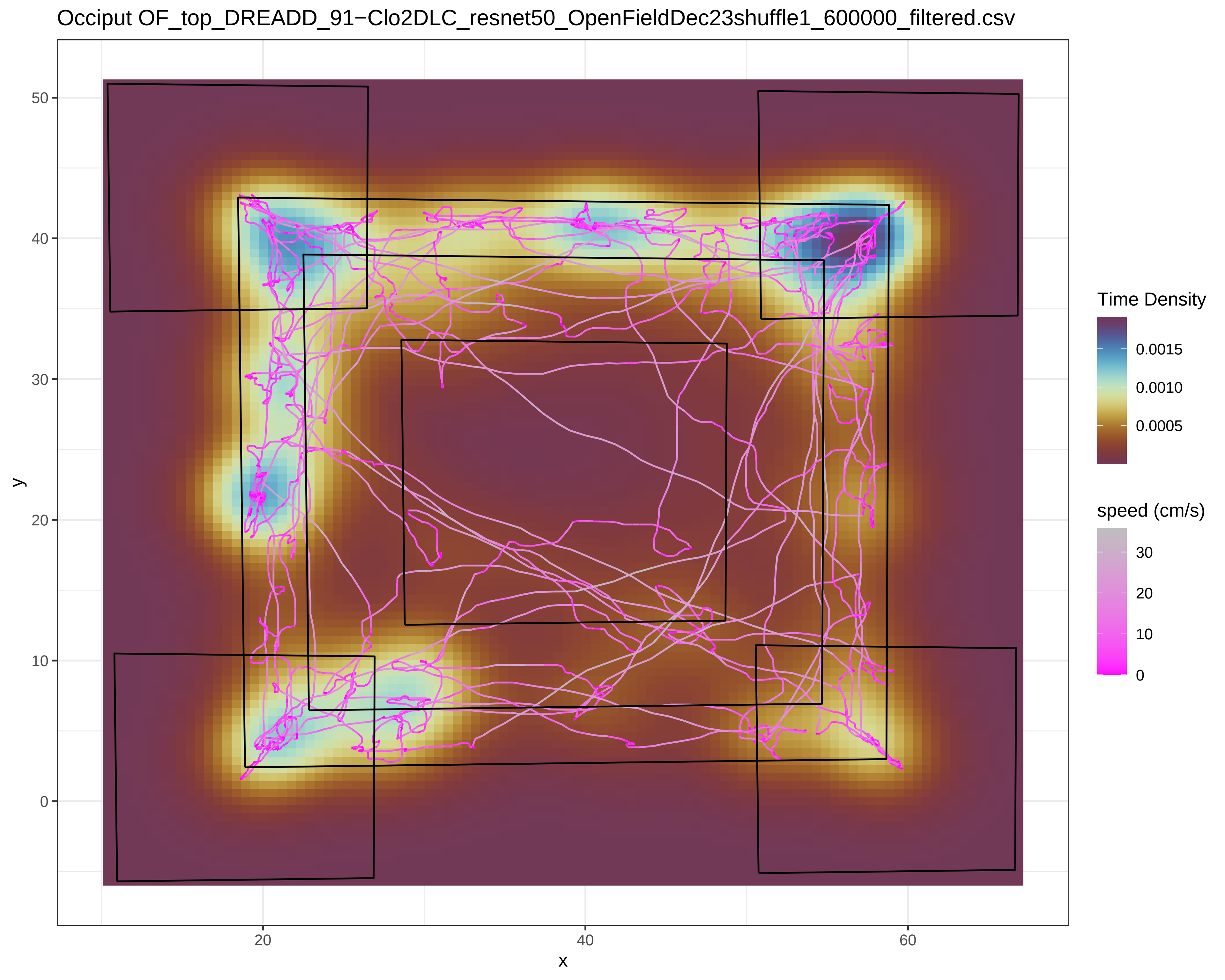

Occiput OF_top_DREADD_91−Clo2DLC_resnet50_OpenFieldDec23shuffle1_600000_filtered.csv
y
x
50
40
30
20
10
0
20
40
60
Time Density
0.0015
0.0010
0.0005
speed (cm/s)
30
20
10
0

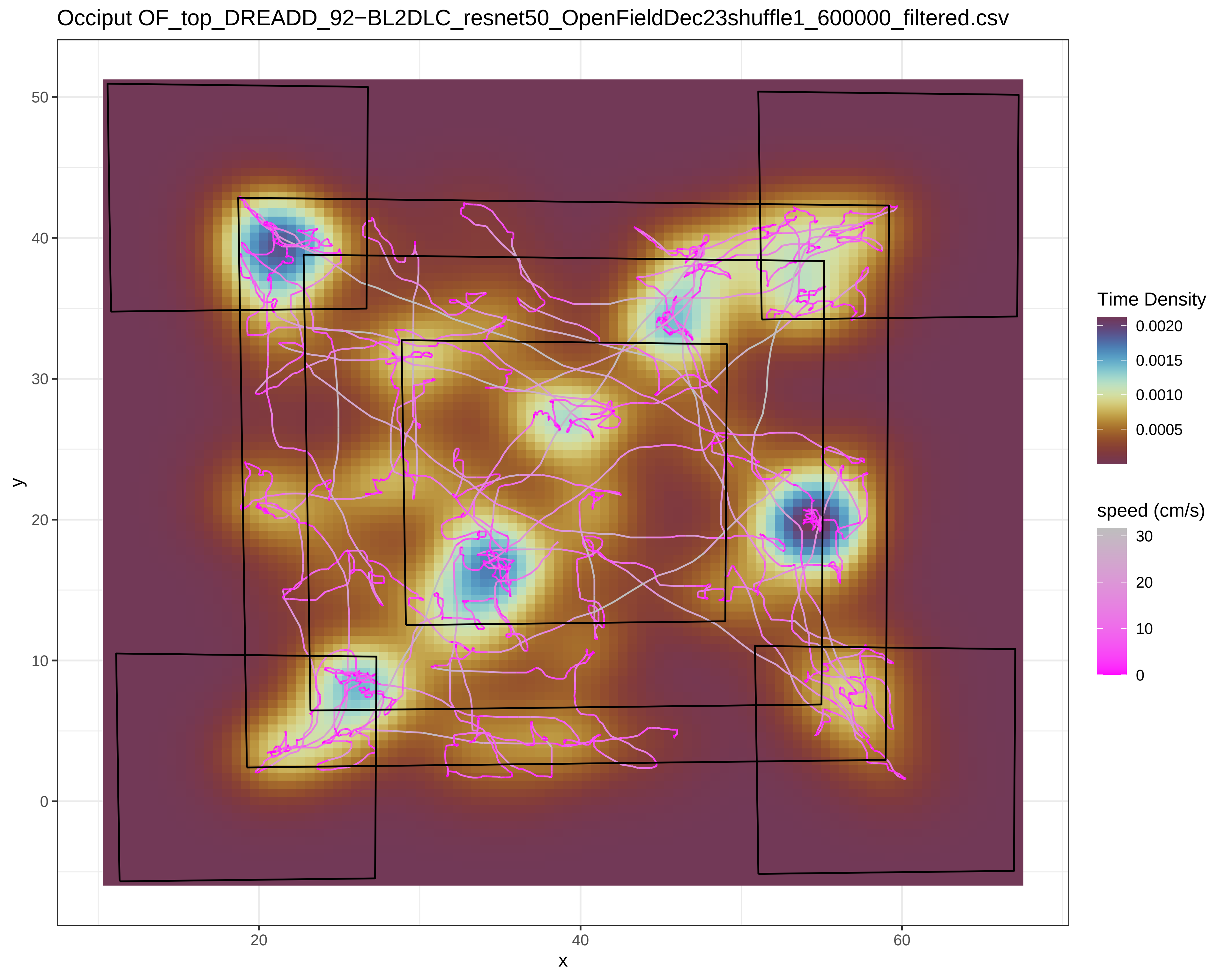

Occiput OF_top_DREADD_92−BL2DLC_resnet50_OpenFieldDec23shuffle1_600000_filtered.csv
y
x
50
40
30
20
10
0
20
40
60
Time Density
0.0020
0.0015
0.0010
0.0005
speed (cm/s)
30
20
10
0

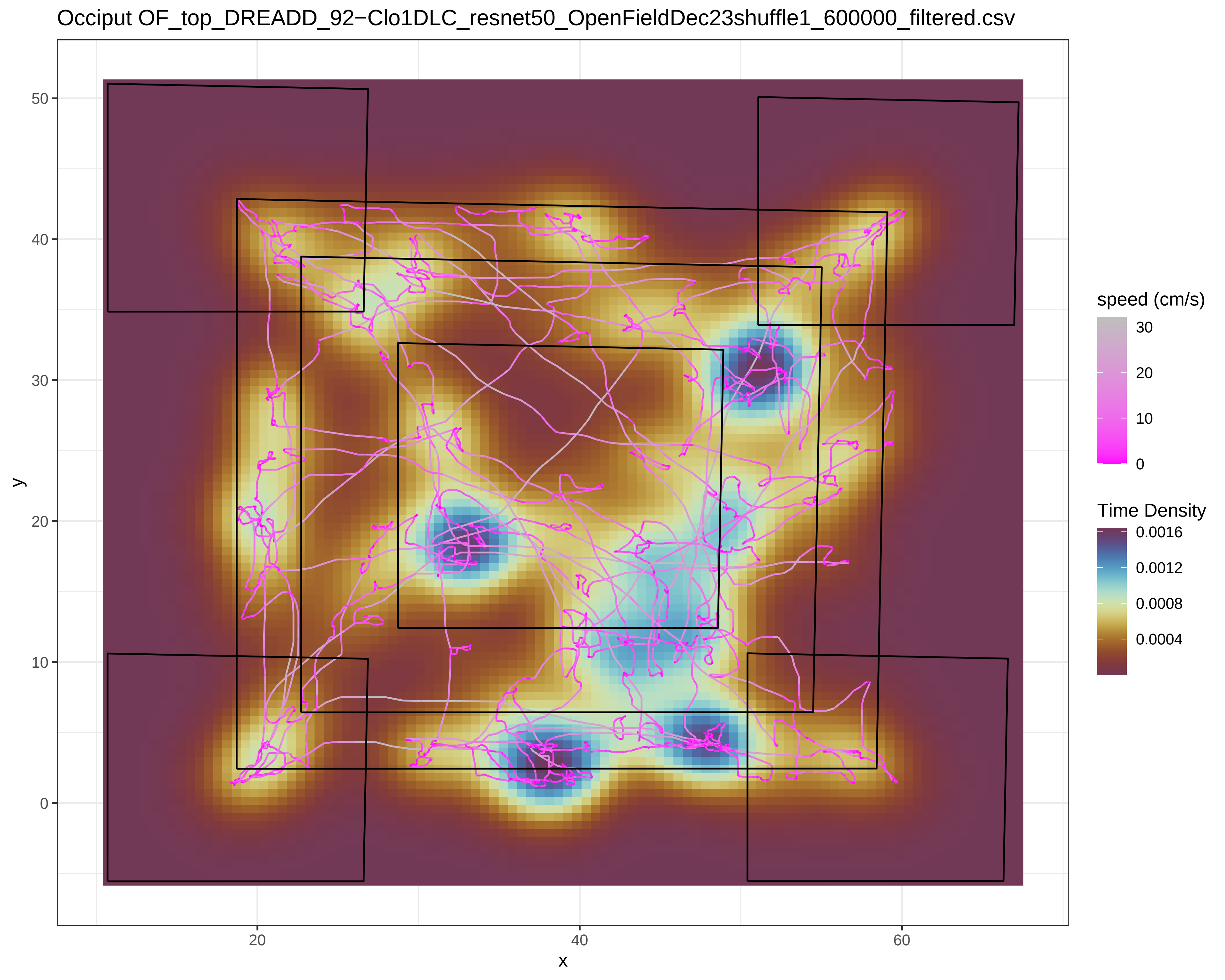
Occiput OF_top_DREADD_92−Clo1DLC_resnet50_OpenFieldDec23shuffle1_600000_filtered.csv
y
x
0
10
20
30
40
50
20
40
60
speed (cm/s)
30
20
10
0
Time Density
0.0016
0.0012
0.0008
0.0004

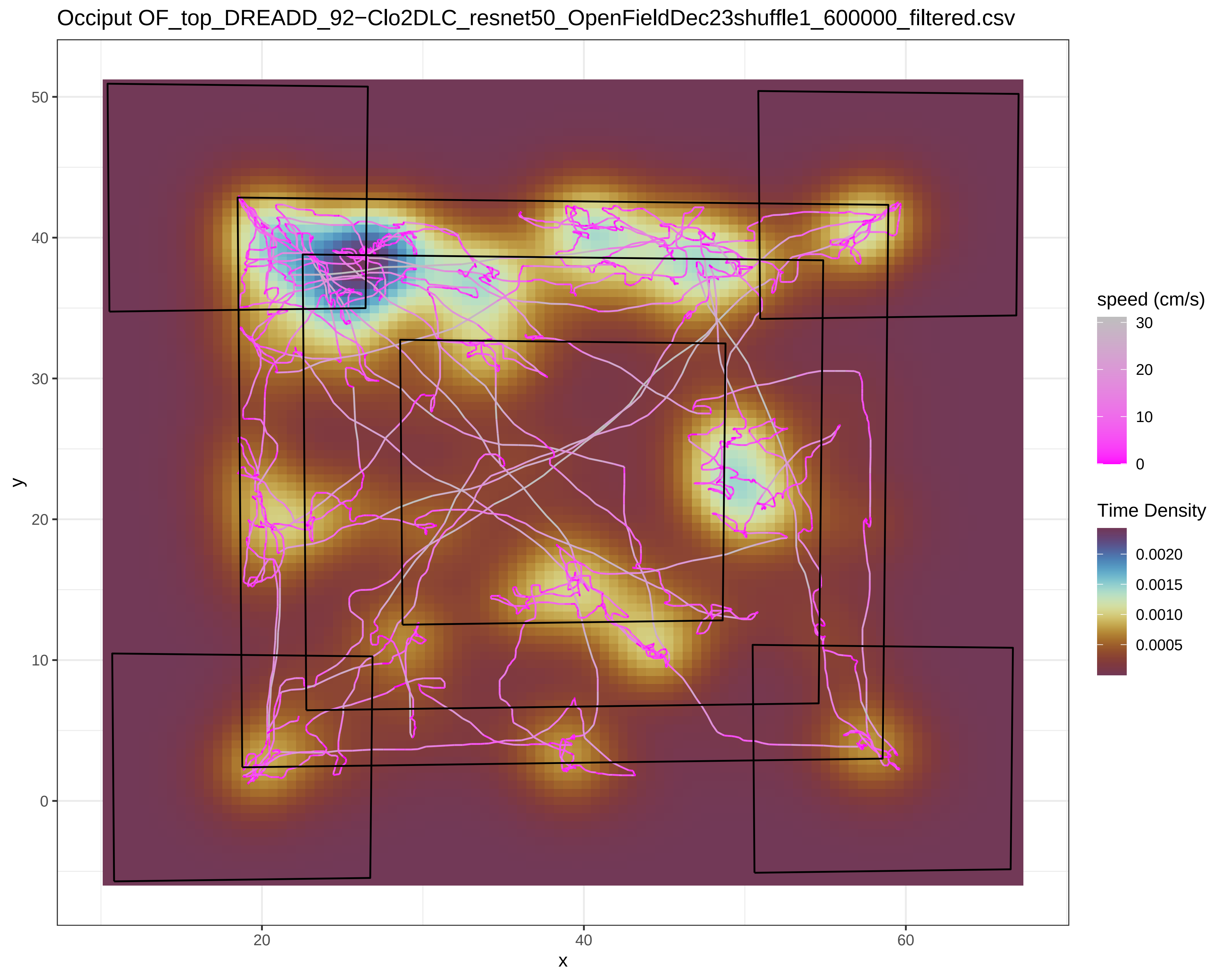

Occiput OF_top_DREADD_92−Clo2DLC_resnet50_OpenFieldDec23shuffle1_600000_filtered.csv
speed (cm/s)
30
20
10
0
Time Density
0.0020
0.0015
0.0010
0.0005
y
50
40
30
20
10
0
x
20
40
60

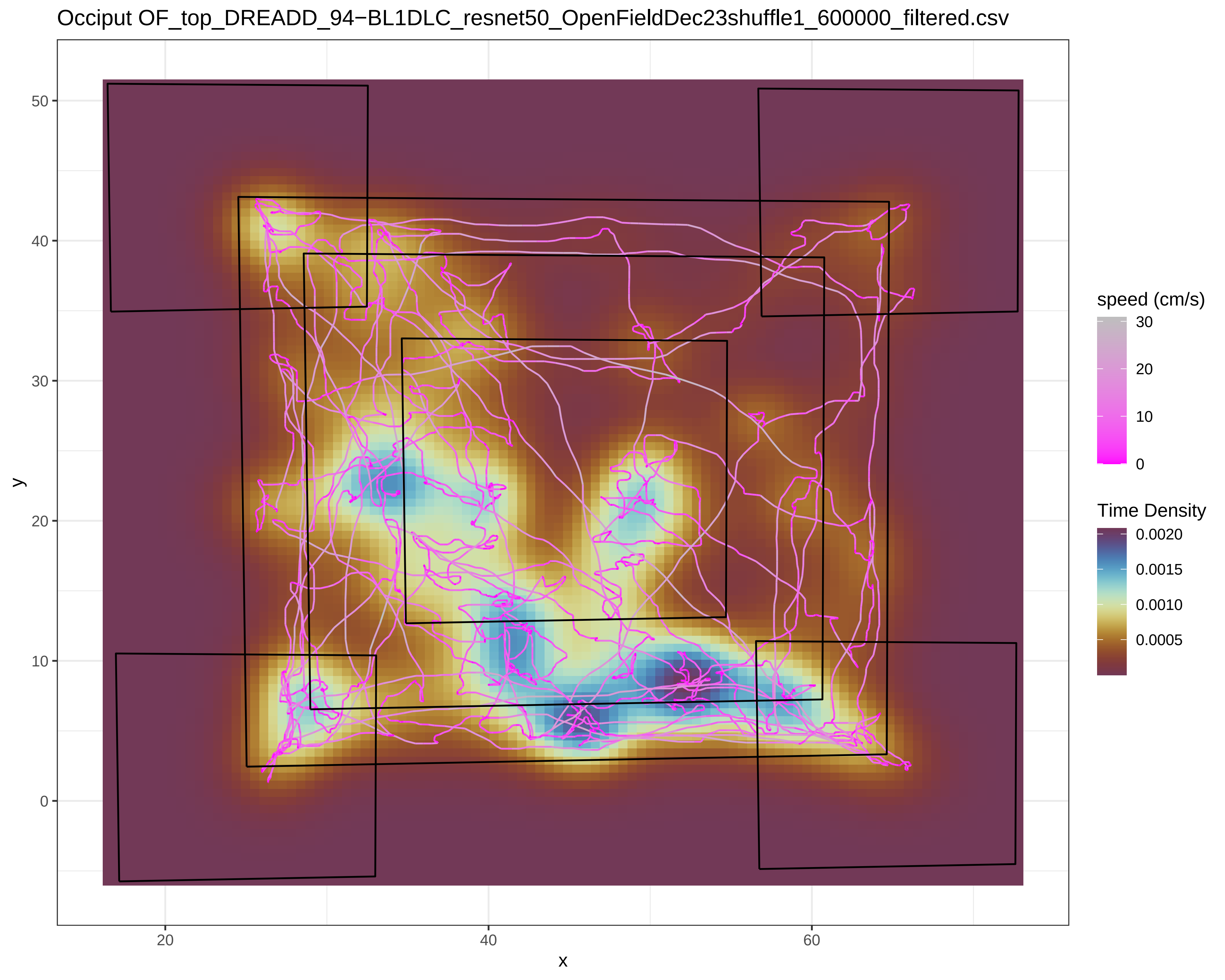
Occiput OF_top_DREADD_94−BL1DLC_resnet50_OpenFieldDec23shuffle1_600000_filtered.csv
y
x
0
10
20
30
40
50
20
40
60
speed (cm/s)
30
20
10
0
Time Density
0.0020
0.0015
0.0010
0.0005

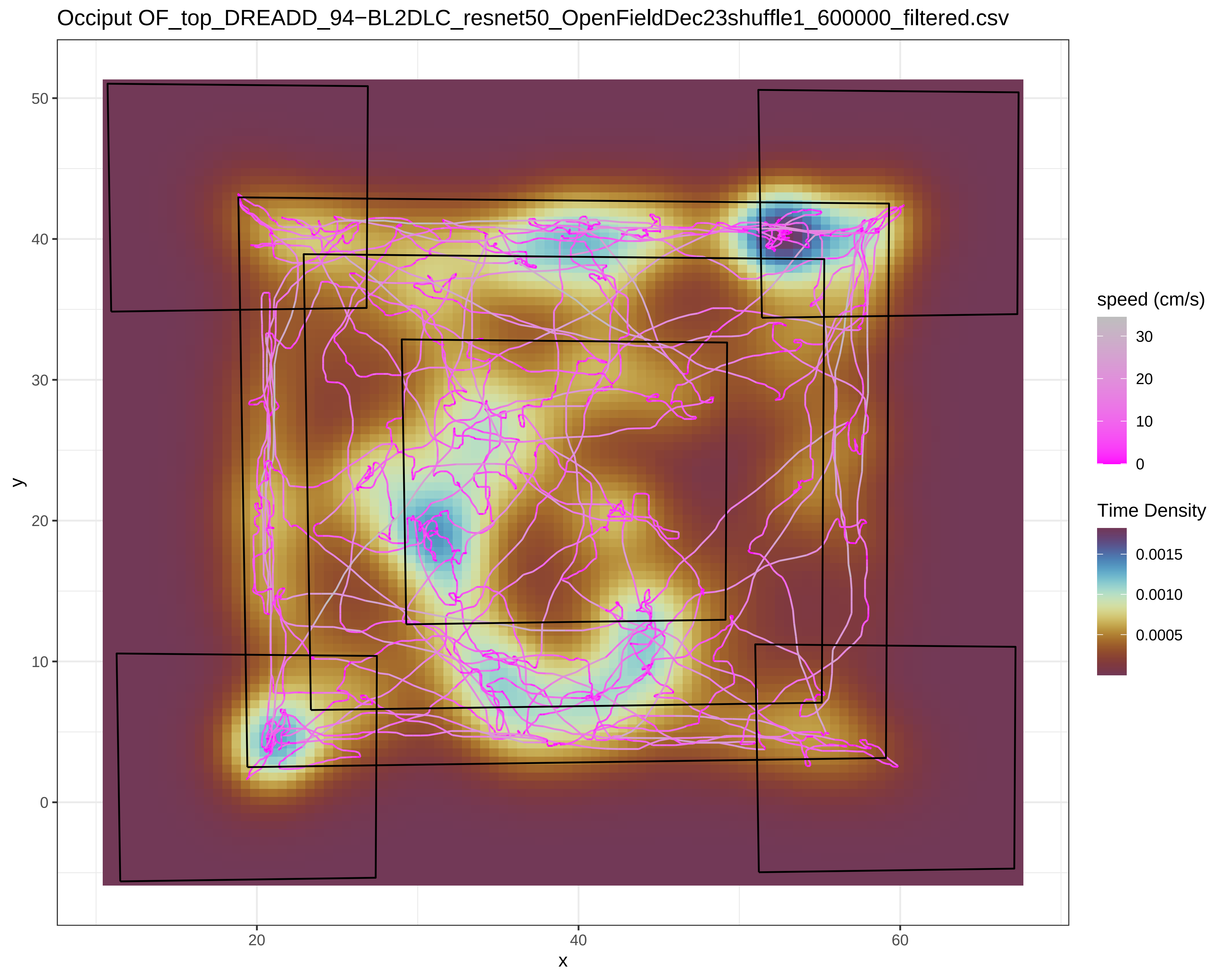

Occiput OF_top_DREADD_94−BL2DLC_resnet50_OpenFieldDec23shuffle1_600000_filtered.csv
y
50
40
30
20
10
0
x
20
40
60
speed (cm/s)
30
20
10
0
Time Density
0.0015
0.0010
0.0005

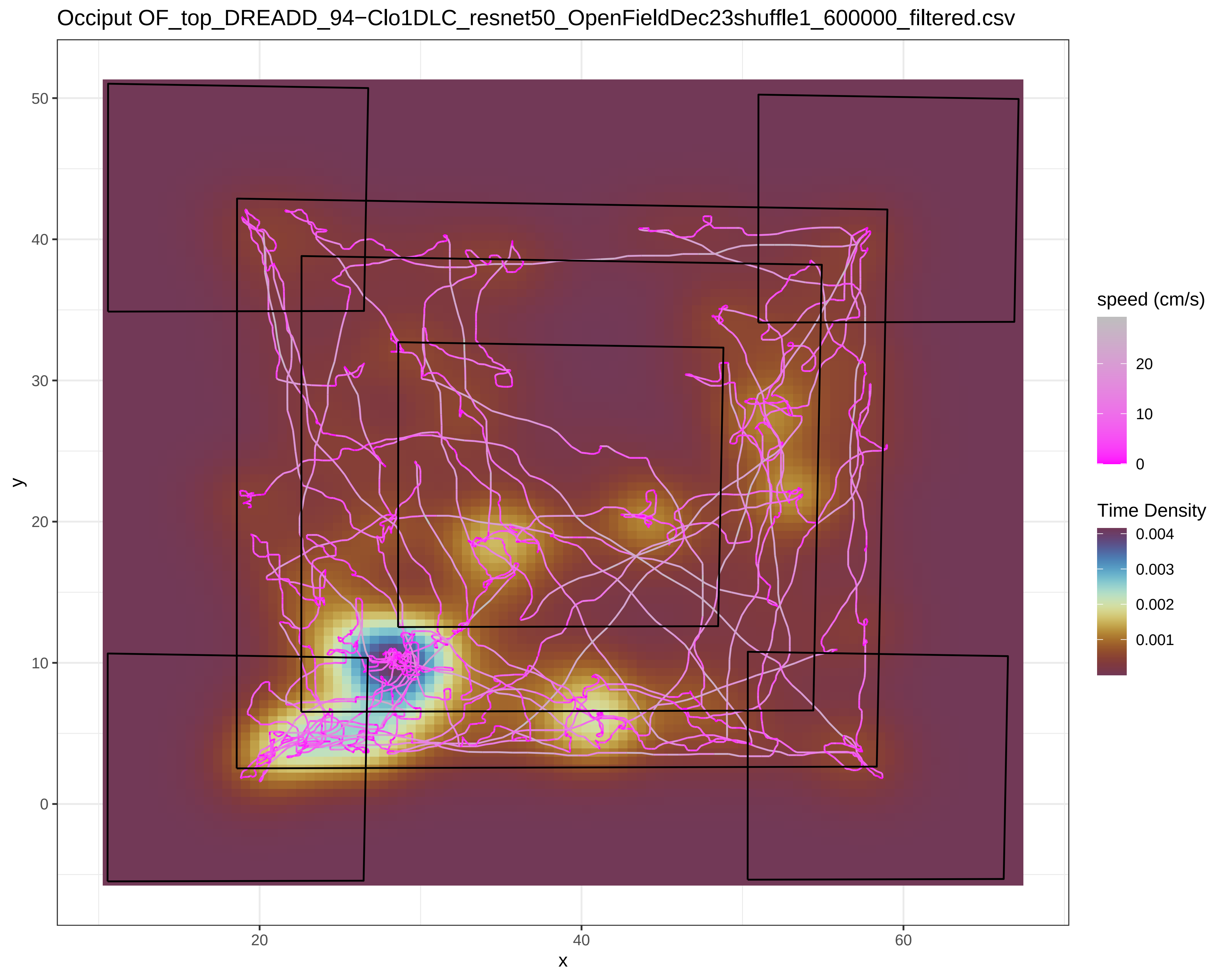

Occiput OF_top_DREADD_94−Clo1DLC_resnet50_OpenFieldDec23shuffle1_600000_filtered.csv
y
x
0
10
20
30
40
50
20
40
60
speed (cm/s)
20
10
0
Time Density
0.004
0.003
0.002
0.001

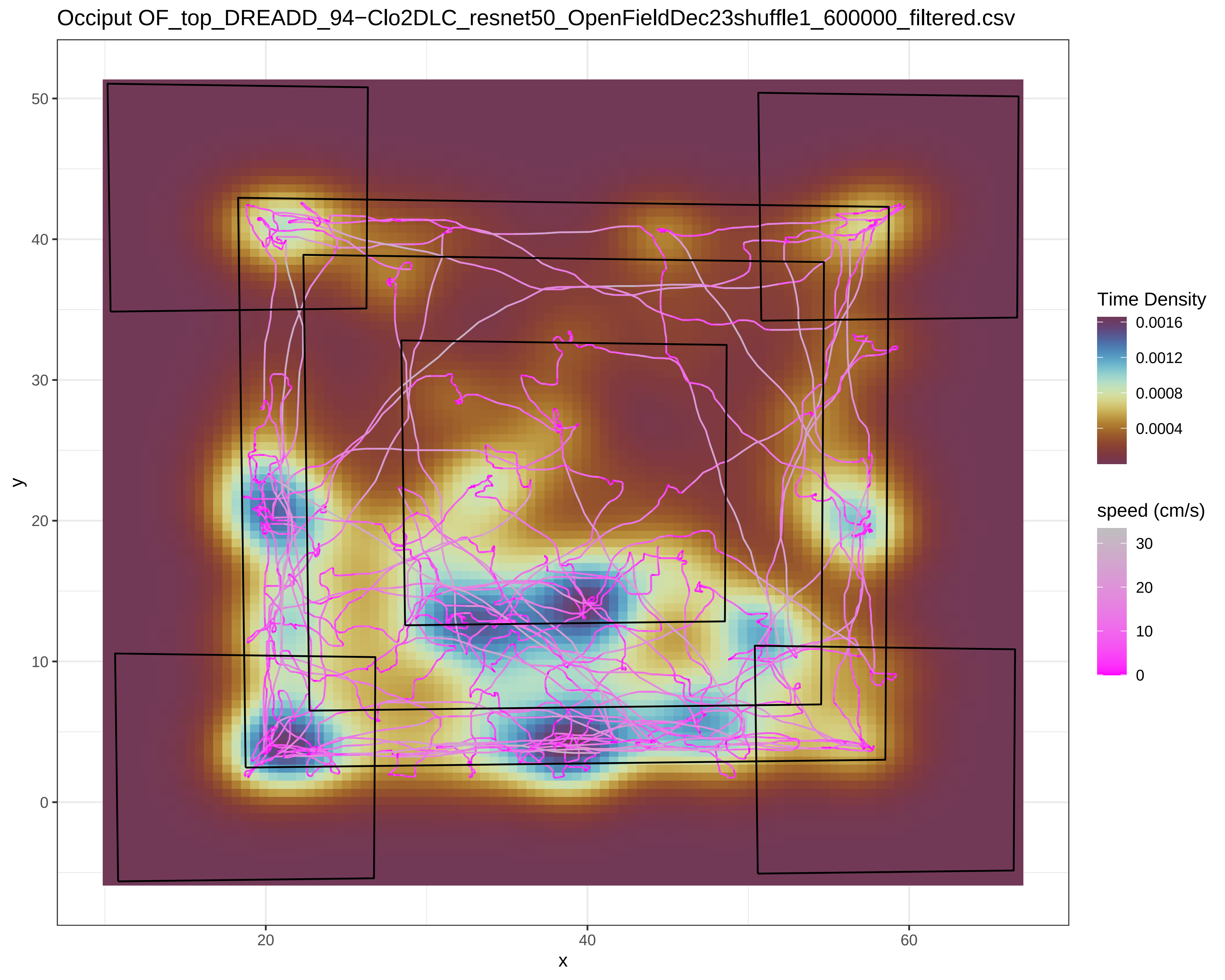
Occiput OF_top_DREADD_94−Clo2DLC_resnet50_OpenFieldDec23shuffle1_600000_filtered.csv
y
x
50
40
30
20
10
0
20
40
60
Time Density
0.0016
0.0012
0.0008
0.0004
speed (cm/s)
30
20
10
0
