## SupplementaryFile_S3 for "Chemo- and optogenetic activation of hypothalamic Foxb1-expressing neurons and their terminal endings in the rostral-dorsolateral PAG leads to tachypnea, bradycardia, and immobility"

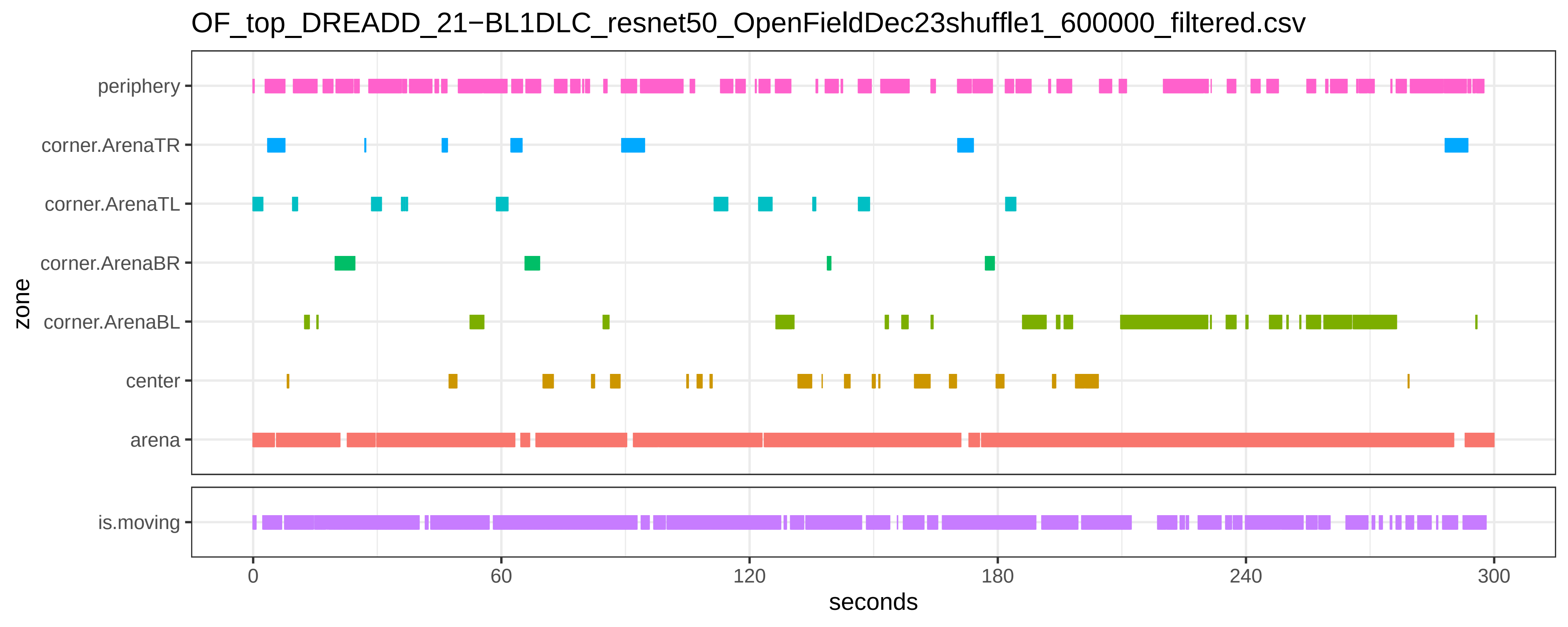
OF_top_DREADD_21−BL1DLC_resnet50_OpenFieldDec23shuffle1_600000_filtered.csv
periphery
corner.ArenaTR
corner.ArenaTL
corner.ArenaBR
corner.ArenaBL
center
arena
is.moving
zone
0
60
120
180
240
300
seconds

OF_top_DREADD_21−BL2DLC_resnet50_OpenFieldDec23shuffle1_600000_filtered.csv

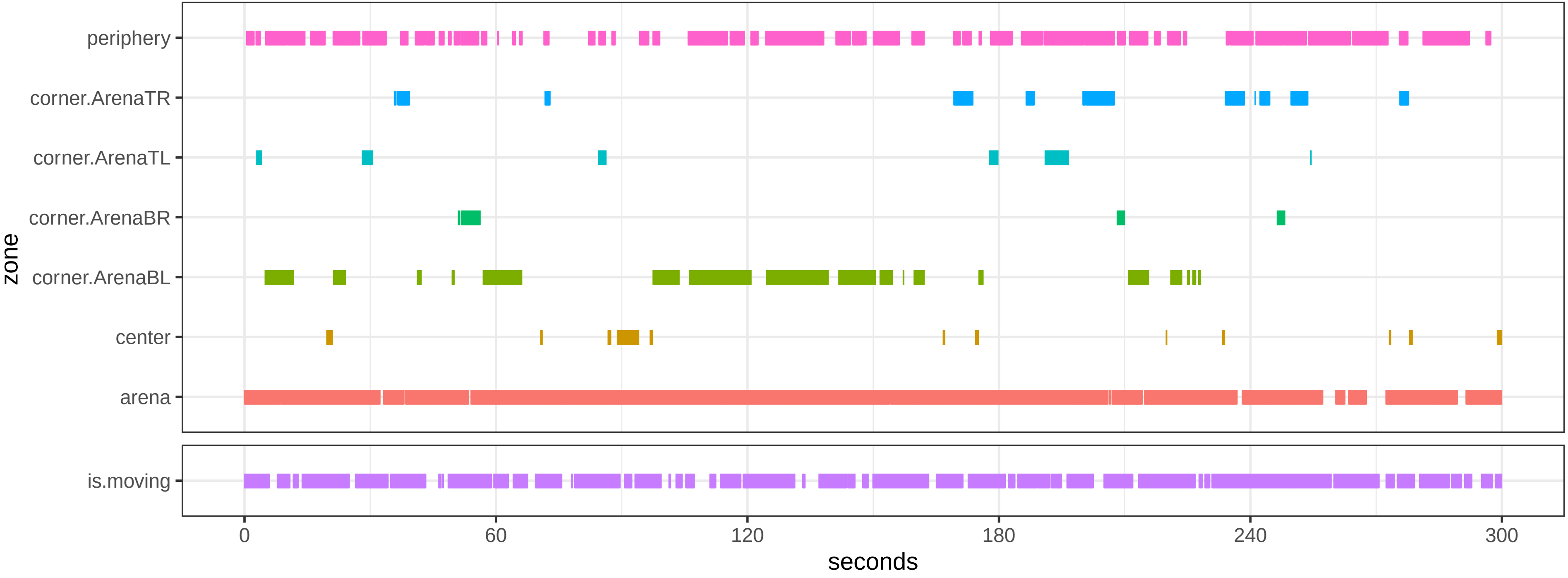

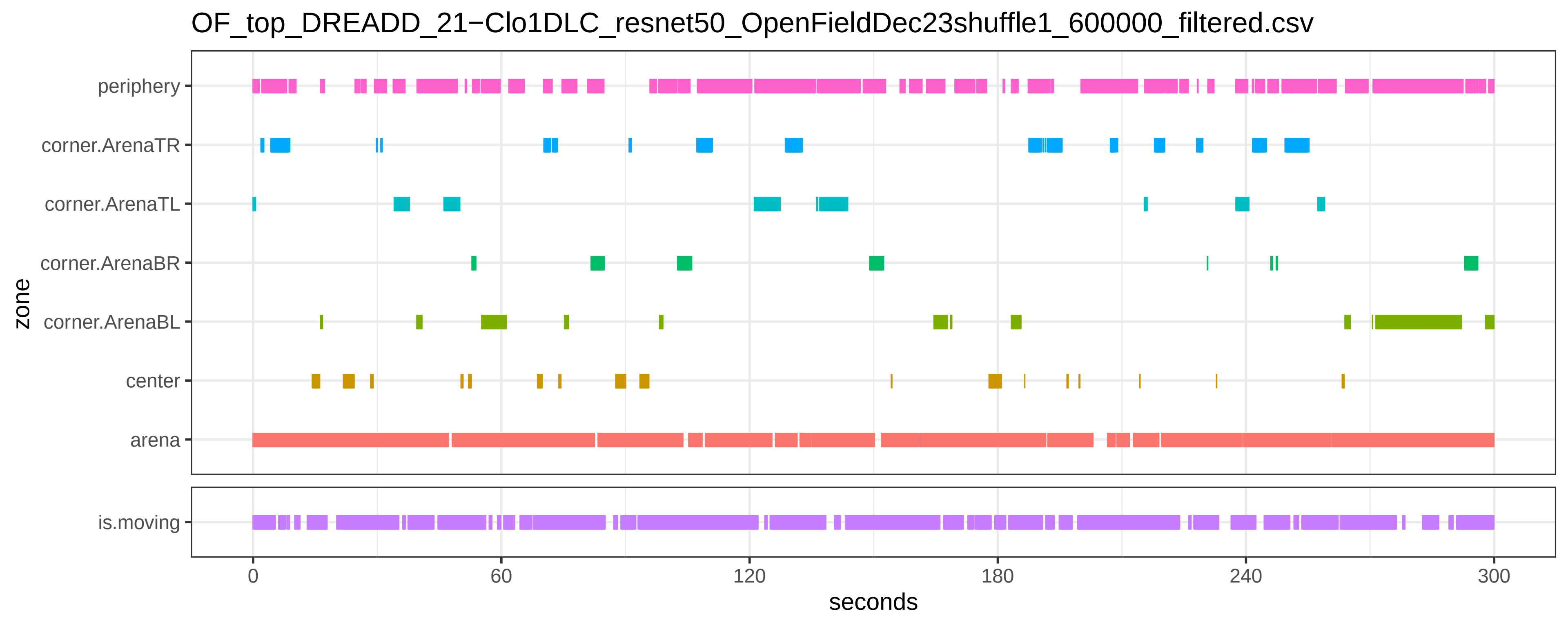
OF_top_DREADD_21−Clo1DLC_resnet50_OpenFieldDec23shuffle1_600000_filtered.csv
periphery
corner.ArenaTR
corner.ArenaTL
corner.ArenaBR
corner.ArenaBL
center
arena
zone
is.moving
0
60
120
180
240
300
seconds

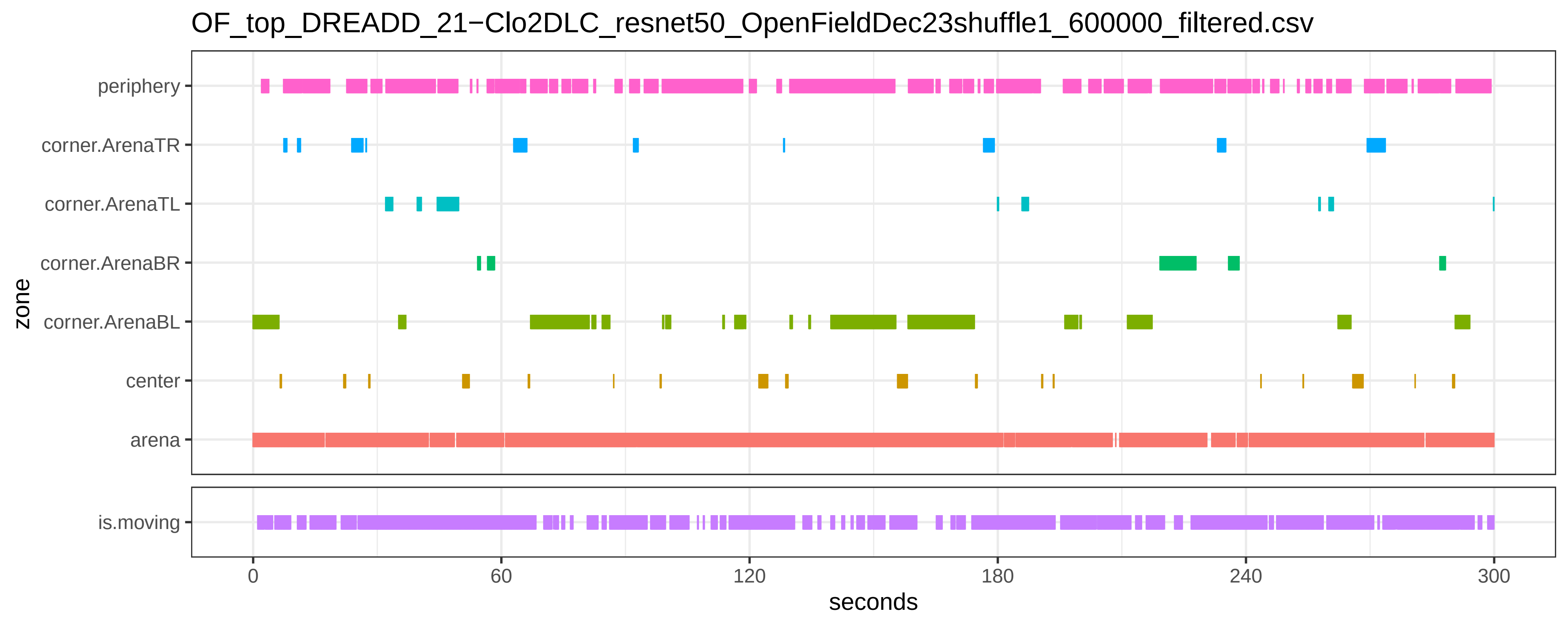

OF_top_DREADD_21−Clo2DLC_resnet50_OpenFieldDec23shuffle1_600000_filtered.csv
periphery
corner.ArenaTR
corner.ArenaTL
corner.ArenaBR
corner.ArenaBL
center
arena
zone
is.moving
0
60
120
180
240
300
seconds

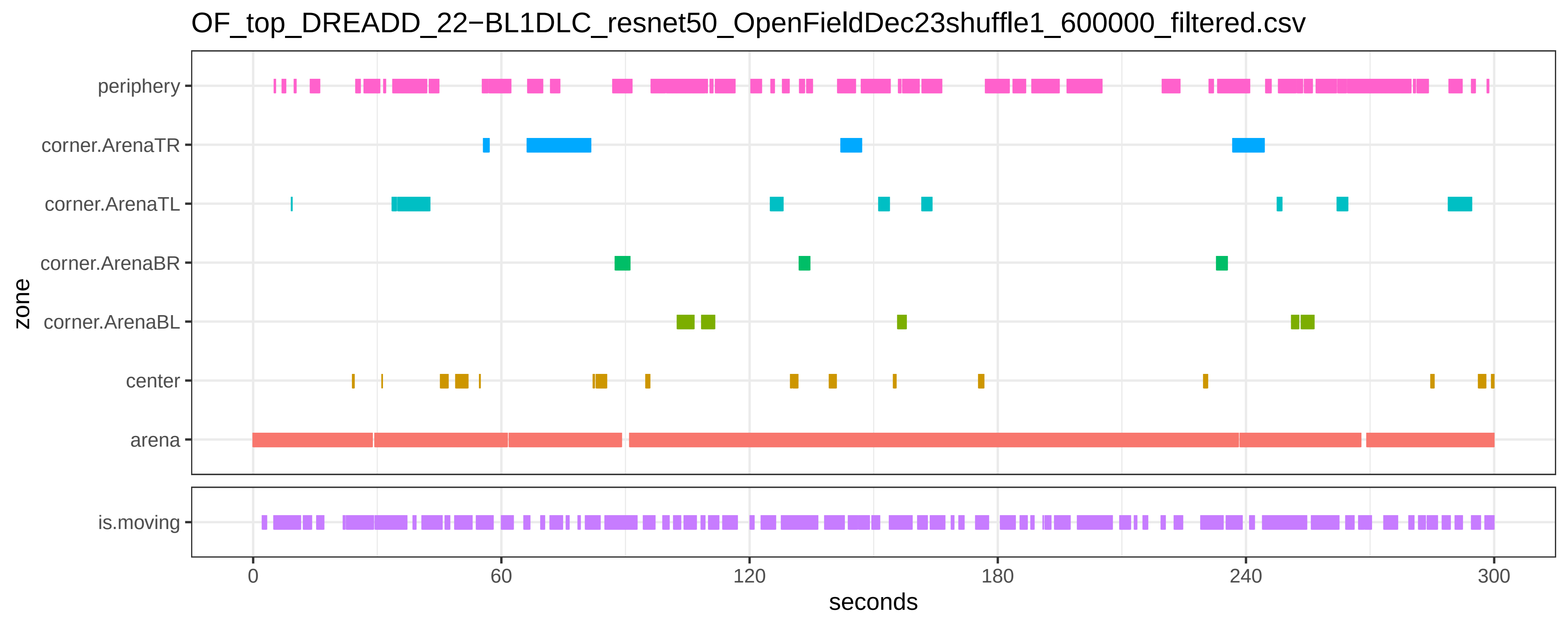
OF_top_DREADD_22−BL1DLC_resnet50_OpenFieldDec23shuffle1_600000_filtered.csv
periphery
corner.ArenaTR
corner.ArenaTL
corner.ArenaBR
corner.ArenaBL
center
arena
zone
is.moving
0
60
120
180
240
300
seconds

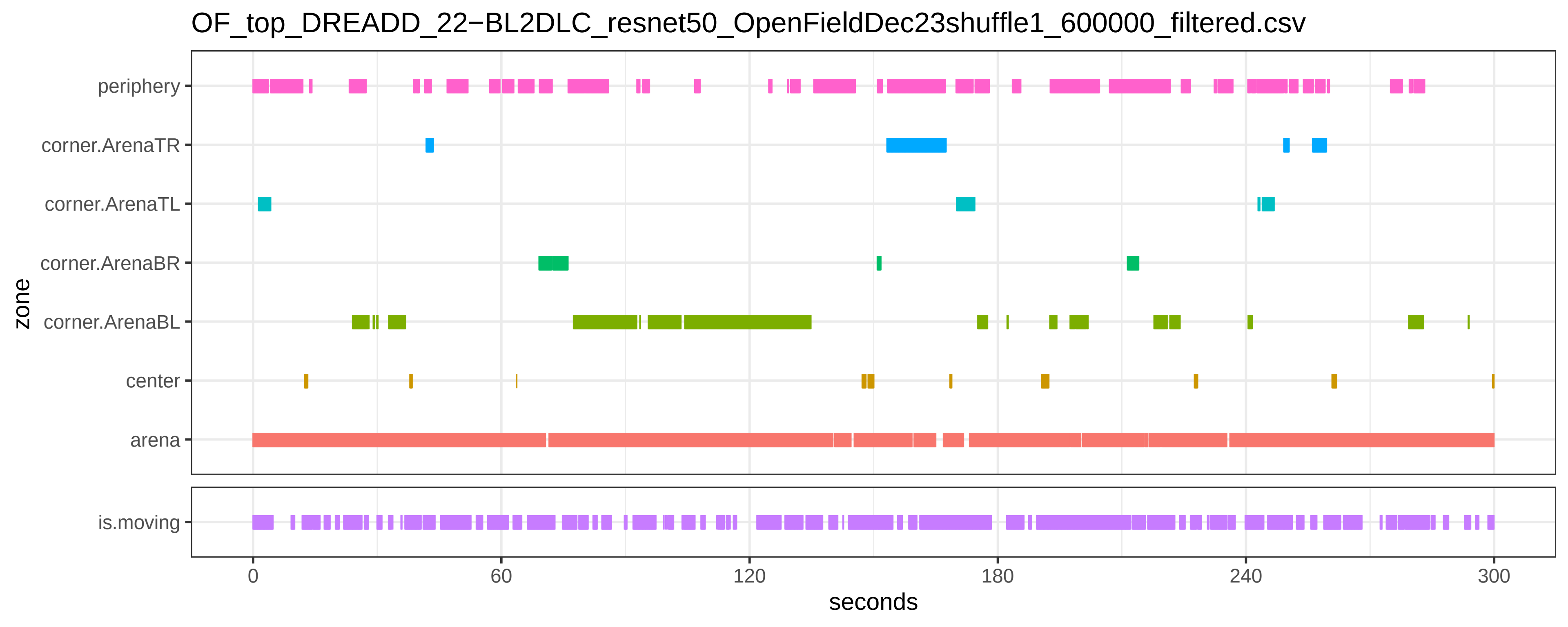
OF_top_DREADD_22−BL2DLC_resnet50_OpenFieldDec23shuffle1_600000_filtered.csv
periphery
corner.ArenaTR
corner.ArenaTL
corner.ArenaBR
corner.ArenaBL
center
arena
is.moving
zone
0
60
120
180
240
300
seconds

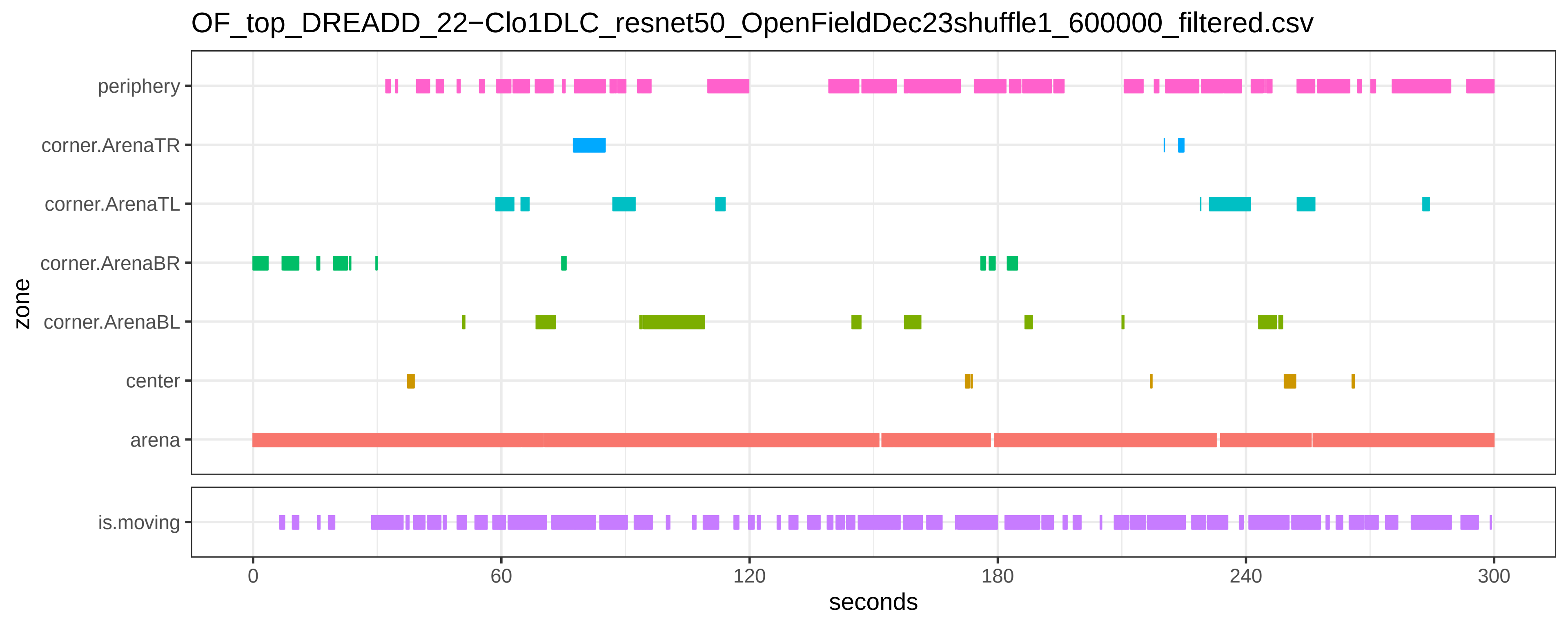
OF_top_DREADD_22−Clo1DLC_resnet50_OpenFieldDec23shuffle1_600000_filtered.csv
periphery
corner.ArenaTR
corner.ArenaTL
corner.ArenaBR
corner.ArenaBL
center
arena
zone
is.moving
0
60
120
180
240
300
seconds

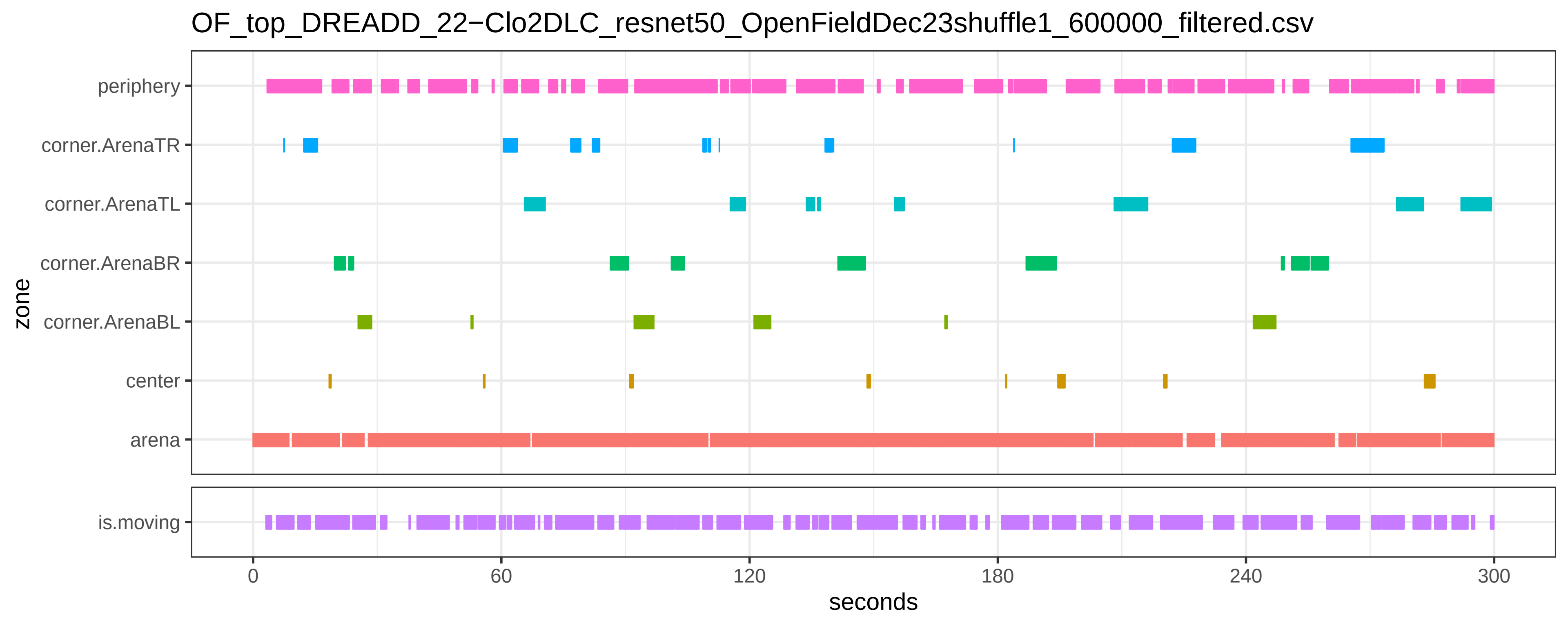

OF_top_DREADD_22−Clo2DLC_resnet50_OpenFieldDec23shuffle1_600000_filtered.csv
periphery
corner.ArenaTR
corner.ArenaTL
corner.ArenaBR
corner.ArenaBL
center
arena
zone
is.moving
0
60
120
180
240
300
seconds

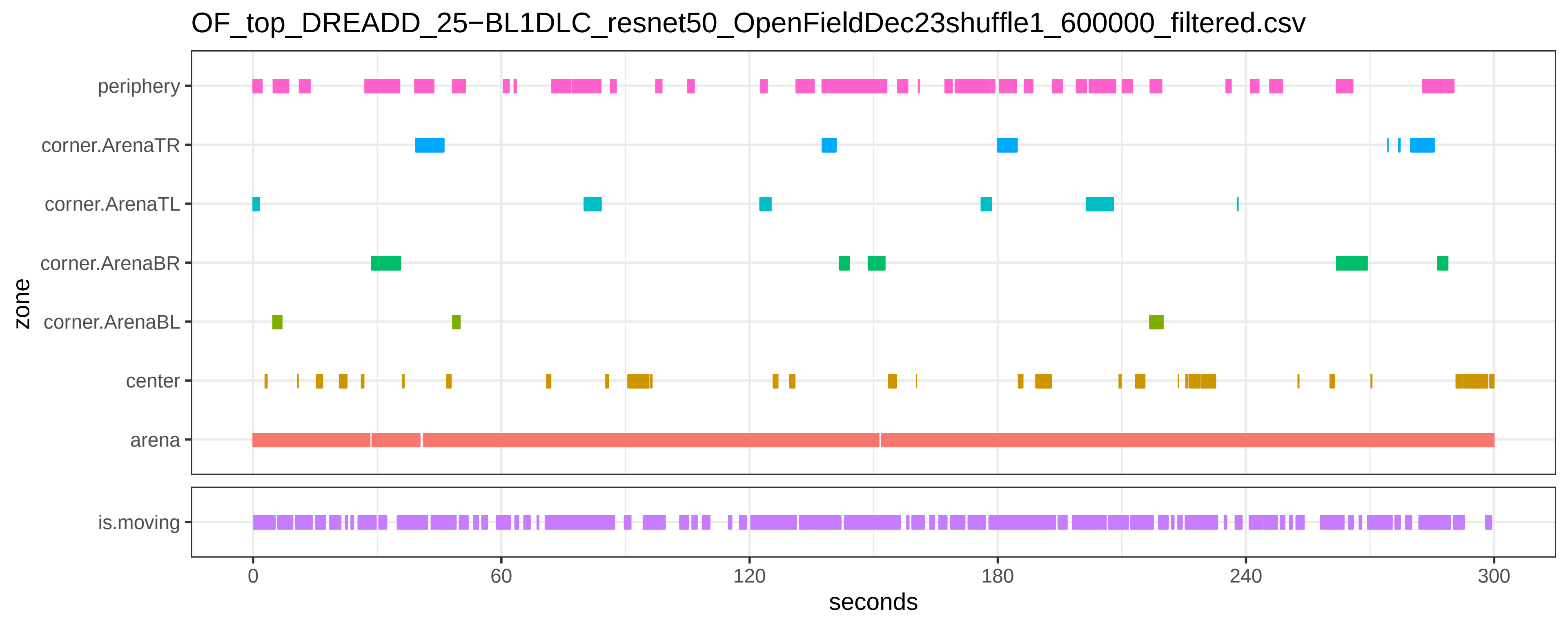
OF_top_DREADD_25−BL1DLC_resnet50_OpenFieldDec23shuffle1_600000_filtered.csv
periphery
corner.ArenaTR
corner.ArenaTL
corner.ArenaBR
corner.ArenaBL
center
arena
zone
is.moving
0
60
120
180
240
300
seconds

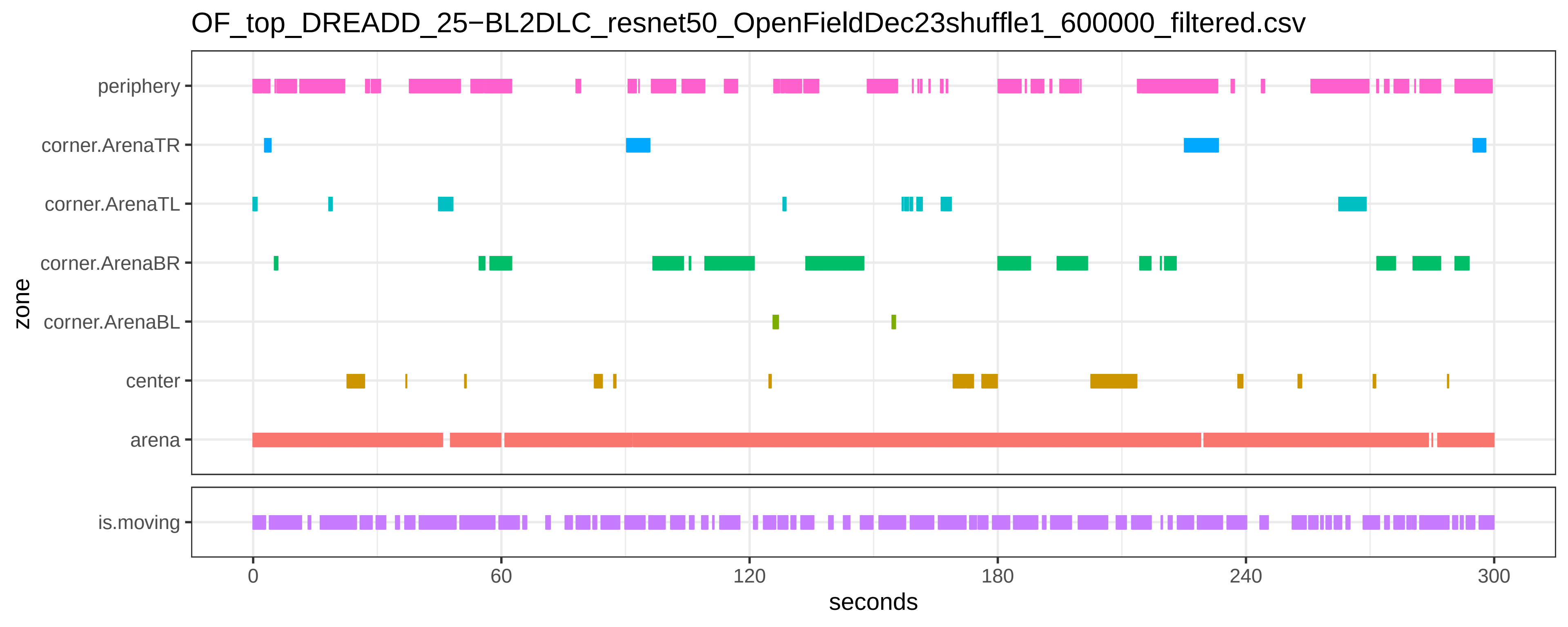

OF_top_DREADD_25−BL2DLC_resnet50_OpenFieldDec23shuffle1_600000_filtered.csv
periphery
corner.ArenaTR
corner.ArenaTL
corner.ArenaBR
corner.ArenaBL
center
arena
zone
is.moving
0
60
120
180
240
300
seconds

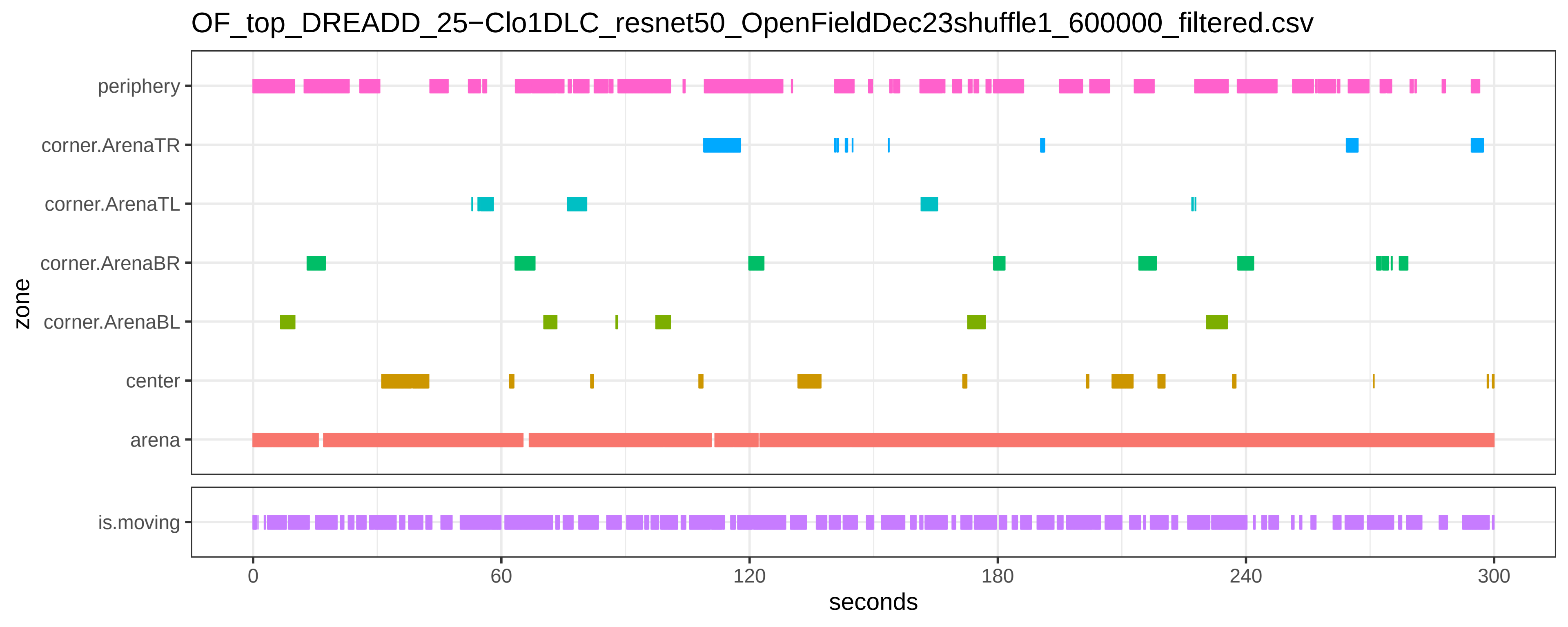
OF_top_DREADD_25−Clo1DLC_resnet50_OpenFieldDec23shuffle1_600000_filtered.csv
zone
periphery
corner.ArenaTR
corner.ArenaTL
corner.ArenaBR
corner.ArenaBL
center
arena
is.moving
0
60
120
180
240
300
seconds

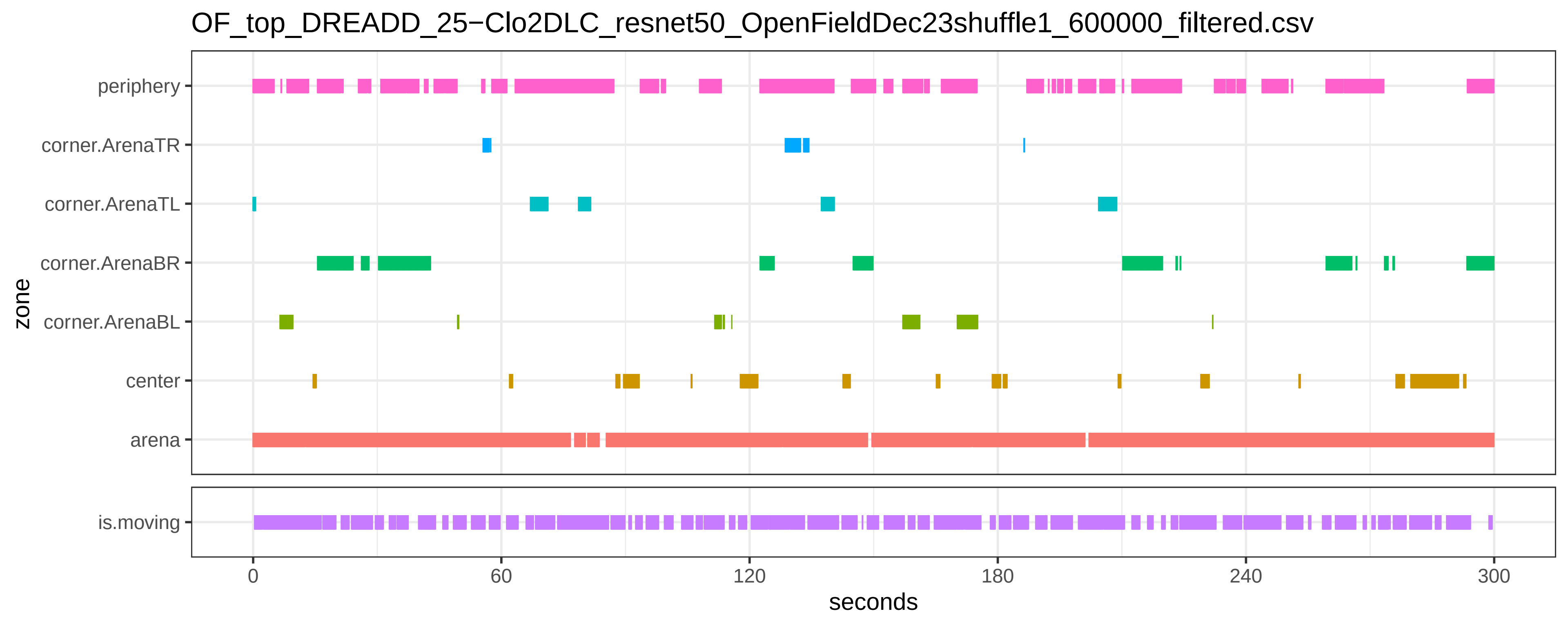
OF_top_DREADD_25−Clo2DLC_resnet50_OpenFieldDec23shuffle1_600000_filtered.csv
periphery
corner.ArenaTR
corner.ArenaTL
corner.ArenaBR
corner.ArenaBL
center
arena
zone
is.moving
0
60
120
180
240
300
seconds

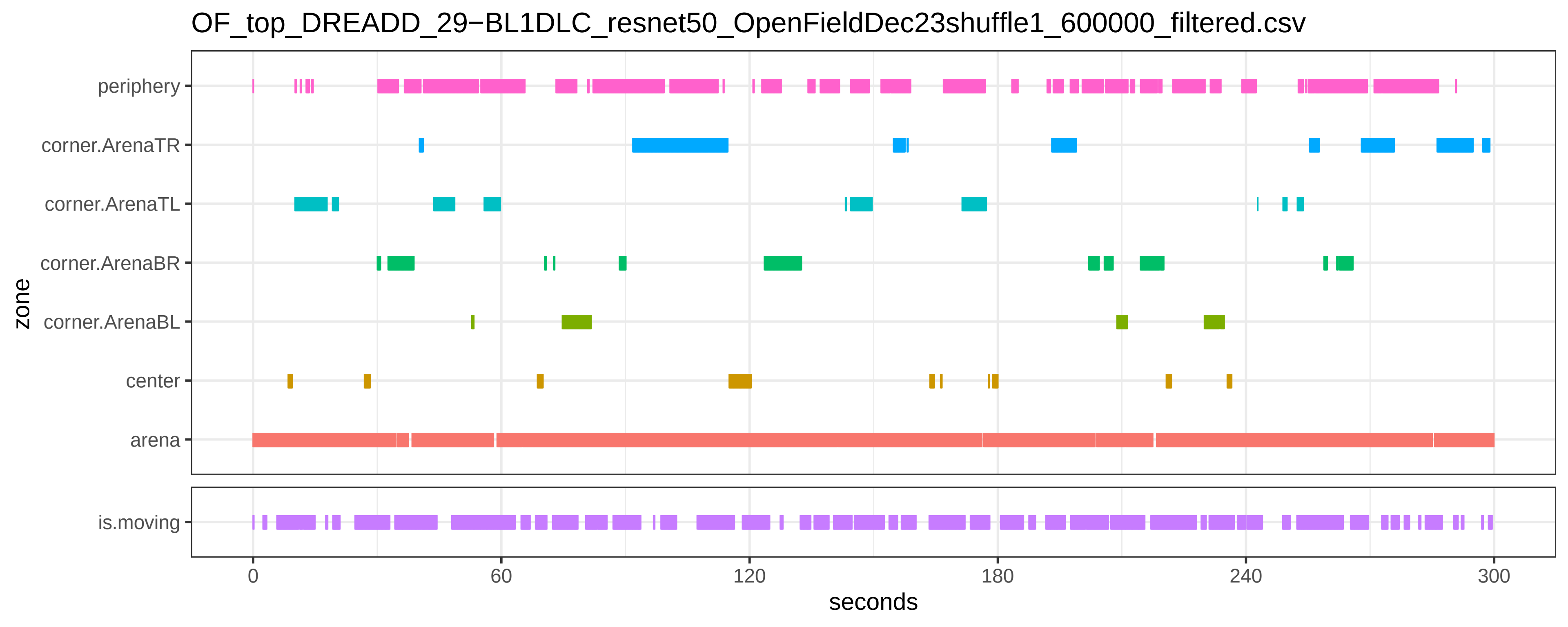
OF_top_DREADD_29−BL1DLC_resnet50_OpenFieldDec23shuffle1_600000_filtered.csv
periphery
corner.ArenaTR
corner.ArenaTL
corner.ArenaBR
corner.ArenaBL
center
arena
zone
is.moving
0
60
120
180
240
300
seconds

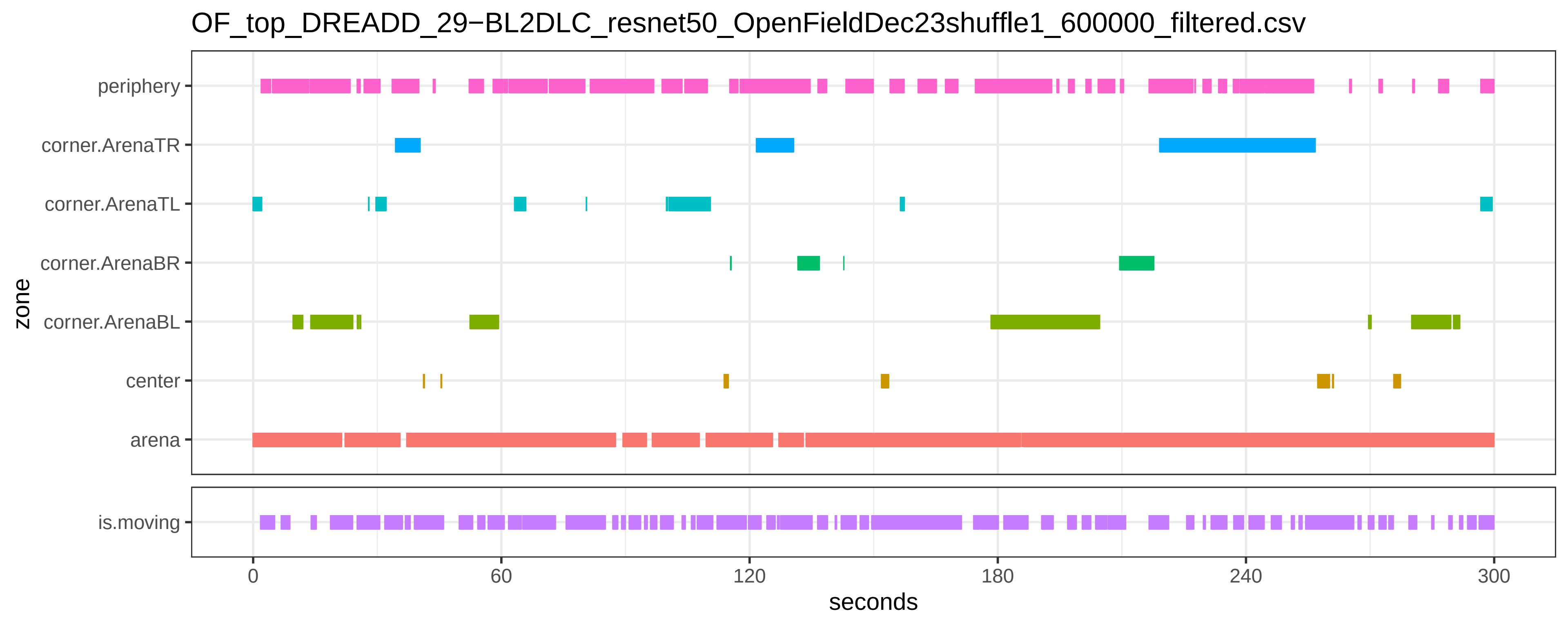

OF_top_DREADD_29−BL2DLC_resnet50_OpenFieldDec23shuffle1_600000_filtered.csv
periphery
corner.ArenaTR
corner.ArenaTL
corner.ArenaBR
corner.ArenaBL
center
arena
is.moving
zone
0
60
120
180
240
300
seconds

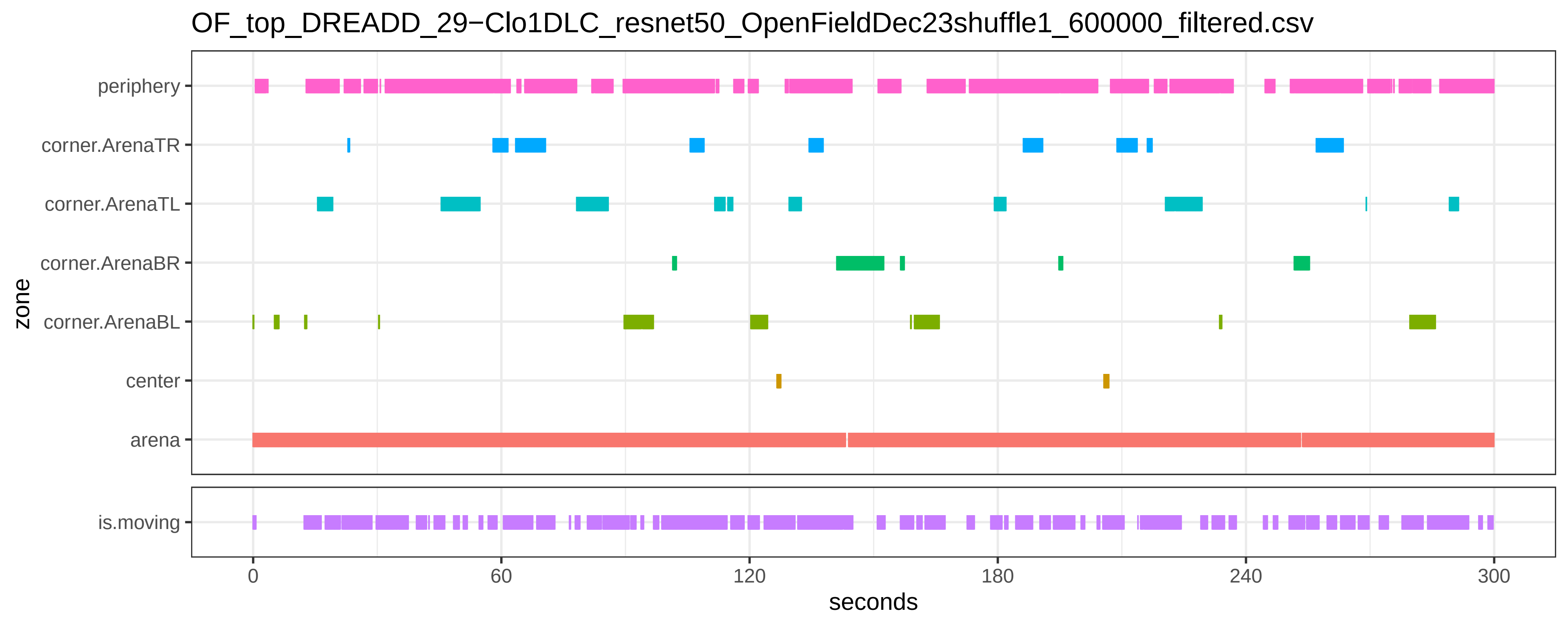

OF_top_DREADD_29−Clo1DLC_resnet50_OpenFieldDec23shuffle1_600000_filtered.csv
periphery
corner.ArenaTR
corner.ArenaTL
corner.ArenaBR
corner.ArenaBL
center
arena
zone
is.moving
0
60
120
180
240
300
seconds

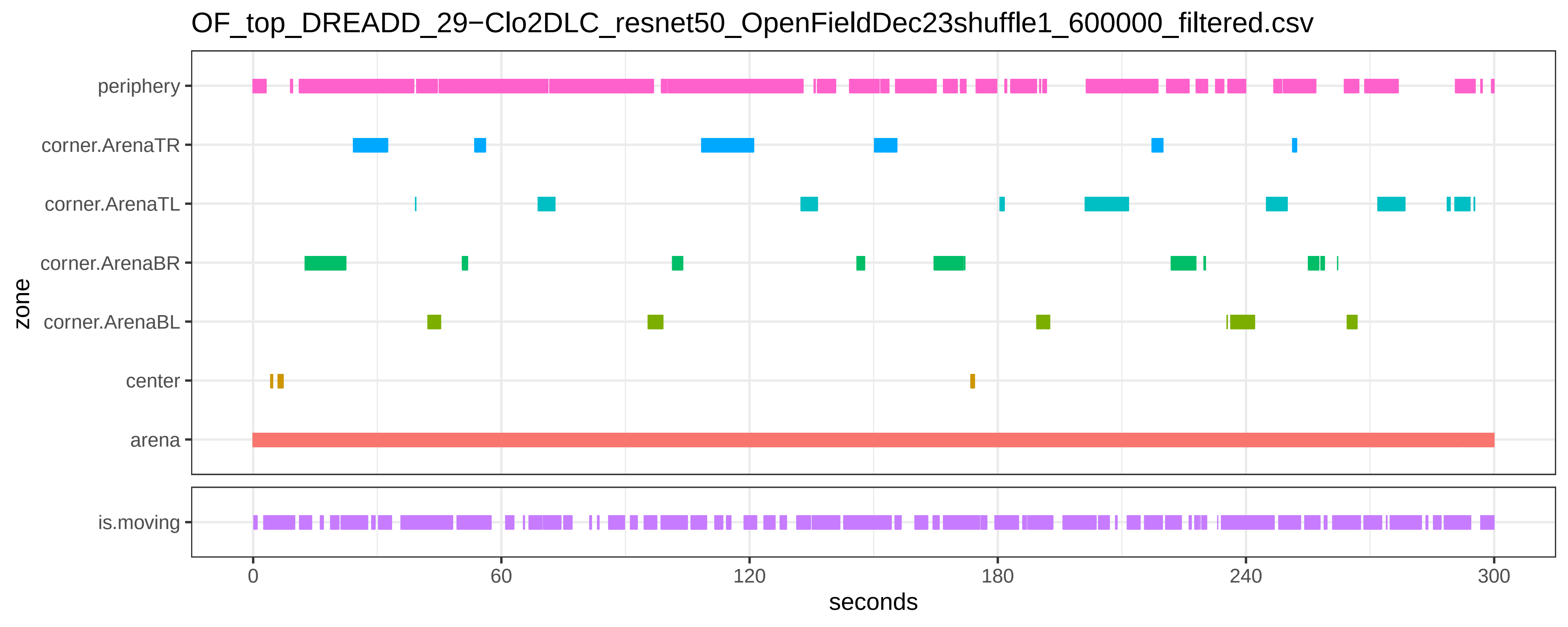

OF_top_DREADD_29−Clo2DLC_resnet50_OpenFieldDec23shuffle1_600000_filtered.csv
zone
periphery
corner.ArenaTR
corner.ArenaTL
corner.ArenaBR
corner.ArenaBL
center
arena
is.moving
0
60
120
180
240
300
seconds

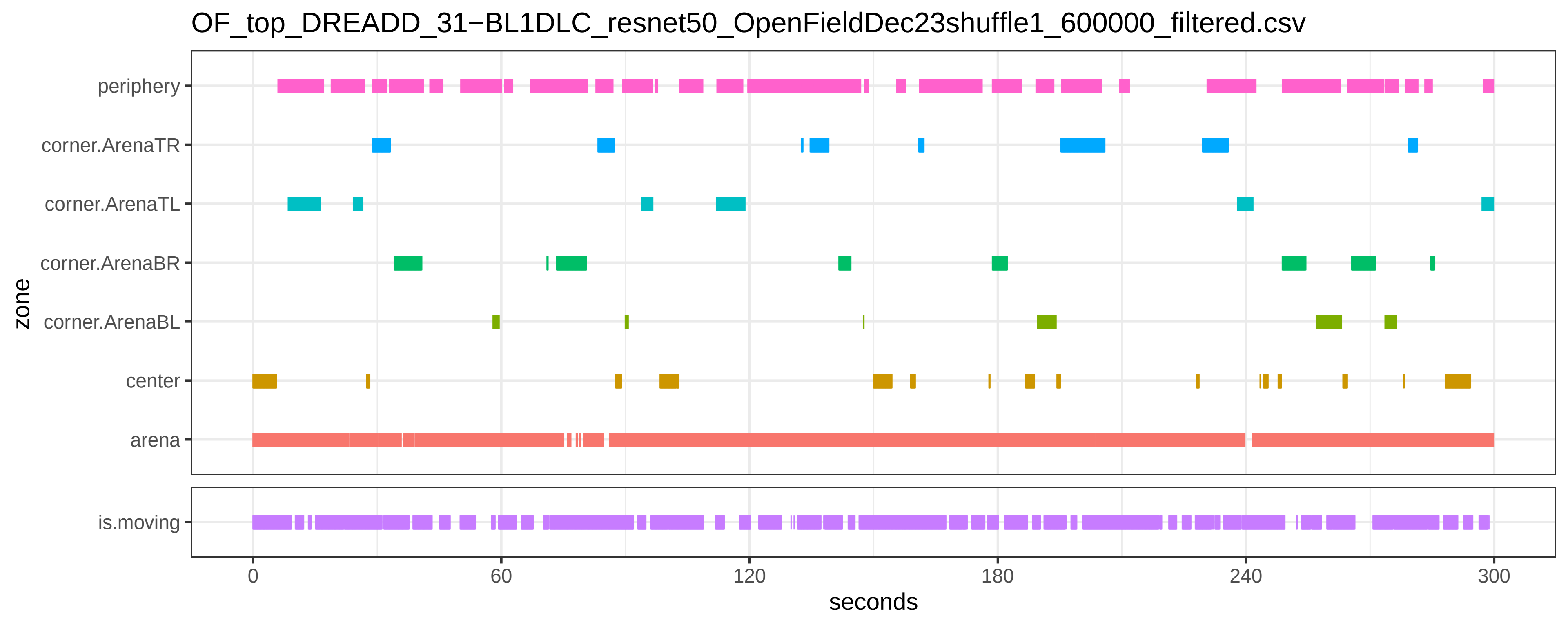
OF_top_DREADD_31−BL1DLC_resnet50_OpenFieldDec23shuffle1_600000_filtered.csv
periphery
corner.ArenaTR
corner.ArenaTL
corner.ArenaBR
corner.ArenaBL
center
arena
zone
is.moving
0
60
120
180
240
300
seconds

OF_top_DREADD_31−BL2DLC_resnet50_OpenFieldDec23shuffle1_600000_filtered.csv

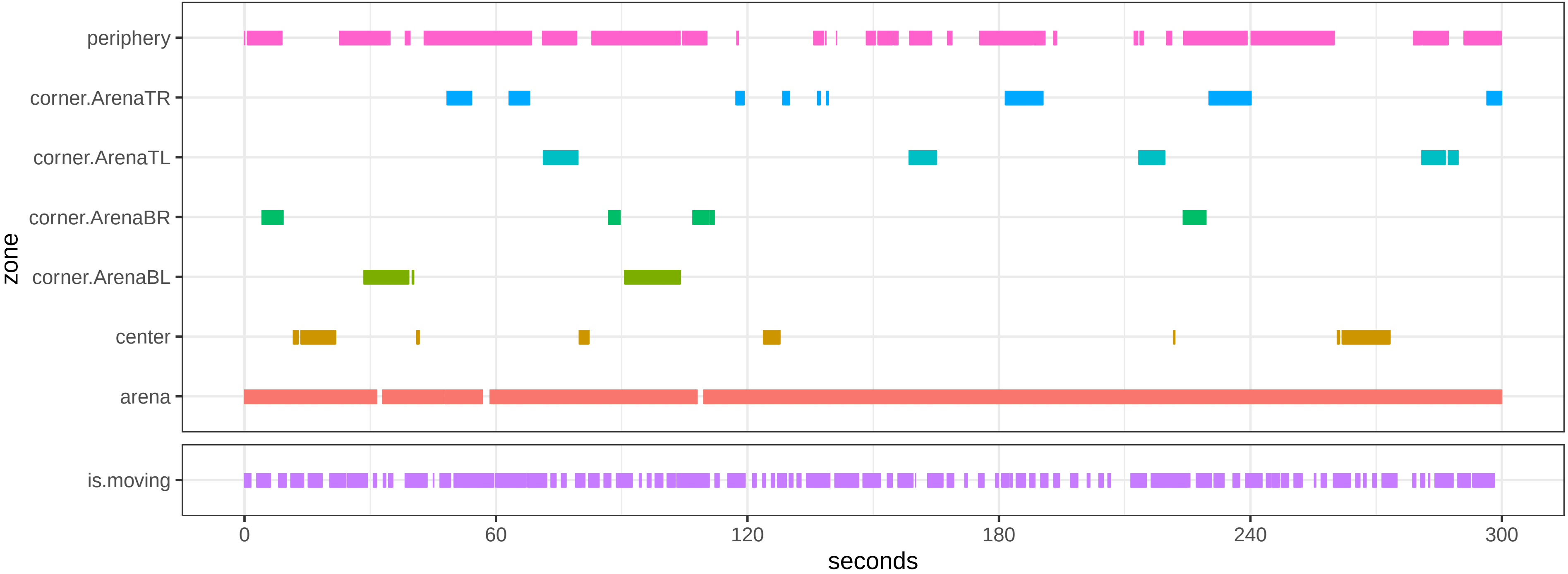

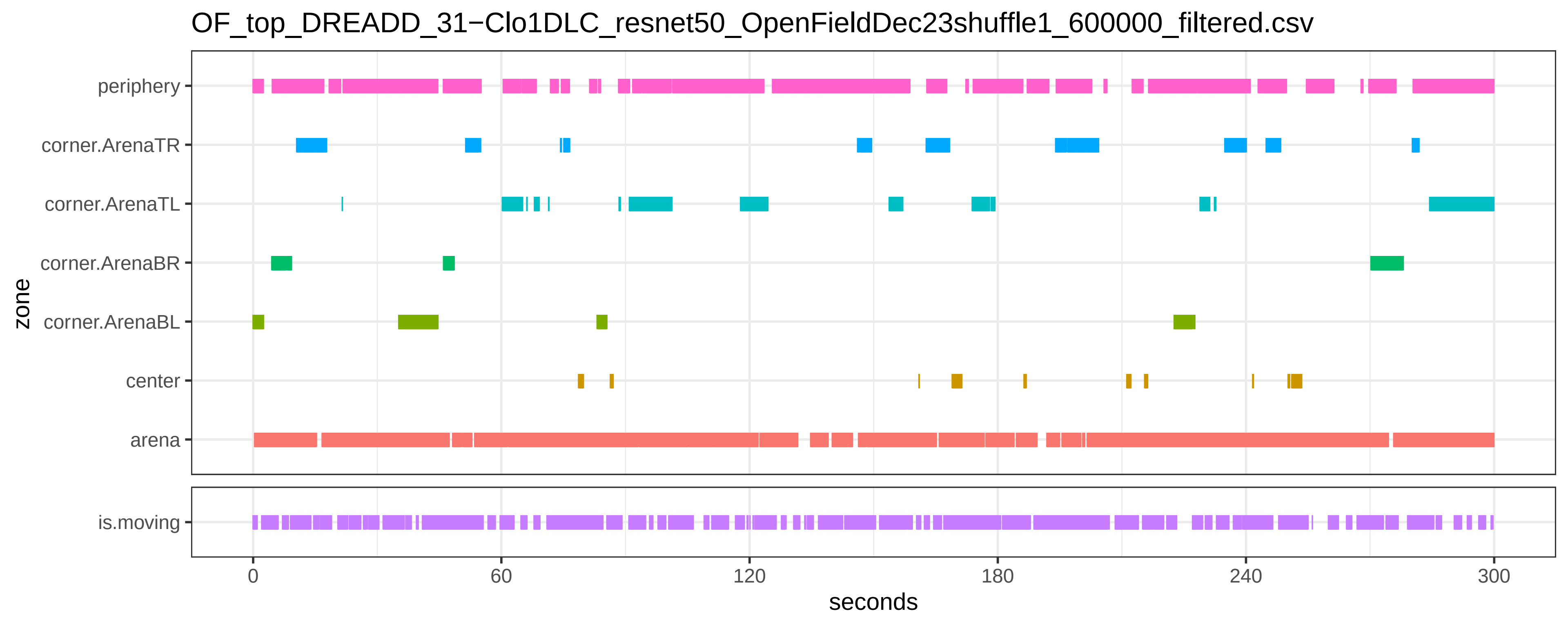
OF_top_DREADD_31−Clo1DLC_resnet50_OpenFieldDec23shuffle1_600000_filtered.csv
periphery
corner.ArenaTR
corner.ArenaTL
corner.ArenaBR
corner.ArenaBL
center
arena
zone
is.moving
0
60
120
180
240
300
seconds

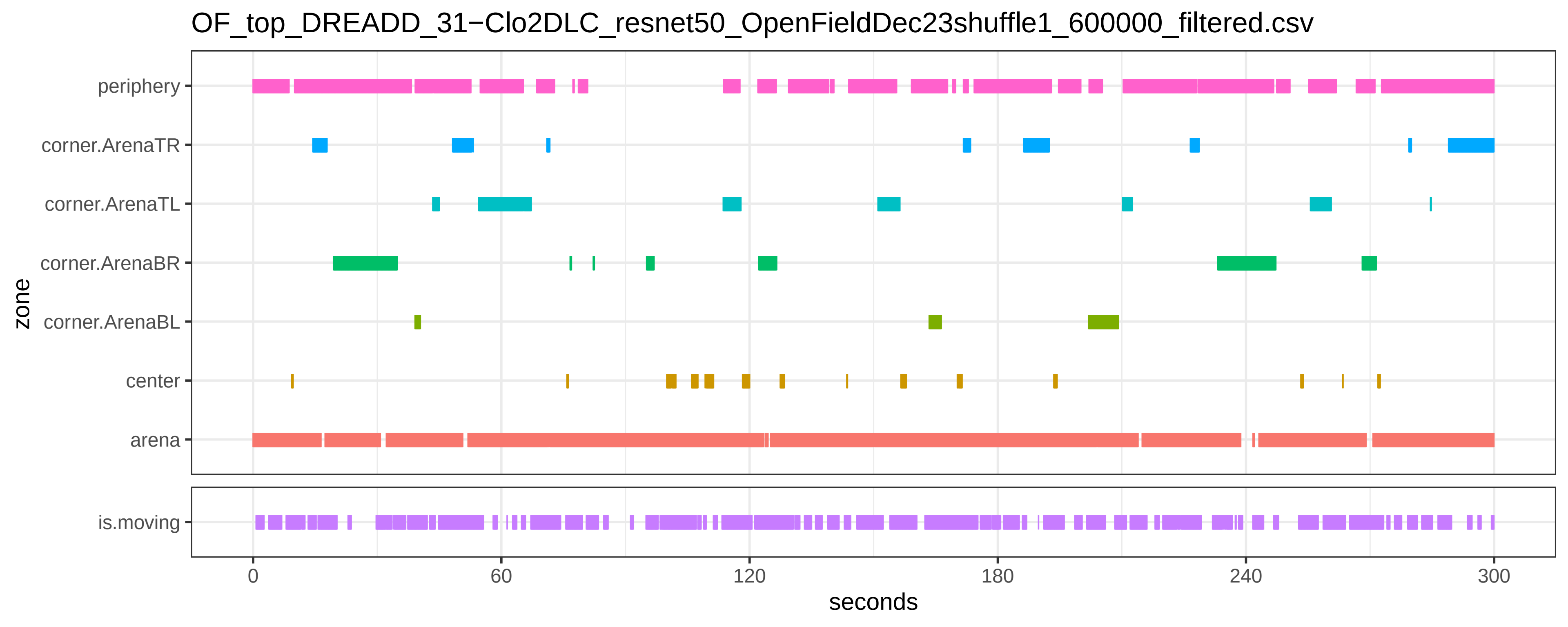

OF_top_DREADD_31−Clo2DLC_resnet50_OpenFieldDec23shuffle1_600000_filtered.csv
zone
periphery
corner.ArenaTR
corner.ArenaTL
corner.ArenaBR
corner.ArenaBL
center
arena
is.moving
0
60
120
180
240
300
seconds

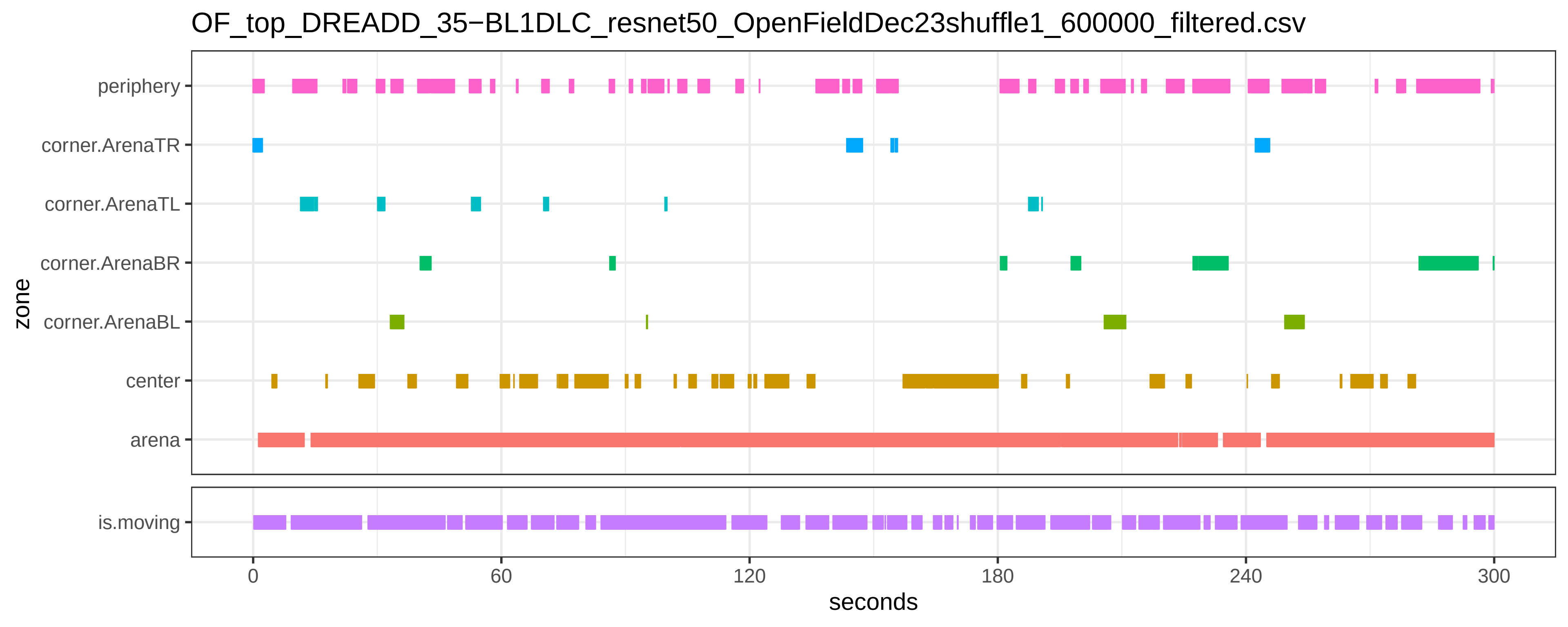

OF_top_DREADD_35−BL1DLC_resnet50_OpenFieldDec23shuffle1_600000_filtered.csv
periphery
corner.ArenaTR
corner.ArenaTL
corner.ArenaBR
corner.ArenaBL
center
arena
zone
is.moving
0
60
120
180
240
300
seconds

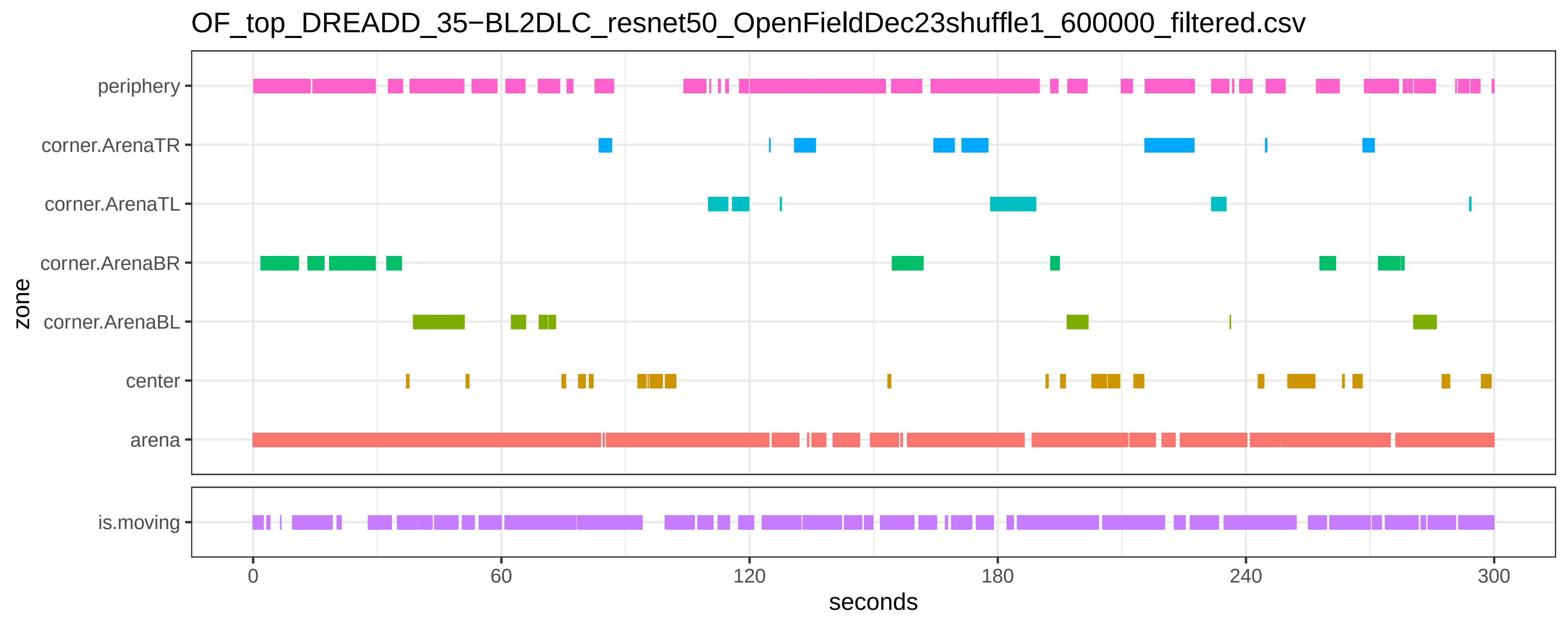

OF_top_DREADD_35−BL2DLC_resnet50_OpenFieldDec23shuffle1_600000_filtered.csv
periphery
corner.ArenaTR
corner.ArenaTL
corner.ArenaBR
corner.ArenaBL
center
arena
zone
is.moving
0
60
120
180
240
300
seconds

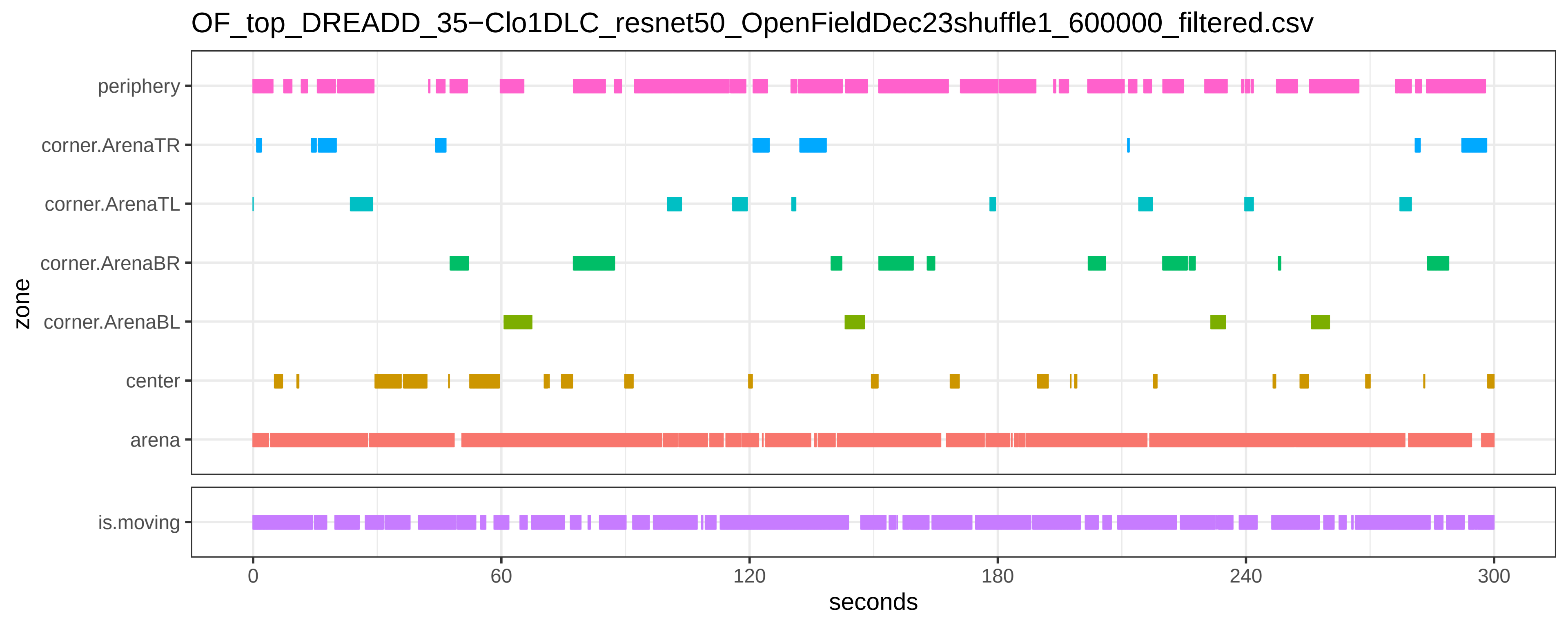

OF_top_DREADD_35−Clo1DLC_resnet50_OpenFieldDec23shuffle1_600000_filtered.csv
periphery
corner.ArenaTR
corner.ArenaTL
corner.ArenaBR
corner.ArenaBL
center
arena
zone
is.moving
0
60
120
180
240
300
seconds

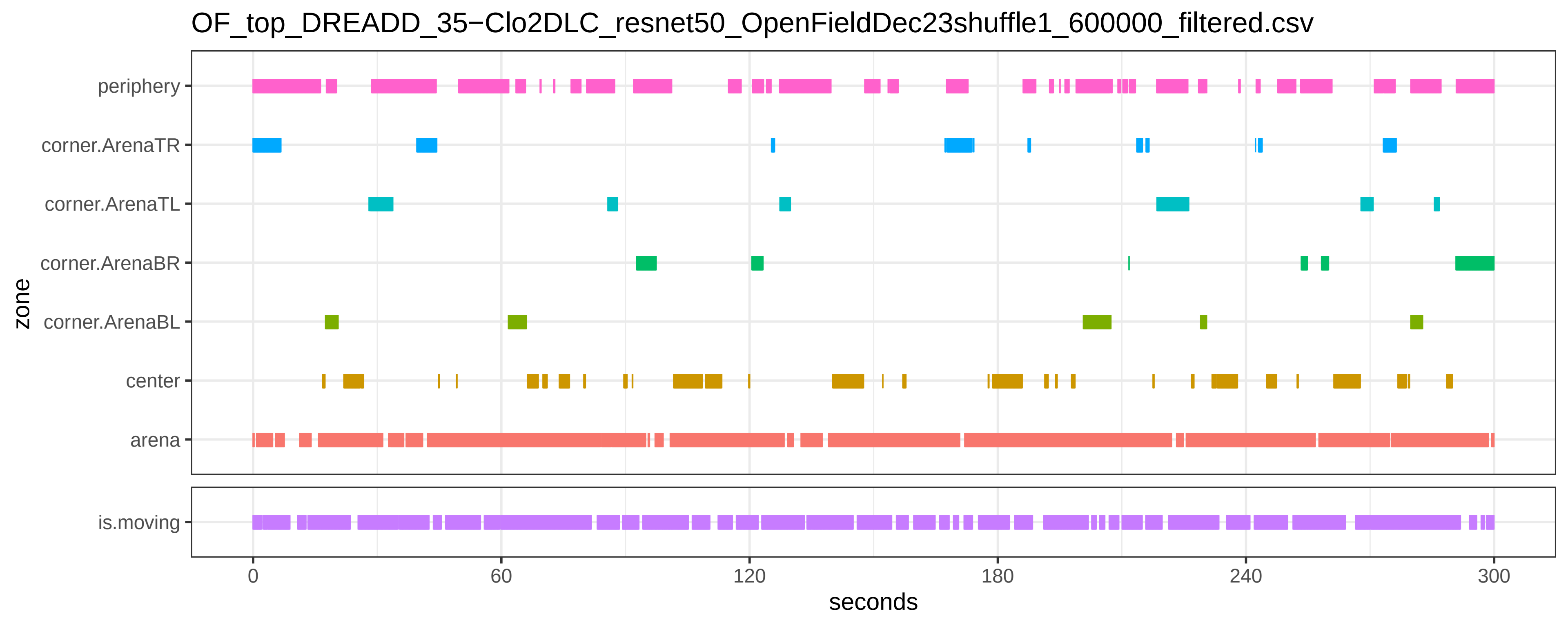
OF_top_DREADD_35−Clo2DLC_resnet50_OpenFieldDec23shuffle1_600000_filtered.csv
periphery
corner.ArenaTR
corner.ArenaTL
corner.ArenaBR
corner.ArenaBL
center
arena
zone
is.moving
0
60
120
180
240
300
seconds

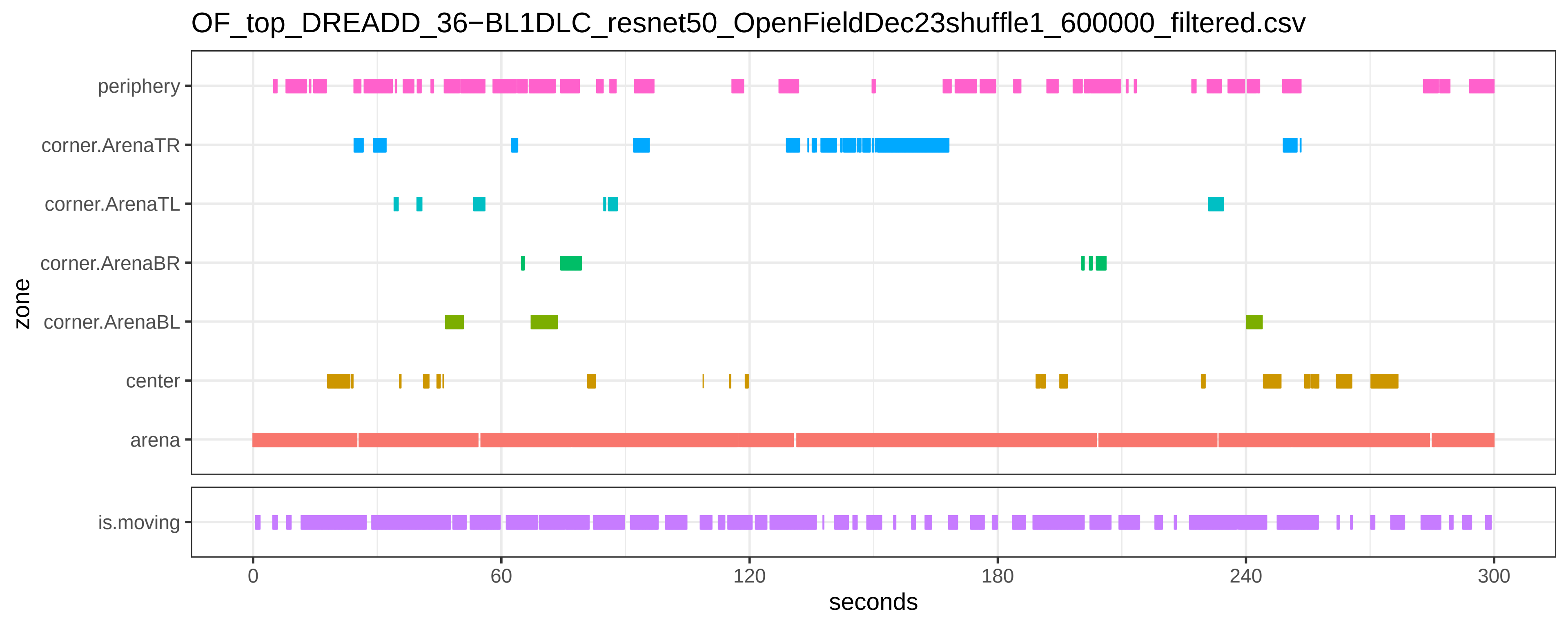
OF_top_DREADD_36−BL1DLC_resnet50_OpenFieldDec23shuffle1_600000_filtered.csv
periphery
corner.ArenaTR
corner.ArenaTL
corner.ArenaBR
corner.ArenaBL
center
arena
zone
is.moving
0
60
120
180
240
300
seconds

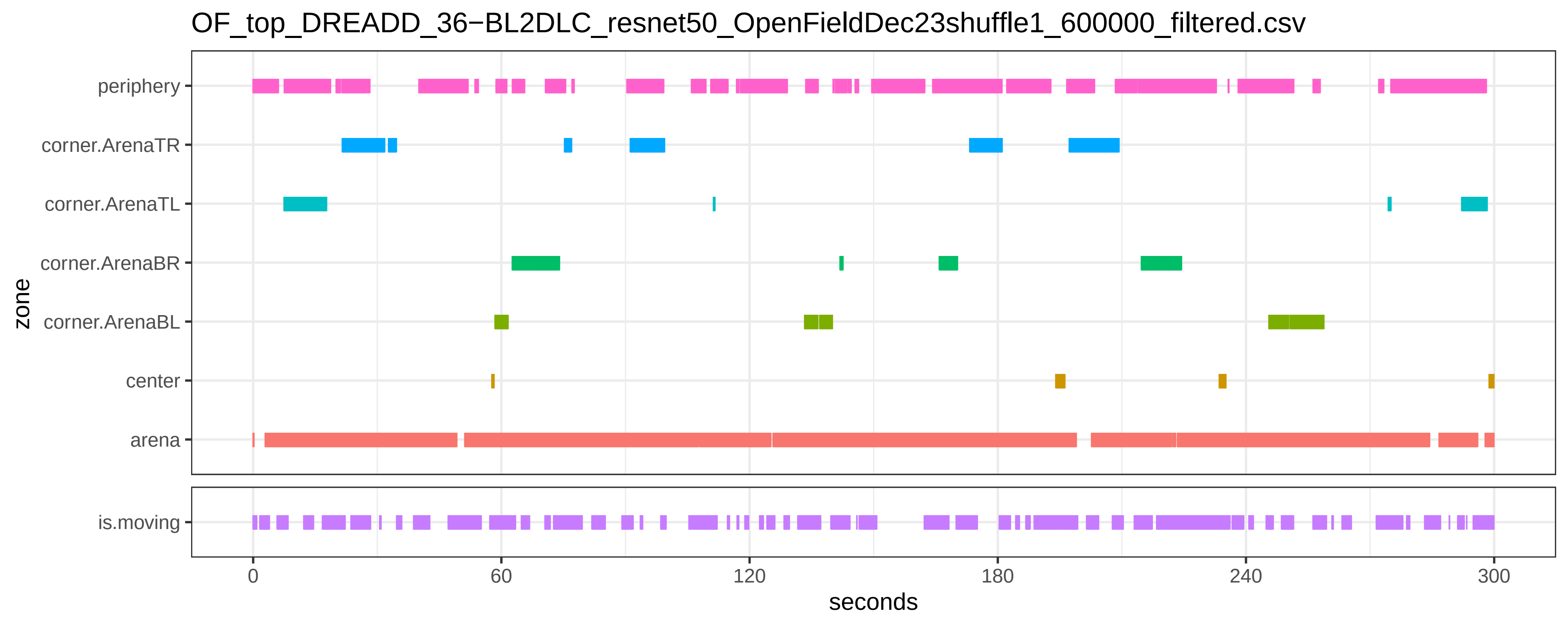

OF_top_DREADD_36−BL2DLC_resnet50_OpenFieldDec23shuffle1_600000_filtered.csv
periphery
corner.ArenaTR
corner.ArenaTL
corner.ArenaBR
corner.ArenaBL
center
arena
zone
is.moving
0
60
120
180
240
300
seconds

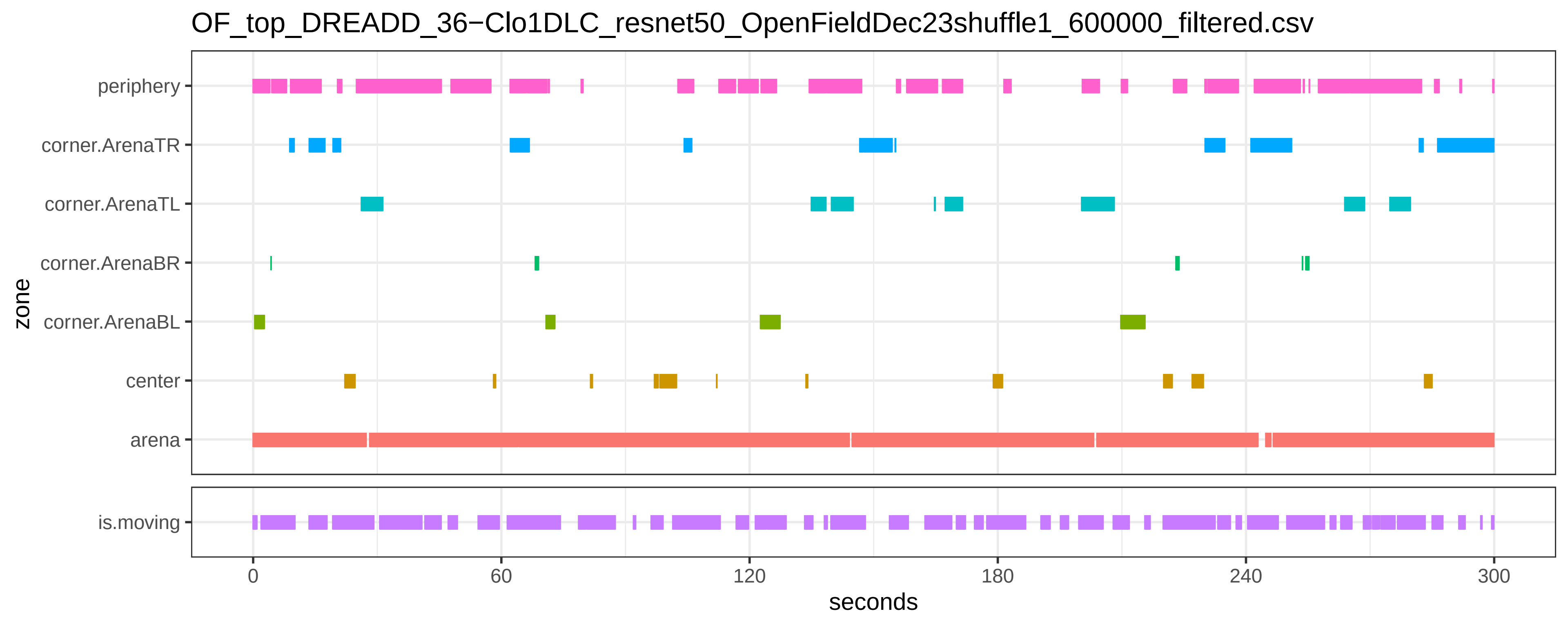
OF_top_DREADD_36−Clo1DLC_resnet50_OpenFieldDec23shuffle1_600000_filtered.csv
zone
periphery
corner.ArenaTR
corner.ArenaTL
corner.ArenaBR
corner.ArenaBL
center
arena
is.moving
0
60
120
180
240
300
seconds

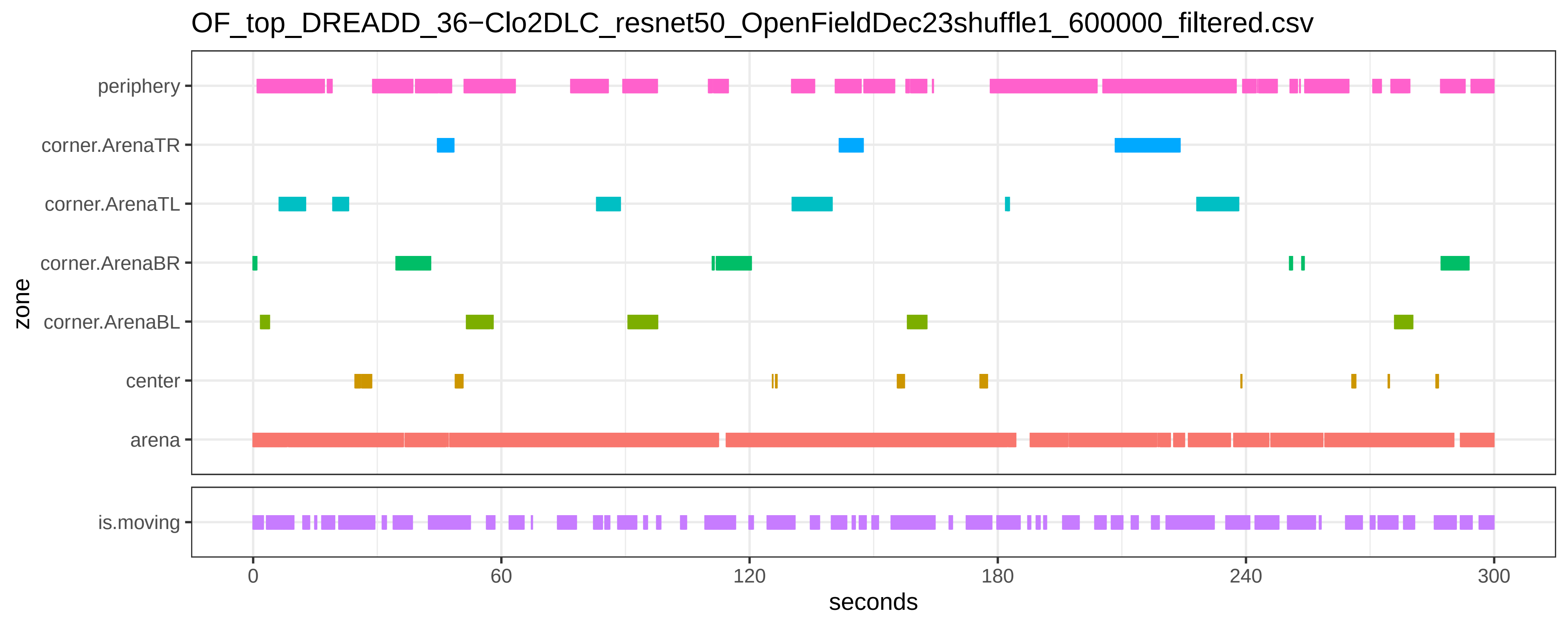

OF_top_DREADD_36−Clo2DLC_resnet50_OpenFieldDec23shuffle1_600000_filtered.csv
periphery
corner.ArenaTR
corner.ArenaTL
corner.ArenaBR
corner.ArenaBL
center
arena
zone
is.moving
0
60
120
180
240
300
seconds

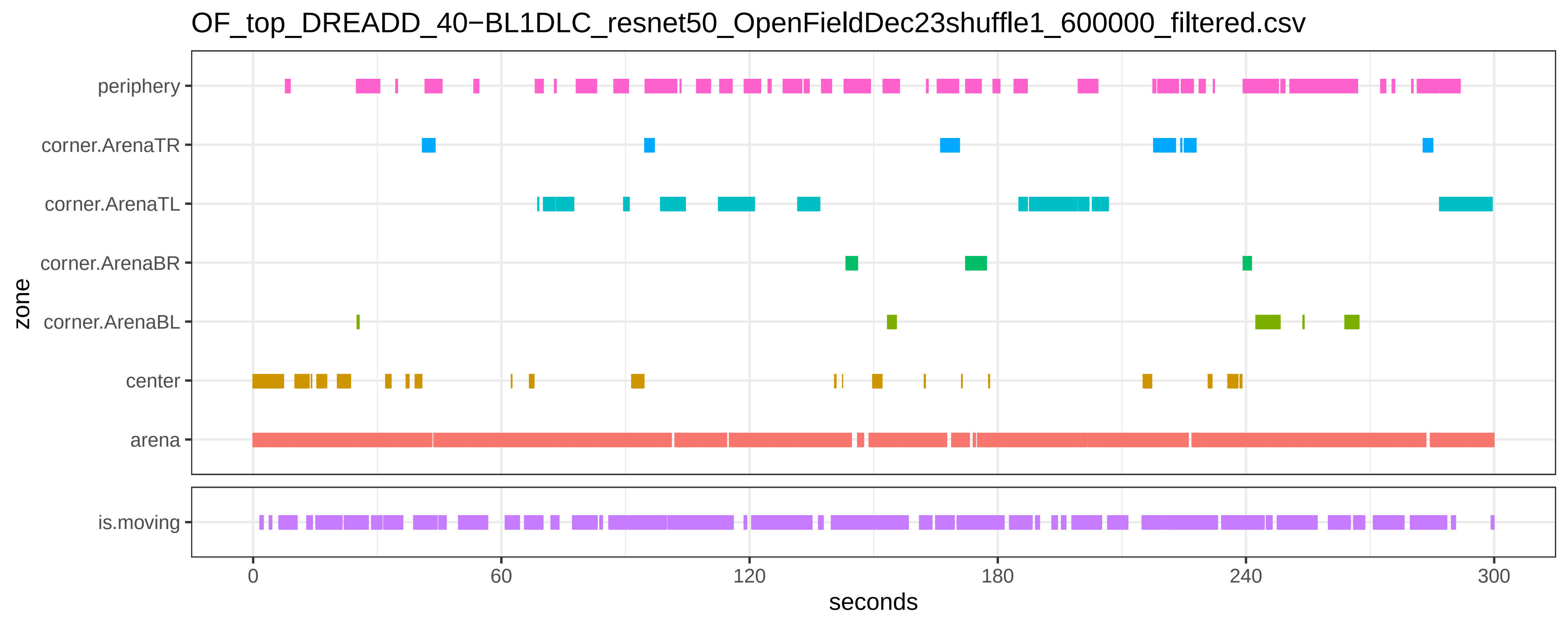

OF_top_DREADD_40−BL1DLC_resnet50_OpenFieldDec23shuffle1_600000_filtered.csv
periphery
corner.ArenaTR
corner.ArenaTL
corner.ArenaBR
corner.ArenaBL
center
arena
zone
is.moving
0
60
120
180
240
300
seconds

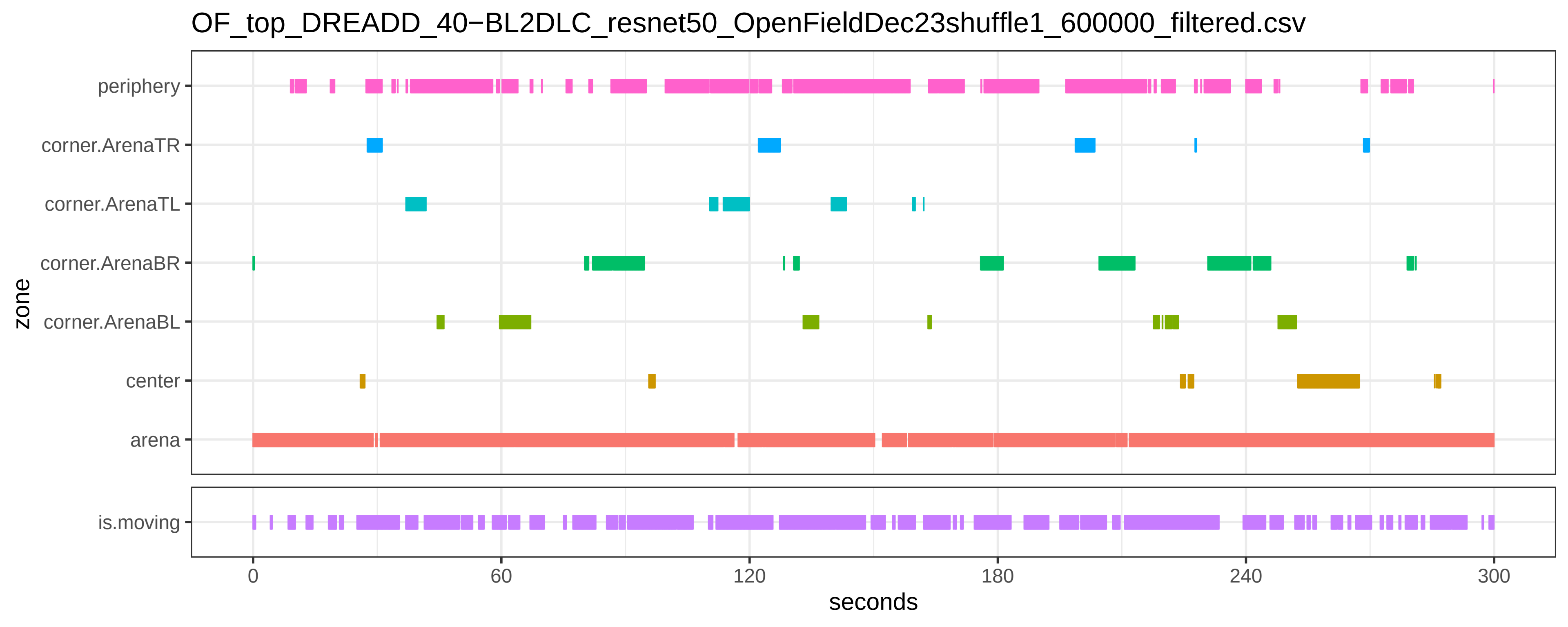
OF_top_DREADD_40−BL2DLC_resnet50_OpenFieldDec23shuffle1_600000_filtered.csv
periphery
corner.ArenaTR
corner.ArenaTL
corner.ArenaBR
corner.ArenaBL
center
arena
zone
is.moving
0
60
120
180
240
300
seconds

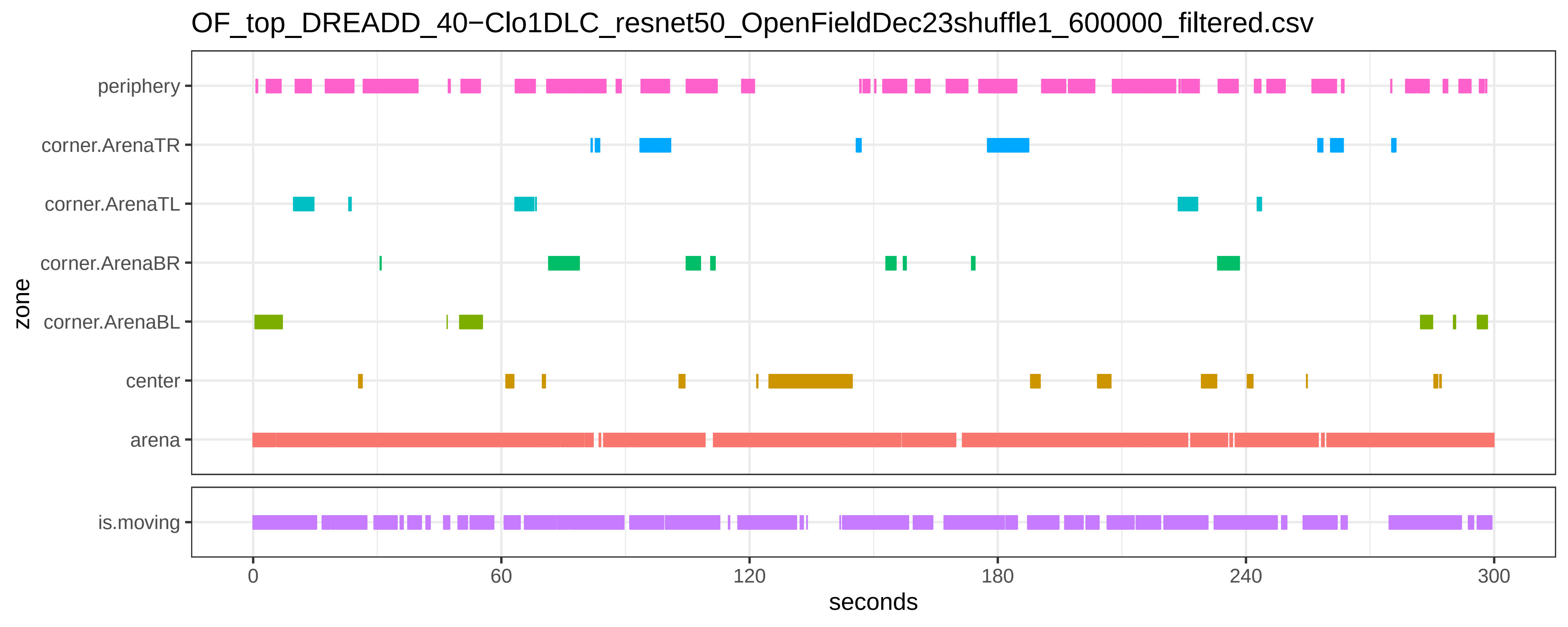
OF_top_DREADD_40−Clo1DLC_resnet50_OpenFieldDec23shuffle1_600000_filtered.csv
periphery
corner.ArenaTR
corner.ArenaTL
corner.ArenaBR
corner.ArenaBL
center
arena
zone
is.moving
0
60
120
180
240
300
seconds

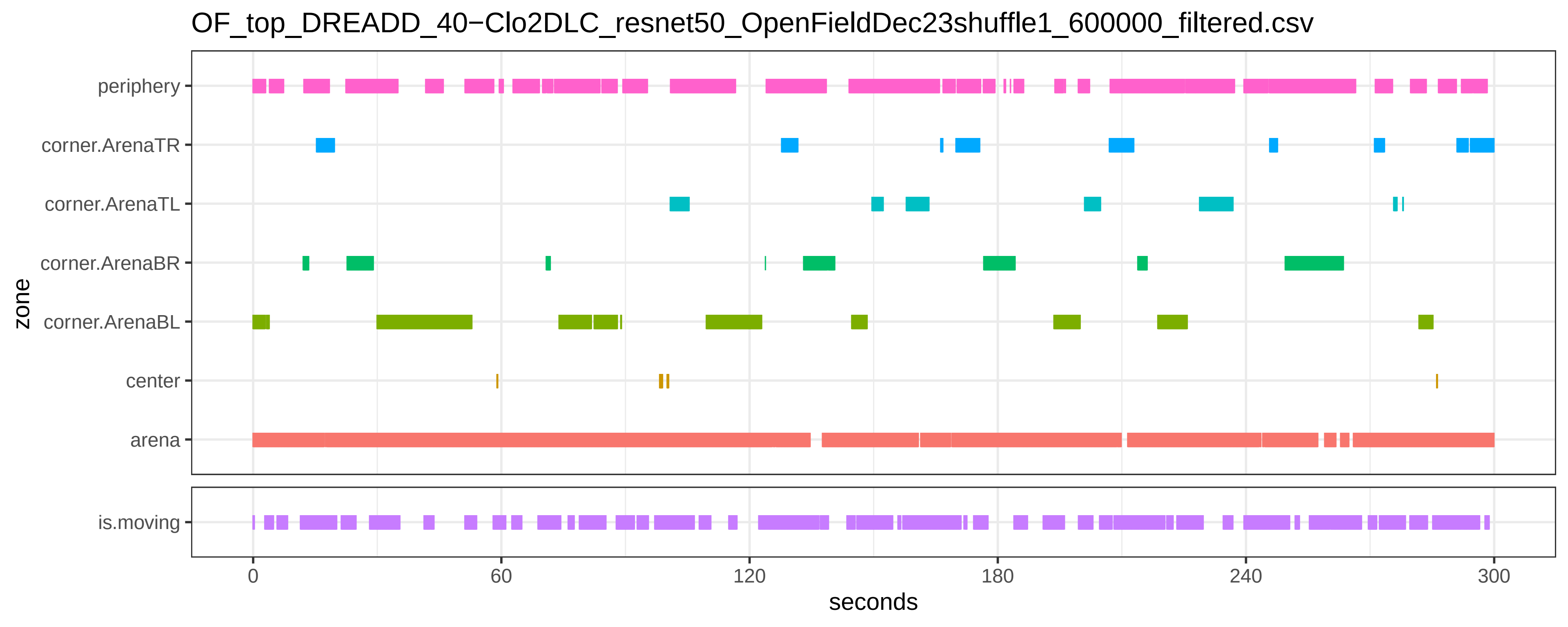
OF_top_DREADD_40−Clo2DLC_resnet50_OpenFieldDec23shuffle1_600000_filtered.csv
periphery
corner.ArenaTR
corner.ArenaTL
corner.ArenaBR
corner.ArenaBL
center
arena
zone
is.moving
0
60
120
180
240
300
seconds

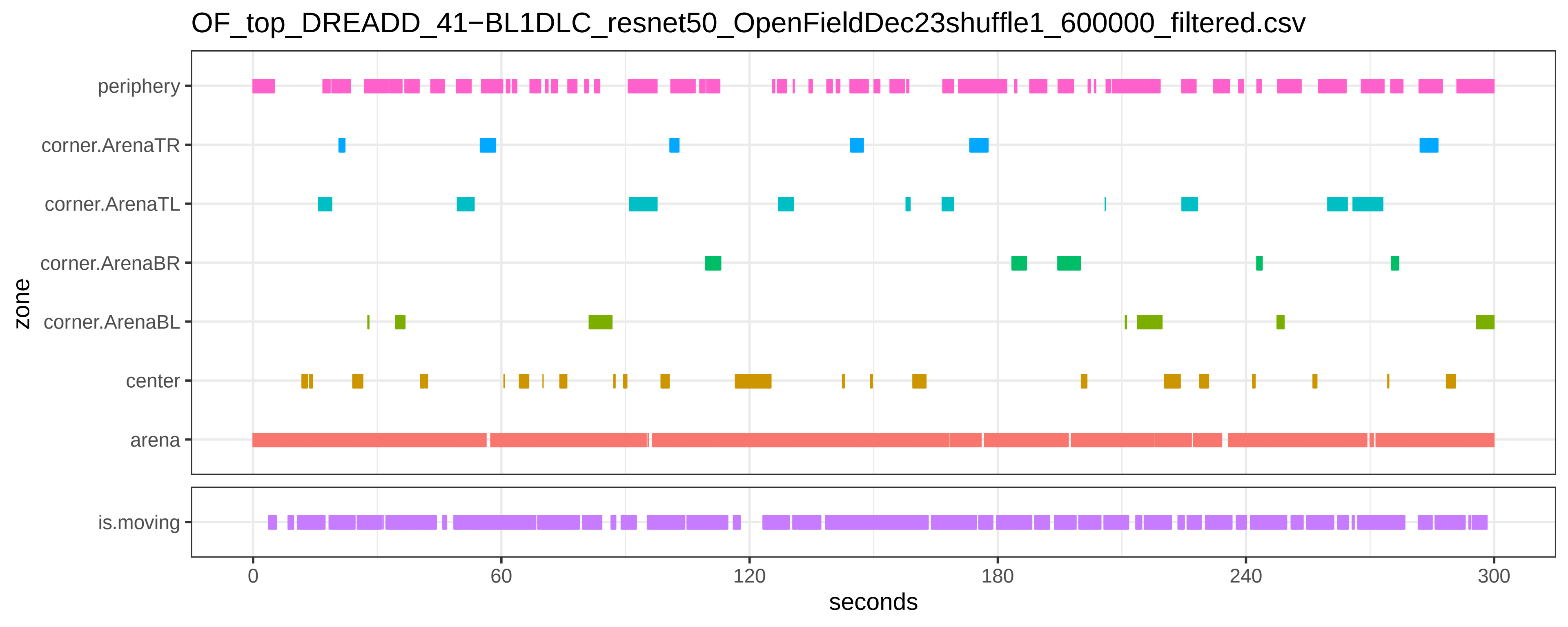

OF_top_DREADD_41−BL1DLC_resnet50_OpenFieldDec23shuffle1_600000_filtered.csv
periphery
corner.ArenaTR
corner.ArenaTL
corner.ArenaBR
corner.ArenaBL
center
arena
zone
is.moving
0
60
120
180
240
300
seconds

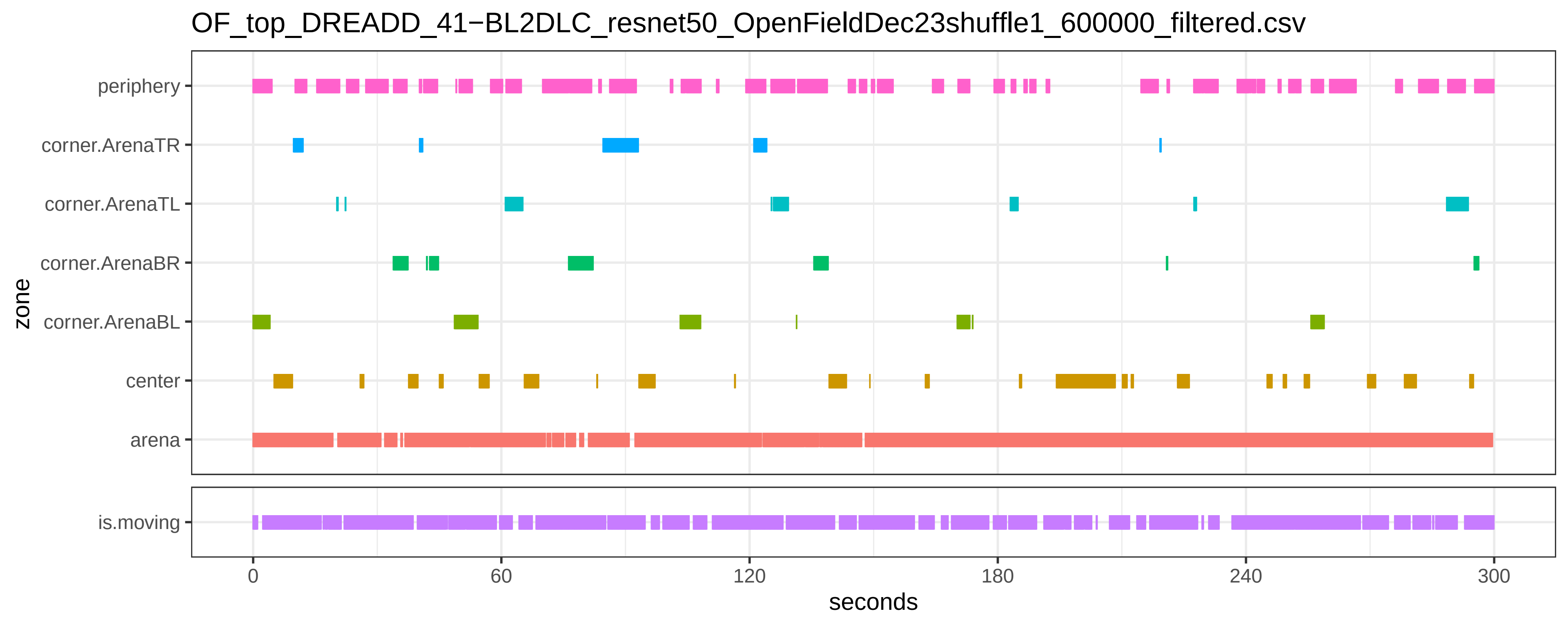
OF_top_DREADD_41−BL2DLC_resnet50_OpenFieldDec23shuffle1_600000_filtered.csv
zone
periphery
corner.ArenaTR
corner.ArenaTL
corner.ArenaBR
corner.ArenaBL
center
arena
is.moving
0
60
120
180
240
300
seconds

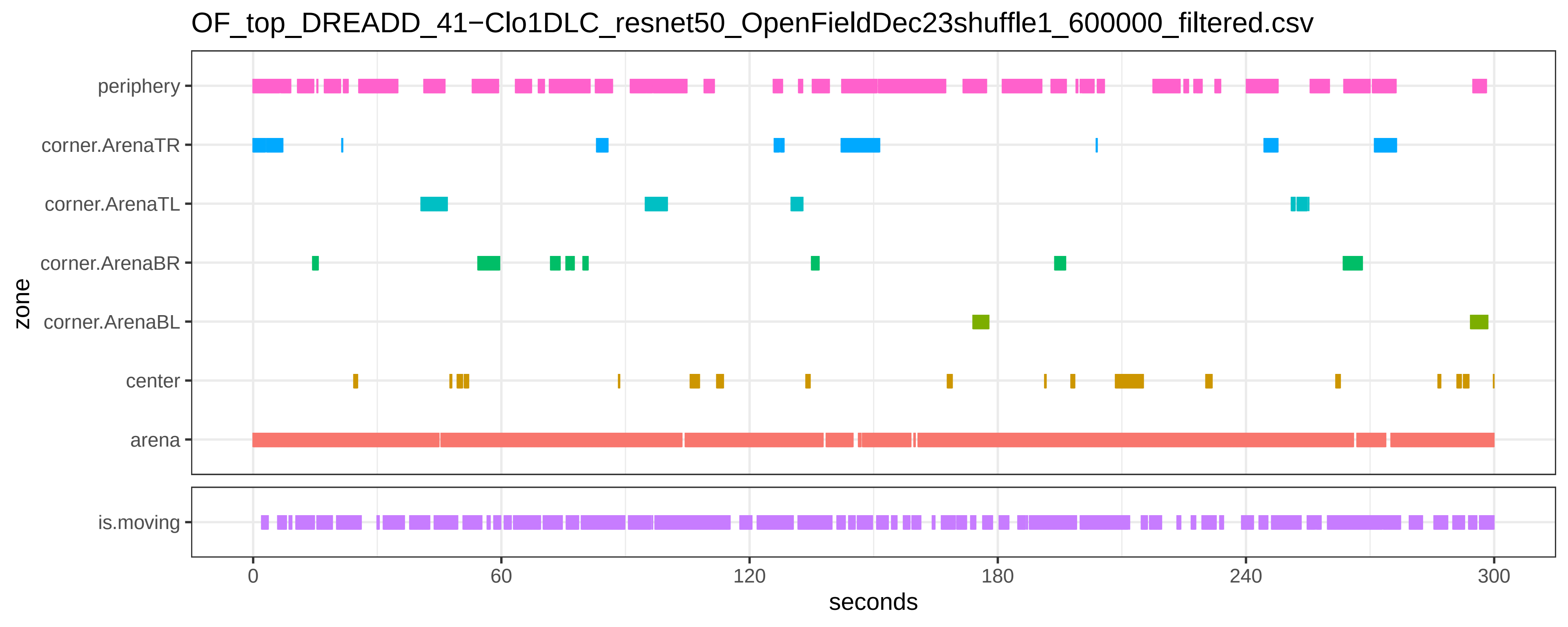

OF_top_DREADD_41−Clo1DLC_resnet50_OpenFieldDec23shuffle1_600000_filtered.csv
periphery
corner.ArenaTR
corner.ArenaTL
corner.ArenaBR
corner.ArenaBL
center
arena
is.moving
zone
0
60
120
180
240
300
seconds

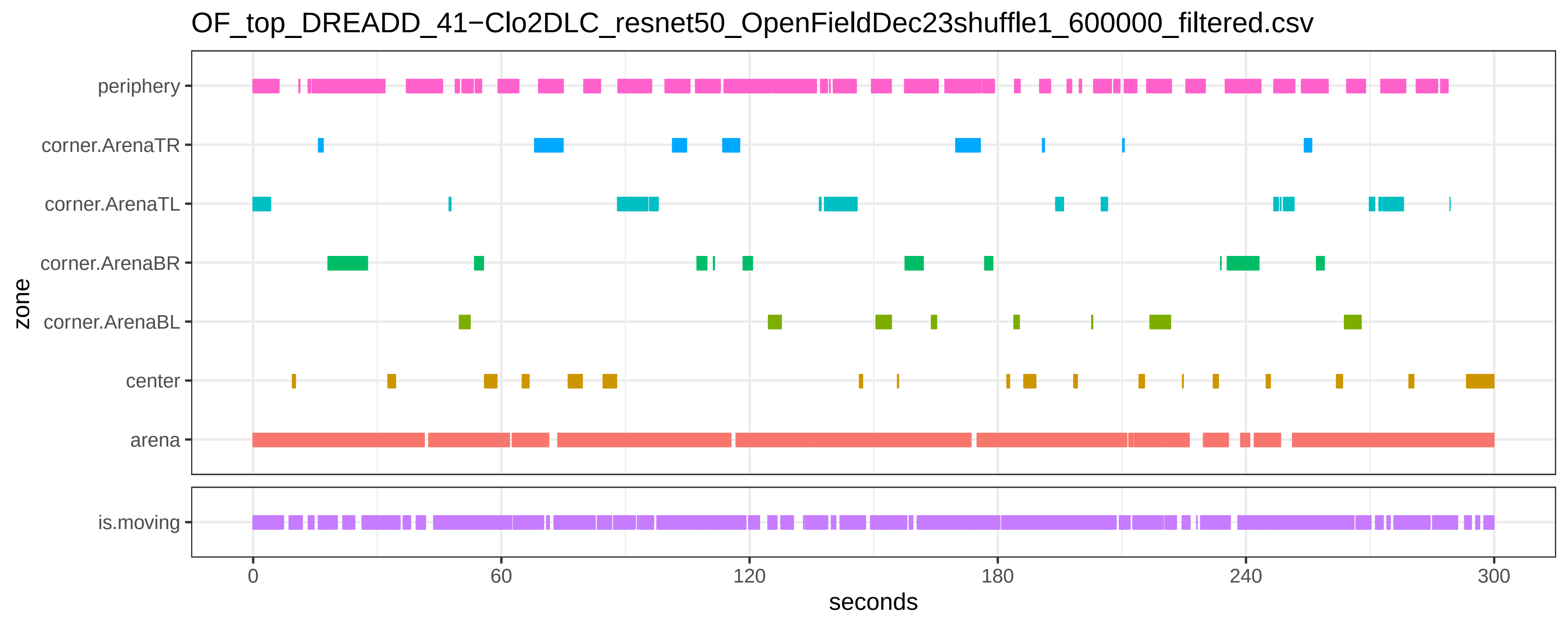

OF_top_DREADD_41−Clo2DLC_resnet50_OpenFieldDec23shuffle1_600000_filtered.csv
periphery
corner.ArenaTR
corner.ArenaTL
corner.ArenaBR
corner.ArenaBL
center
arena
zone
is.moving
0
60
120
180
240
300
seconds

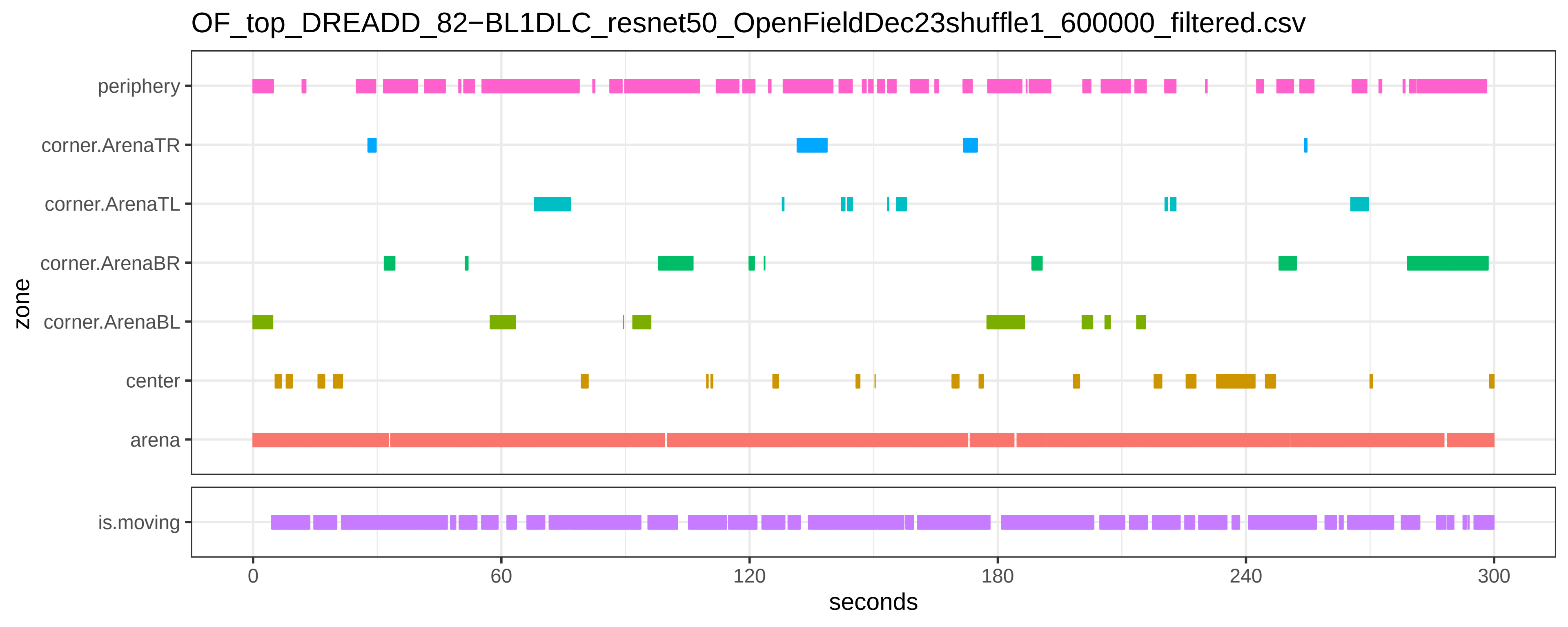
OF_top_DREADD_82−BL1DLC_resnet50_OpenFieldDec23shuffle1_600000_filtered.csv
zone
periphery
corner.ArenaTR
corner.ArenaTL
corner.ArenaBR
corner.ArenaBL
center
arena
is.moving
0
60
120
180
240
300
seconds

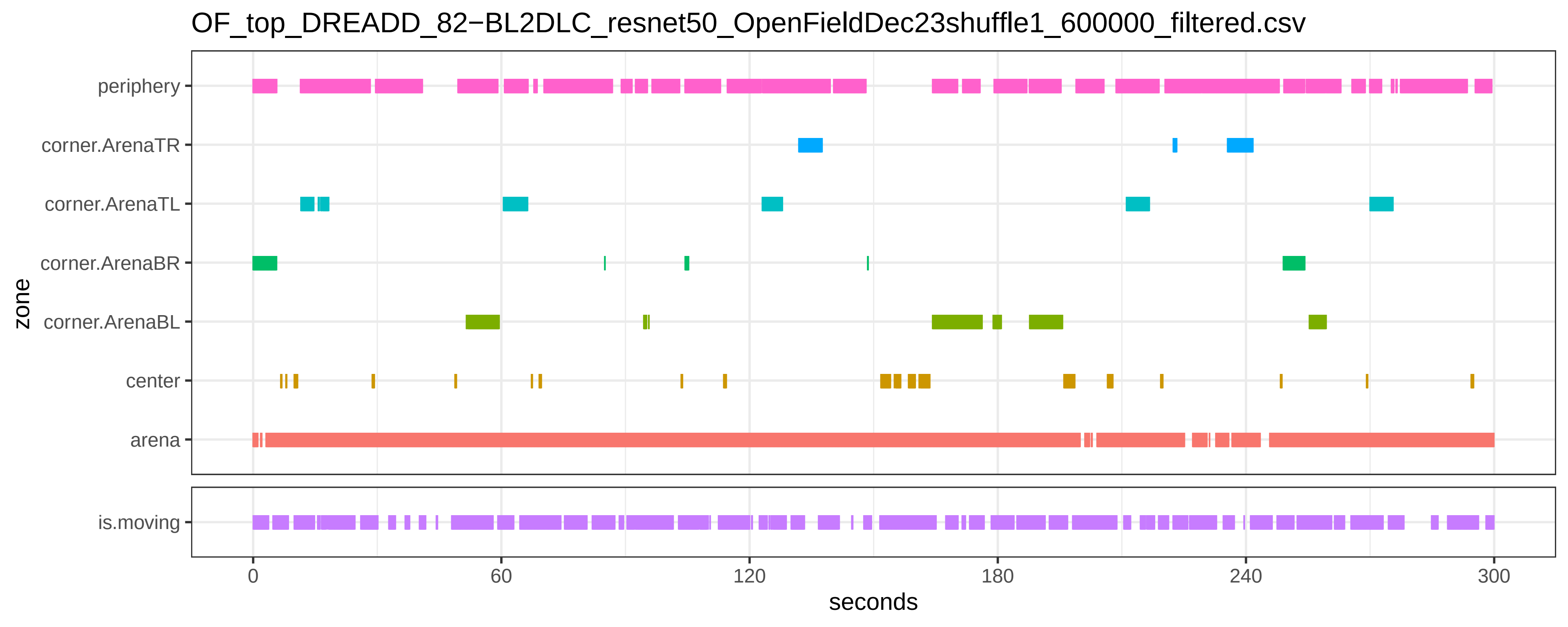

OF_top_DREADD_82−BL2DLC_resnet50_OpenFieldDec23shuffle1_600000_filtered.csv
periphery
corner.ArenaTR
corner.ArenaTL
corner.ArenaBR
corner.ArenaBL
center
arena
zone
is.moving
0
60
120
180
240
300
seconds

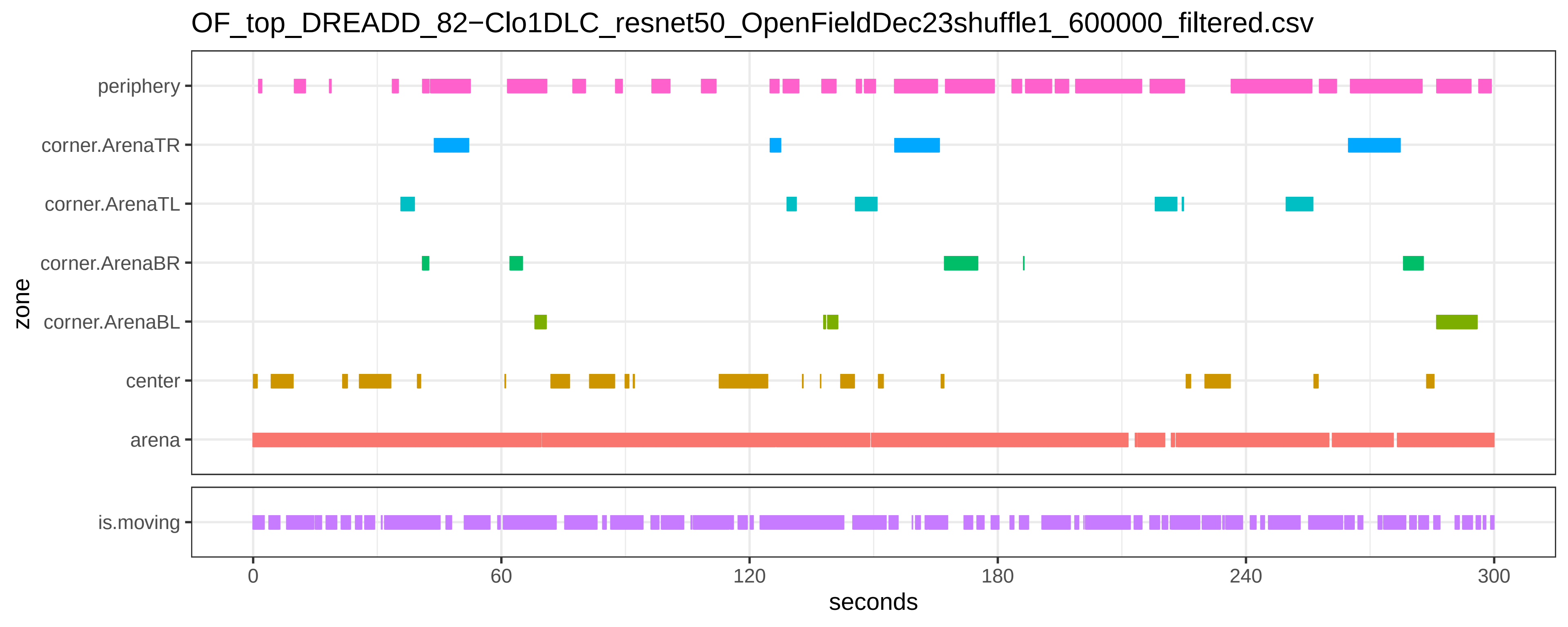
OF_top_DREADD_82−Clo1DLC_resnet50_OpenFieldDec23shuffle1_600000_filtered.csv
periphery
corner.ArenaTR
corner.ArenaTL
corner.ArenaBR
corner.ArenaBL
center
arena
is.moving
zone
0
60
120
180
240
300
seconds

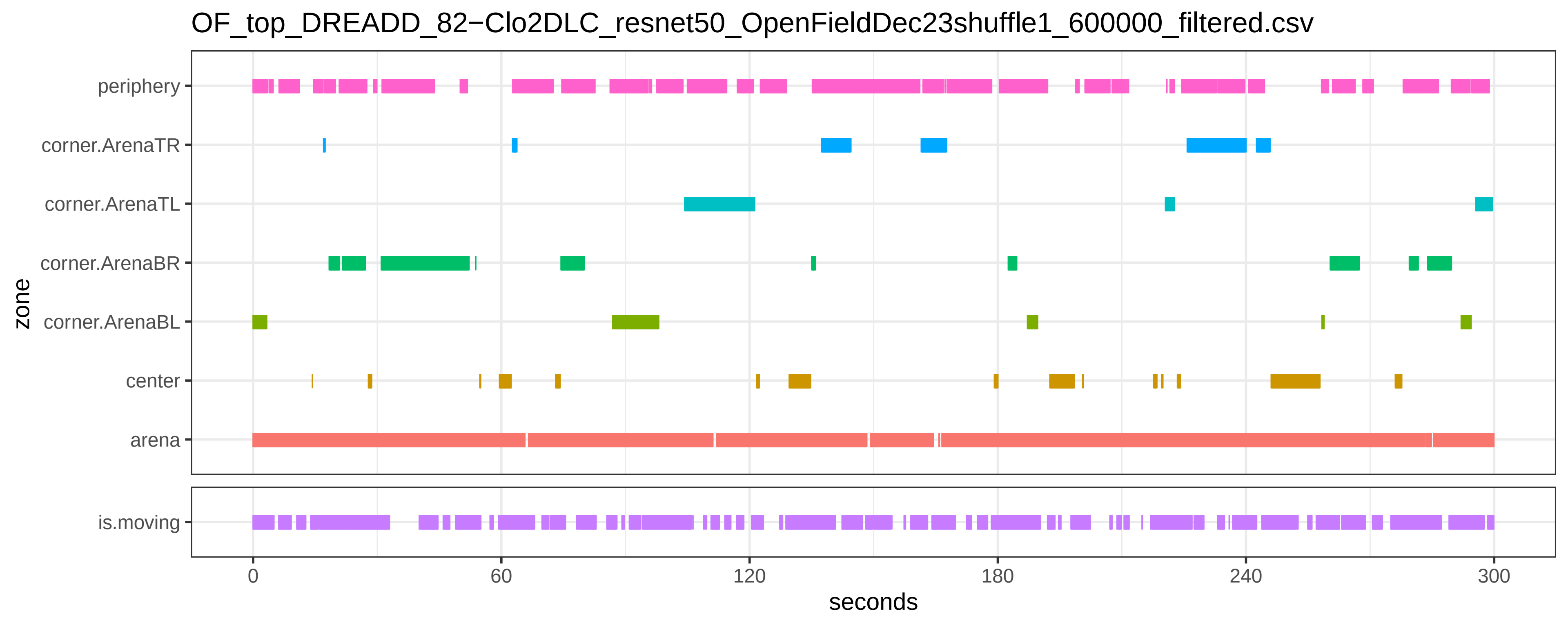

OF_top_DREADD_82−Clo2DLC_resnet50_OpenFieldDec23shuffle1_600000_filtered.csv
periphery
corner.ArenaTR
corner.ArenaTL
corner.ArenaBR
corner.ArenaBL
center
arena
zone
is.moving
0
60
120
180
240
300
seconds

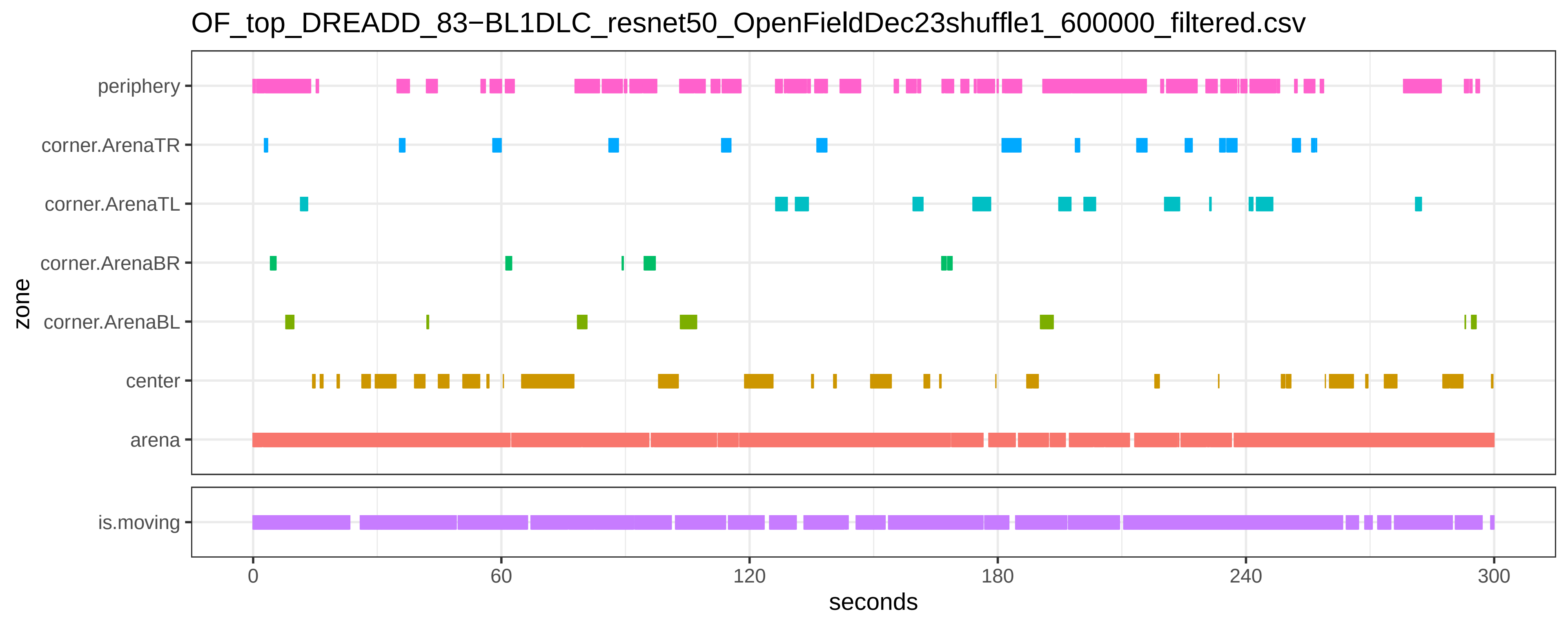

OF_top_DREADD_83−BL1DLC_resnet50_OpenFieldDec23shuffle1_600000_filtered.csv
periphery
corner.ArenaTR
corner.ArenaTL
corner.ArenaBR
corner.ArenaBL
center
arena
is.moving
zone
0
60
120
180
240
300
seconds

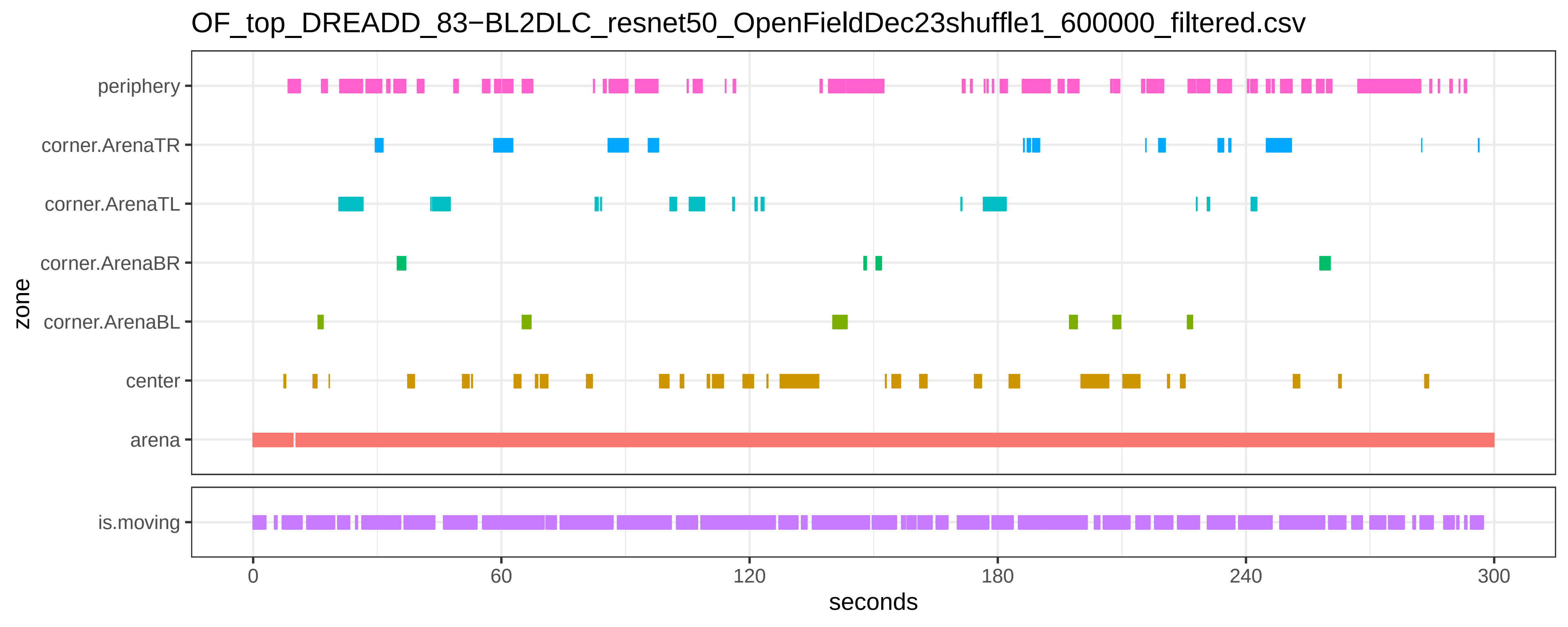
OF_top_DREADD_83−BL2DLC_resnet50_OpenFieldDec23shuffle1_600000_filtered.csv
zone
periphery
corner.ArenaTR
corner.ArenaTL
corner.ArenaBR
corner.ArenaBL
center
arena
is.moving
0
60
120
180
240
300
seconds

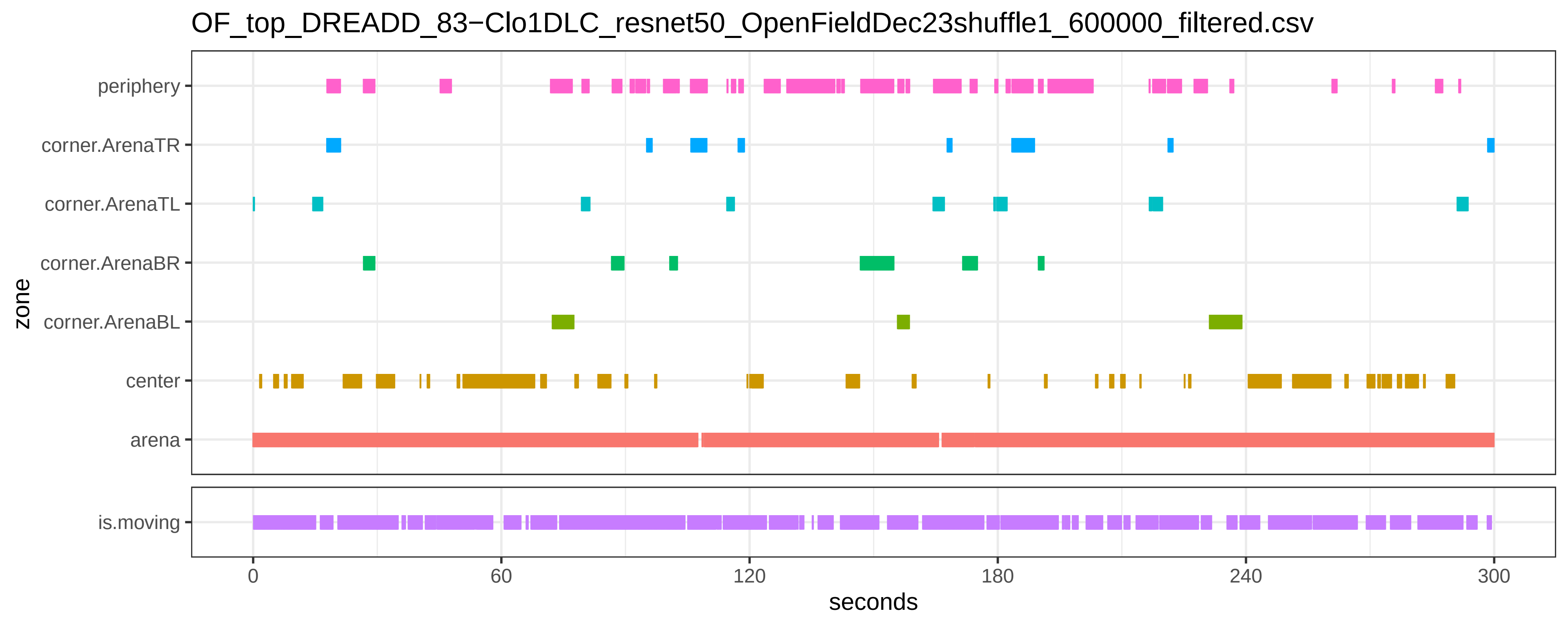

OF_top_DREADD_83−Clo1DLC_resnet50_OpenFieldDec23shuffle1_600000_filtered.csv
periphery
corner.ArenaTR
corner.ArenaTL
corner.ArenaBR
corner.ArenaBL
center
arena
zone
is.moving
0
60
120
180
240
300
seconds

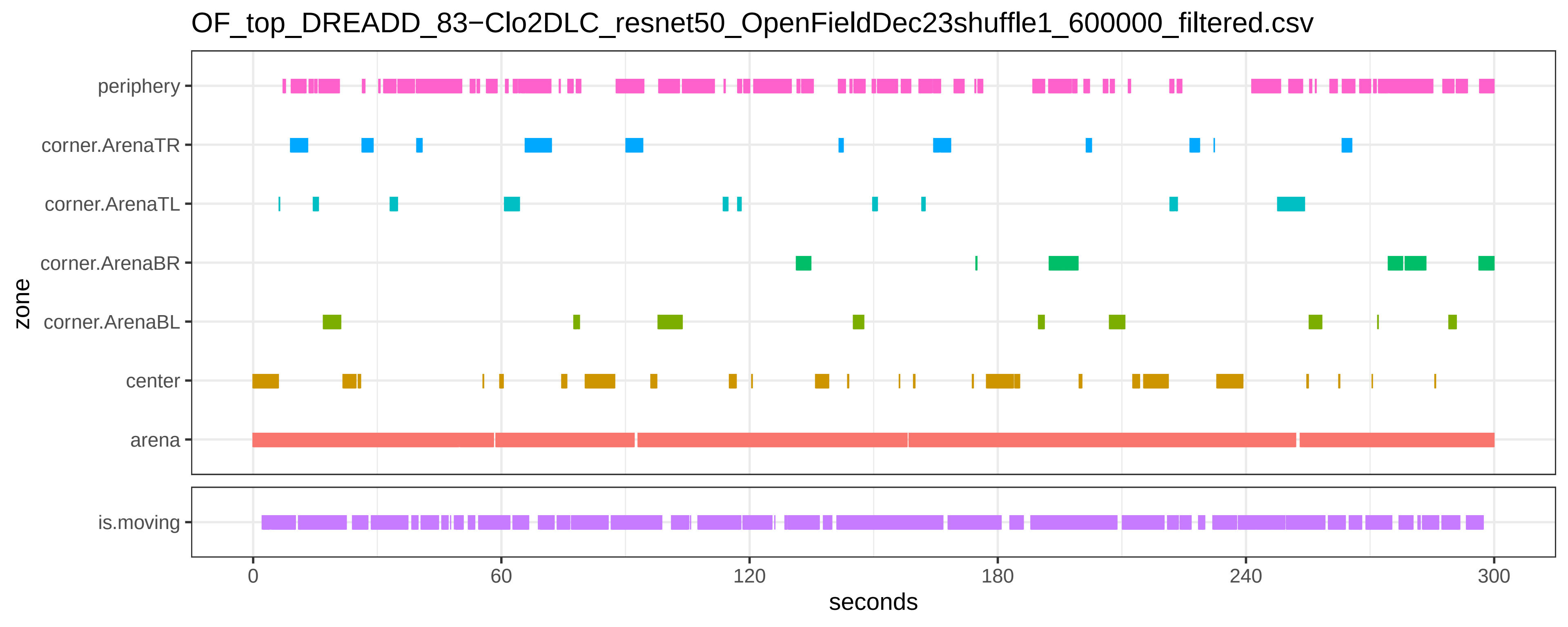

OF_top_DREADD_83−Clo2DLC_resnet50_OpenFieldDec23shuffle1_600000_filtered.csv
periphery
corner.ArenaTR
corner.ArenaTL
corner.ArenaBR
corner.ArenaBL
center
arena
zone
is.moving
0
60
120
180
240
300
seconds

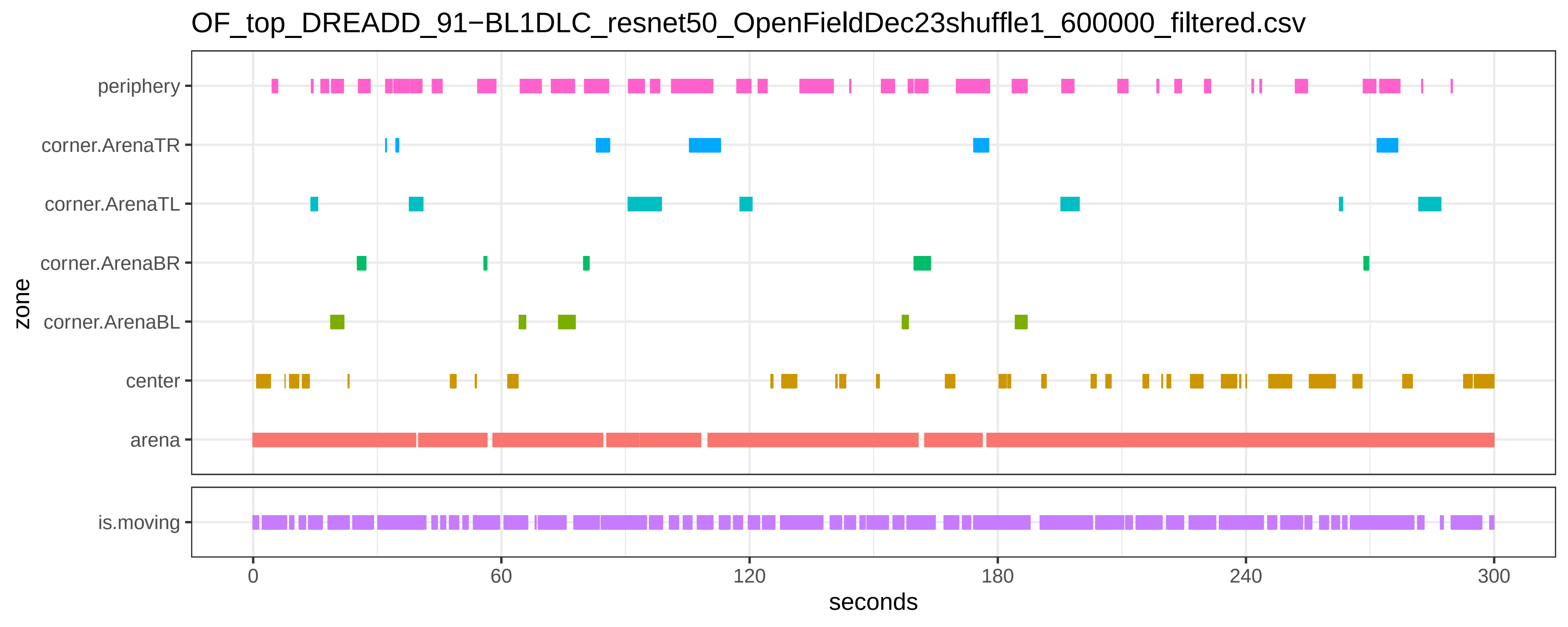
OF_top_DREADD_91−BL1DLC_resnet50_OpenFieldDec23shuffle1_600000_filtered.csv
periphery
corner.ArenaTR
corner.ArenaTL
corner.ArenaBR
corner.ArenaBL
center
arena
zone
is.moving
0
60
120
180
240
300
seconds

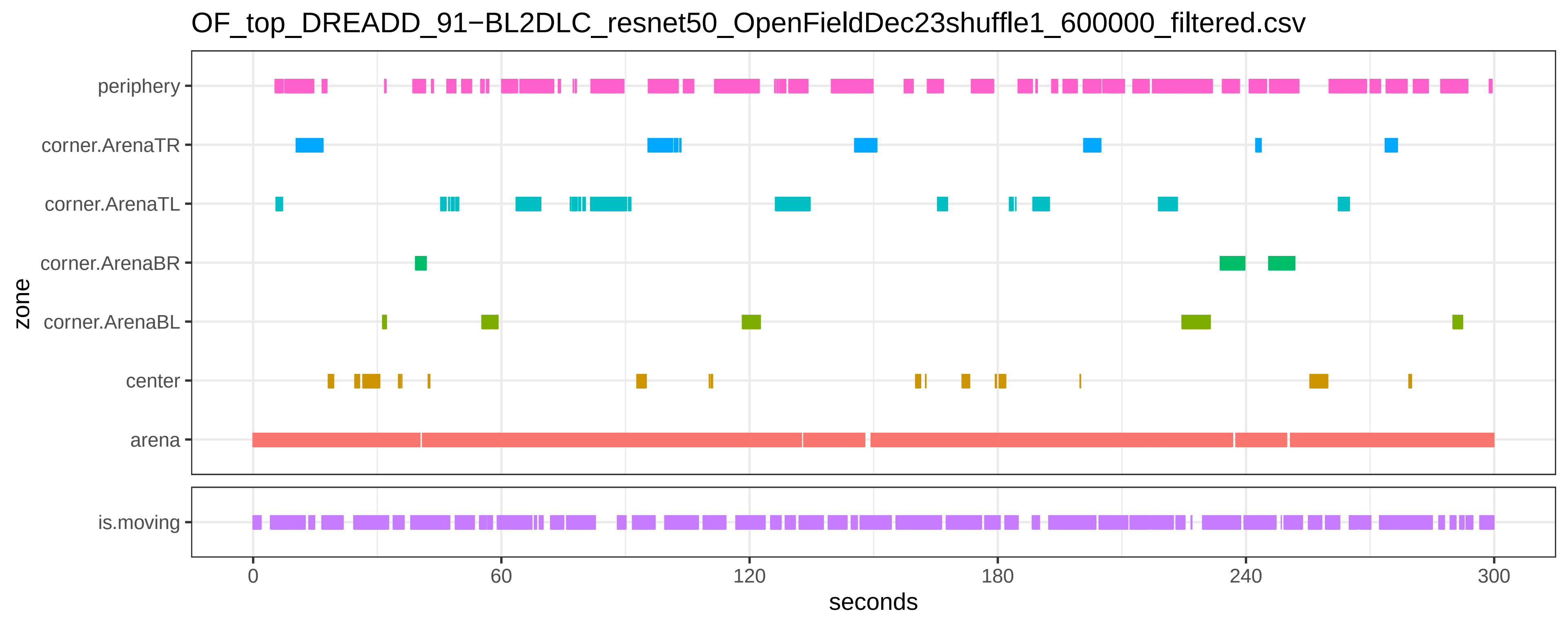
OF_top_DREADD_91−BL2DLC_resnet50_OpenFieldDec23shuffle1_600000_filtered.csv
periphery
corner.ArenaTR
corner.ArenaTL
corner.ArenaBR
corner.ArenaBL
center
arena
zone
is.moving
0
60
120
180
240
300
seconds

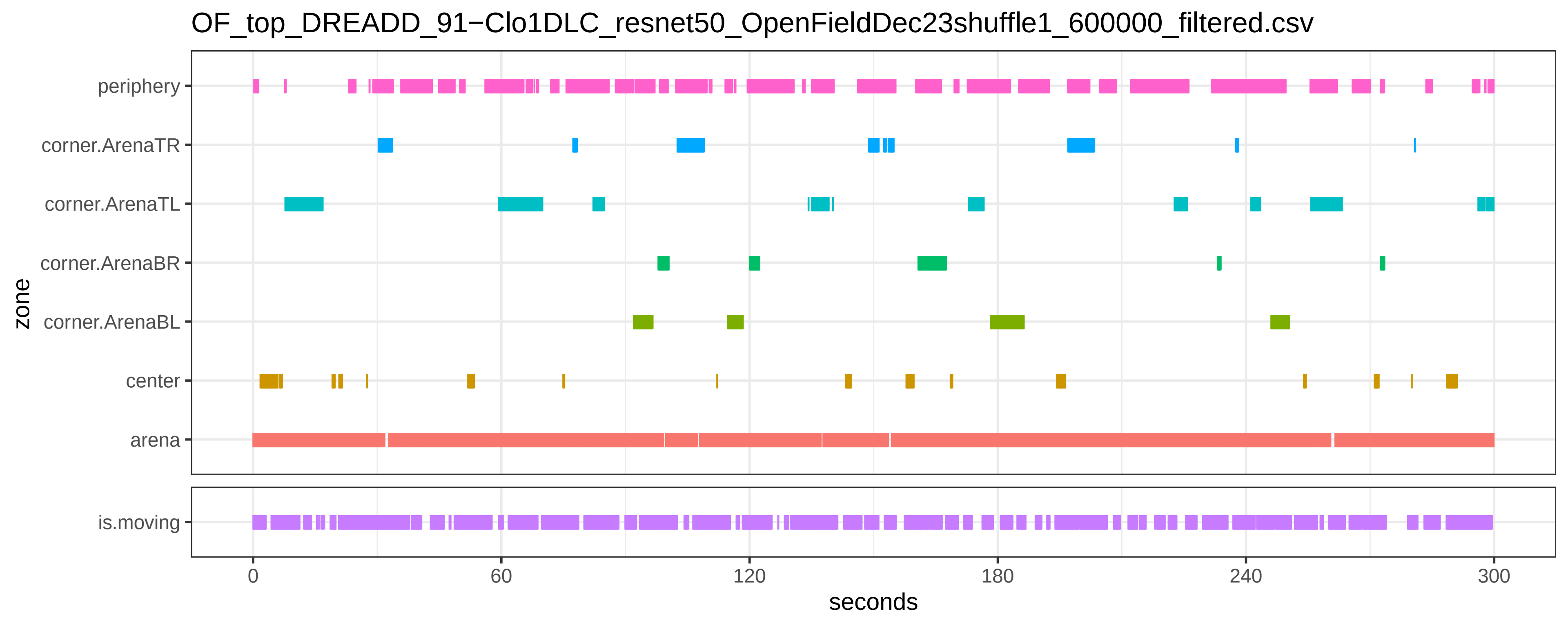

OF_top_DREADD_91−Clo1DLC_resnet50_OpenFieldDec23shuffle1_600000_filtered.csv
zone
periphery
corner.ArenaTR
corner.ArenaTL
corner.ArenaBR
corner.ArenaBL
center
arena
is.moving
0
60
120
180
240
300
seconds

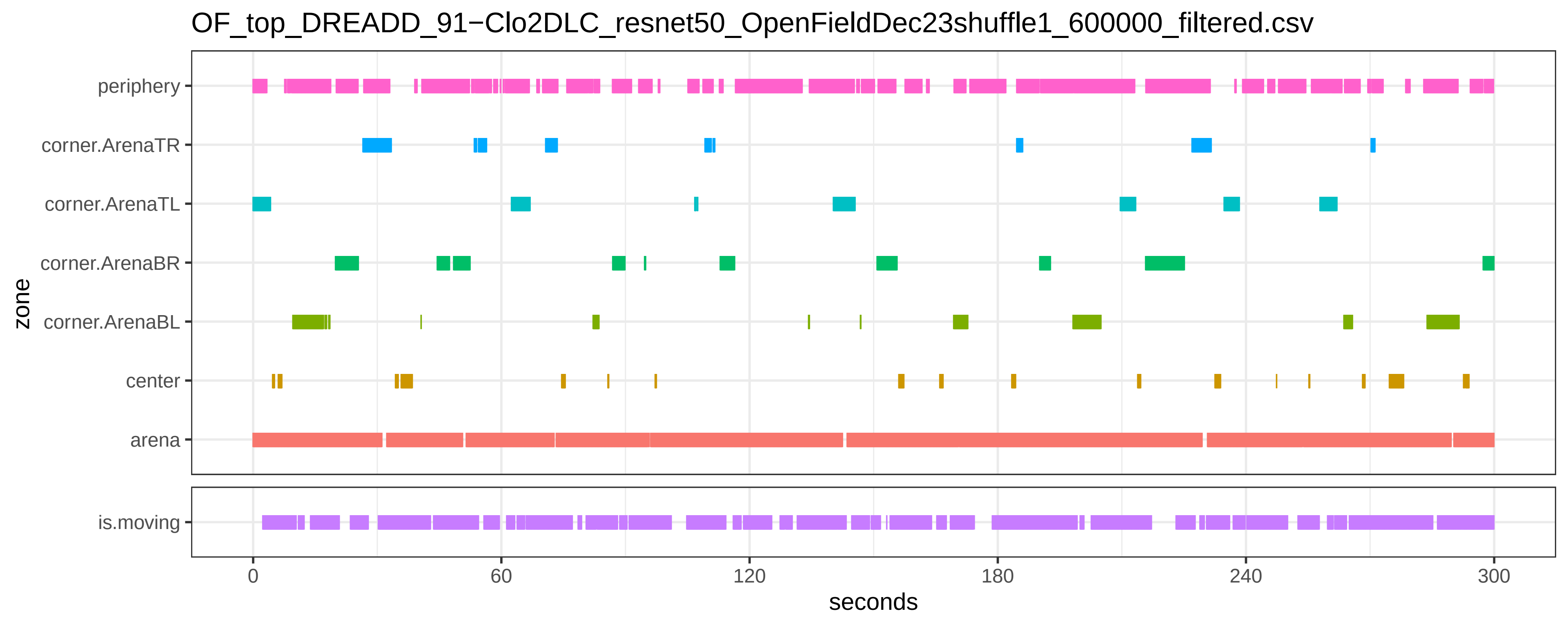
OF_top_DREADD_91−Clo2DLC_resnet50_OpenFieldDec23shuffle1_600000_filtered.csv
periphery
corner.ArenaTR
corner.ArenaTL
corner.ArenaBR
corner.ArenaBL
center
arena
zone
is.moving
0
60
120
180
240
300
seconds

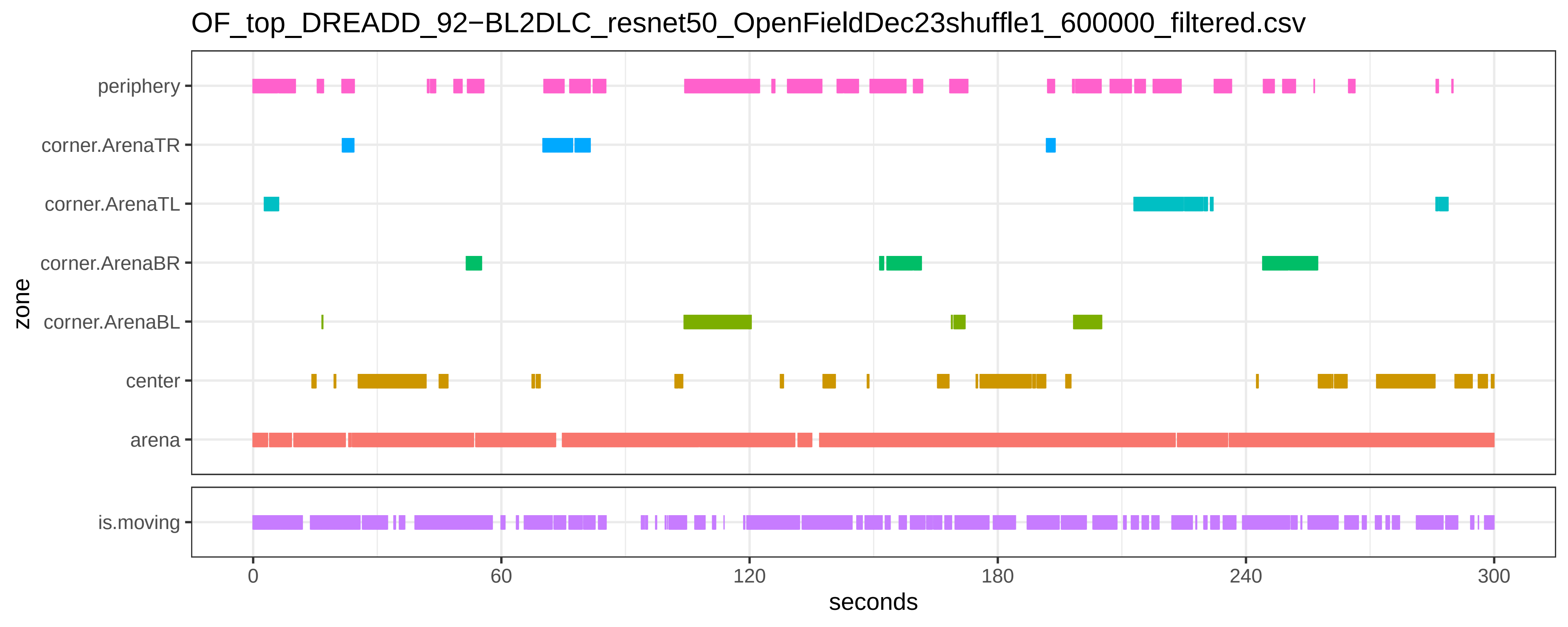

OF_top_DREADD_92−BL2DLC_resnet50_OpenFieldDec23shuffle1_600000_filtered.csv
zone
periphery
corner.ArenaTR
corner.ArenaTL
corner.ArenaBR
corner.ArenaBL
center
arena
is.moving
0
60
120
180
240
300
seconds

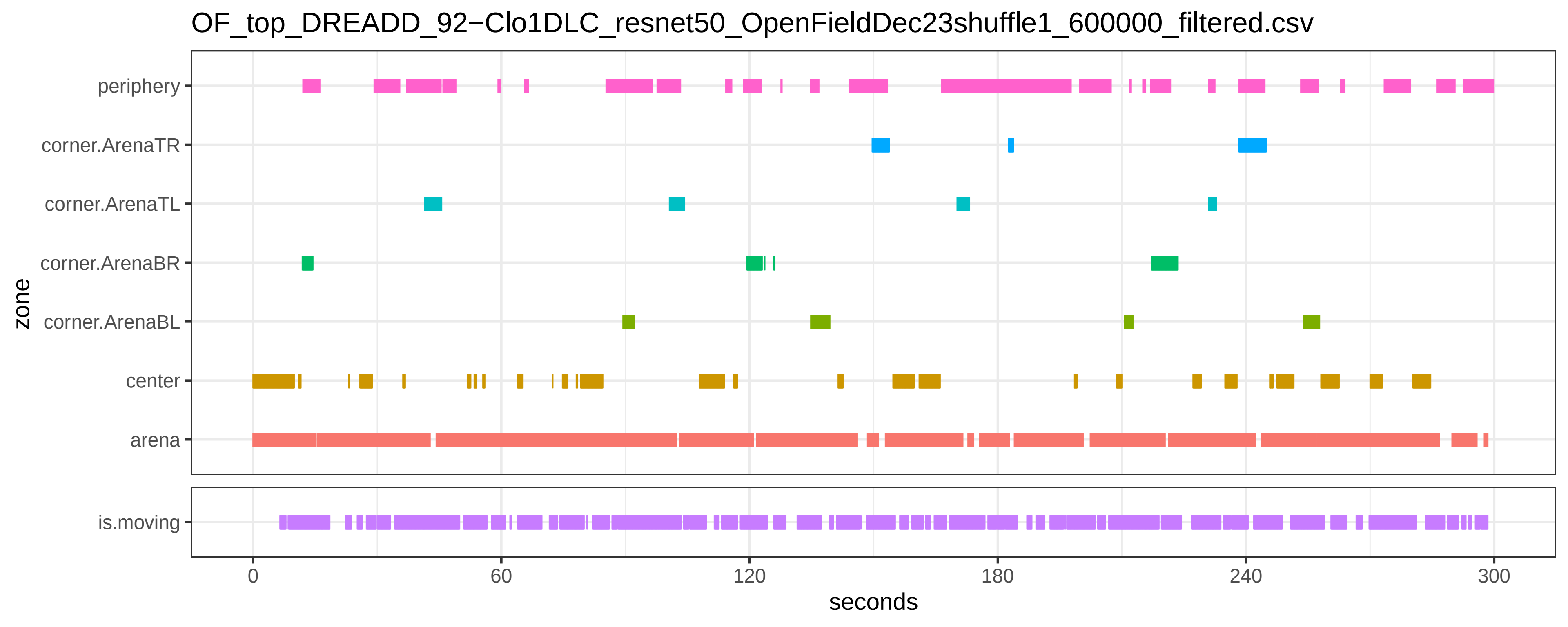
OF_top_DREADD_92−Clo1DLC_resnet50_OpenFieldDec23shuffle1_600000_filtered.csv
periphery
corner.ArenaTR
corner.ArenaTL
corner.ArenaBR
corner.ArenaBL
center
arena
is.moving
zone
0
60
120
180
240
300
seconds

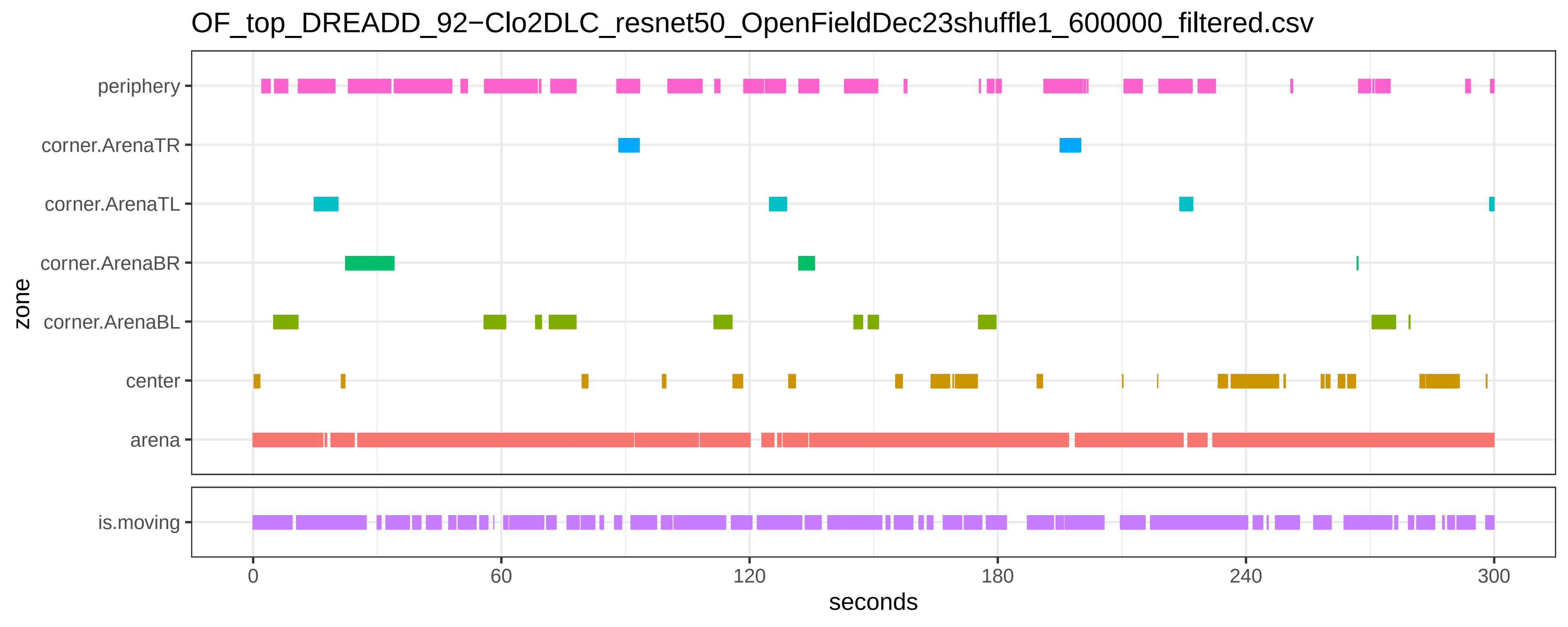
OF_top_DREADD_92−Clo2DLC_resnet50_OpenFieldDec23shuffle1_600000_filtered.csv
periphery
corner.ArenaTR
corner.ArenaTL
corner.ArenaBR
corner.ArenaBL
center
arena
zone
is.moving
0
60
120
180
240
300
seconds

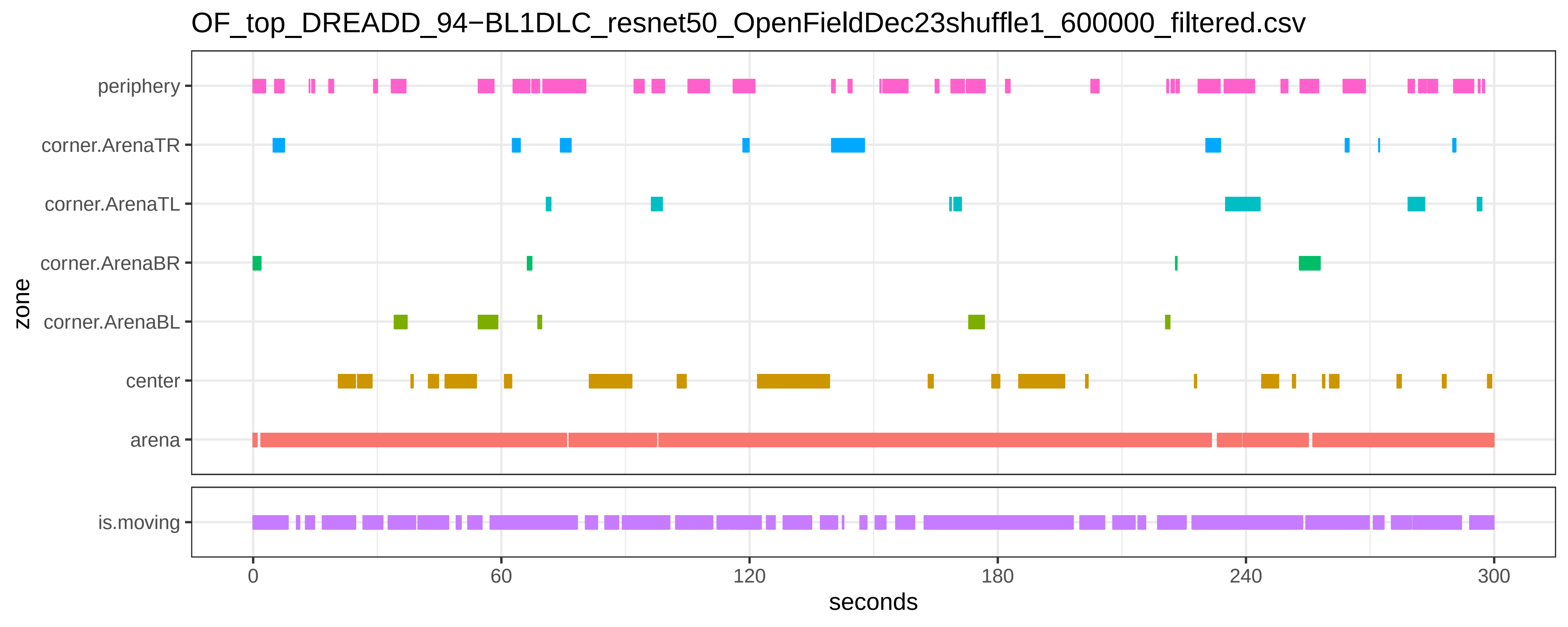

OF_top_DREADD_94−BL1DLC_resnet50_OpenFieldDec23shuffle1_600000_filtered.csv
periphery
corner.ArenaTR
corner.ArenaTL
corner.ArenaBR
corner.ArenaBL
center
arena
is.moving
zone
0
60
120
180
240
300
seconds

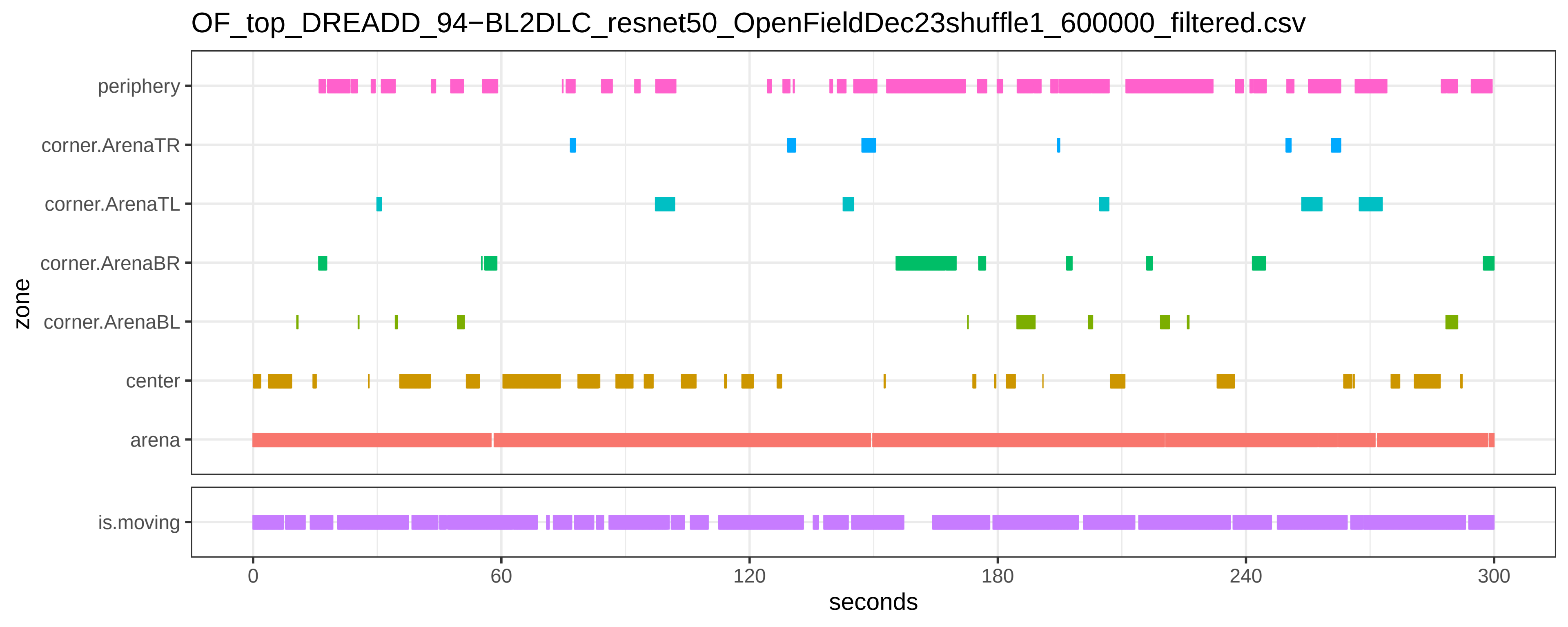

OF_top_DREADD_94−BL2DLC_resnet50_OpenFieldDec23shuffle1_600000_filtered.csv
periphery
corner.ArenaTR
corner.ArenaTL
corner.ArenaBR
corner.ArenaBL
center
arena
is.moving
zone
0
60
120
180
240
300
seconds

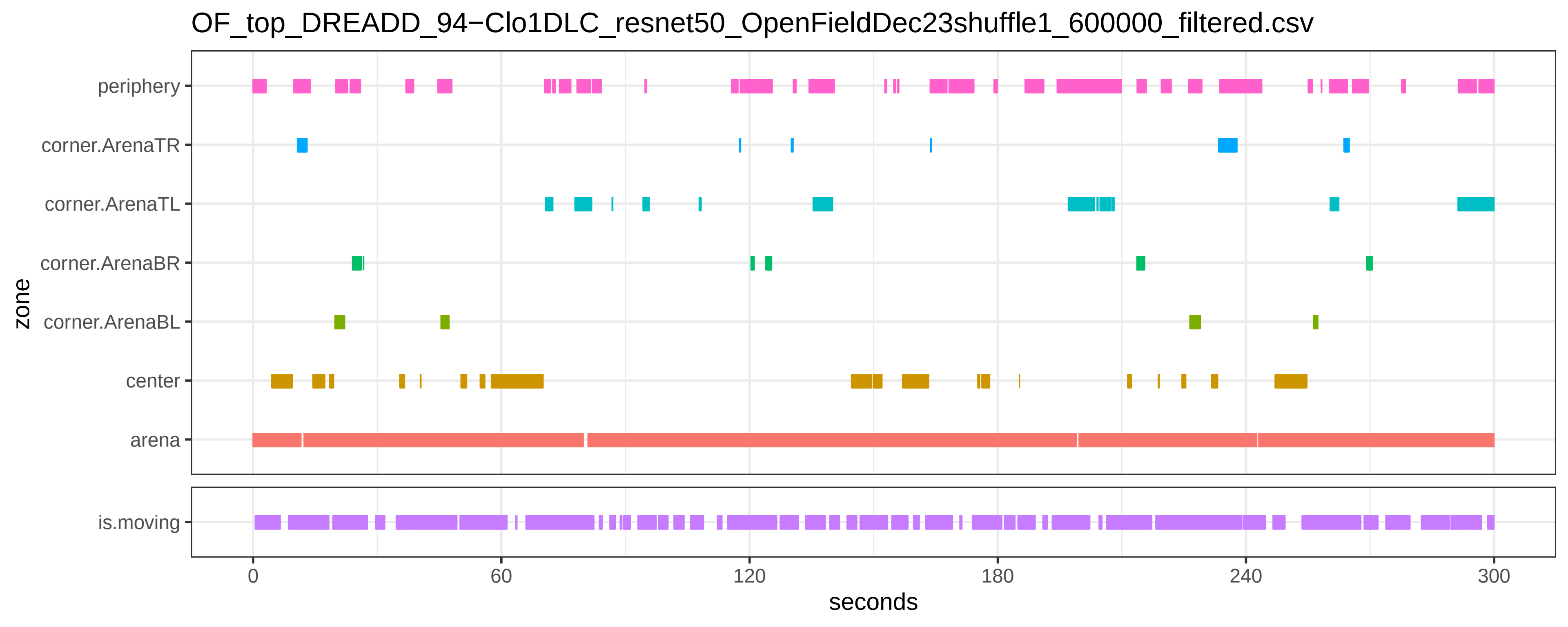
OF_top_DREADD_94−Clo1DLC_resnet50_OpenFieldDec23shuffle1_600000_filtered.csv
periphery
corner.ArenaTR
corner.ArenaTL
corner.ArenaBR
corner.ArenaBL
center
arena
is.moving
zone
0
60
120
180
240
300
seconds

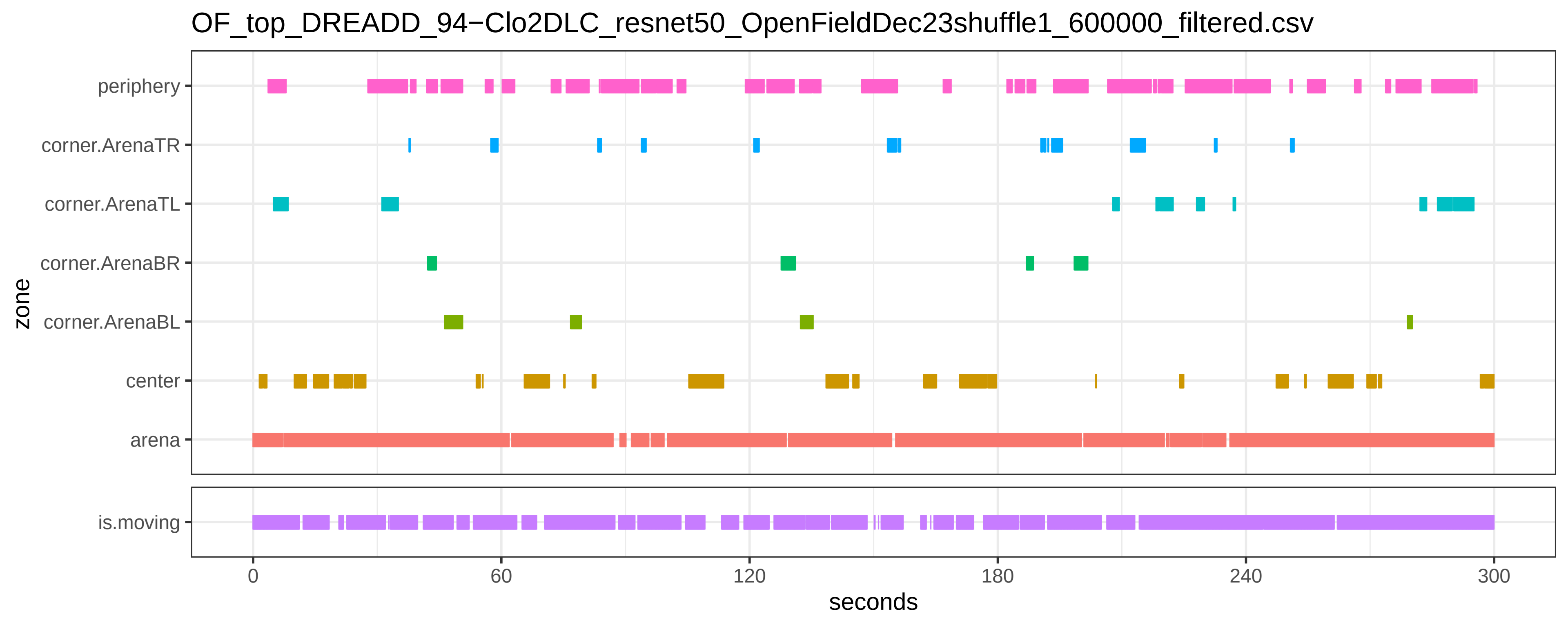
OF_top_DREADD_94−Clo2DLC_resnet50_OpenFieldDec23shuffle1_600000_filtered.csv
periphery
corner.ArenaTR
corner.ArenaTL
corner.ArenaBR
corner.ArenaBL
center
arena
zone
is.moving
0
60
120
180
240
300
seconds
