## SupplementaryFile_S4 for "Chemo- and optogenetic activation of hypothalamic Foxb1-expressing neurons and their terminal endings in the rostral-dorsolateral PAG leads to tachypnea, bradycardia, and immobility"

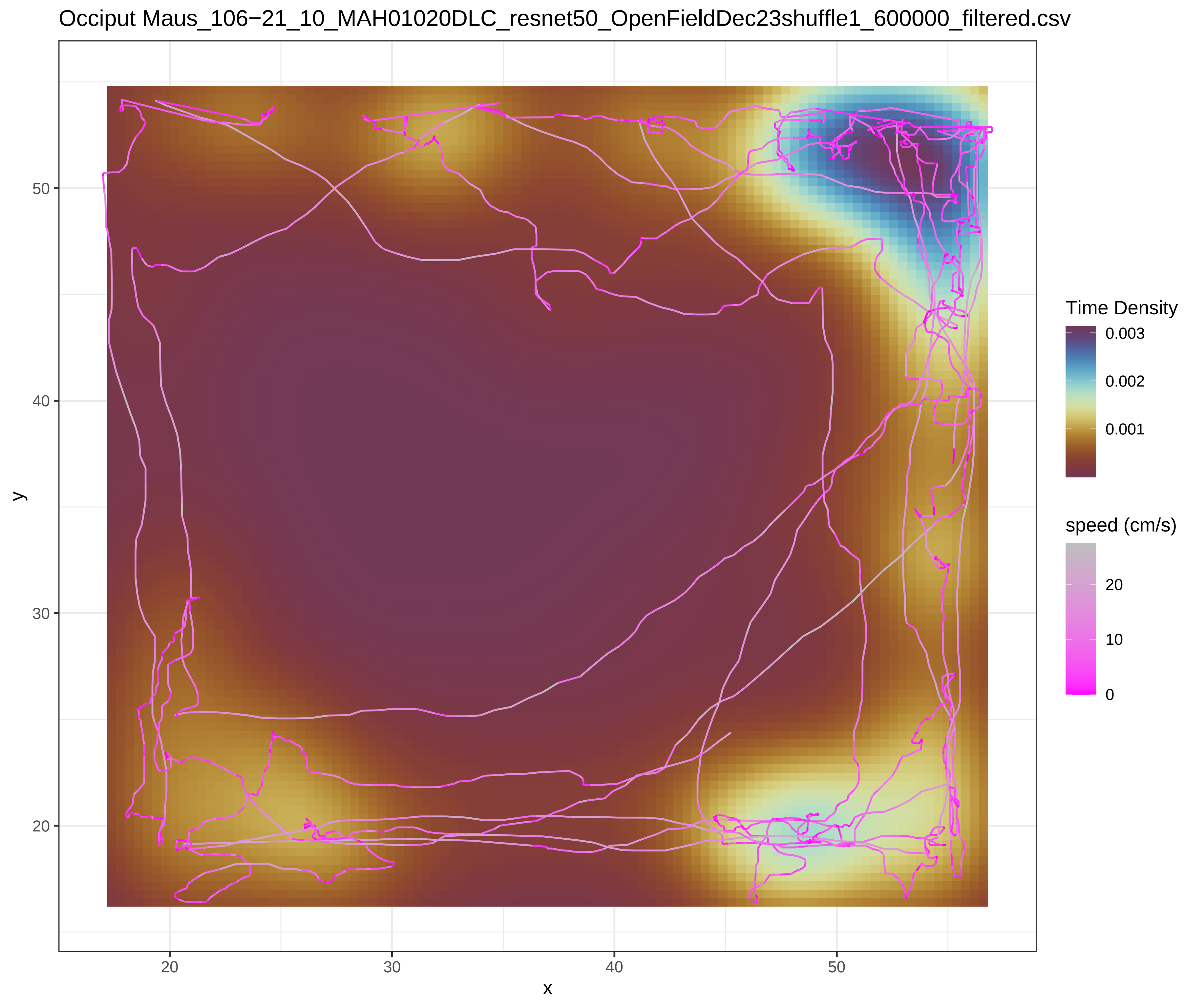

Occiput Maus_106−21_10_MAH01020DLC_resnet50_OpenFieldDec23shuffle1_600000_filtered.csv
Time Density
0.003
0.002
0.001
speed (cm/s)
20
10
0
y
50
40
30
20
x
20
30
40
50


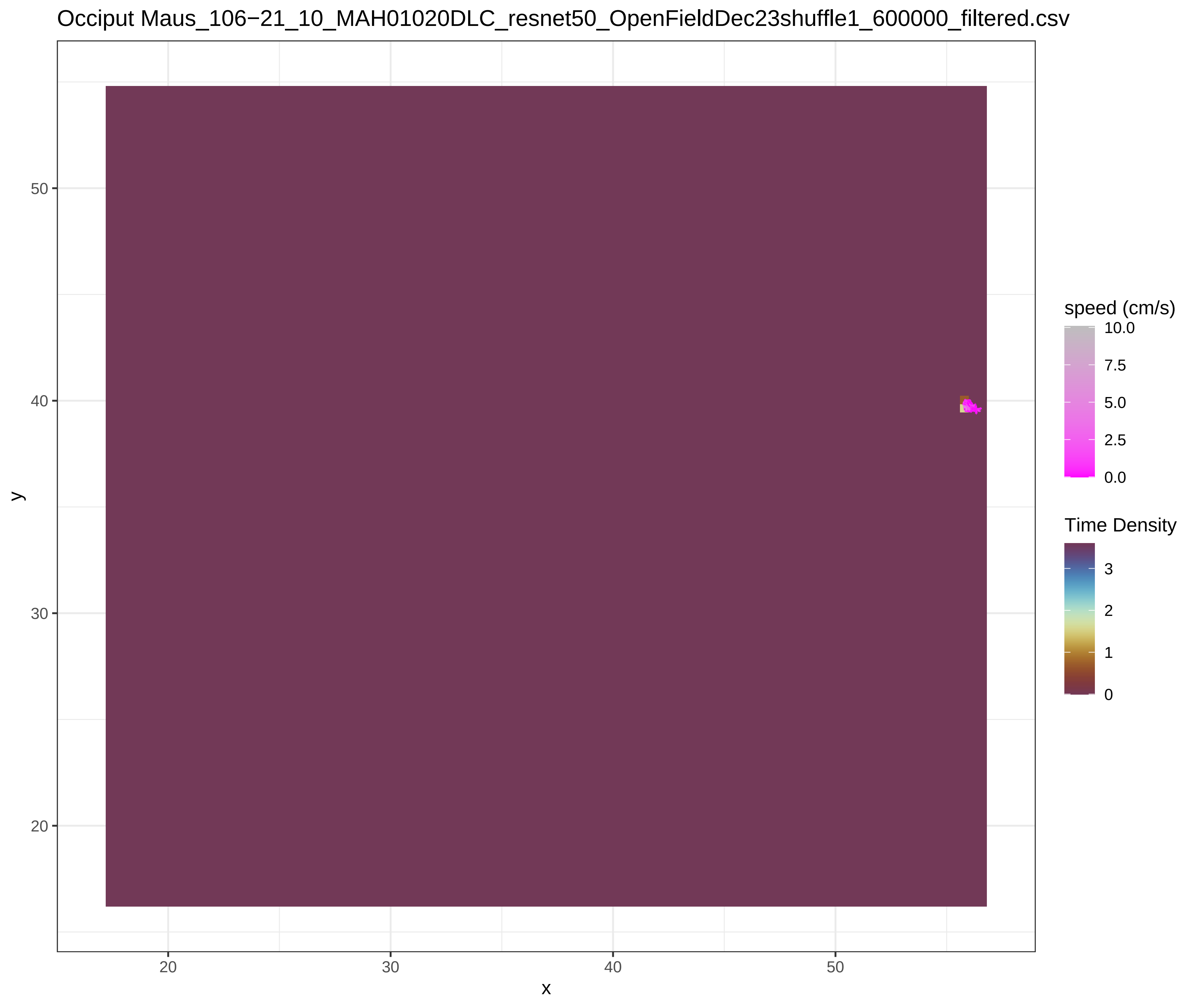

Occiput Maus_106−21_10_MAH01020DLC_resnet50_OpenFieldDec23shuffle1_600000_filtered.csv
speed (cm/s)
10.0
7.5
5.0
2.5
0.0
Time Density
3
2
1
0
y
50
40
30
20
x
20
30
40
50

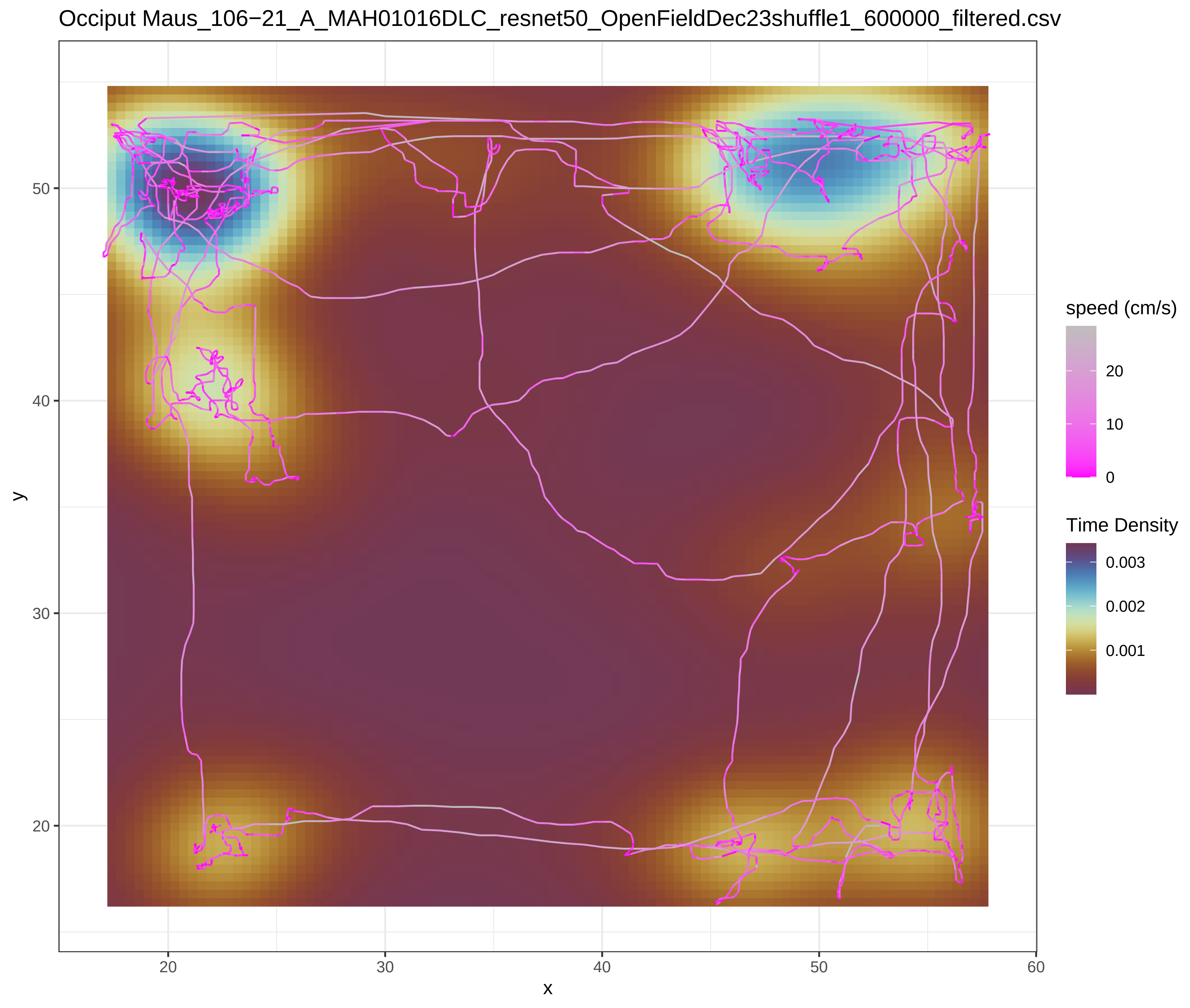

Occiput Maus_106−21_A_MAH01016DLC_resnet50_OpenFieldDec23shuffle1_600000_filtered.csv
speed (cm/s)
20
10
0
Time Density
0.003
0.002
0.001
y
50
40
30
20
x
20
30
40
50
60


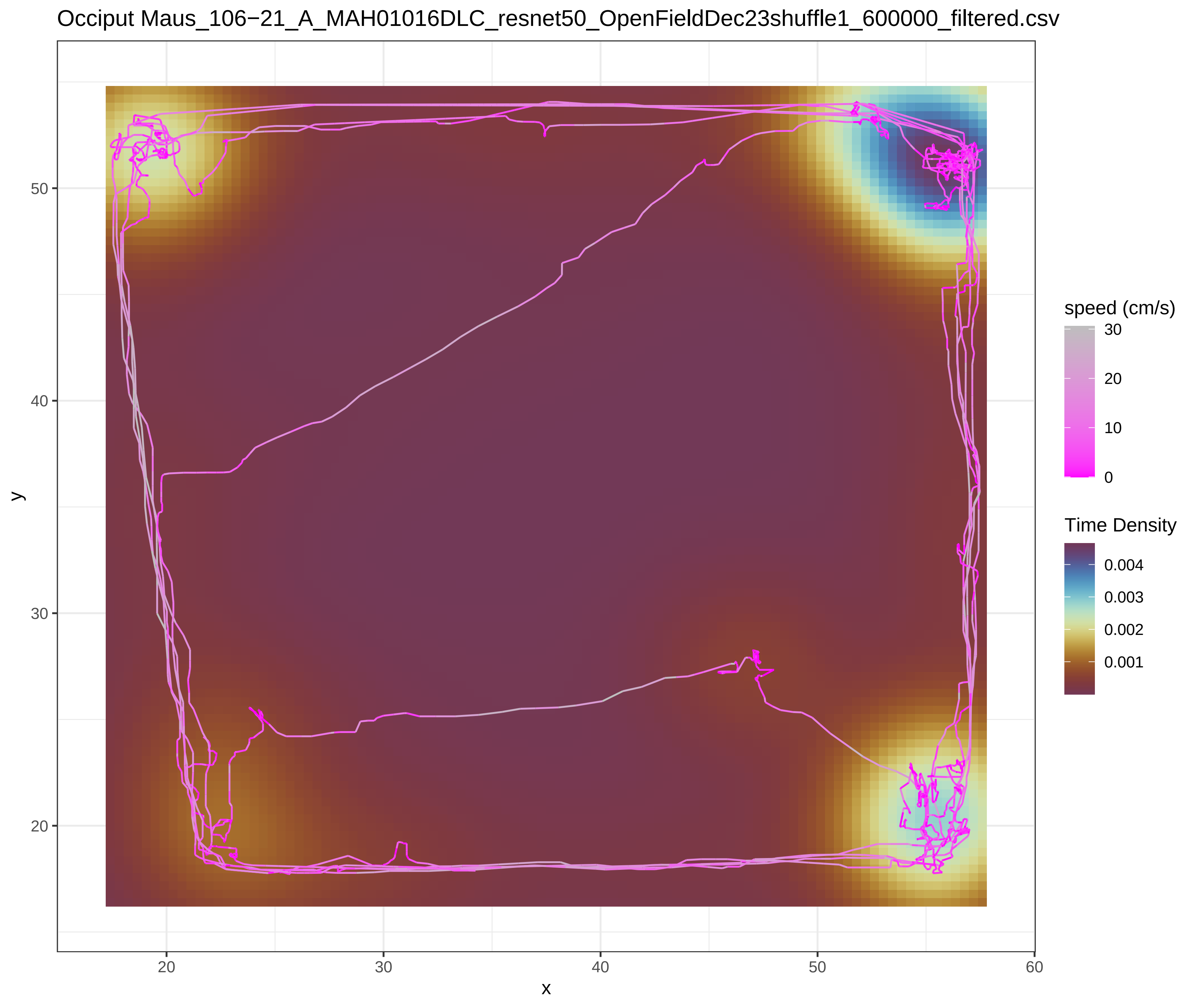

Occiput Maus_106−21_A_MAH01016DLC_resnet50_OpenFieldDec23shuffle1_600000_filtered.csv
speed (cm/s)
30
20
10
0
Time Density
0.004
0.003
0.002
0.001
y
50
40
30
20
x
20
30
40
50
60

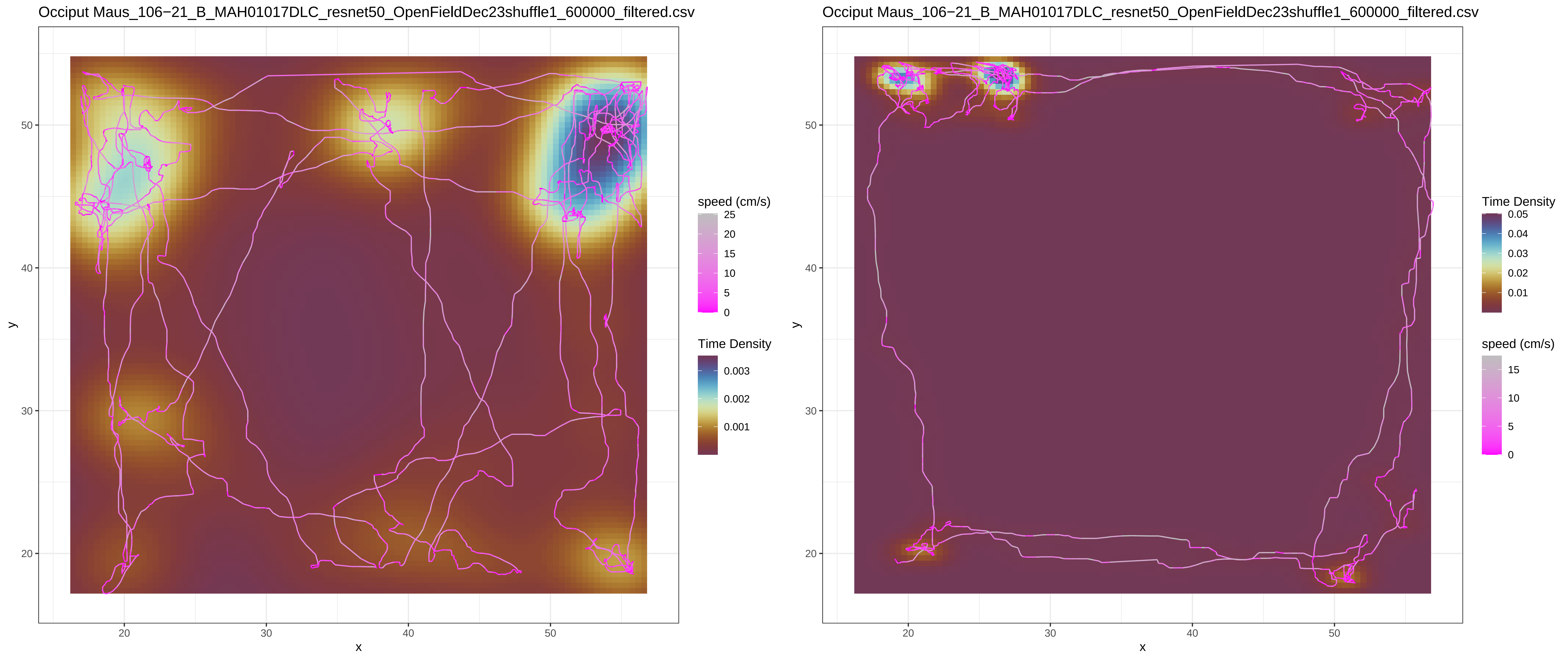

Occiput Maus_106−21_B_MAH01017DLC_resnet50_OpenFieldDec23shuffle1_600000_filtered.csv
speed (cm/s)
25
20
15
10
5
0
Time Density
0.003
0.002
0.001
y
x
50
40
30
20
20
30
40
50
Occiput Maus_106−21_B_MAH01017DLC_resnet50_OpenFieldDec23shuffle1_600000_filtered.csv
Time Density
0.05
0.04
0.03
0.02
0.01
speed (cm/s)
15
10
5
0

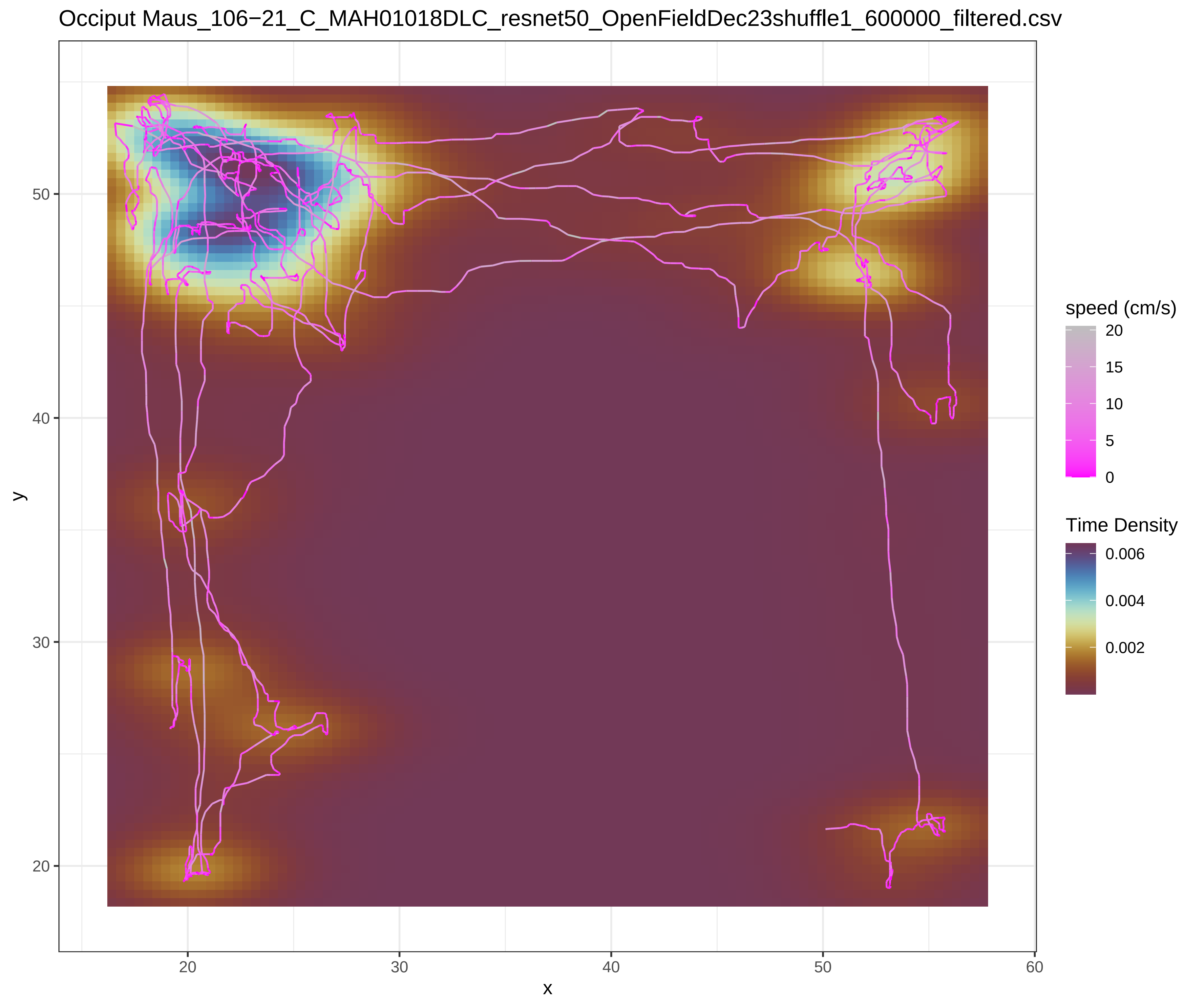
Occiput Maus_106−21_C_MAH01018DLC_resnet50_OpenFieldDec23shuffle1_600000_filtered.csv
y
x
20
30
40
50
60
speed (cm/s)
20
15
10
5
0
Time Density
0.006
0.004
0.002

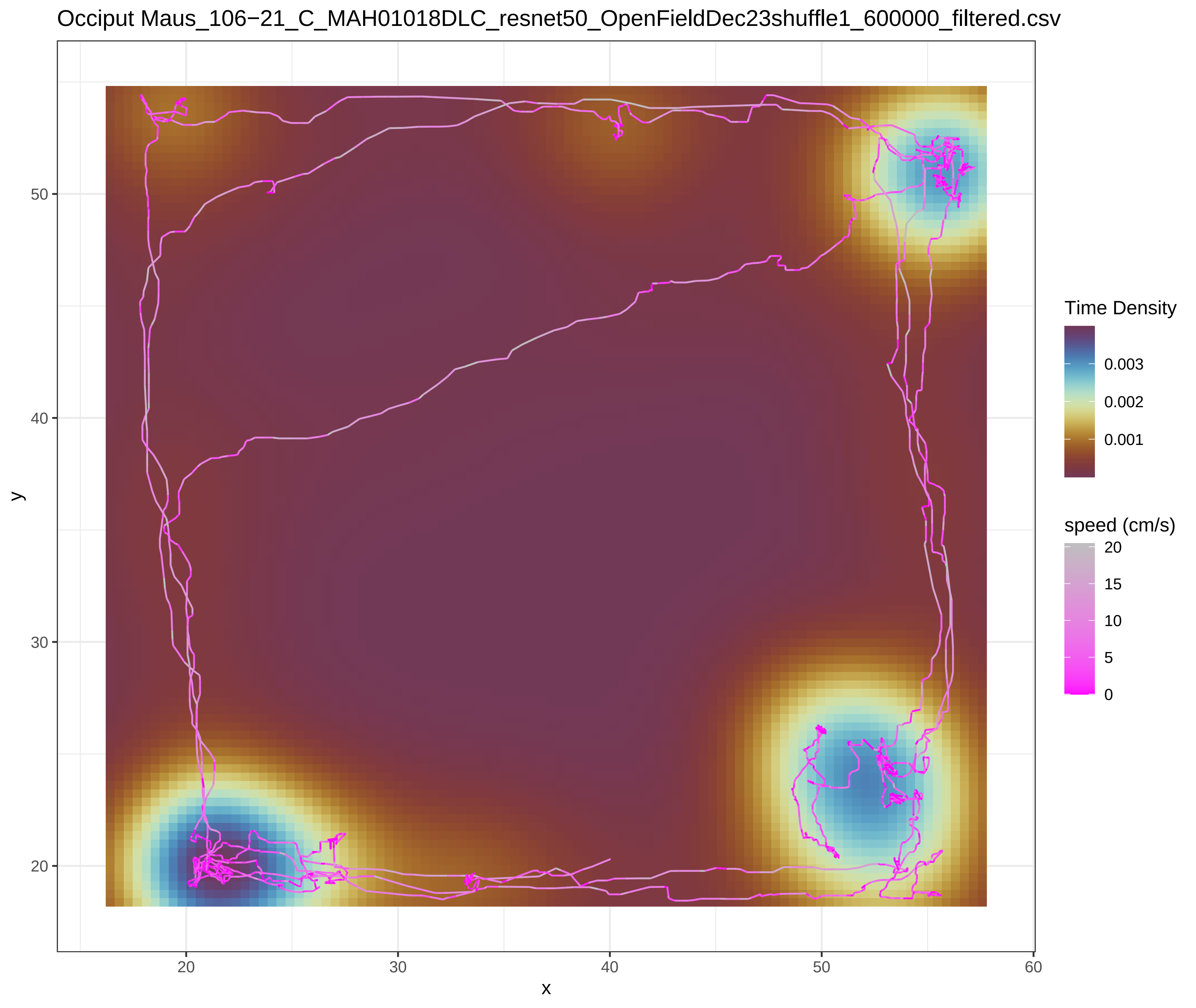
Occiput Maus_106−21_C_MAH01018DLC_resnet50_OpenFieldDec23shuffle1_600000_filtered.csv
y
x
20
30
40
50
60
Time Density
0.003
0.002
0.001
speed (cm/s)
20
15
10
5
0

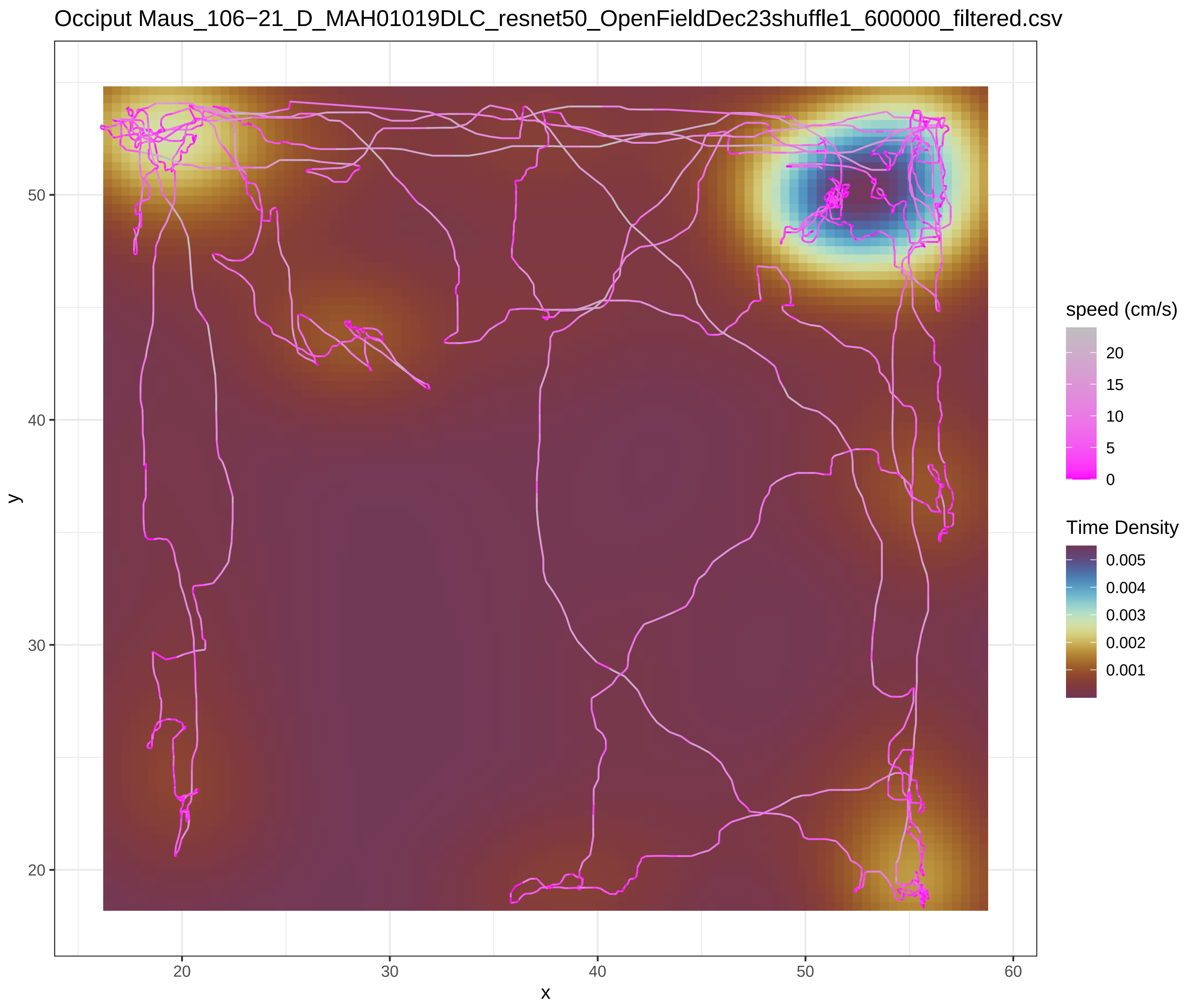
Occiput Maus_106−21_D_MAH01019DLC_resnet50_OpenFieldDec23shuffle1_600000_filtered.csv
speed (cm/s)
20
15
10
5
0
Time Density
0.005
0.004
0.003
0.002
0.001
y
50
40
30
20
x
20
30
40
50
60

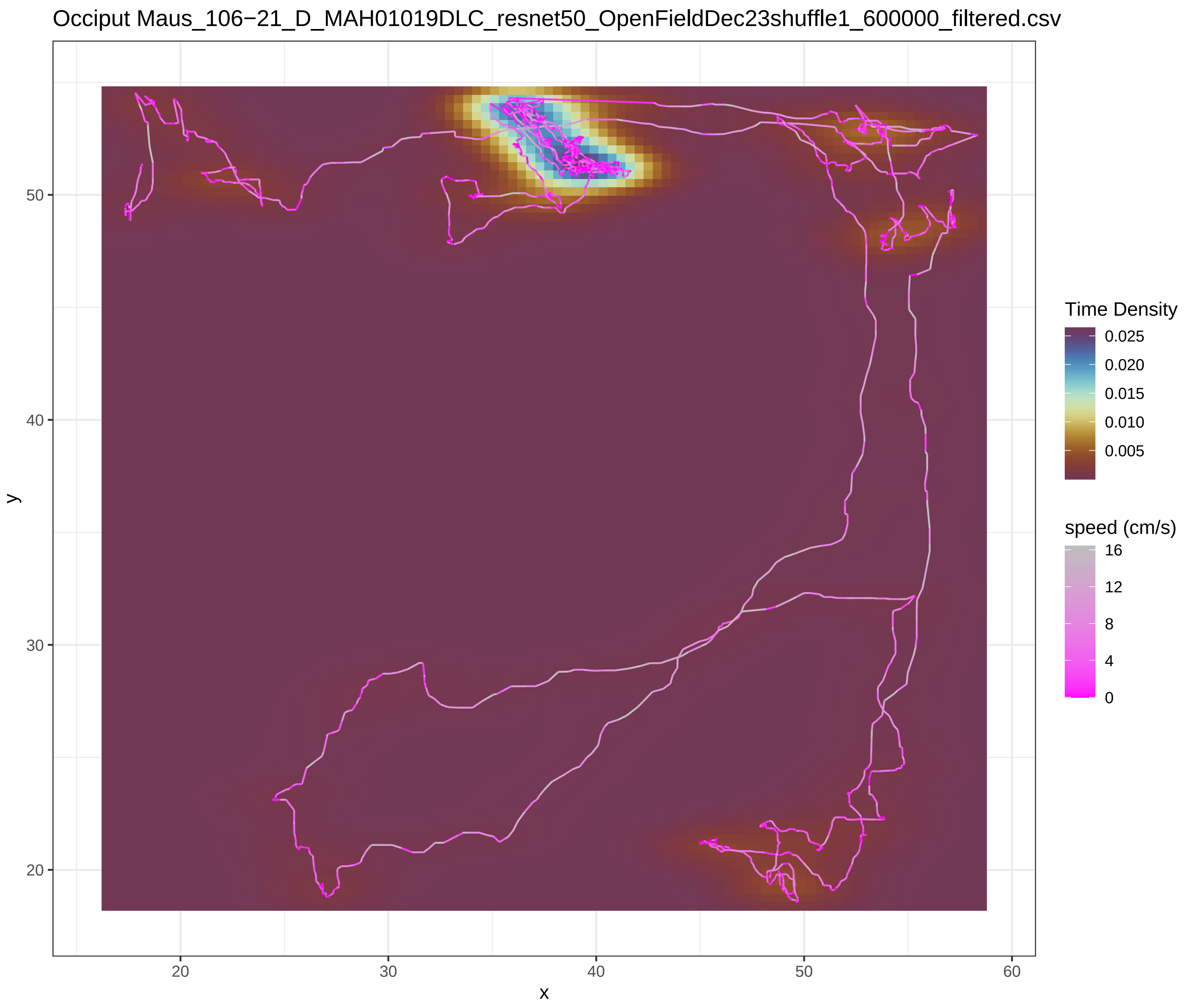
Occiput Maus_106−21_D_MAH01019DLC_resnet50_OpenFieldDec23shuffle1_600000_filtered.csv
Time Density
0.025
0.020
0.015
0.010
0.005
speed (cm/s)
16
12
8
4
0
y
50
40
30
20
x
20
30
40
50
60

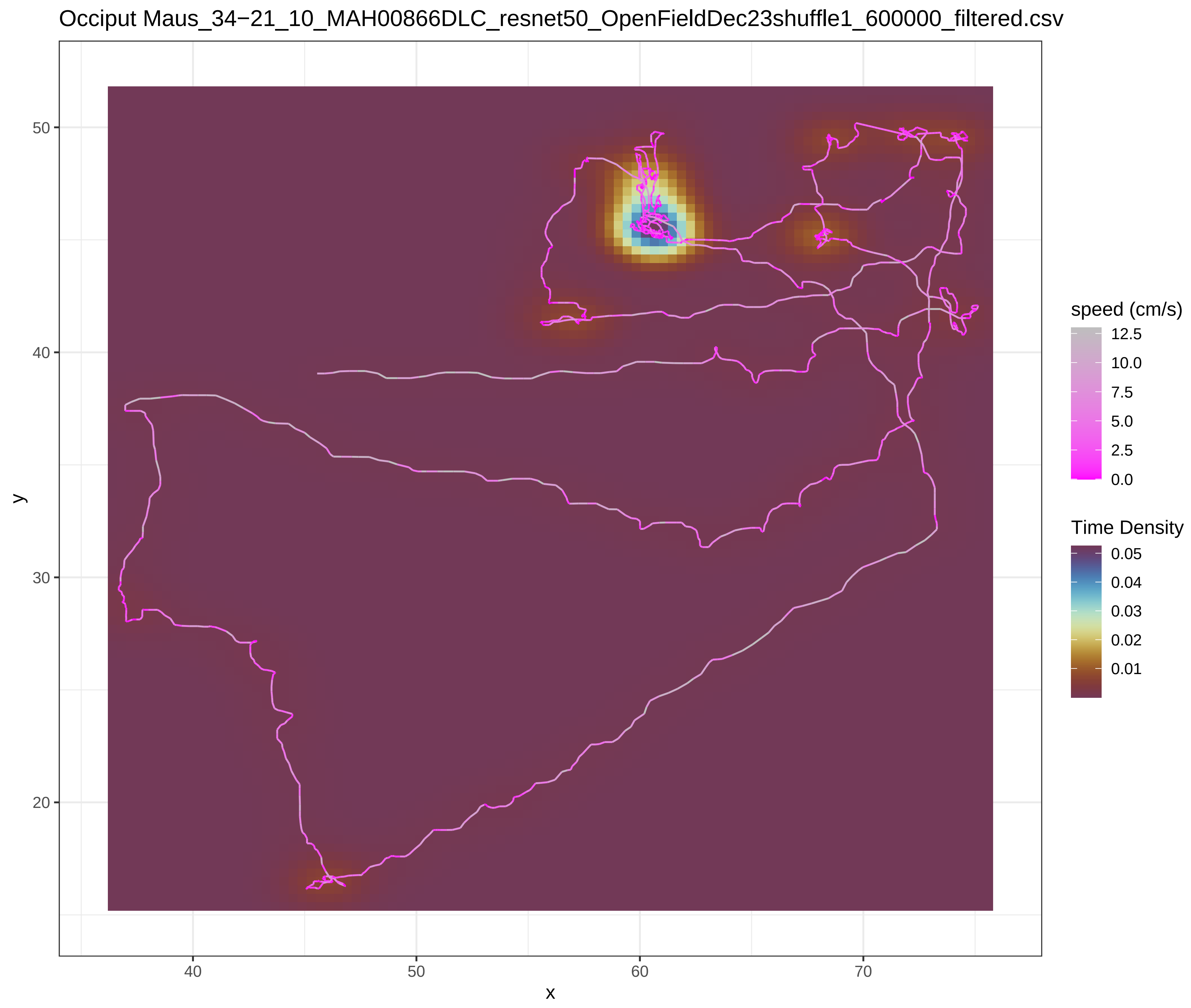
Occiput Maus_34−21_10_MAH00866DLC_resnet50_OpenFieldDec23shuffle1_600000_filtered.csv
speed (cm/s)
12.5
10.0
7.5
5.0
2.5
0.0
Time Density
0.05
0.04
0.03
0.02
0.01
y
x
50
40
30
20
40
50
60
70

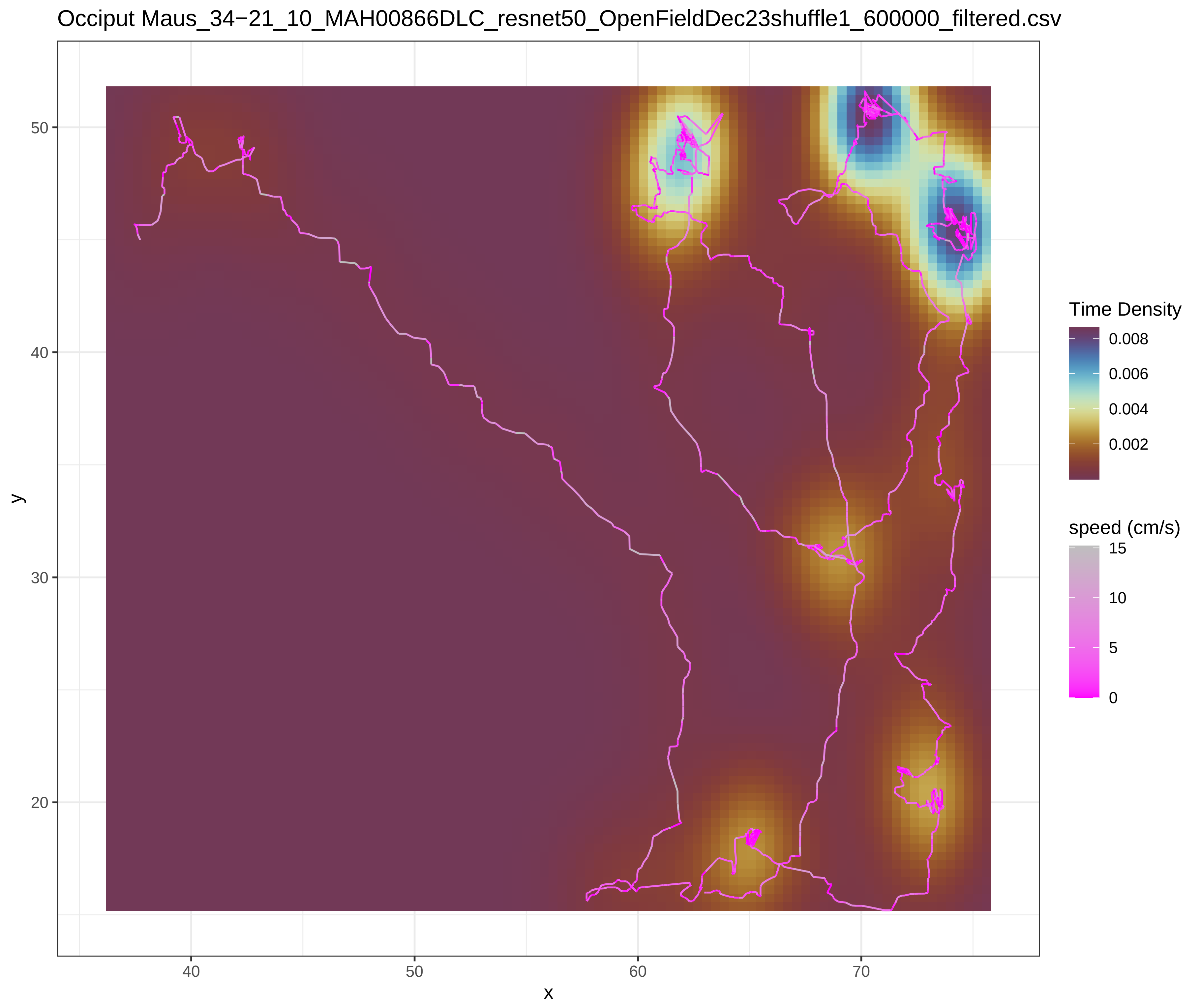
Occiput Maus_34−21_10_MAH00866DLC_resnet50_OpenFieldDec23shuffle1_600000_filtered.csv
Time Density
0.008
0.006
0.004
0.002
speed (cm/s)
15
10
5
0
y
x
50
40
30
20
40
50
60
70

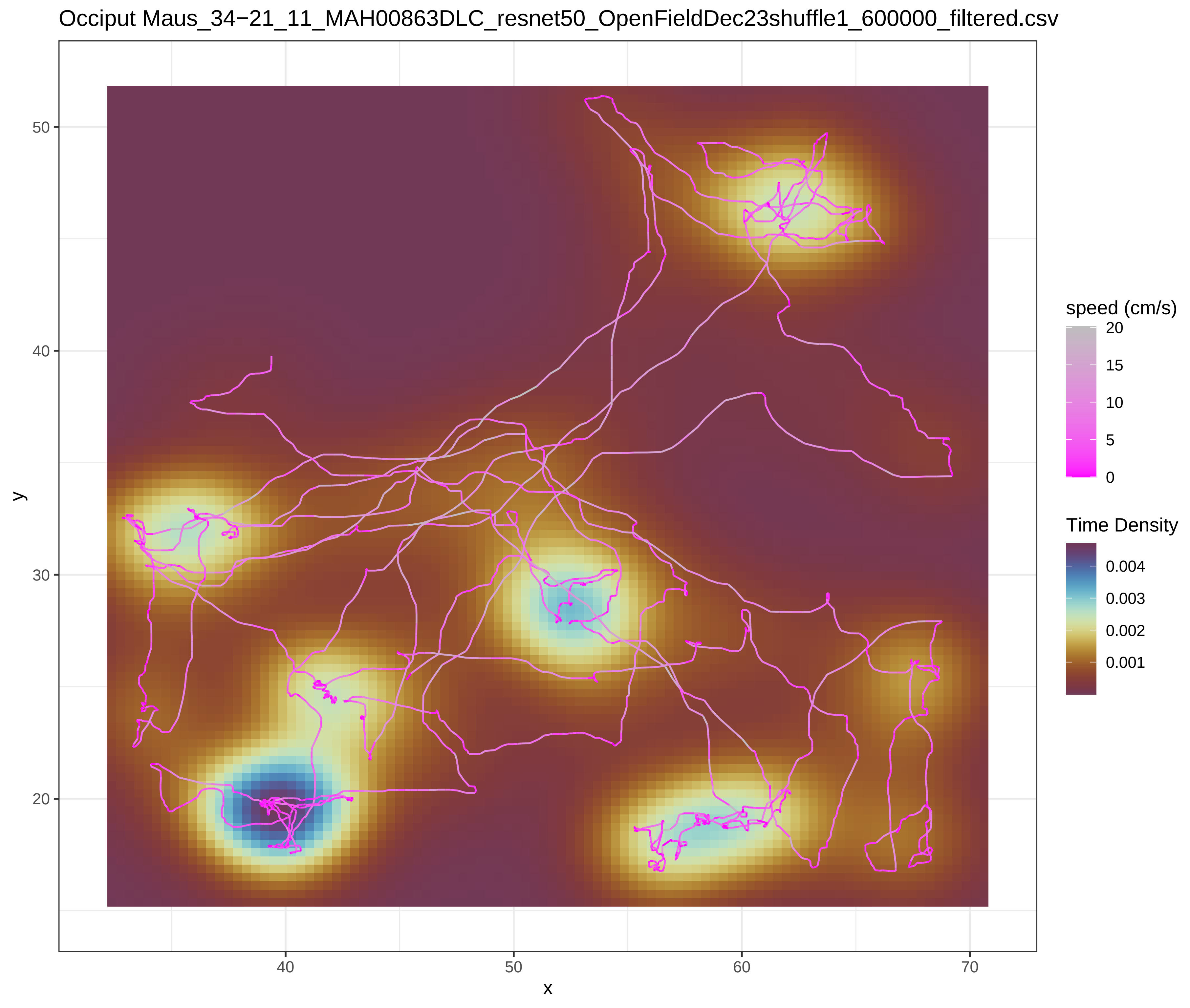
Occiput Maus_34−21_11_MAH00863DLC_resnet50_OpenFieldDec23shuffle1_600000_filtered.csv
y
x
50
40
30
20
40
50
60
70
speed (cm/s)
20
15
10
5
0
Time Density
0.004
0.003
0.002
0.001

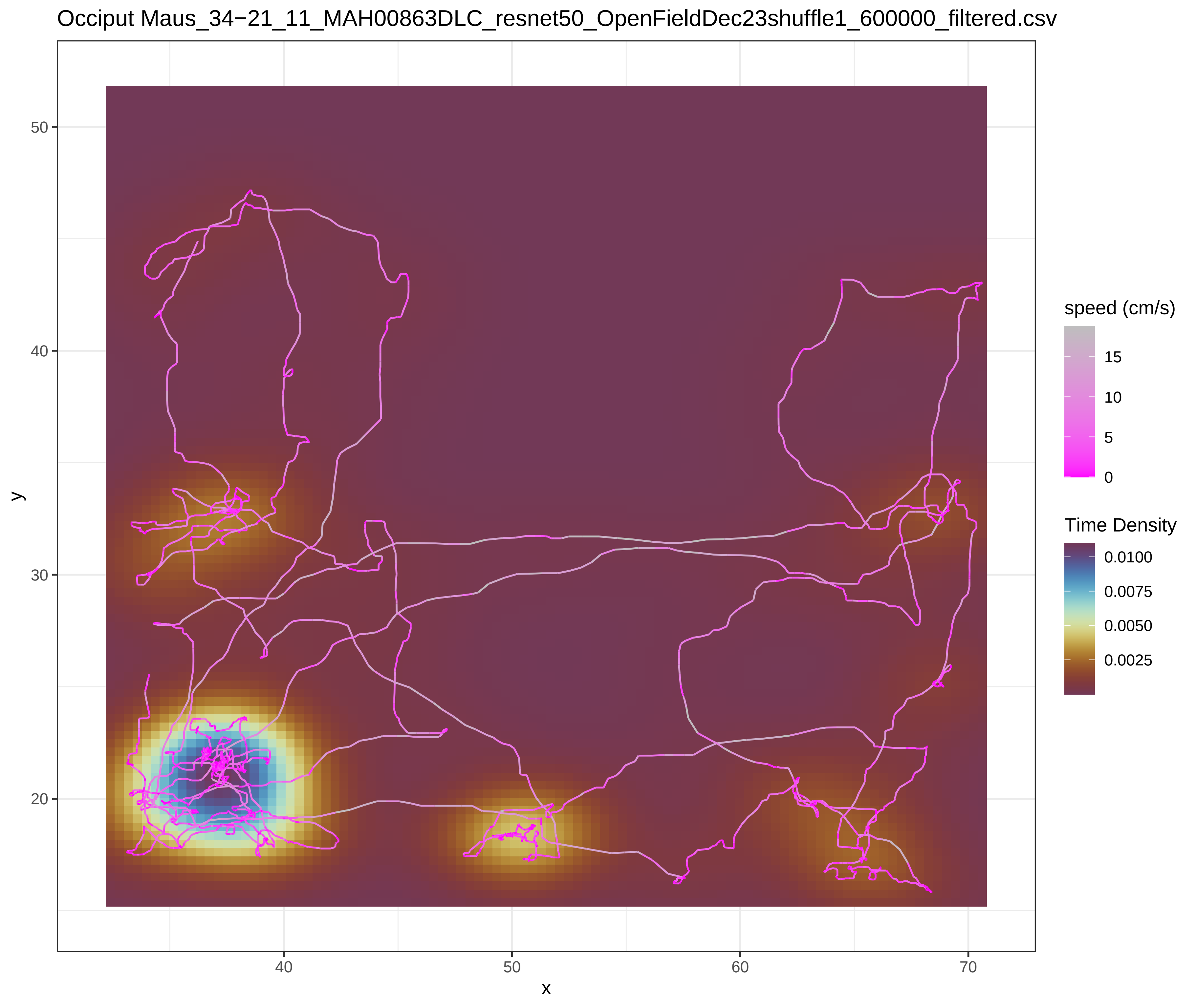
Occiput Maus_34−21_11_MAH00863DLC_resnet50_OpenFieldDec23shuffle1_600000_filtered.csv
y
x
50
40
30
20
40
50
60
70
speed (cm/s)
15
10
5
0
Time Density
0.0100
0.0075
0.0050
0.0025

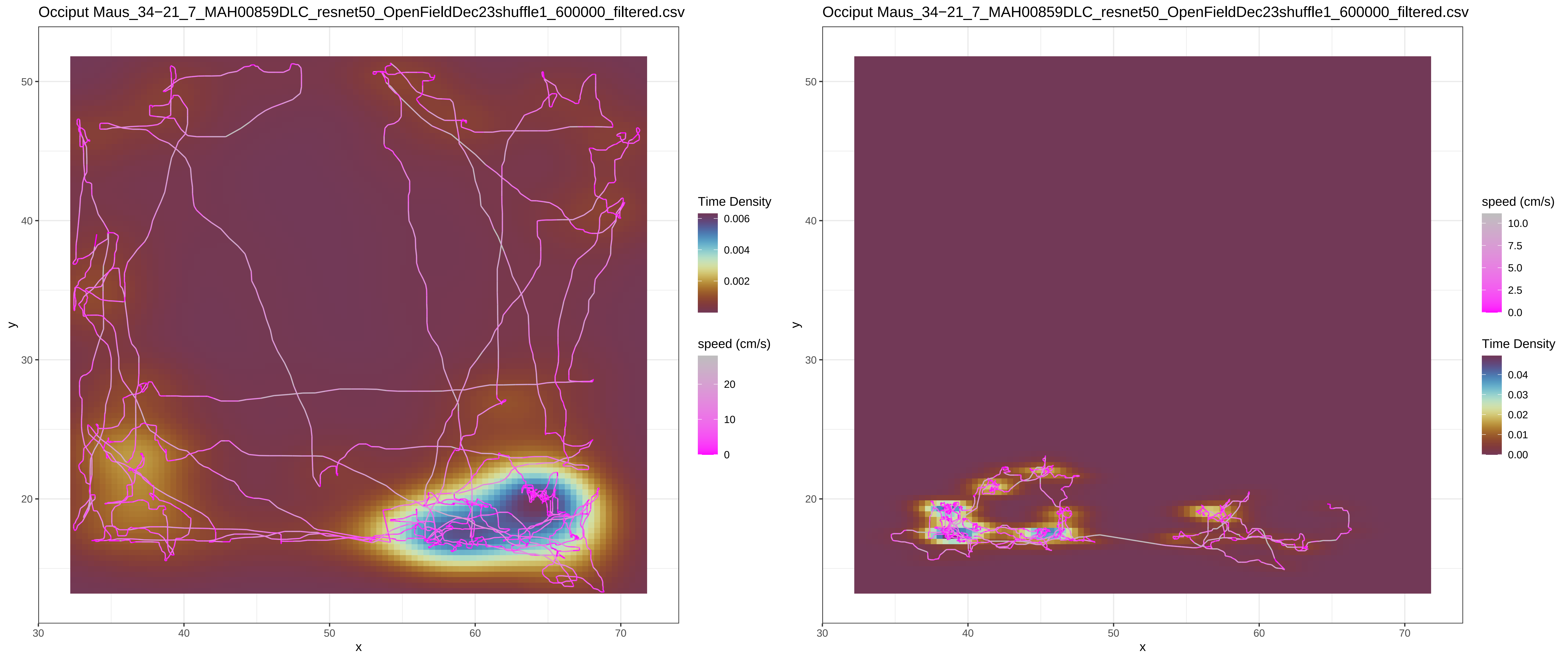

Occiput Maus_34−21_7_MAH00859DLC_resnet50_OpenFieldDec23shuffle1_600000_filtered.csv
Time Density
0.006
0.004
0.002
speed (cm/s)
20
10
0
y
x
Occiput Maus_34−21_7_MAH00859DLC_resnet50_OpenFieldDec23shuffle1_600000_filtered.csv
speed (cm/s)
10.0
7.5
5.0
2.5
0.0
Time Density
0.04
0.03
0.02
0.01
0.00

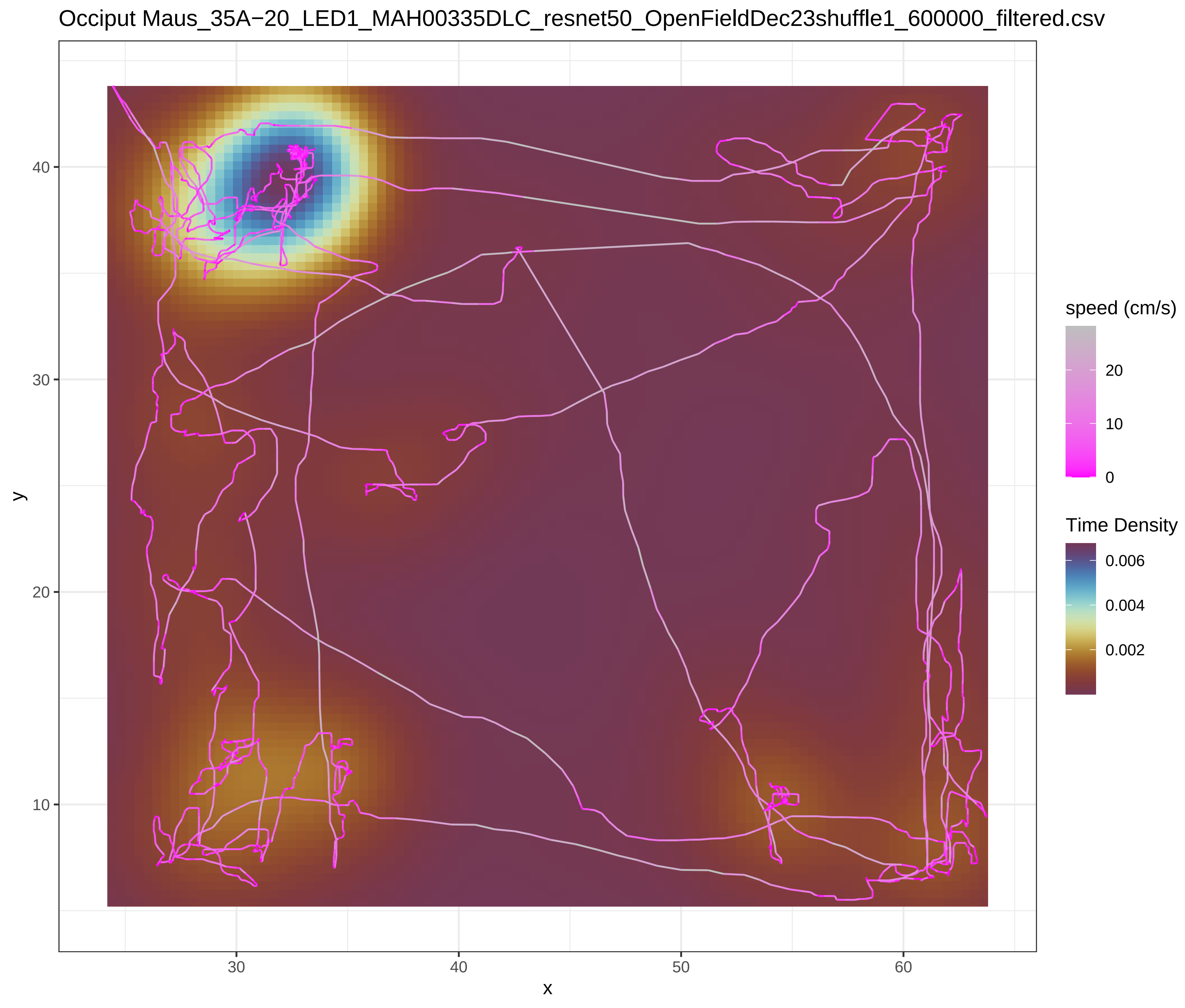
Occiput Maus_35A−20_LED1_MAH00335DLC_resnet50_OpenFieldDec23shuffle1_600000_filtered.csv
y
x
speed (cm/s)
20
10
0
Time Density
0.006
0.004
0.002
10
20
30
40
50
60

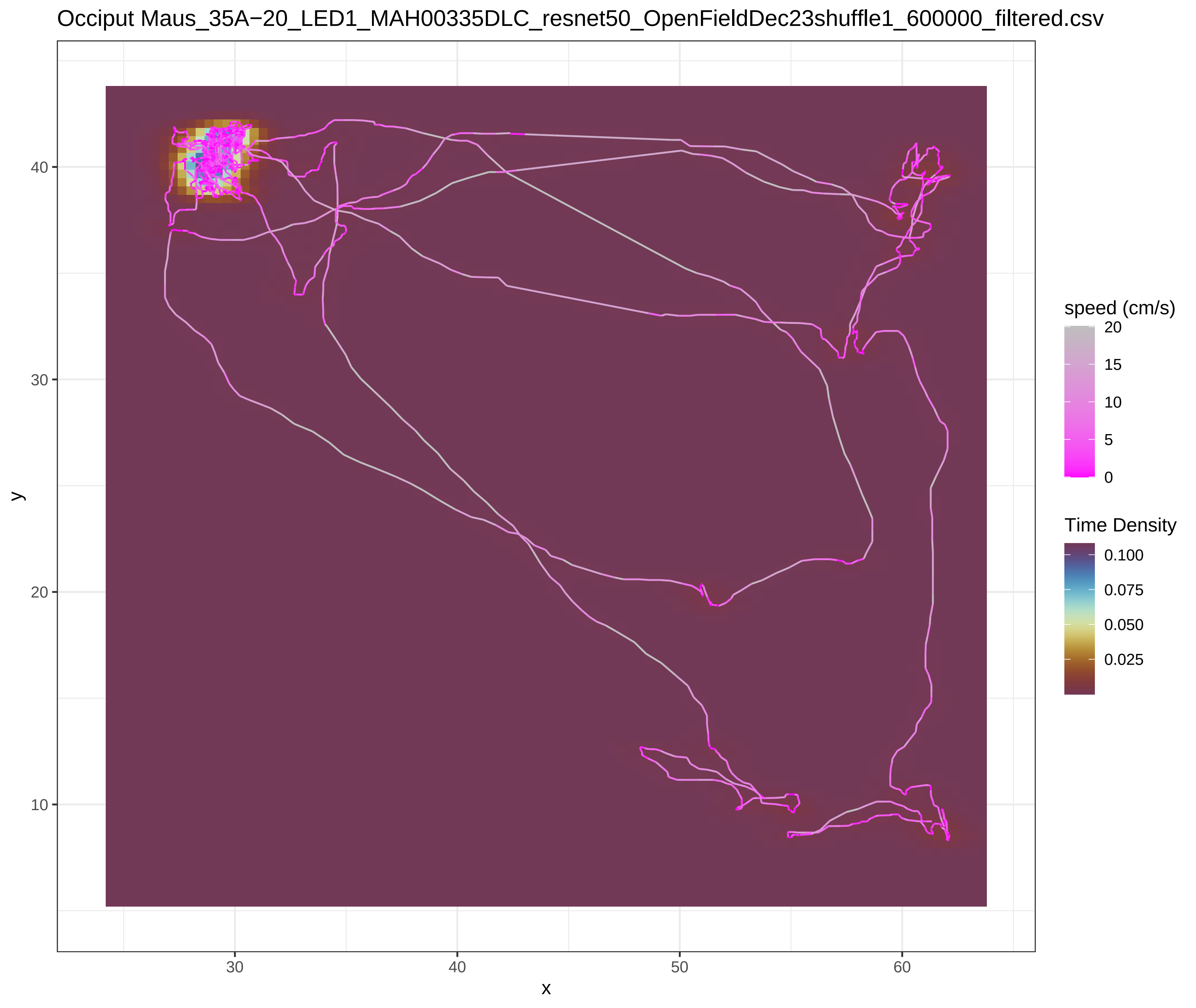
Occiput Maus_35A−20_LED1_MAH00335DLC_resnet50_OpenFieldDec23shuffle1_600000_filtered.csv
y
x
speed (cm/s)
20
15
10
5
0
Time Density
0.100
0.075
0.050
0.025
10
20
30
40
50
60

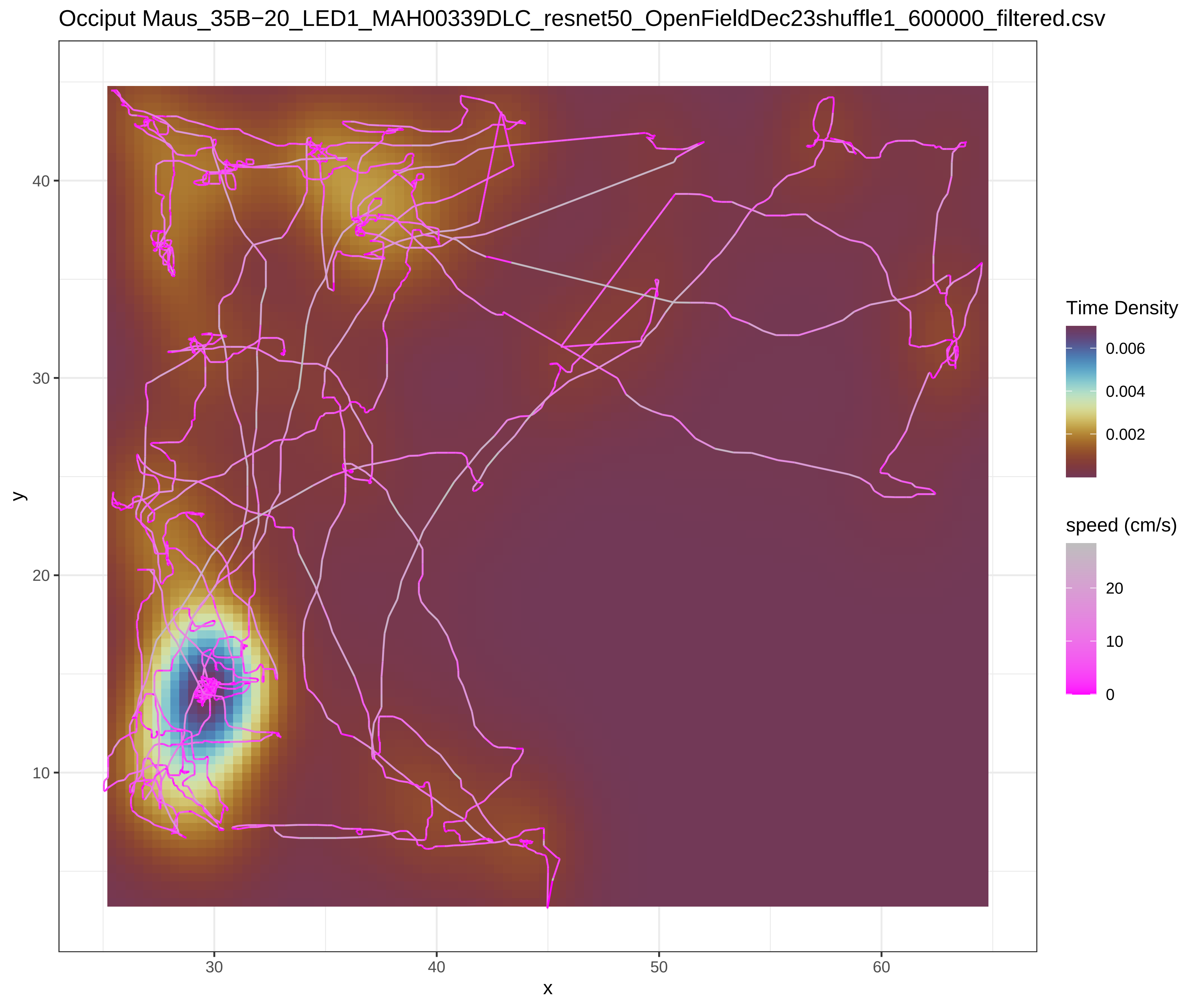
Occiput Maus_35B−20_LED1_MAH00339DLC_resnet50_OpenFieldDec23shuffle1_600000_filtered.csv
Time Density
0.006
0.004
0.002
speed (cm/s)
20
10
0
y
x
40
30
20
10
30
40
50
60

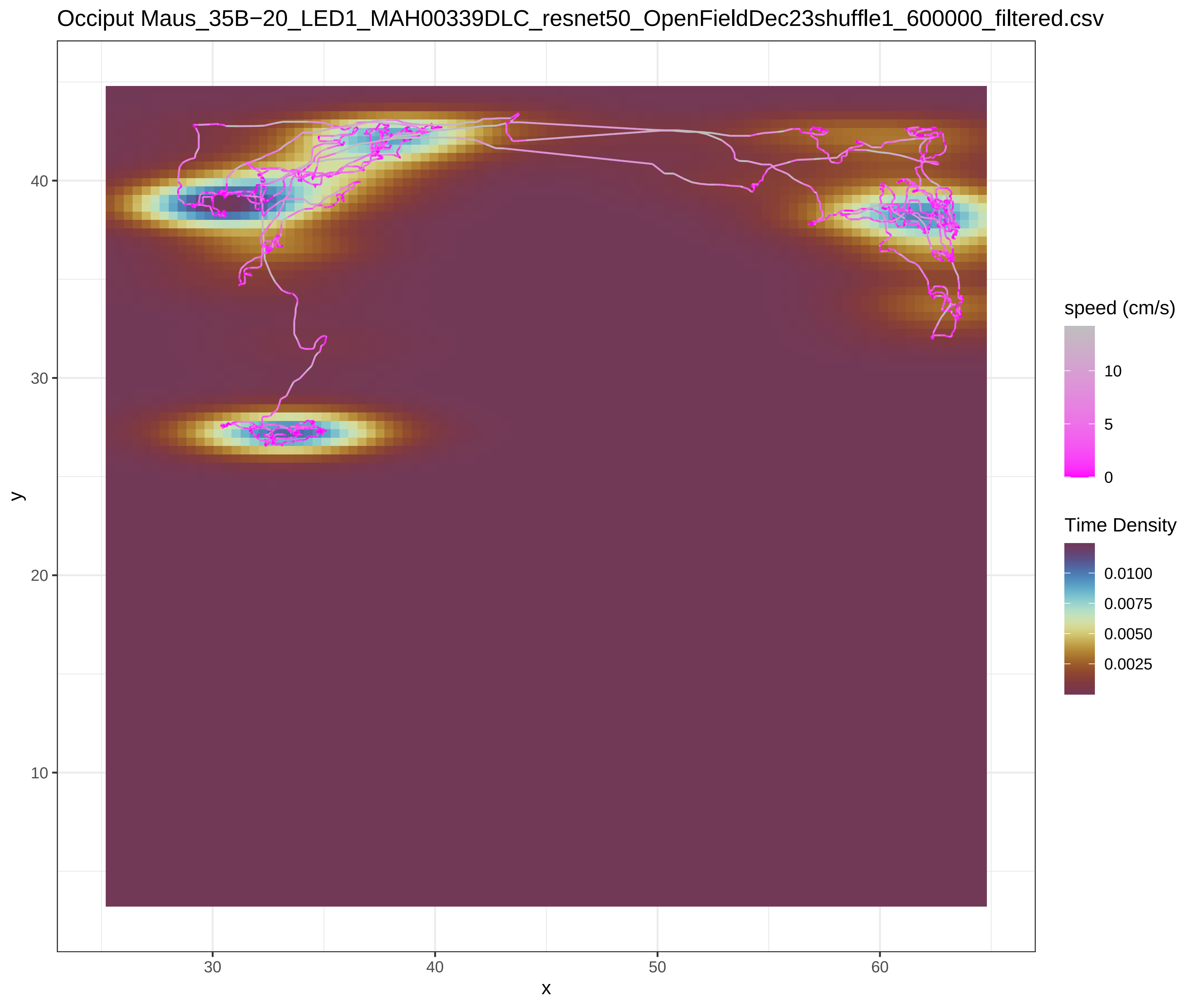
Occiput Maus_35B−20_LED1_MAH00339DLC_resnet50_OpenFieldDec23shuffle1_600000_filtered.csv
speed (cm/s)
10
5
0
Time Density
0.0100
0.0075
0.0050
0.0025
y
x
40
30
20
10
30
40
50
60

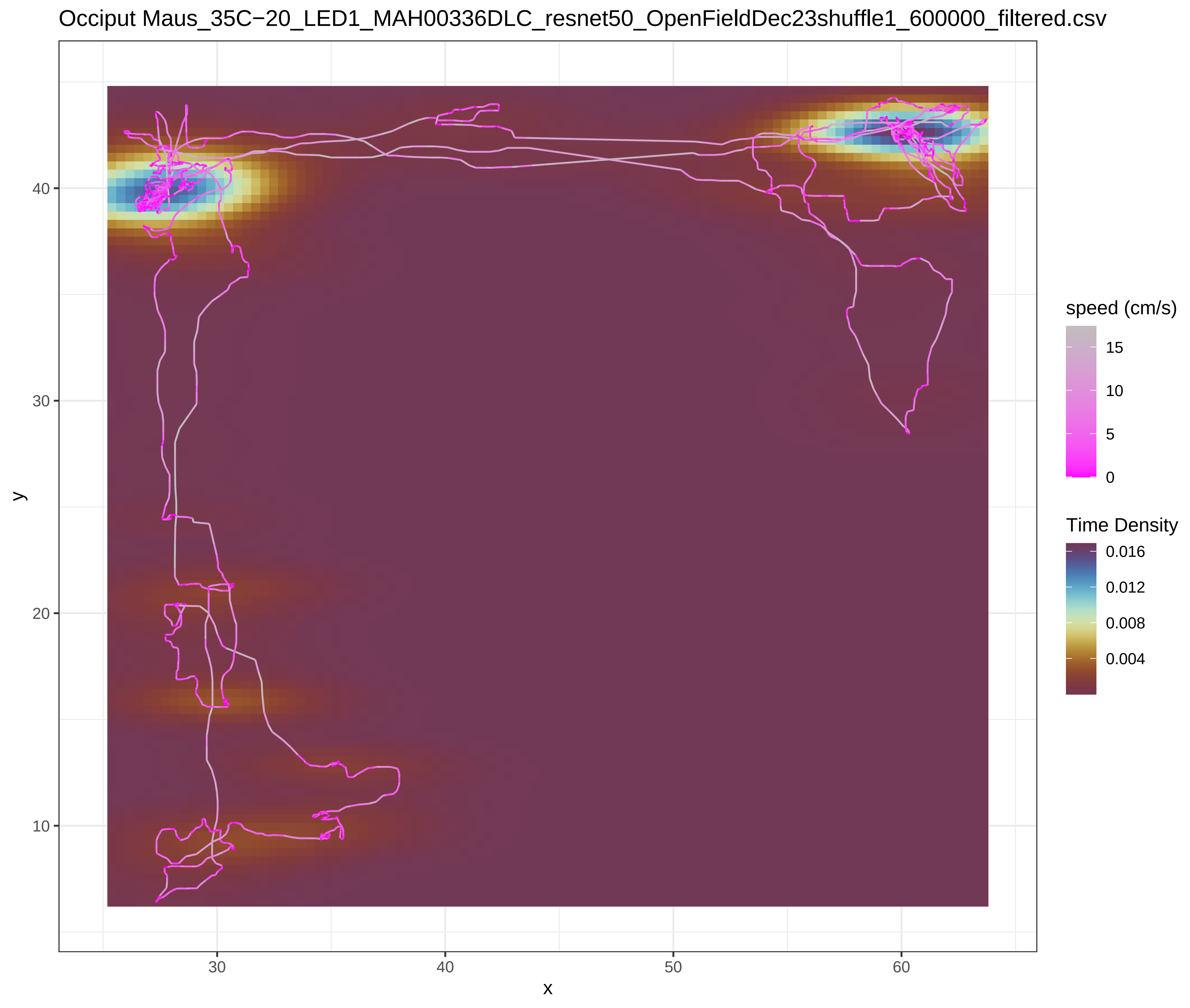

Occiput Maus_35C−20_LED1_MAH00336DLC_resnet50_OpenFieldDec23shuffle1_600000_filtered.csv
y
x
10
20
30
40
30
40
50
60
speed (cm/s)
15
10
5
0
Time Density
0.016
0.012
0.008
0.004


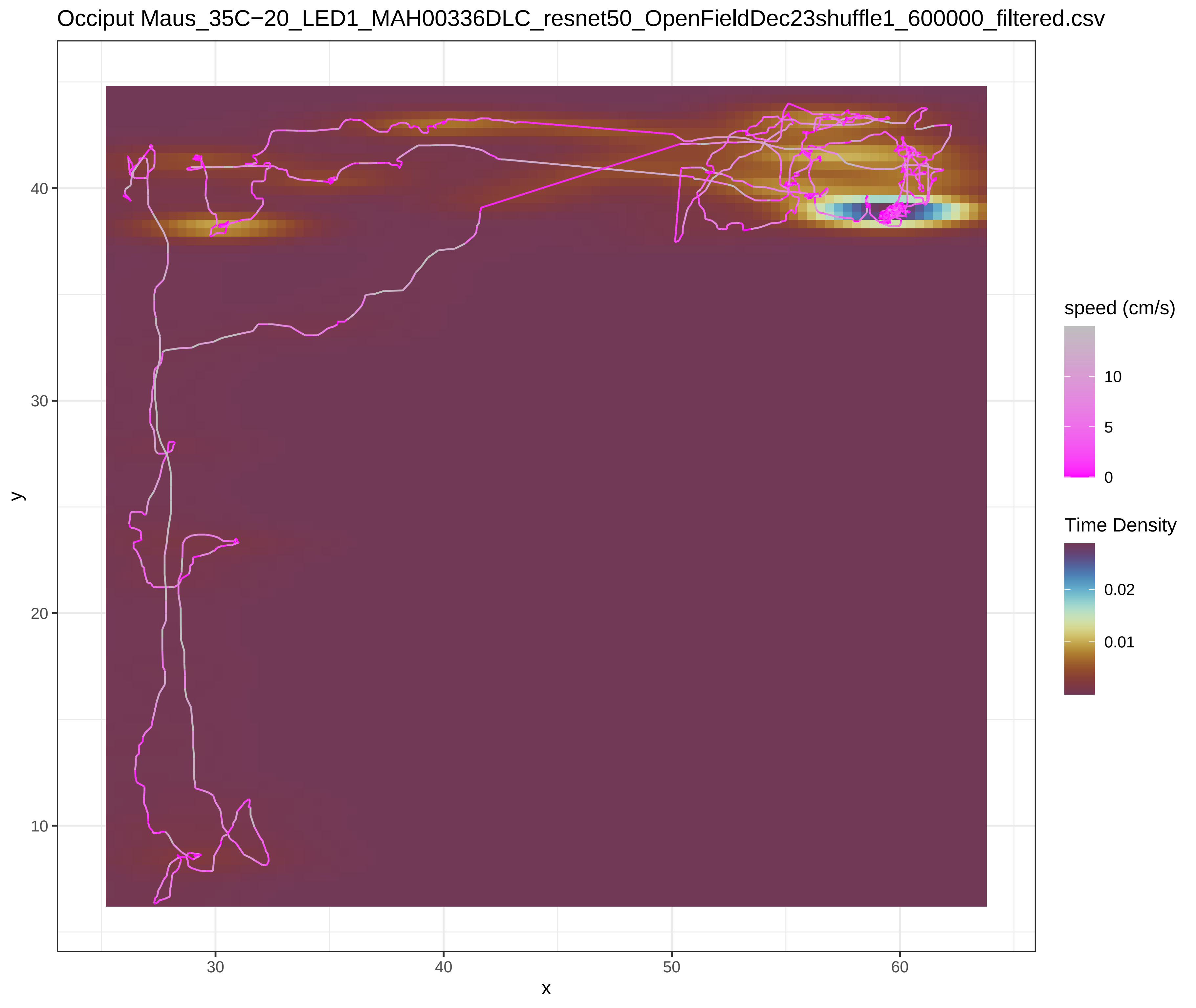

Occiput Maus_35C−20_LED1_MAH00336DLC_resnet50_OpenFieldDec23shuffle1_600000_filtered.csv
y
x
10
20
30
40
30
40
50
60
speed (cm/s)
10
5
0
Time Density
0.02
0.01

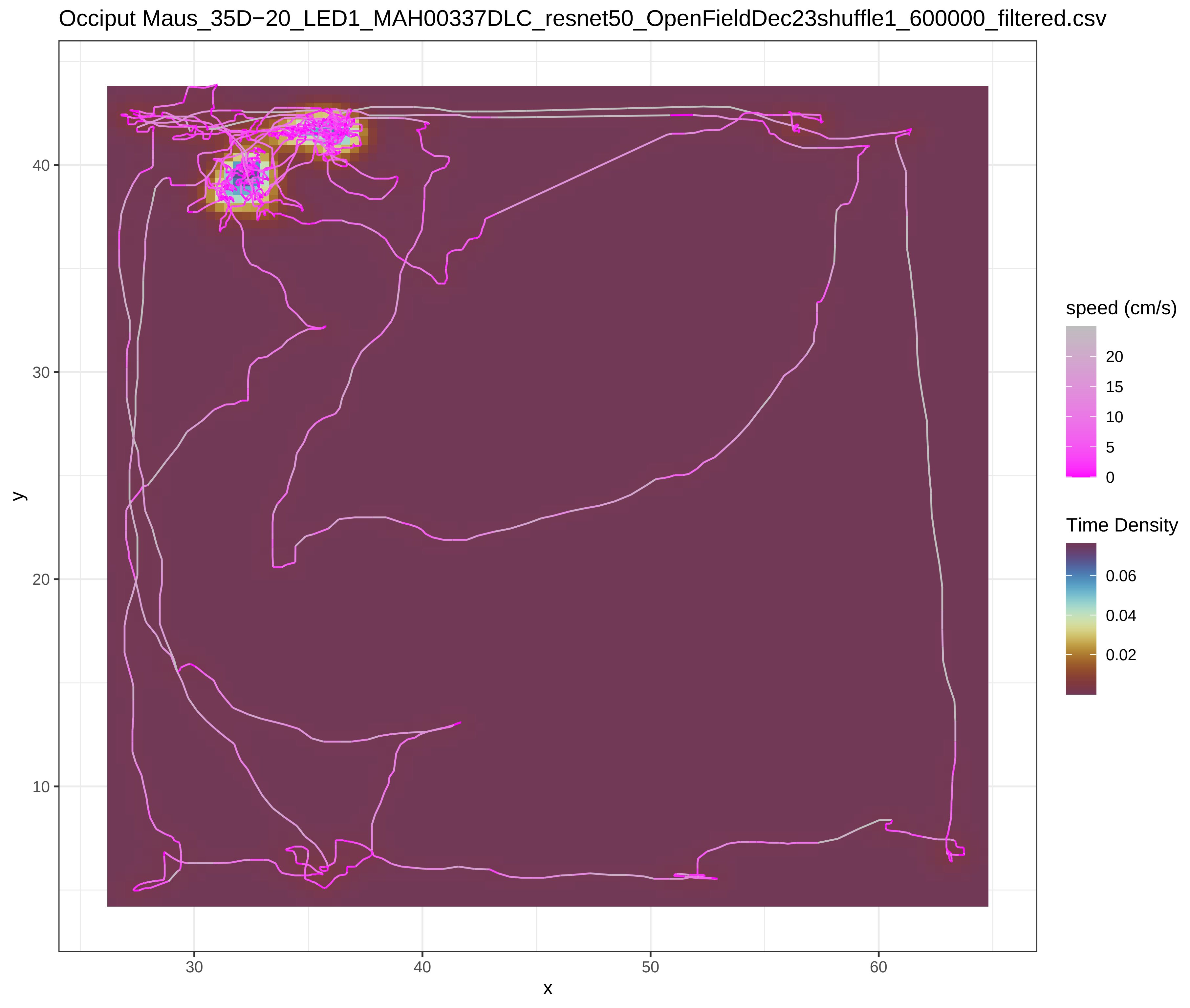

Occiput Maus_35D−20_LED1_MAH00337DLC_resnet50_OpenFieldDec23shuffle1_600000_filtered.csv
speed (cm/s)
20
15
10
5
0
Time Density
0.06
0.04
0.02
x
y
30
40
50
60
10
20
30
40


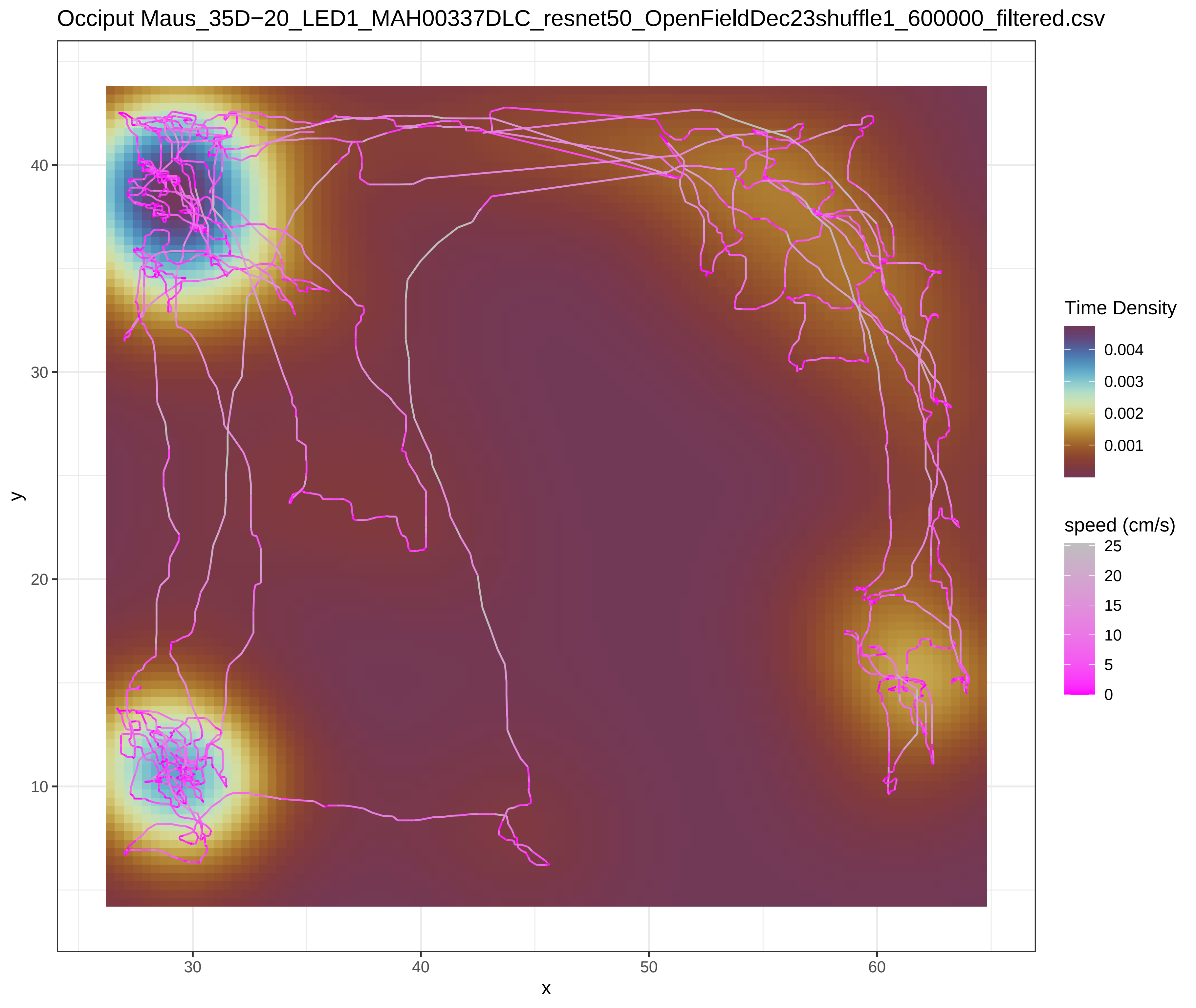

Occiput Maus_35D−20_LED1_MAH00337DLC_resnet50_OpenFieldDec23shuffle1_600000_filtered.csv
Time Density
0.004
0.003
0.002
0.001
speed (cm/s)
25
20
15
10
5
0
x
y
30
40
50
60
10
20
30
40

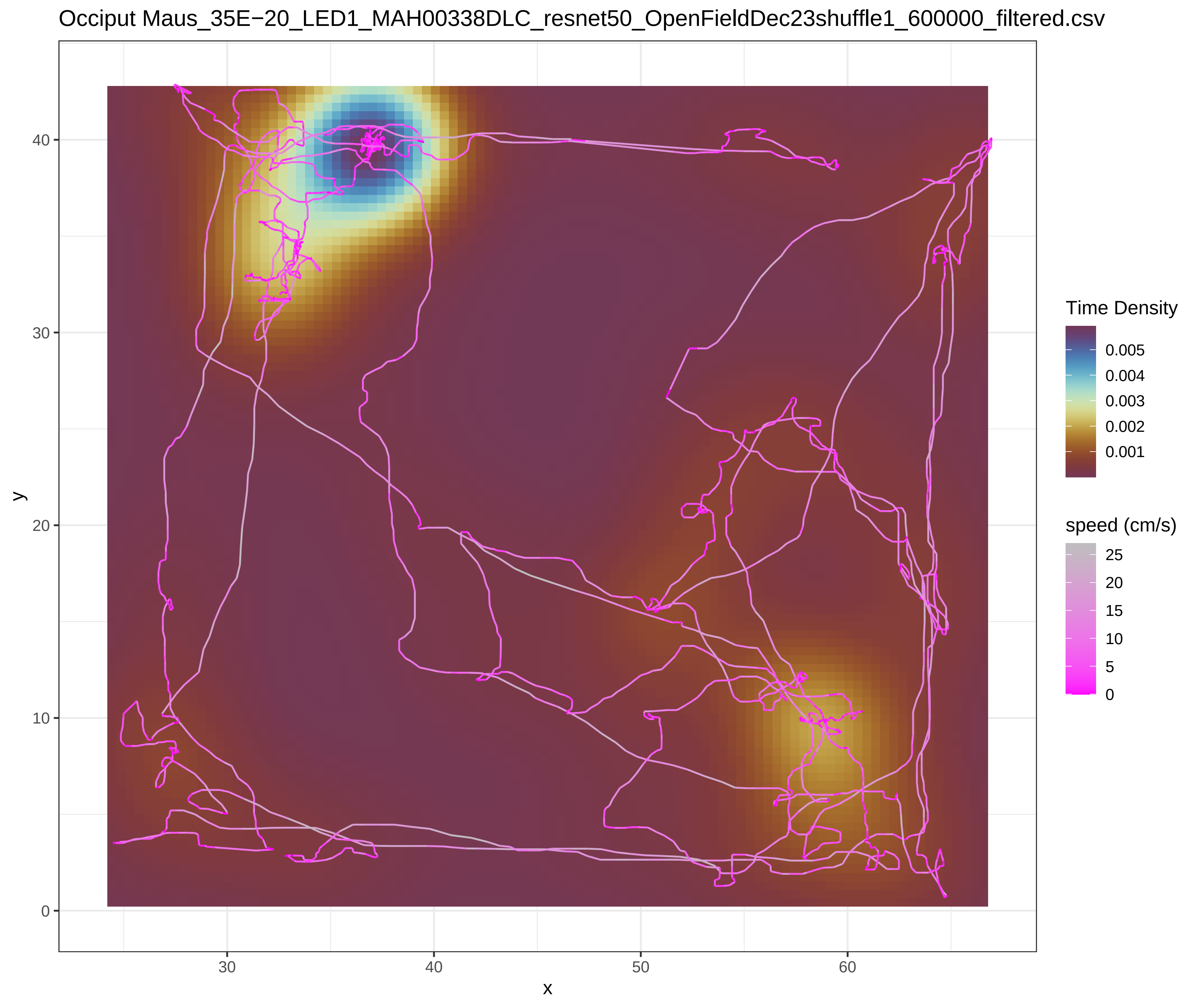
Occiput Maus_35E−20_LED1_MAH00338DLC_resnet50_OpenFieldDec23shuffle1_600000_filtered.csv
Time Density
0.005
0.004
0.003
0.002
0.001
speed (cm/s)
25
20
15
10
5
0
y
x

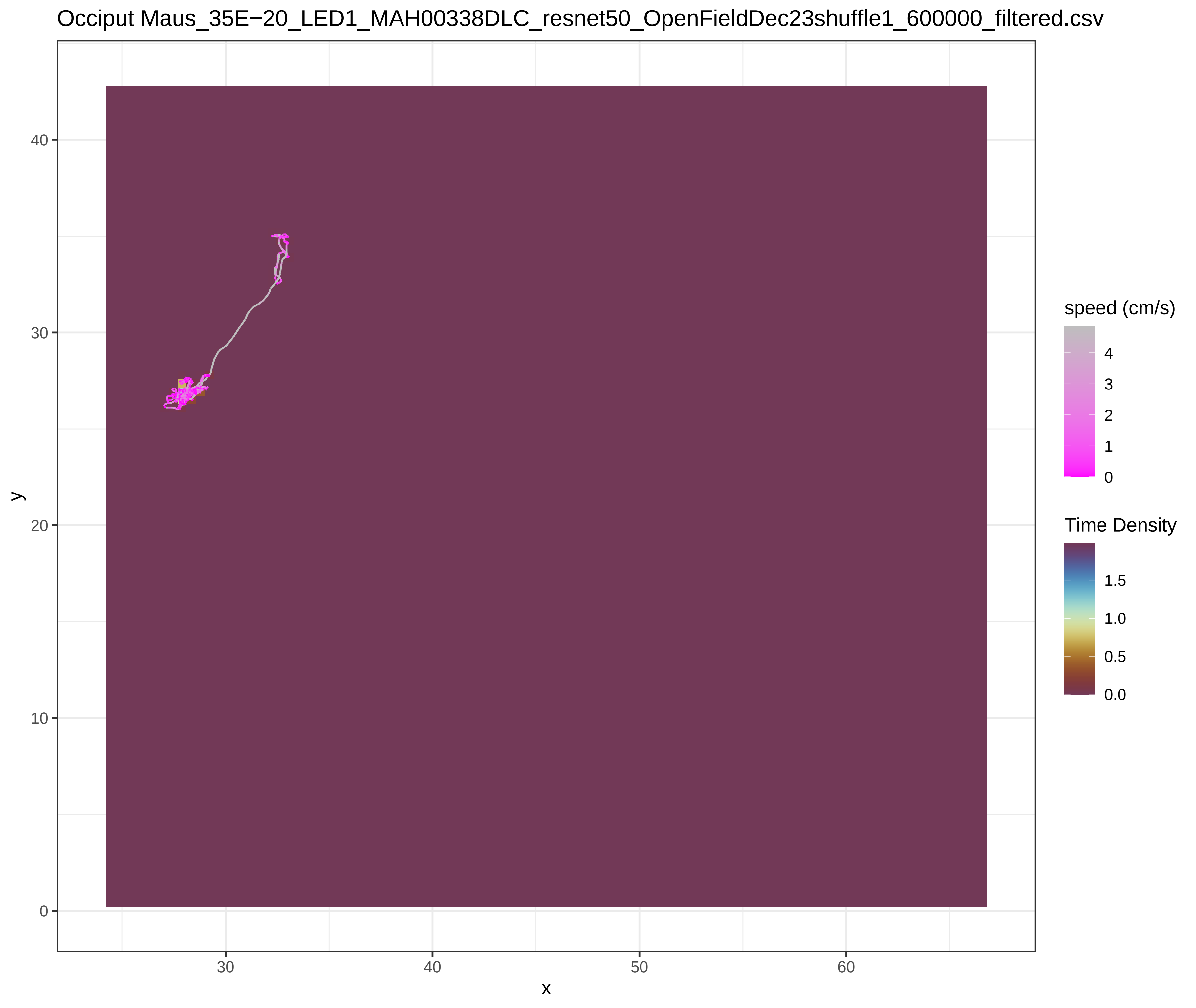
Occiput Maus_35E−20_LED1_MAH00338DLC_resnet50_OpenFieldDec23shuffle1_600000_filtered.csv
speed (cm/s)
4
3
2
1
0
Time Density
1.5
1.0
0.5
0.0
y
x

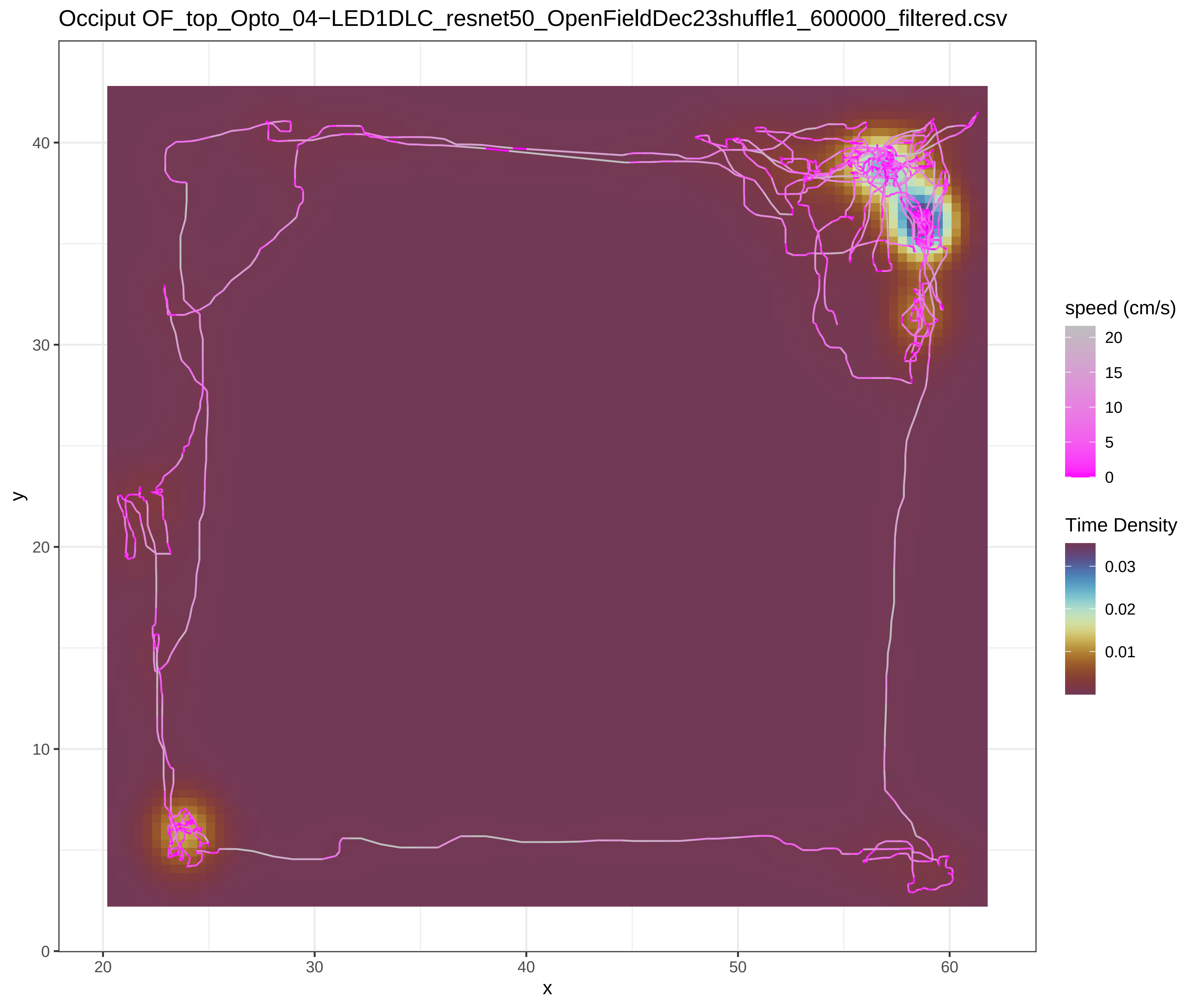
Occiput OF_top_Opto_04−LED1DLC_resnet50_OpenFieldDec23shuffle1_600000_filtered.csv
y
x
speed (cm/s)
20
15
10
5
0
Time Density
0.03
0.02
0.01


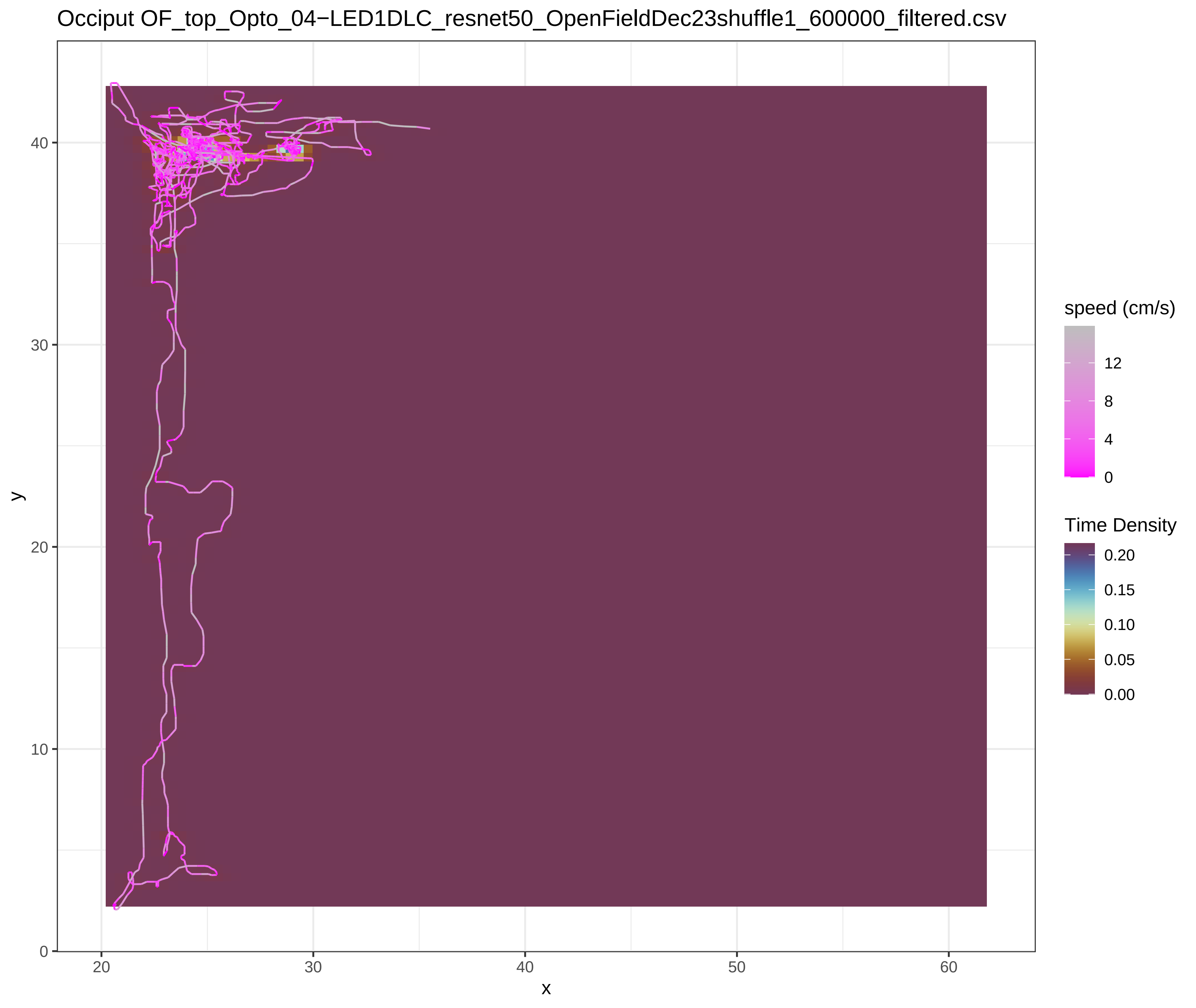
Occiput OF_top_Opto_04−LED1DLC_resnet50_OpenFieldDec23shuffle1_600000_filtered.csv
y
x
speed (cm/s)
12
8
4
0
Time Density
0.20
0.15
0.10
0.05
0.00

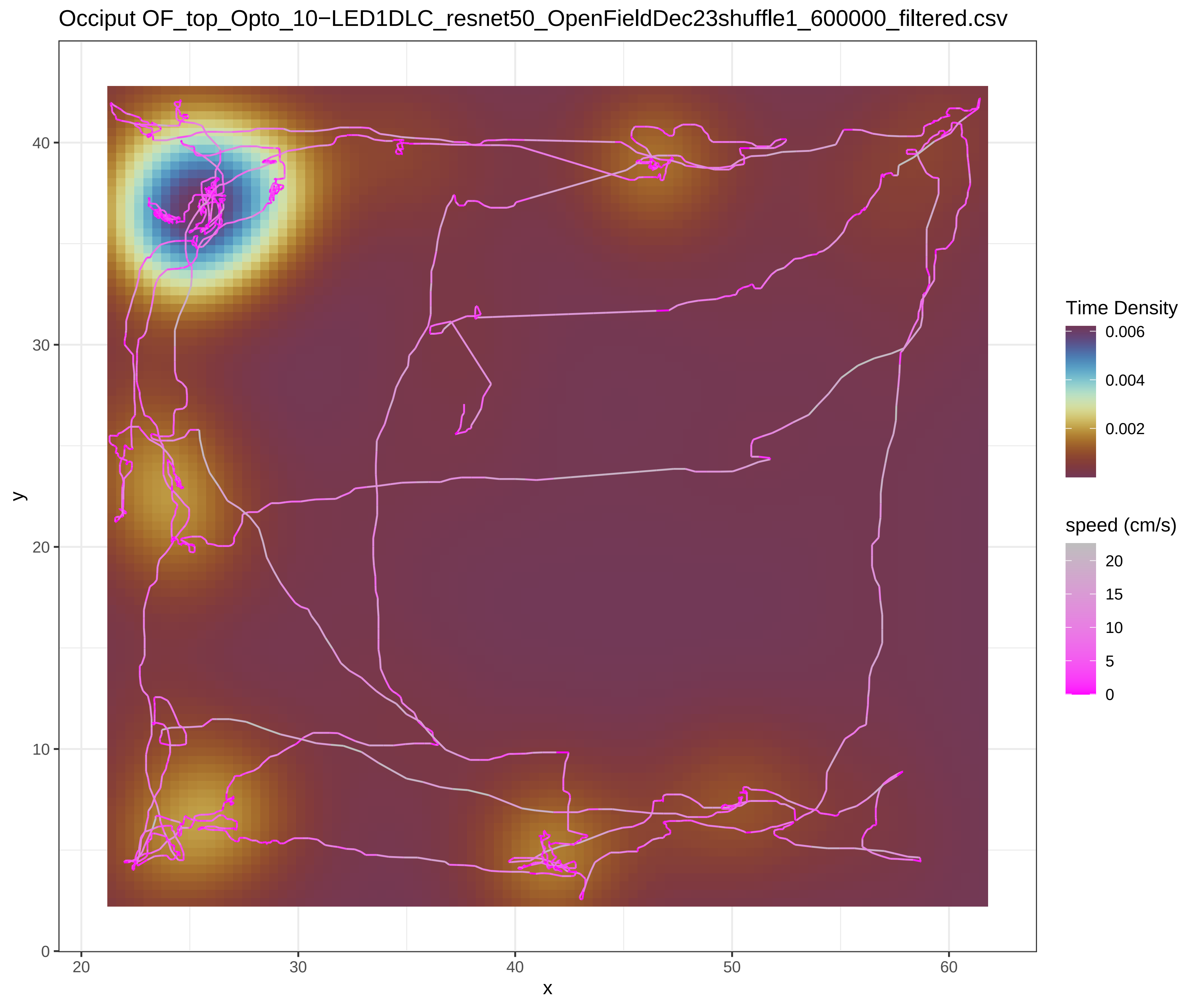
Occiput OF_top_Opto_10−LED1DLC_resnet50_OpenFieldDec23shuffle1_600000_filtered.csv
Time Density
0.006
0.004
0.002
speed (cm/s)
20
15
10
5
0
y
x
40
30
20
10
0
20
30
40
50
60

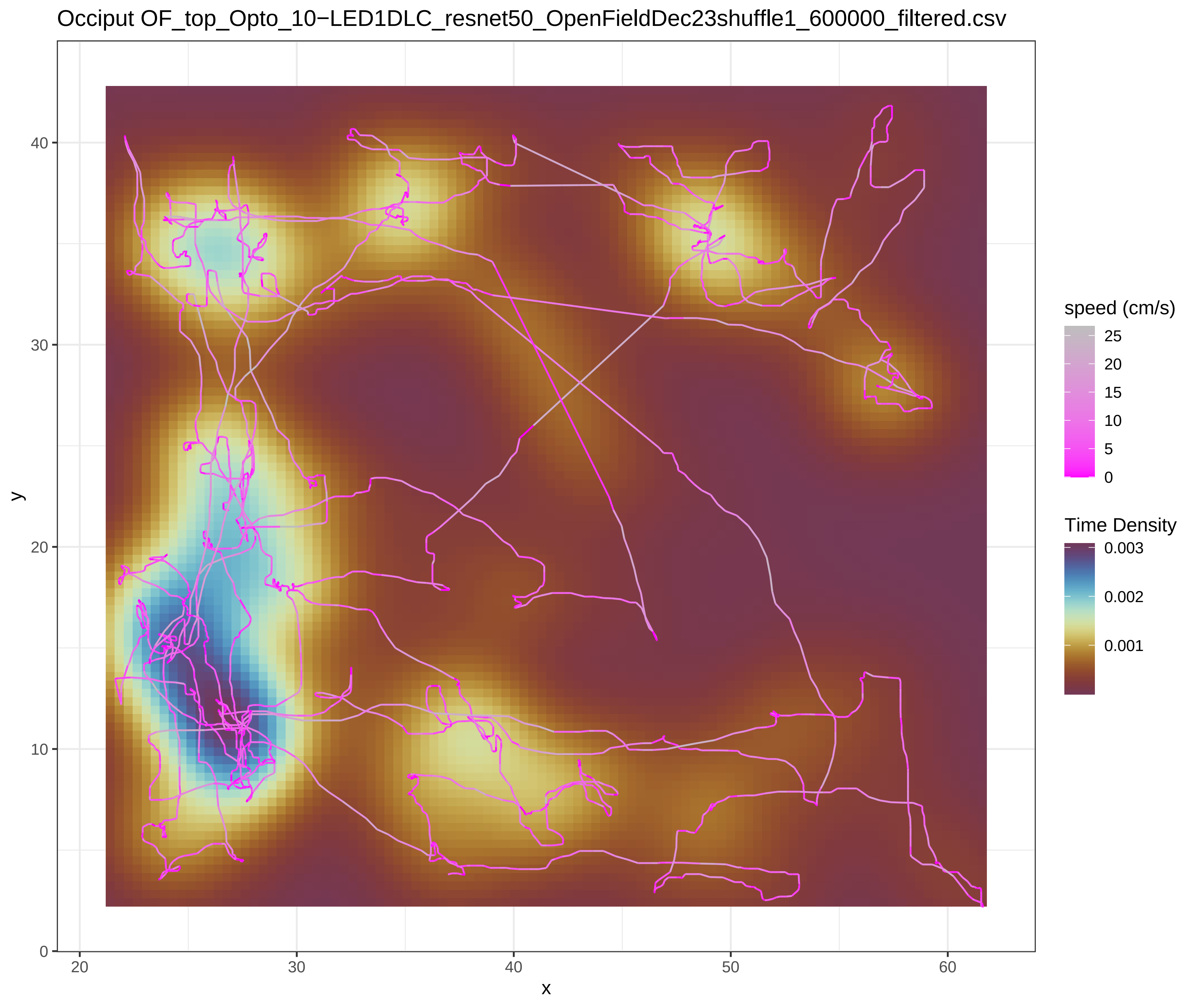
Occiput OF_top_Opto_10−LED1DLC_resnet50_OpenFieldDec23shuffle1_600000_filtered.csv
speed (cm/s)
25
20
15
10
5
0
Time Density
0.003
0.002
0.001
y
x
40
30
20
10
0
20
30
40
50
60

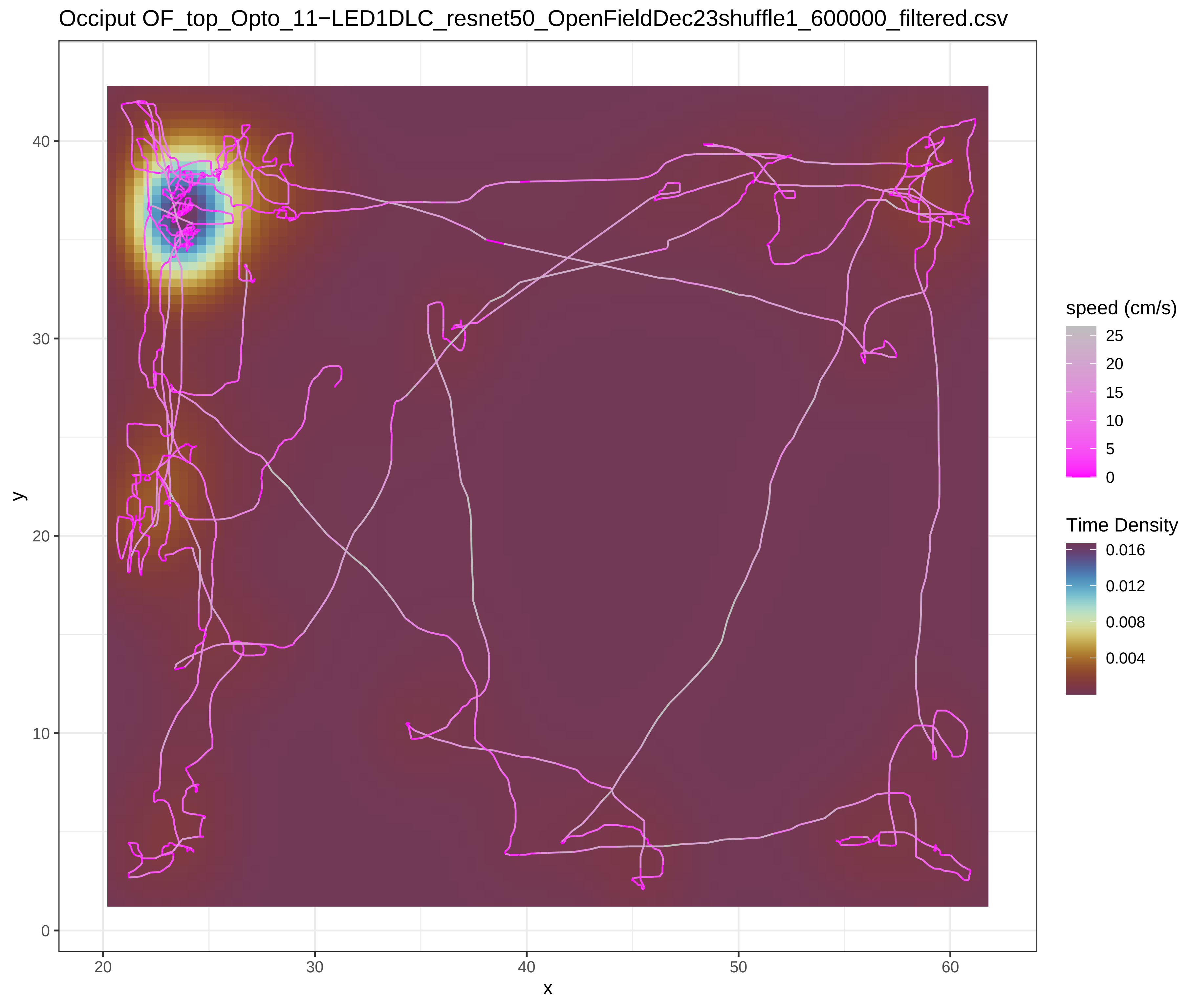
Occiput OF_top_Opto_11−LED1DLC_resnet50_OpenFieldDec23shuffle1_600000_filtered.csv
speed (cm/s)
25
20
15
10
5
0
Time Density
0.016
0.012
0.008
0.004
y
x
40
30
20
10
0
20
30
40
50
60

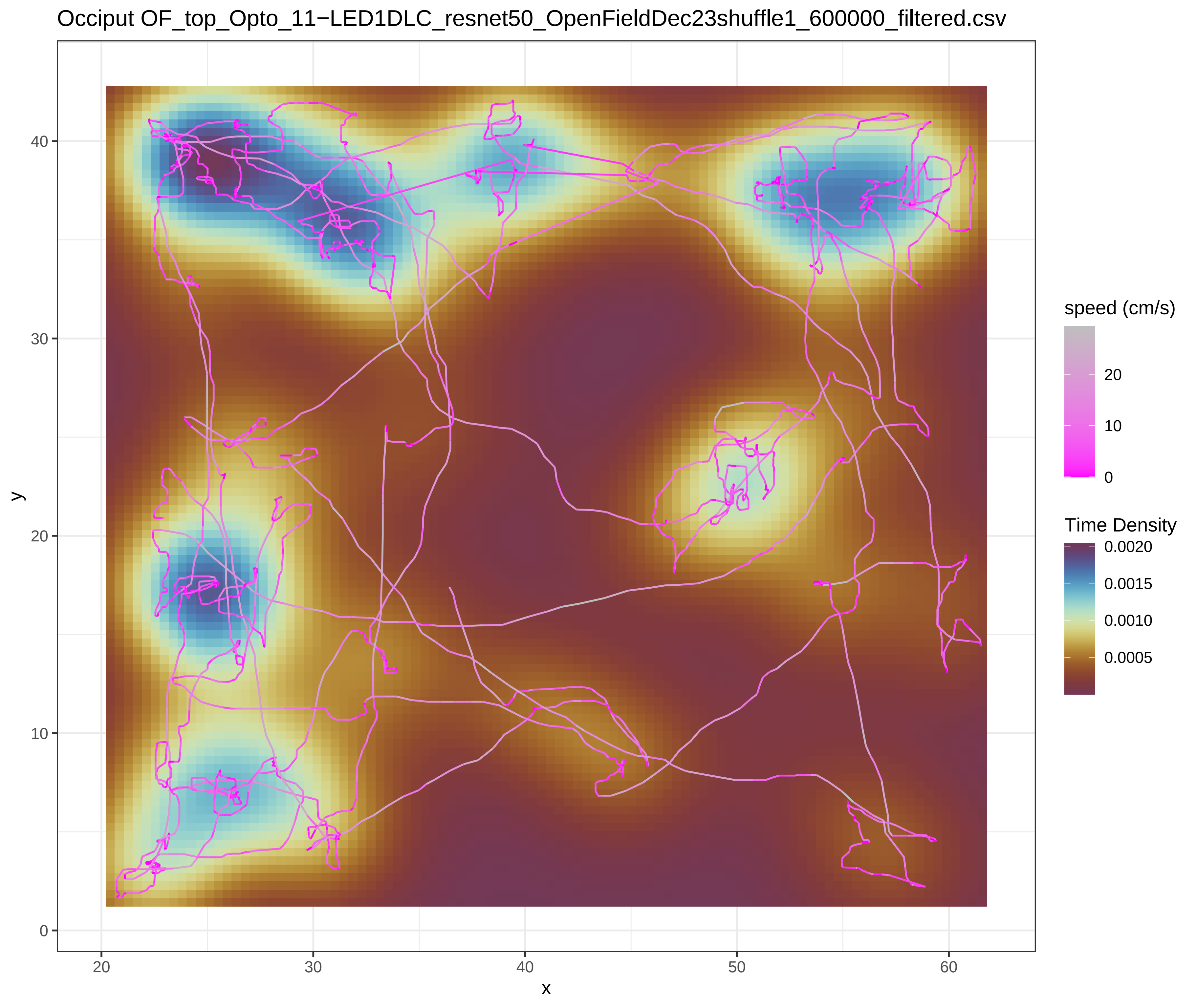
Occiput OF_top_Opto_11−LED1DLC_resnet50_OpenFieldDec23shuffle1_600000_filtered.csv
speed (cm/s)
20
10
0
Time Density
0.0020
0.0015
0.0010
0.0005
y
x
40
30
20
10
0
20
30
40
50
60

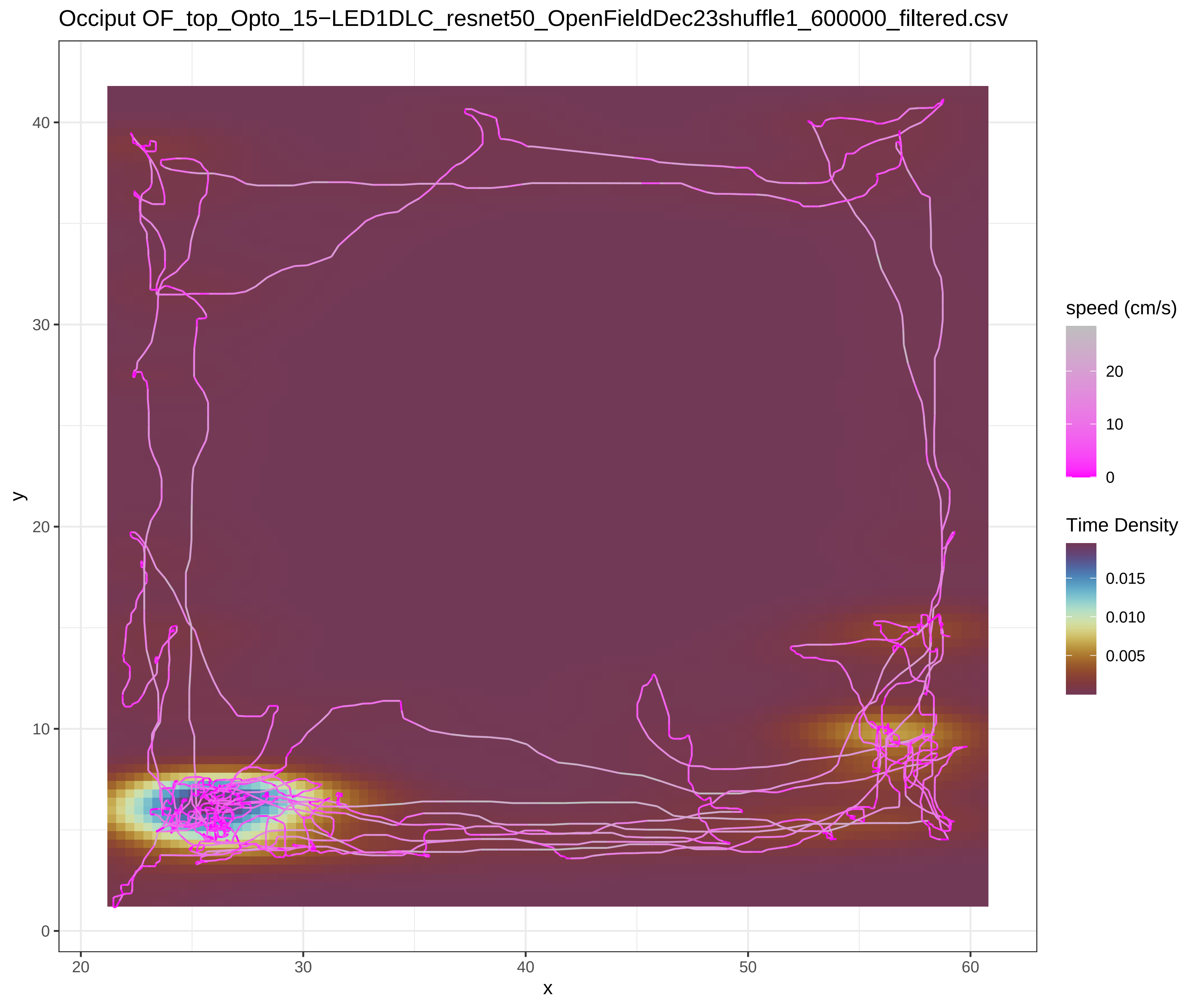
Occiput OF_top_Opto_15−LED1DLC_resnet50_OpenFieldDec23shuffle1_600000_filtered.csv
speed (cm/s)
20
10
0
Time Density
0.015
0.010
0.005
y
x
0
10
20
30
40
20
30
40
50
60

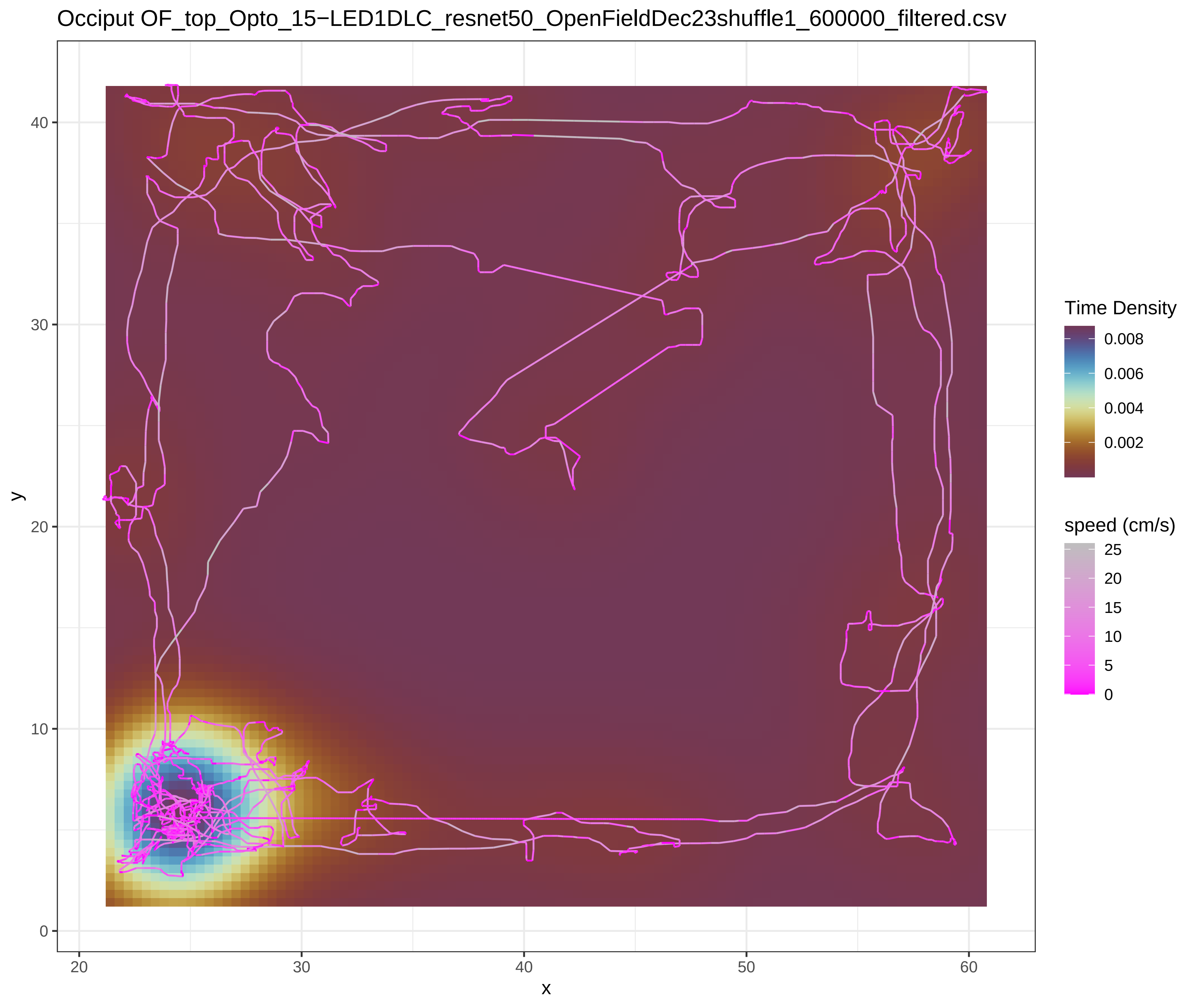
Occiput OF_top_Opto_15−LED1DLC_resnet50_OpenFieldDec23shuffle1_600000_filtered.csv
Time Density
0.008
0.006
0.004
0.002
speed (cm/s)
25
20
15
10
5
0
y
x
0
10
20
30
40
20
30
40
50
60

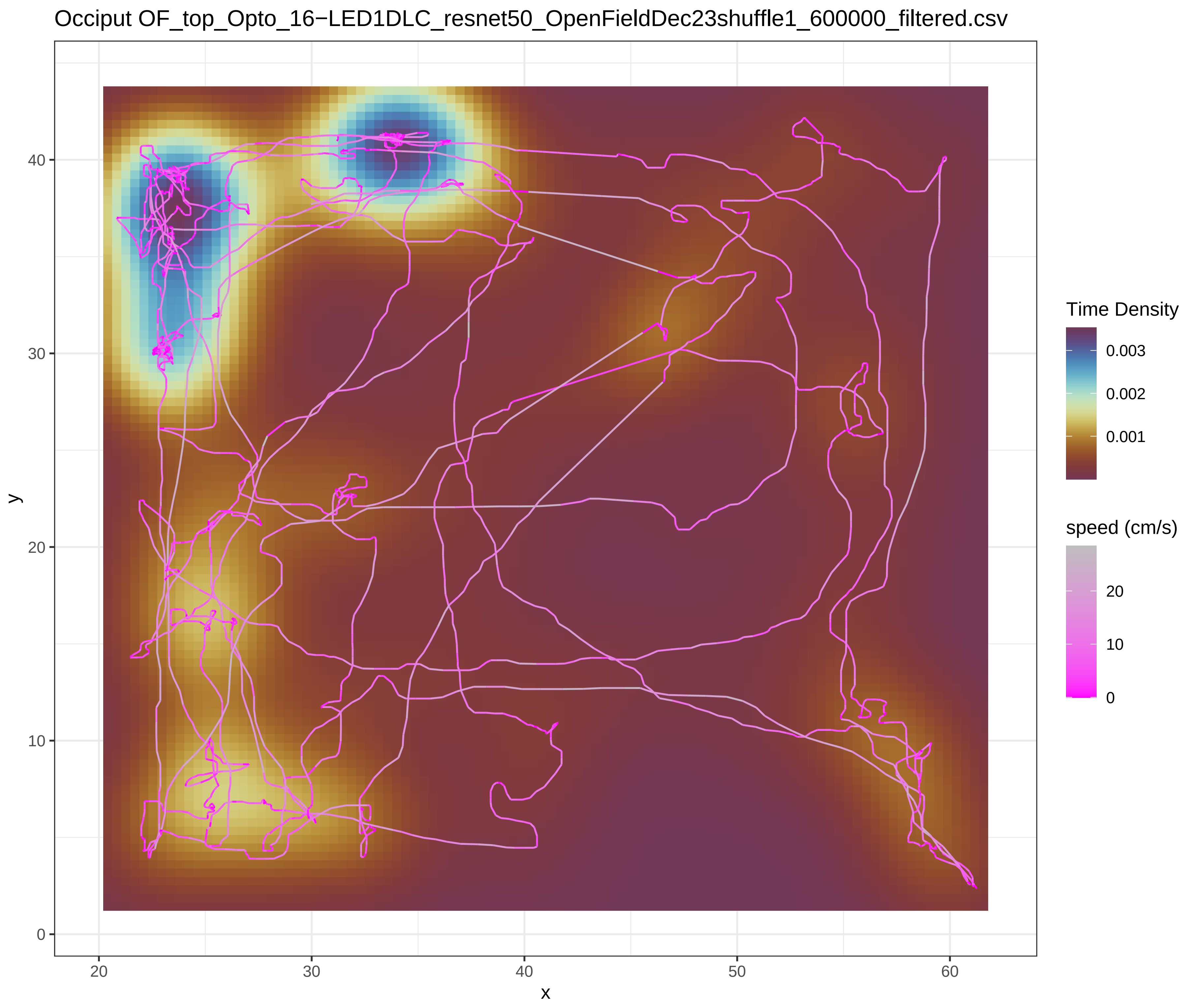

Occiput OF_top_Opto_16−LED1DLC_resnet50_OpenFieldDec23shuffle1_600000_filtered.csv
Time Density
0.003
0.002
0.001
speed (cm/s)
20
10
0
y
x


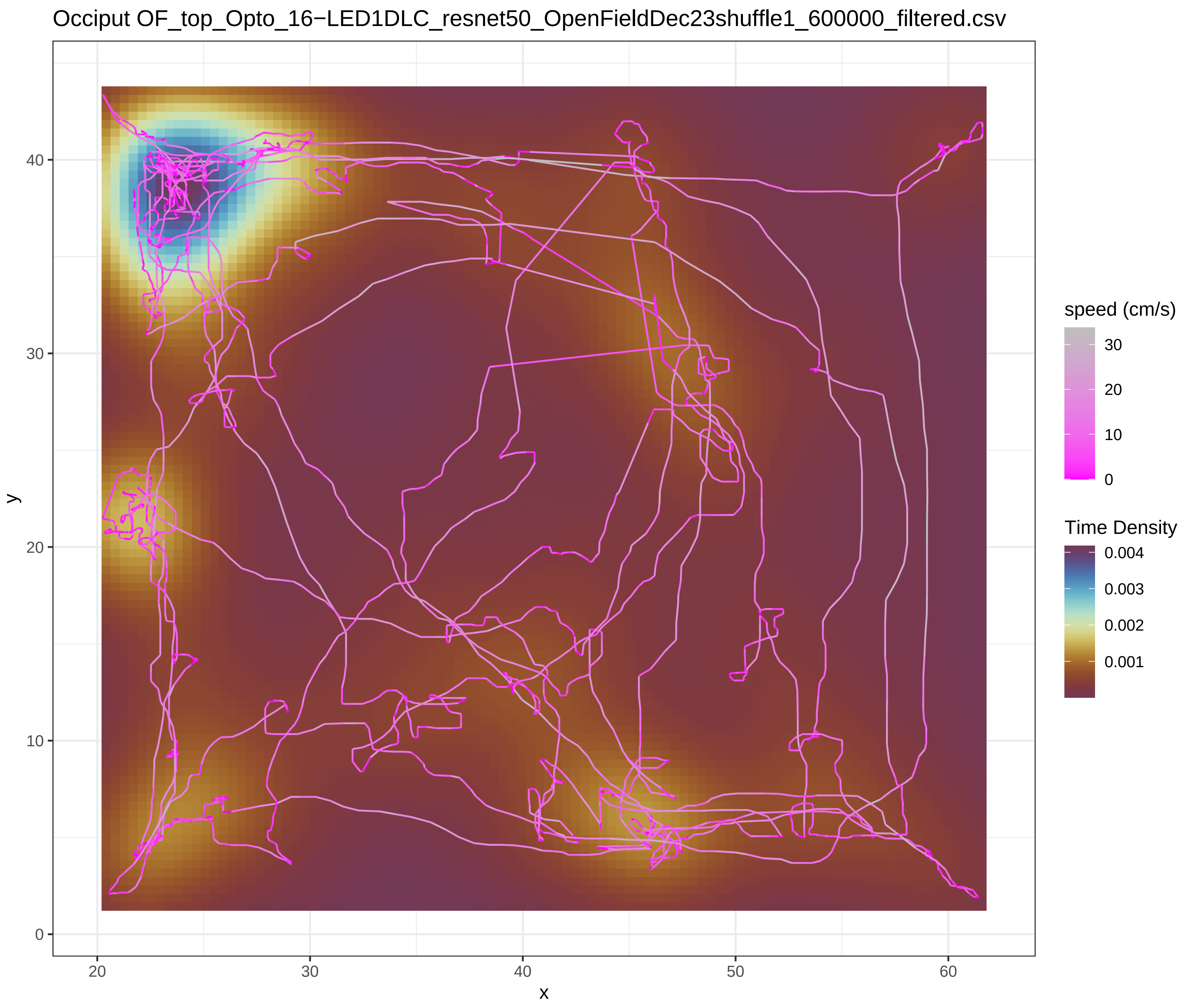

Occiput OF_top_Opto_16−LED1DLC_resnet50_OpenFieldDec23shuffle1_600000_filtered.csv
speed (cm/s)
30
20
10
0
Time Density
0.004
0.003
0.002
0.001
y
x

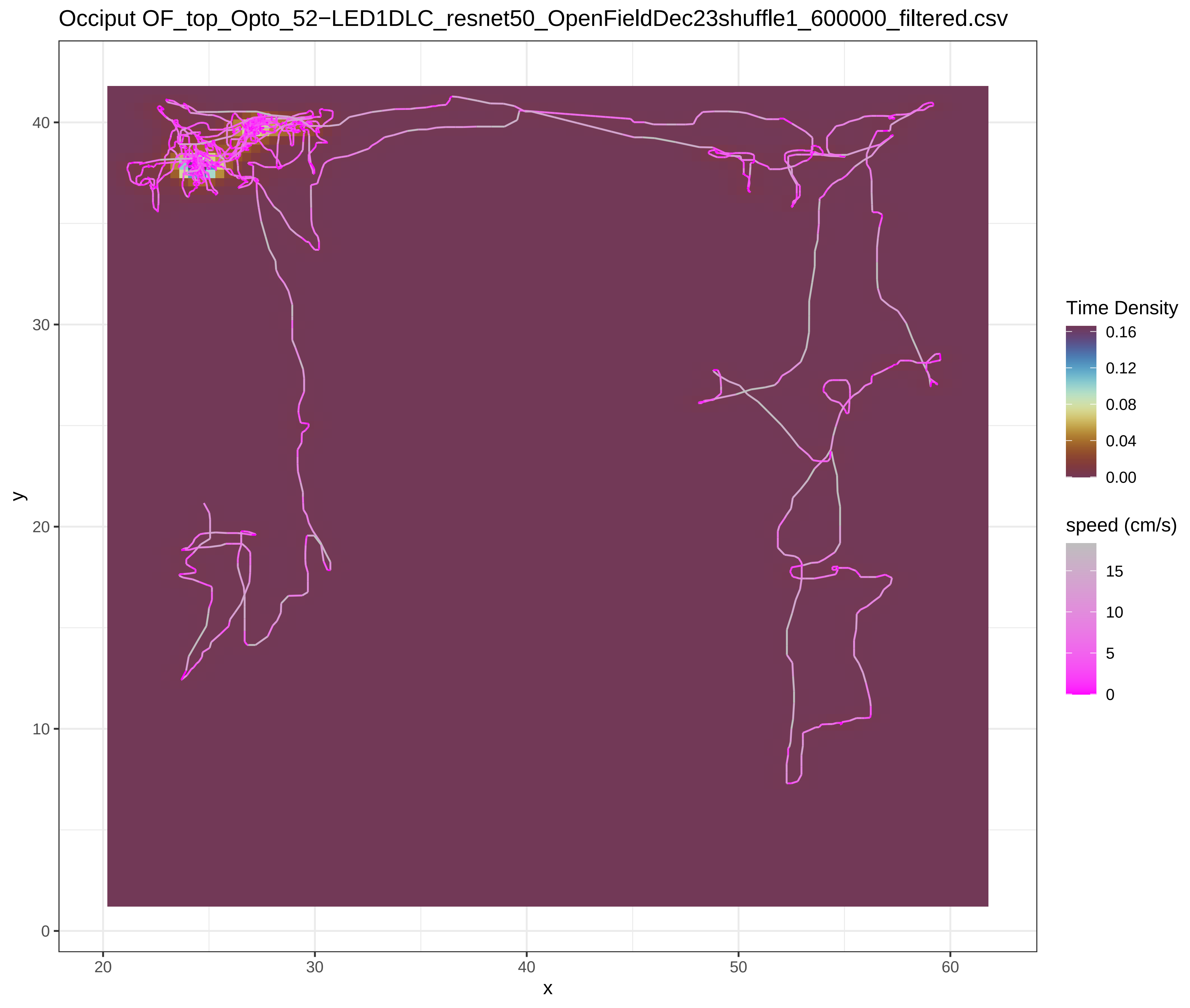
Occiput OF_top_Opto_52−LED1DLC_resnet50_OpenFieldDec23shuffle1_600000_filtered.csv
Time Density
0.16
0.12
0.08
0.04
0.00
speed (cm/s)
15
10
5
0
y
x
40
30
20
10
0
20
30
40
50
60

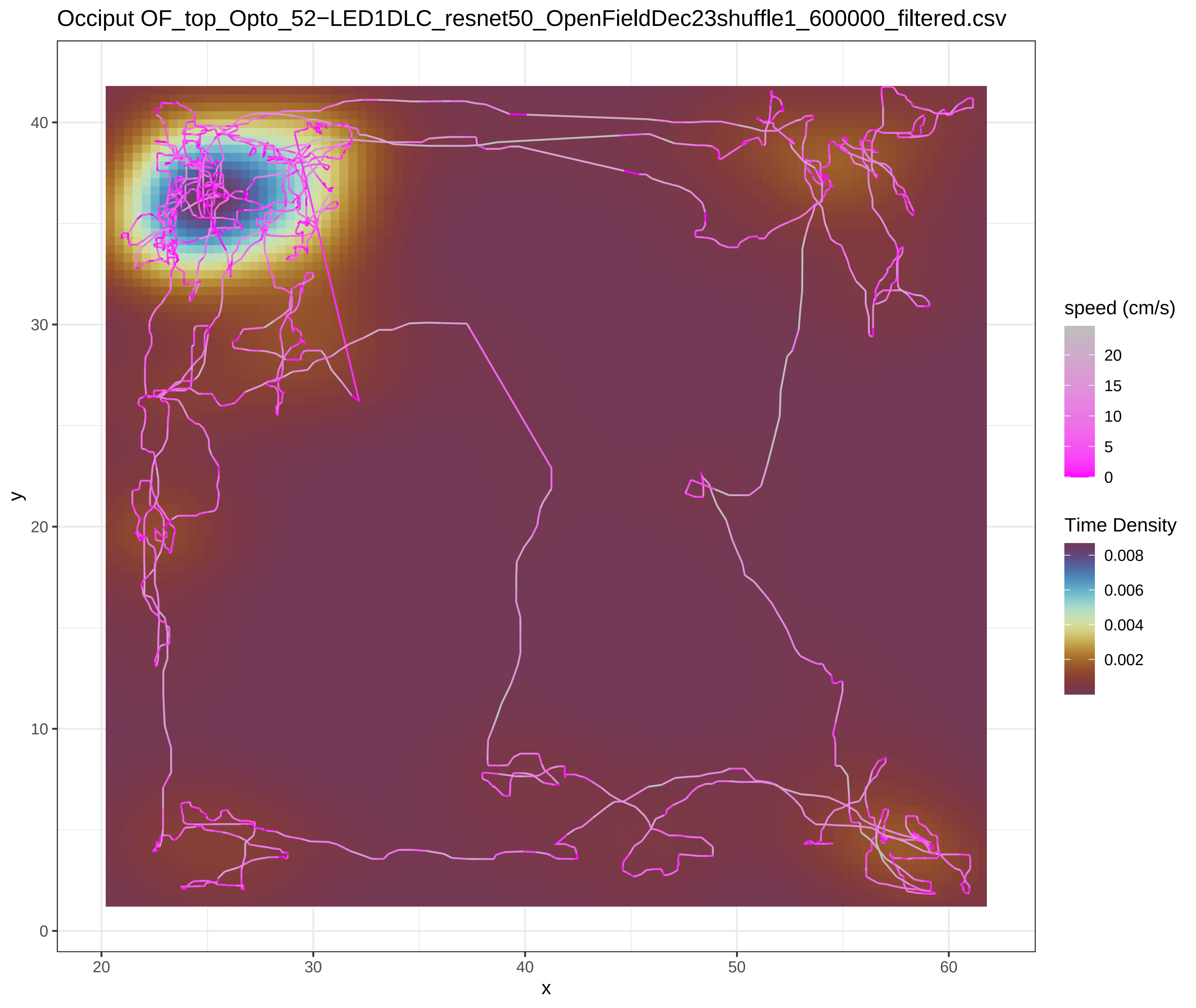
Occiput OF_top_Opto_52−LED1DLC_resnet50_OpenFieldDec23shuffle1_600000_filtered.csv
speed (cm/s)
20
15
10
5
0
Time Density
0.008
0.006
0.004
0.002
y
x
40
30
20
10
0
20
30
40
50
60

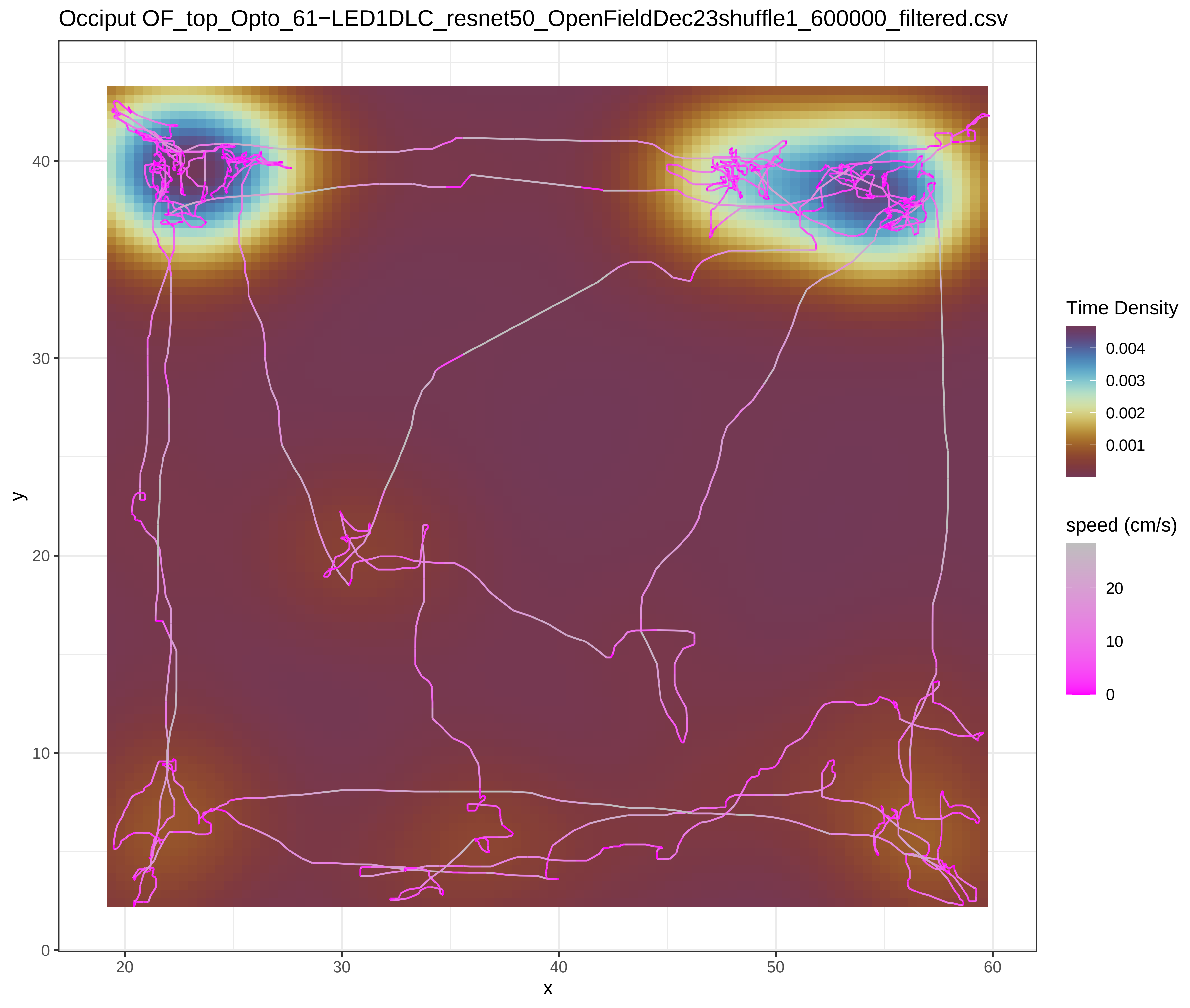
Occiput OF_top_Opto_61−LED1DLC_resnet50_OpenFieldDec23shuffle1_600000_filtered.csv
Time Density
0.004
0.003
0.002
0.001
speed (cm/s)
20
10
0
x
y
20
30
40
50
60
0
10
20
30
40

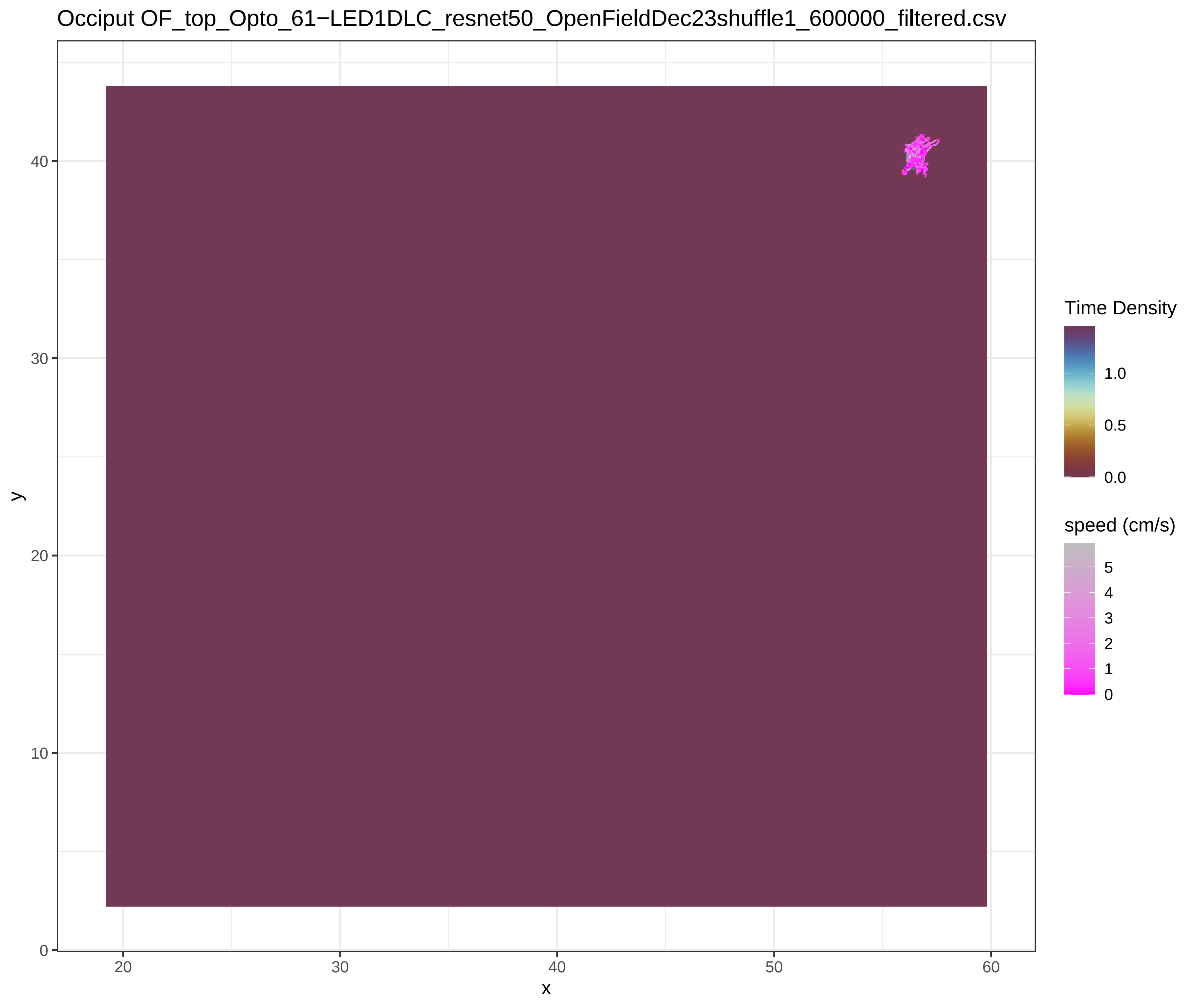
Occiput OF_top_Opto_61−LED1DLC_resnet50_OpenFieldDec23shuffle1_600000_filtered.csv
Time Density
1.0
0.5
0.0
speed (cm/s)
5
4
3
2
1
0
x
y
20
30
40
50
60
0
10
20
30
40

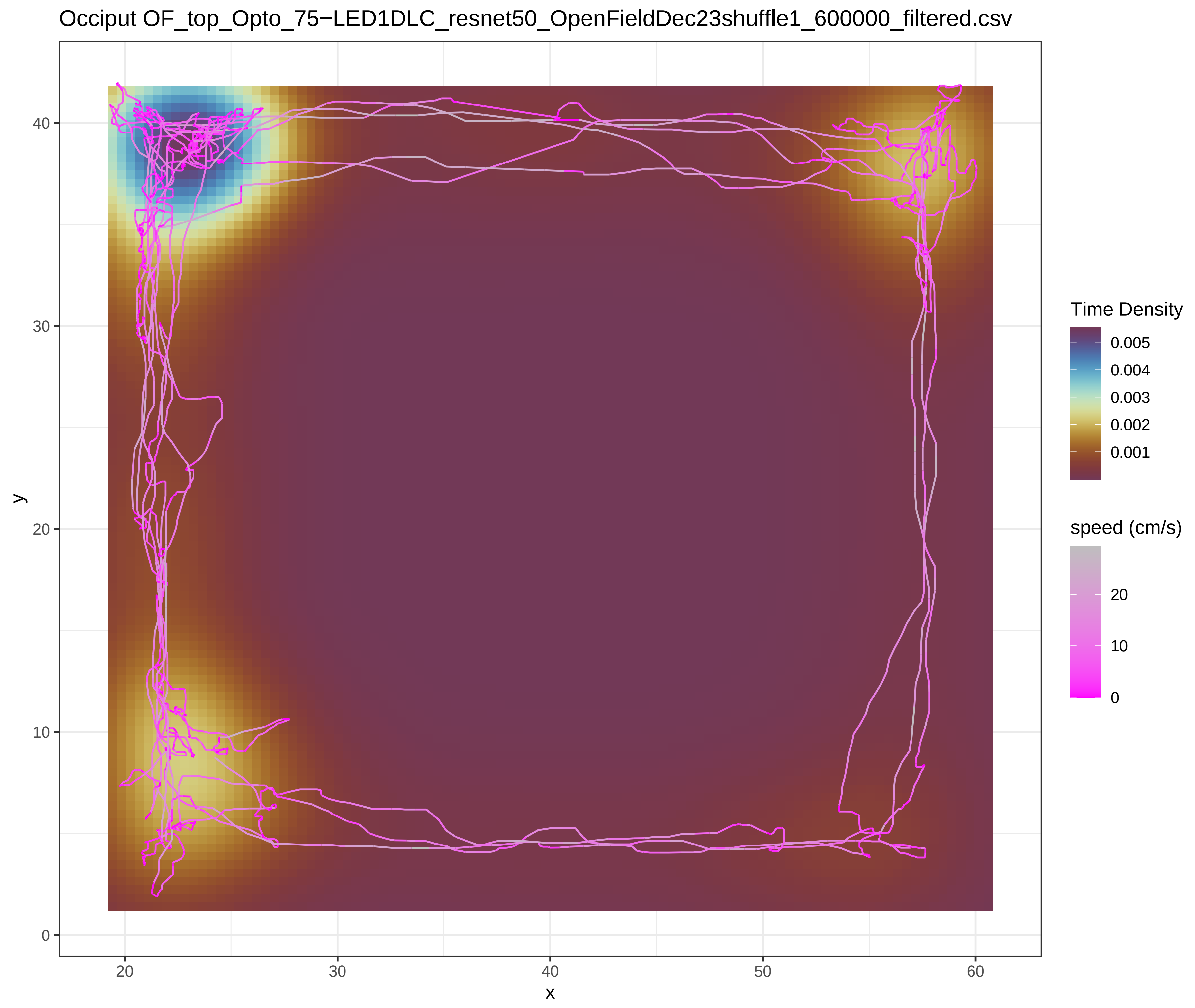

Occiput OF_top_Opto_75−LED1DLC_resnet50_OpenFieldDec23shuffle1_600000_filtered.csv
Time Density
0.005
0.004
0.003
0.002
0.001
speed (cm/s)
20
10
0
x
y
20
30
40
50
60
0
10
20
30
40


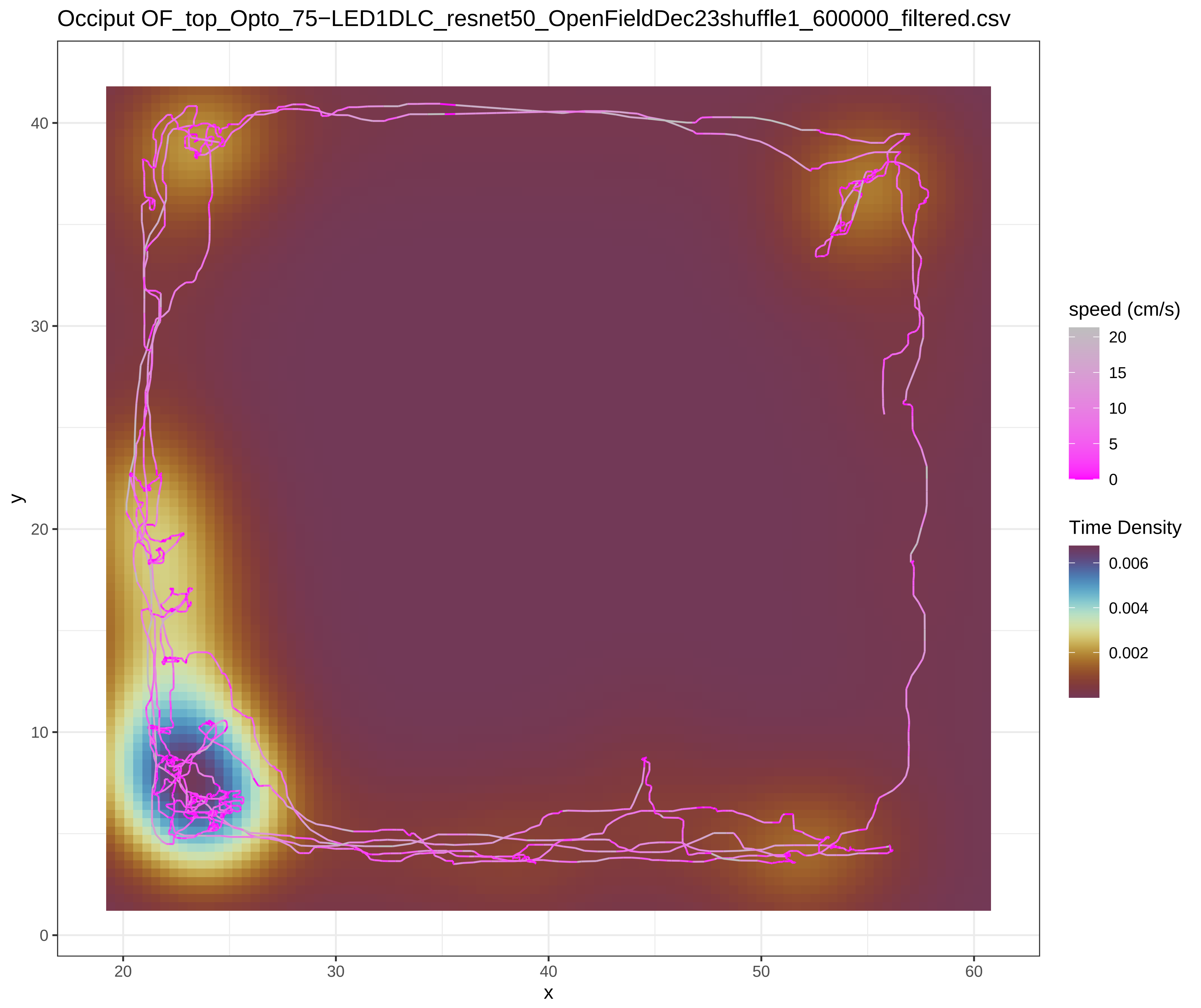

Occiput OF_top_Opto_75−LED1DLC_resnet50_OpenFieldDec23shuffle1_600000_filtered.csv
speed (cm/s)
20
15
10
5
0
Time Density
0.006
0.004
0.002
x
y
20
30
40
50
60
0
10
20
30
40

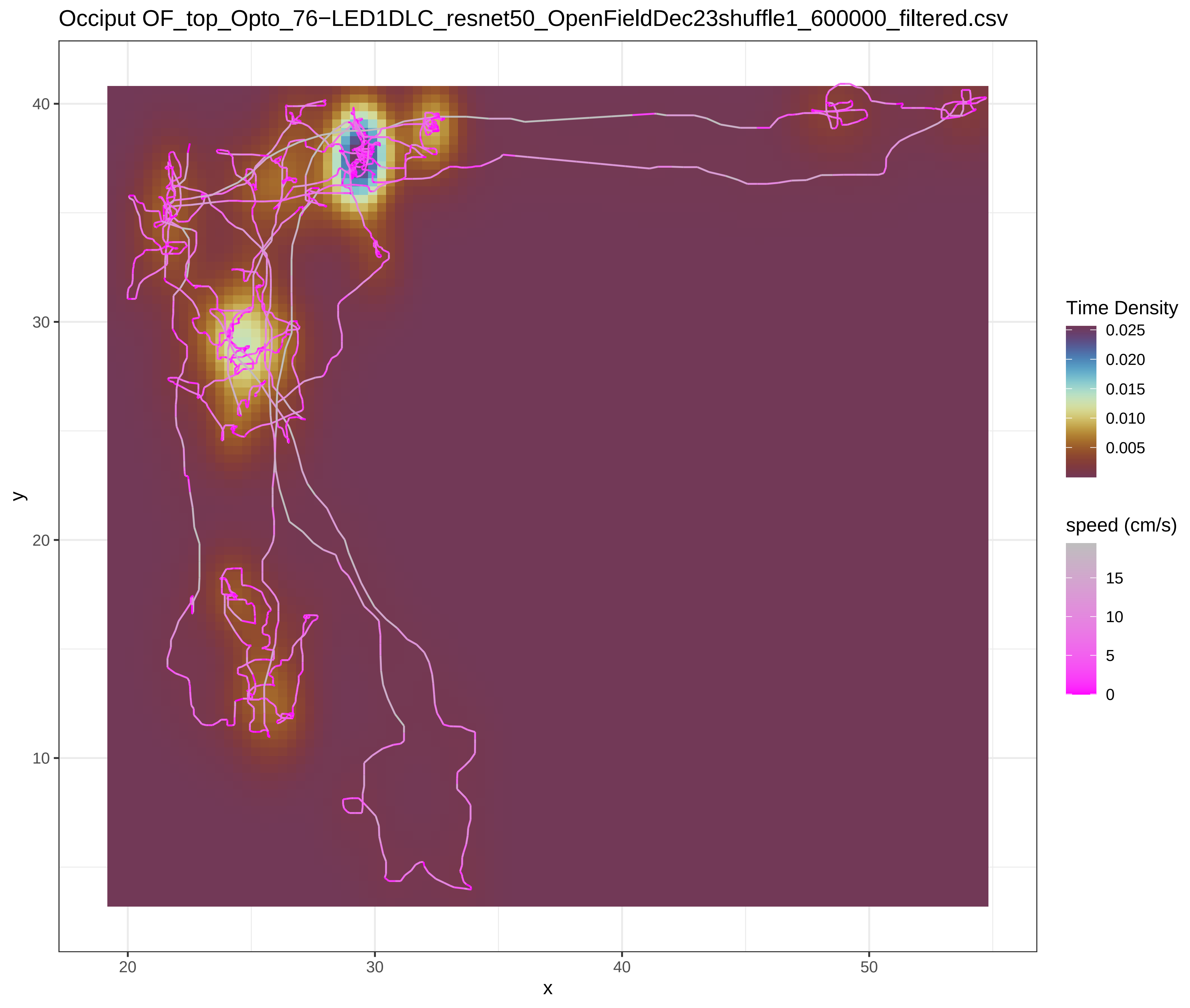

Occiput OF_top_Opto_76−LED1DLC_resnet50_OpenFieldDec23shuffle1_600000_filtered.csv
Time Density
0.025
0.020
0.015
0.010
0.005
speed (cm/s)
15
10
5
0
y
x
40
30
20
10
20
30
40
50


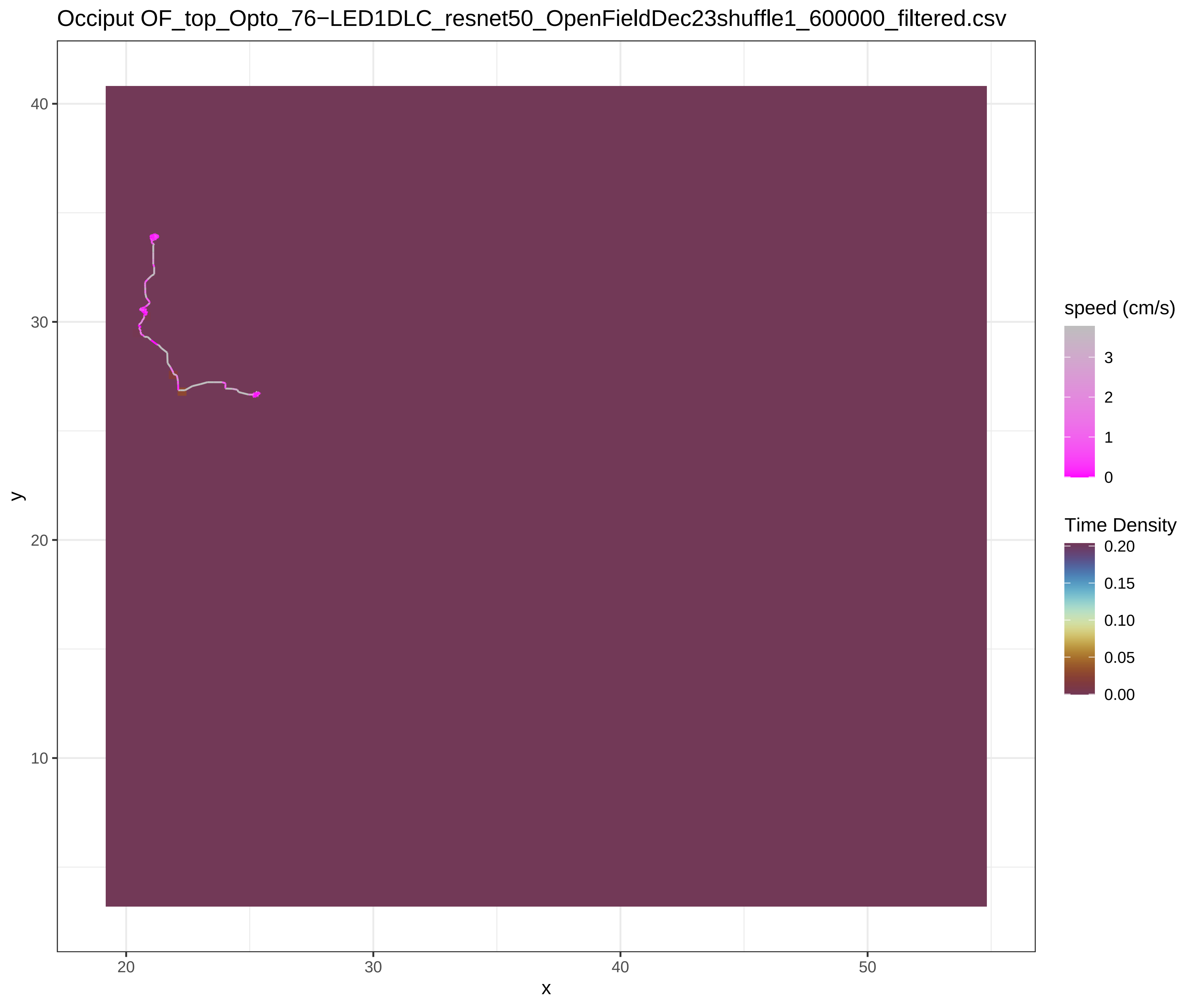

Occiput OF_top_Opto_76−LED1DLC_resnet50_OpenFieldDec23shuffle1_600000_filtered.csv
speed (cm/s)
3
2
1
0
Time Density
0.20
0.15
0.10
0.05
0.00
y
x
40
30
20
10
20
30
40
50

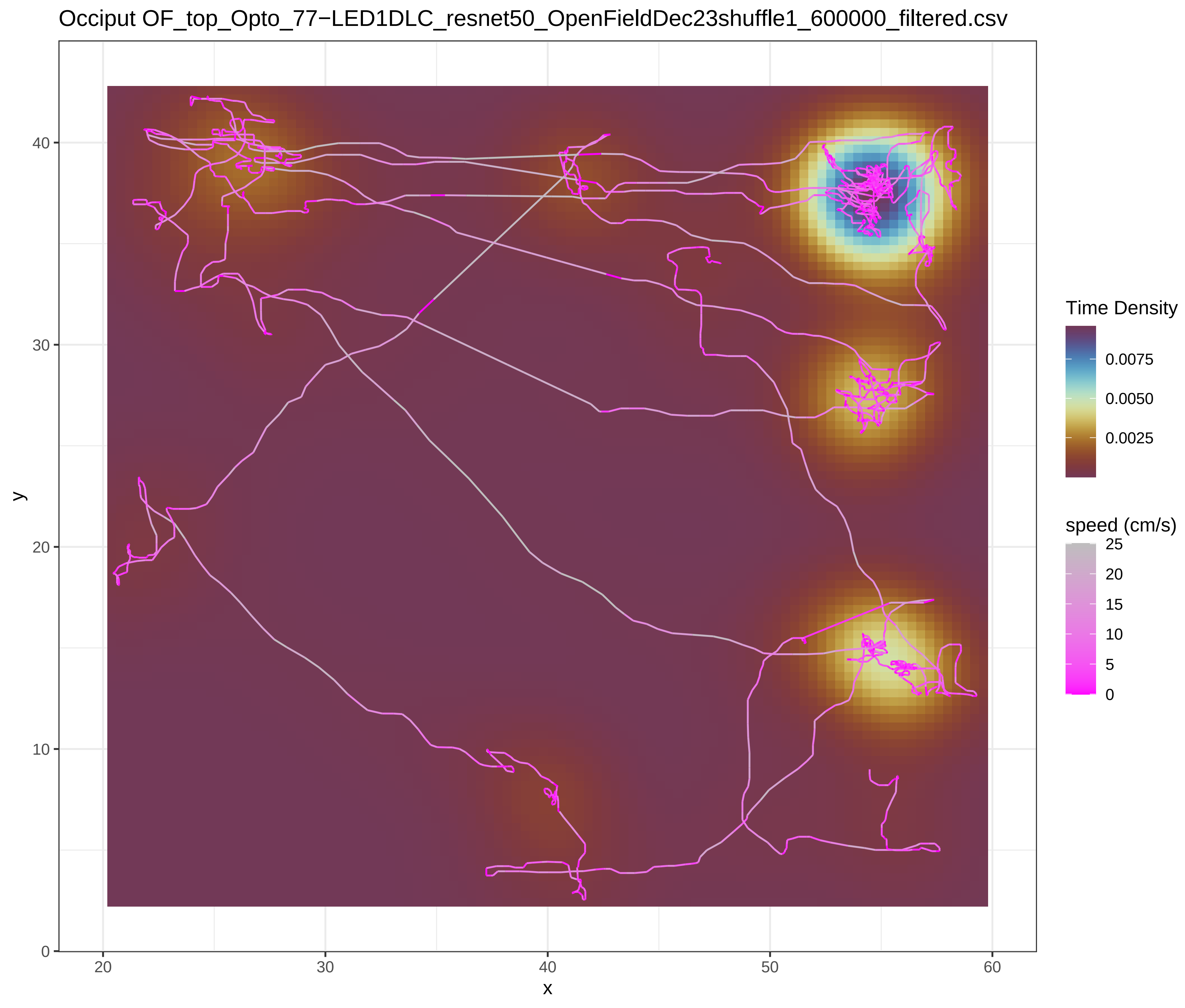
Occiput OF_top_Opto_77−LED1DLC_resnet50_OpenFieldDec23shuffle1_600000_filtered.csv
Time Density
0.0075
0.0050
0.0025
speed (cm/s)
25
20
15
10
5
0
y
x
0
10
20
30
40
20
30
40
50
60

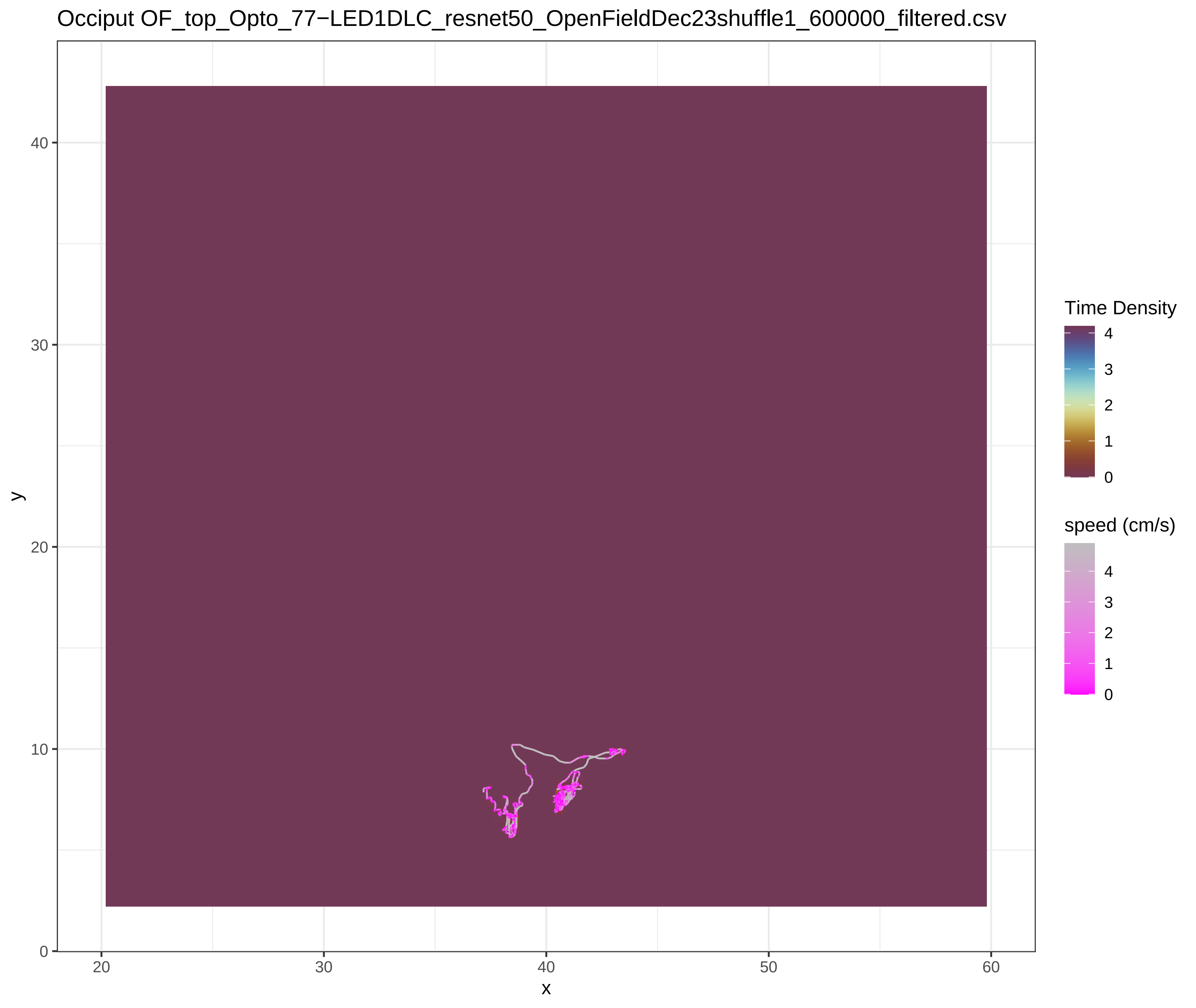
Occiput OF_top_Opto_77−LED1DLC_resnet50_OpenFieldDec23shuffle1_600000_filtered.csv
Time Density
4
3
2
1
0
speed (cm/s)
4
3
2
1
0
y
x
0
10
20
30
40
20
30
40
50
60
