## SupplementaryFile_S5 for "Chemo- and optogenetic activation of hypothalamic Foxb1-expressing neurons and their terminal endings in the rostral-dorsolateral PAG leads to tachypnea, bradycardia, and immobility"

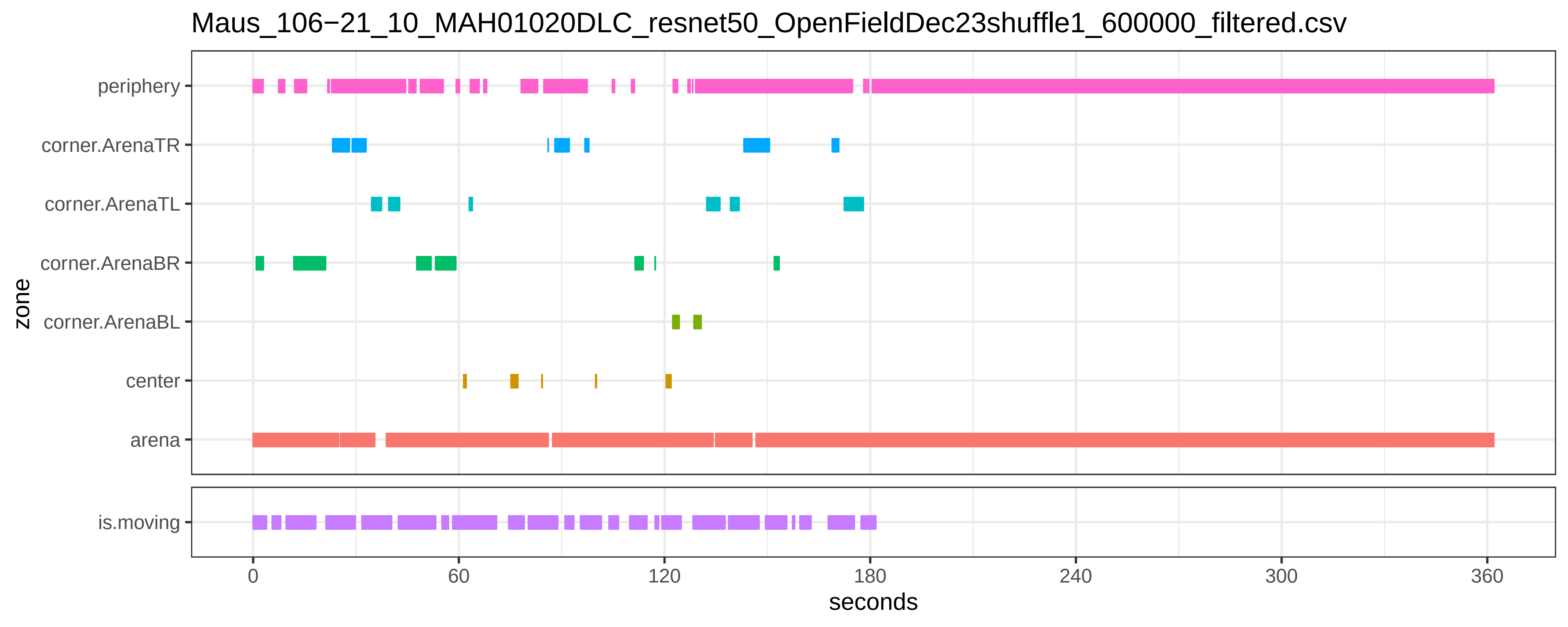
Maus_106−21_10_MAH01020DLC_resnet50_OpenFieldDec23shuffle1_600000_filtered.csv
periphery
corner.ArenaTR
corner.ArenaTL
corner.ArenaBR
corner.ArenaBL
center
arena
zone
is.moving
0
60
120
180
240
300
360
seconds

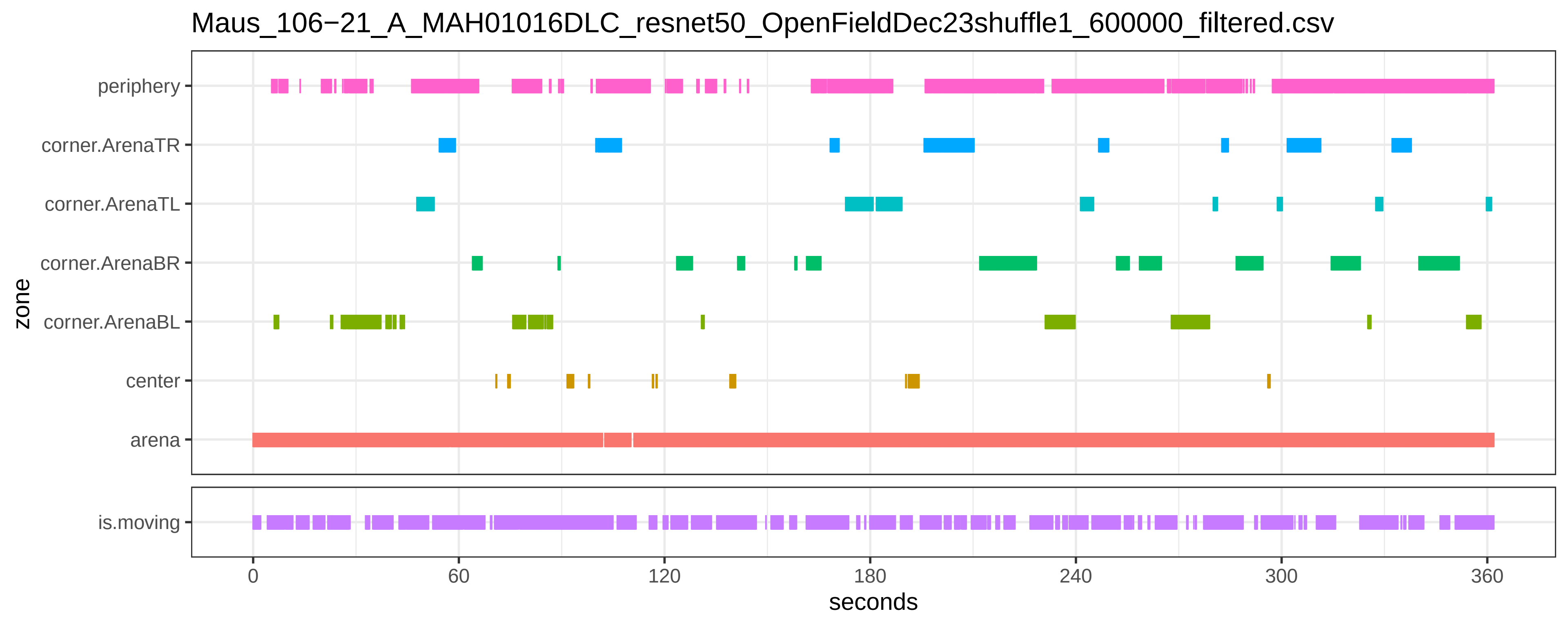

Maus_106−21_A_MAH01016DLC_resnet50_OpenFieldDec23shuffle1_600000_filtered.csv
periphery
corner.ArenaTR
corner.ArenaTL
corner.ArenaBR
corner.ArenaBL
center
arena
is.moving
zone
0
60
120
180
240
300
360
seconds

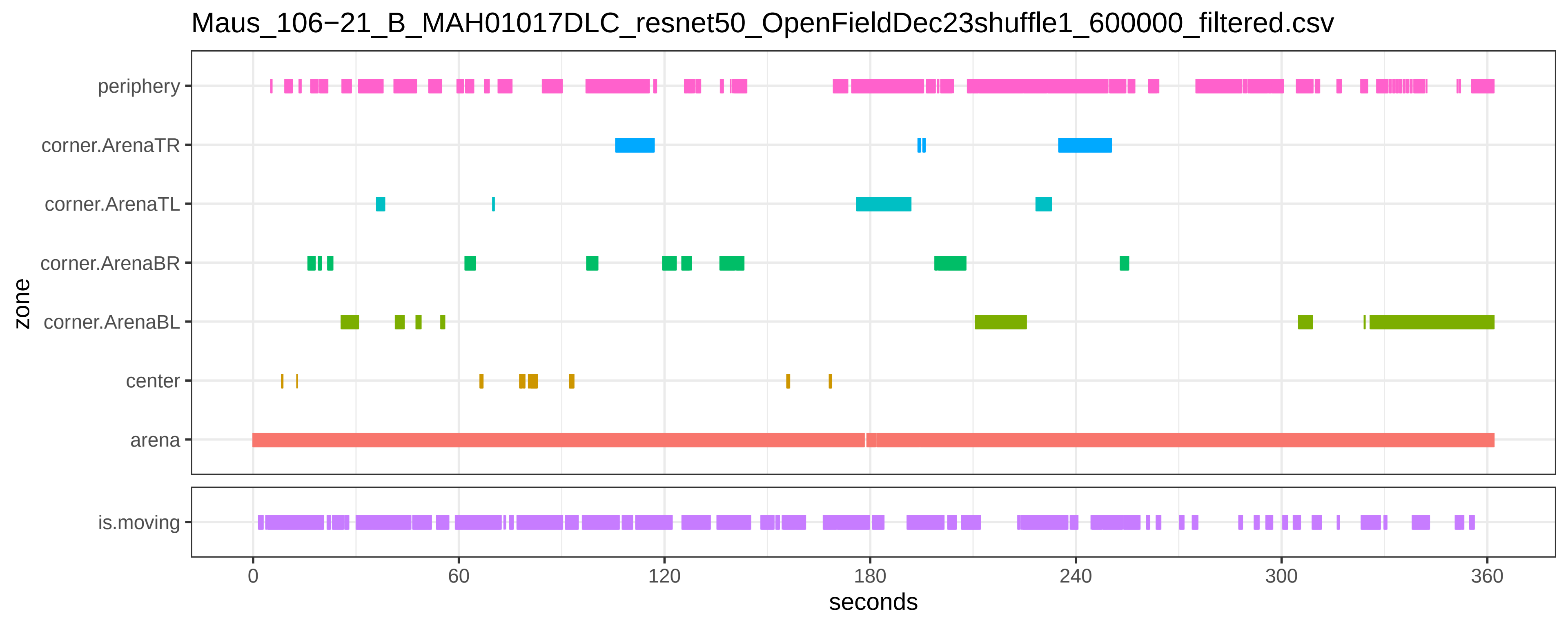

Maus_106−21_B_MAH01017DLC_resnet50_OpenFieldDec23shuffle1_600000_filtered.csv
periphery
corner.ArenaTR
corner.ArenaTL
corner.ArenaBR
corner.ArenaBL
center
arena
zone
is.moving
0
60
120
180
240
300
360
seconds

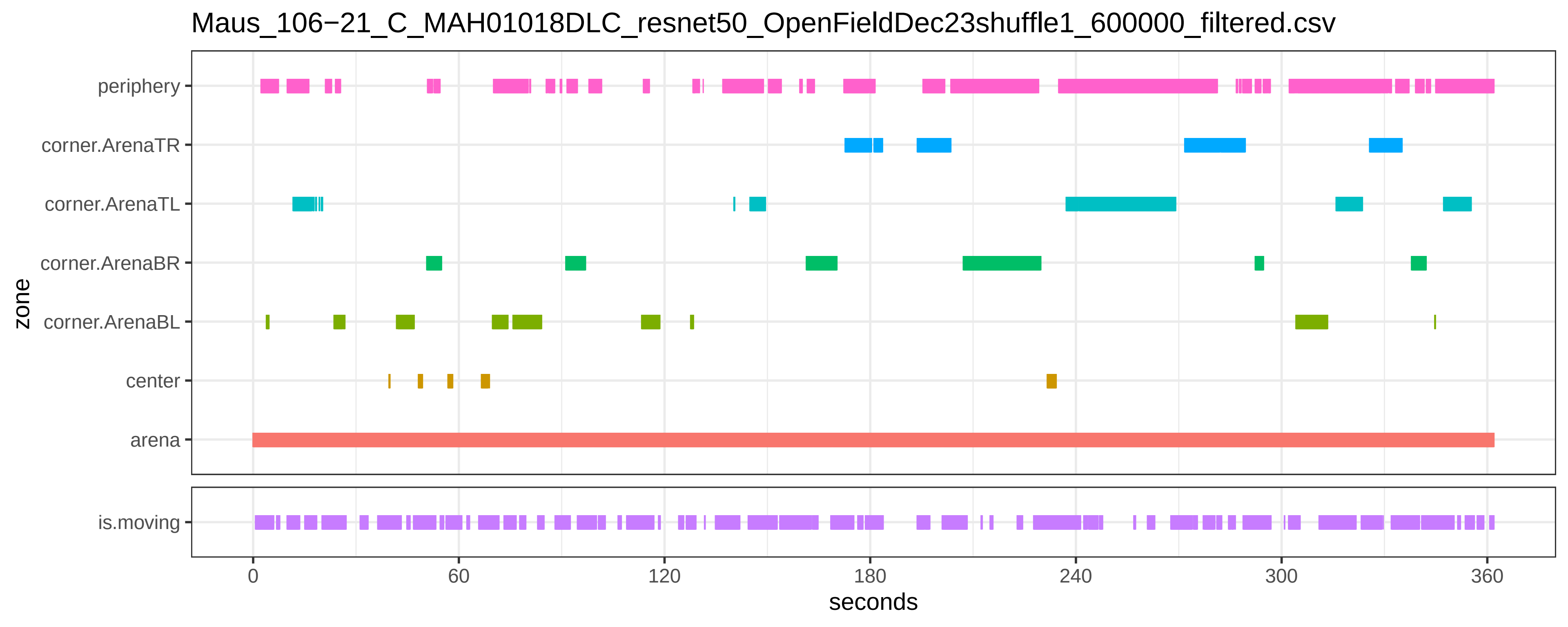
Maus_106−21_C_MAH01018DLC_resnet50_OpenFieldDec23shuffle1_600000_filtered.csv
periphery
corner.ArenaTR
corner.ArenaTL
corner.ArenaBR
corner.ArenaBL
center
arena
zone
is.moving
0
60
120
180
240
300
360
seconds

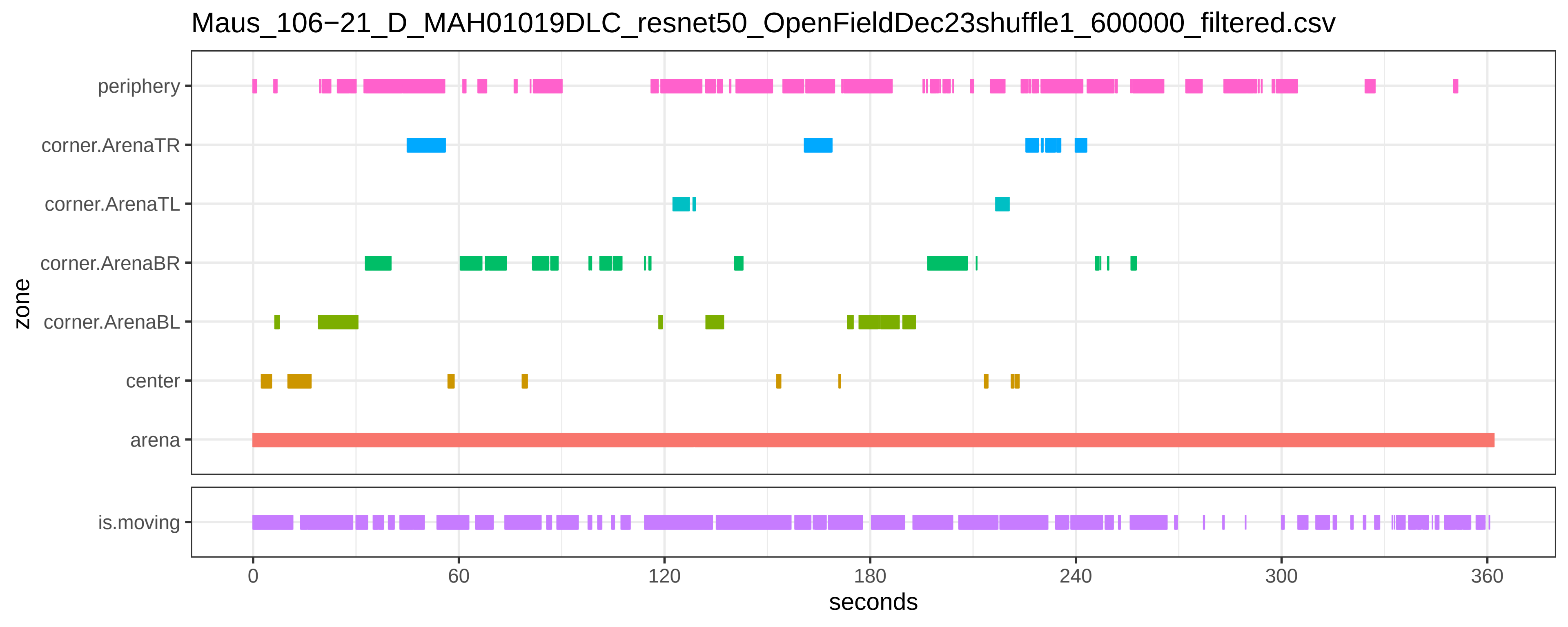
Maus_106−21_D_MAH01019DLC_resnet50_OpenFieldDec23shuffle1_600000_filtered.csv
periphery
corner.ArenaTR
corner.ArenaTL
corner.ArenaBR
corner.ArenaBL
center
arena
is.moving
zone
0
60
120
180
240
300
360
seconds

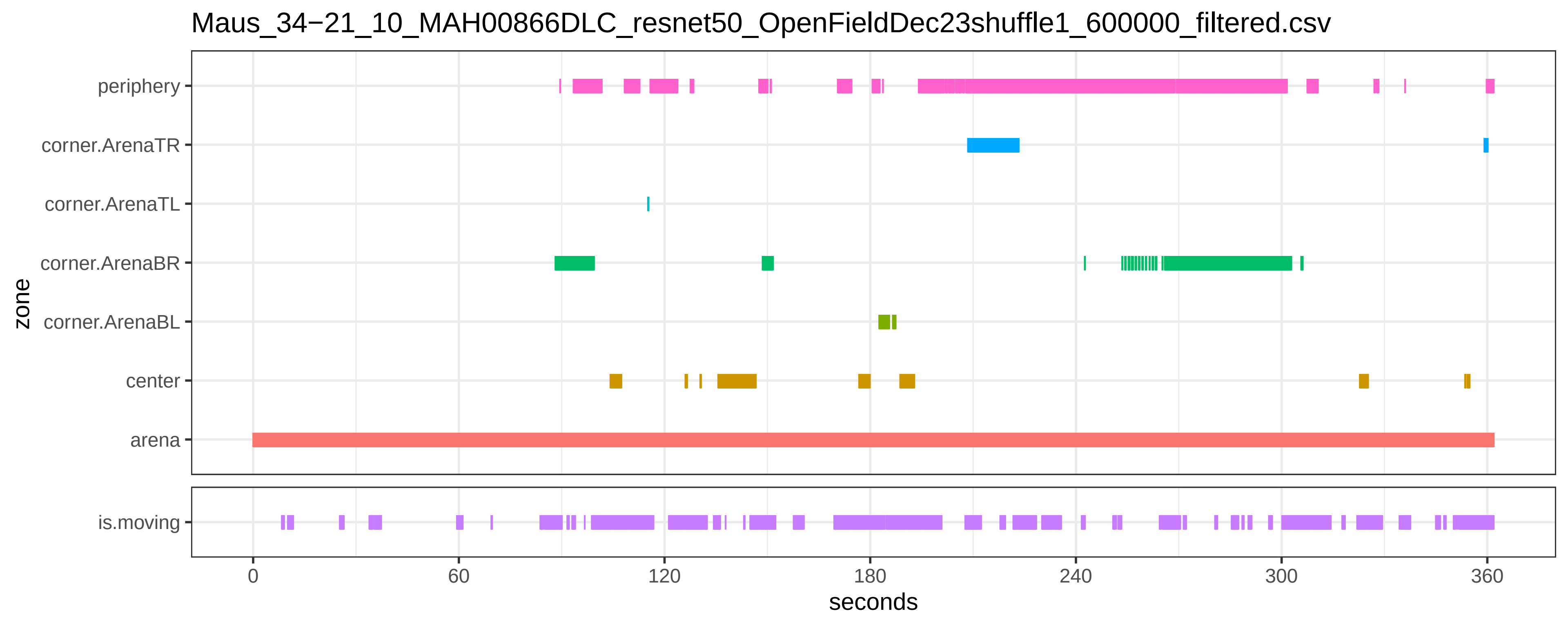
Maus_34−21_10_MAH00866DLC_resnet50_OpenFieldDec23shuffle1_600000_filtered.csv
periphery
corner.ArenaTR
corner.ArenaTL
corner.ArenaBR
corner.ArenaBL
center
arena
zone
is.moving
0
60
120
180
240
300
360
seconds

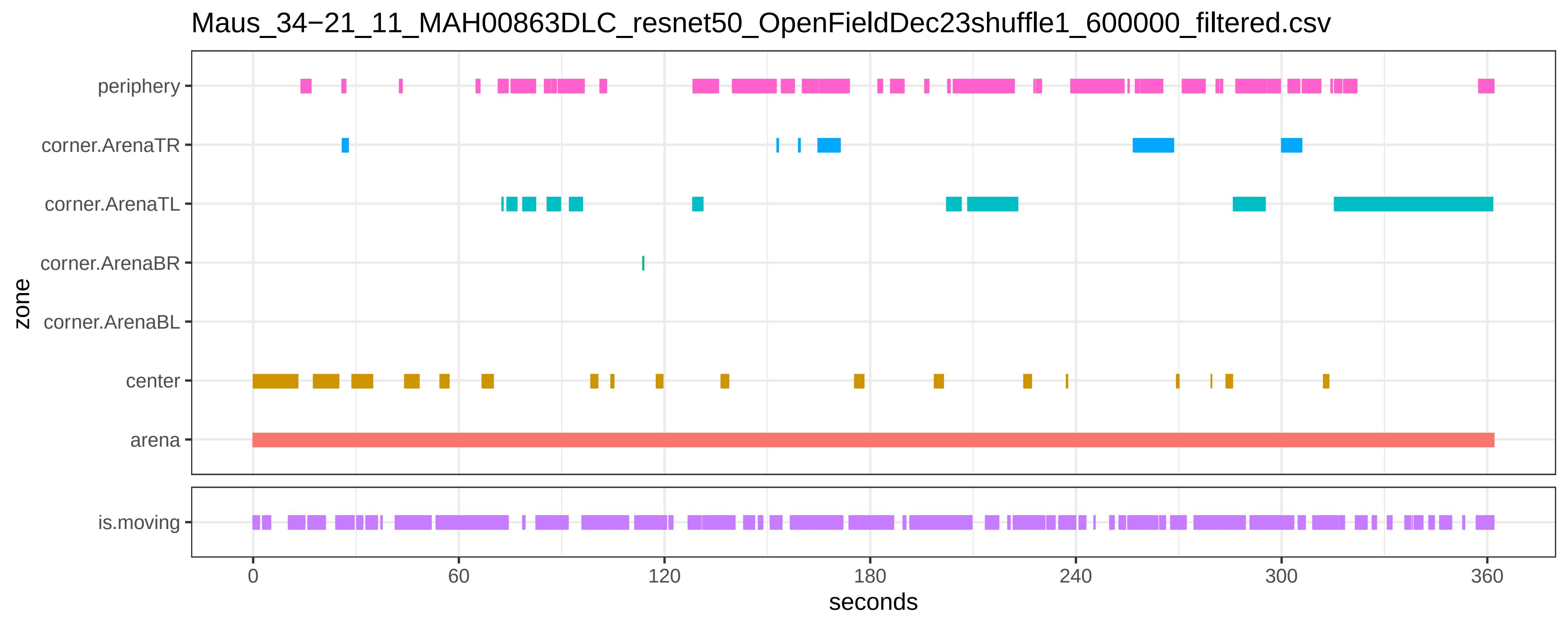
Maus_34−21_11_MAH00863DLC_resnet50_OpenFieldDec23shuffle1_600000_filtered.csv
periphery
corner.ArenaTR
corner.ArenaTL
corner.ArenaBR
corner.ArenaBL
center
arena
zone
is.moving
0
60
120
180
240
300
360
seconds

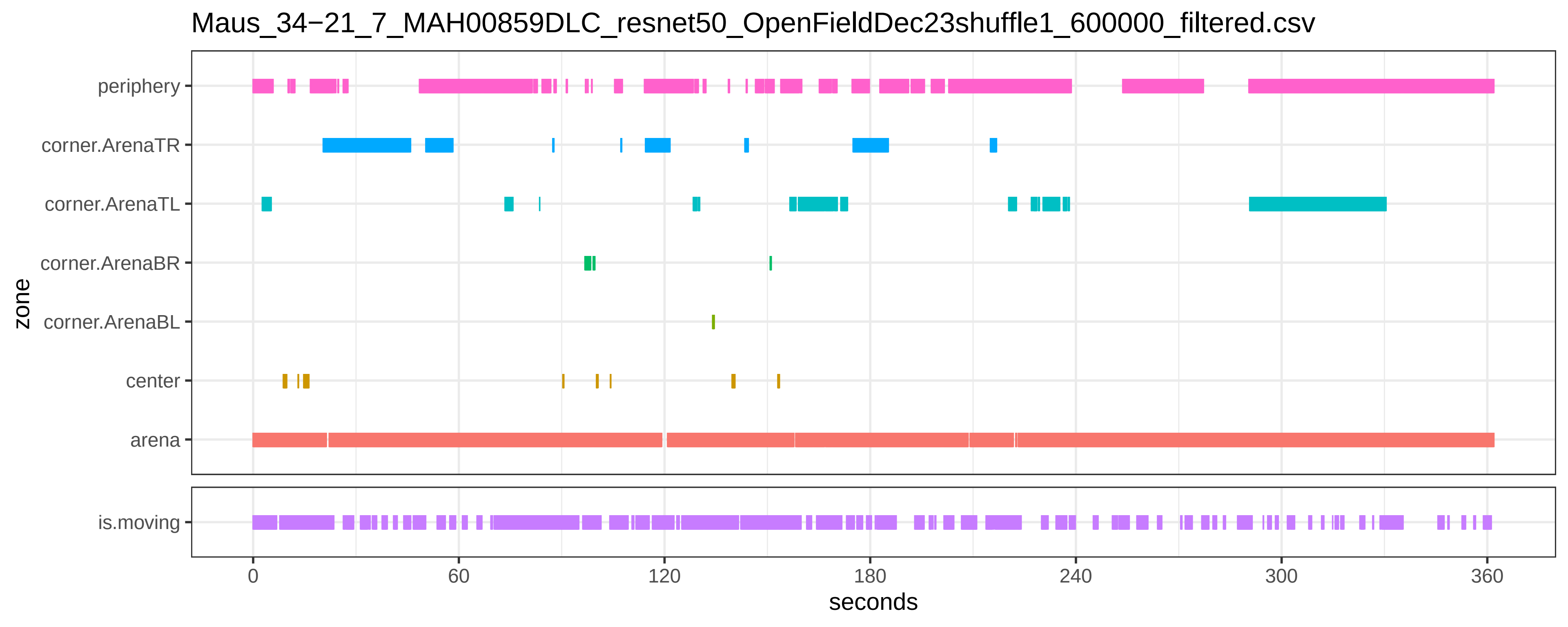

Maus_34−21_7_MAH00859DLC_resnet50_OpenFieldDec23shuffle1_600000_filtered.csv
periphery
corner.ArenaTR
corner.ArenaTL
corner.ArenaBR
corner.ArenaBL
center
arena
is.moving
zone
0
60
120
180
240
300
360
seconds

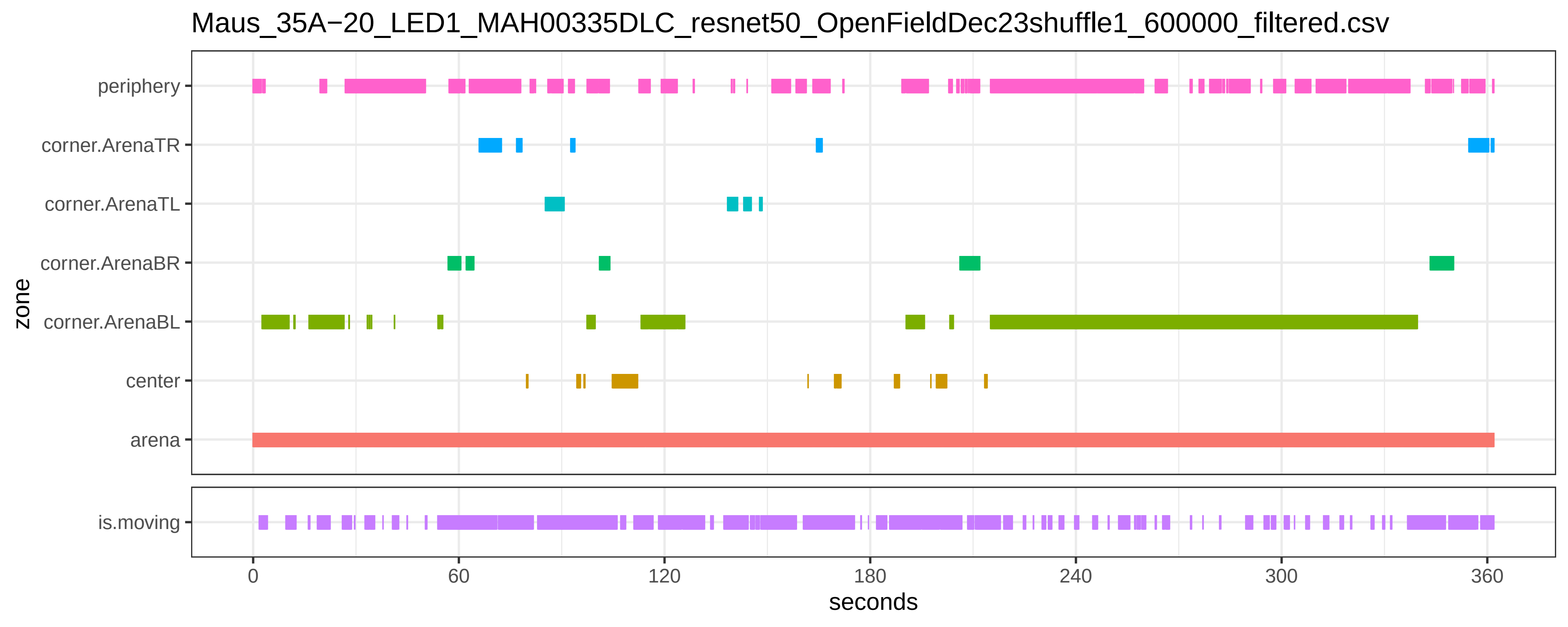

Maus_35A−20_LED1_MAH00335DLC_resnet50_OpenFieldDec23shuffle1_600000_filtered.csv
periphery
corner.ArenaTR
corner.ArenaTL
corner.ArenaBR
corner.ArenaBL
center
arena
zone
is.moving
0
60
120
180
240
300
360
seconds

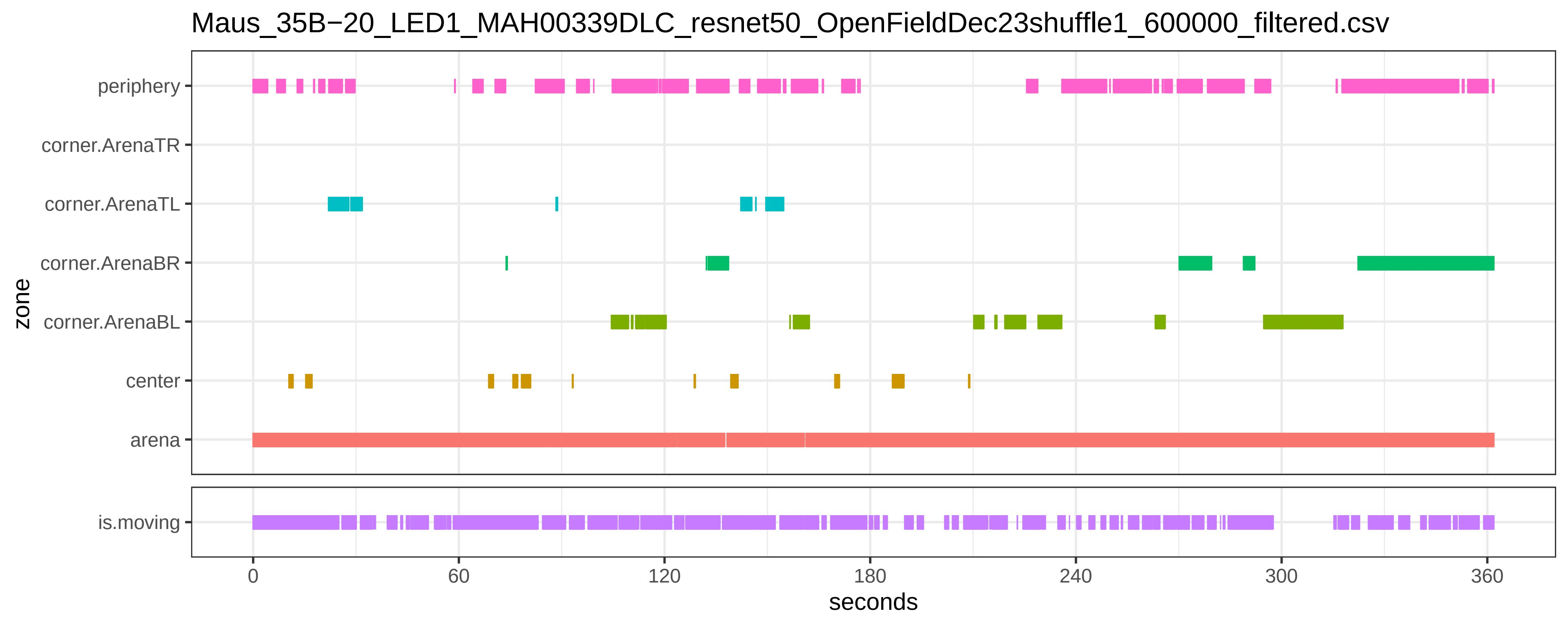

Maus_35B−20_LED1_MAH00339DLC_resnet50_OpenFieldDec23shuffle1_600000_filtered.csv
periphery
corner.ArenaTR
corner.ArenaTL
corner.ArenaBR
corner.ArenaBL
center
arena
zone
is.moving
0
60
120
180
240
300
360
seconds

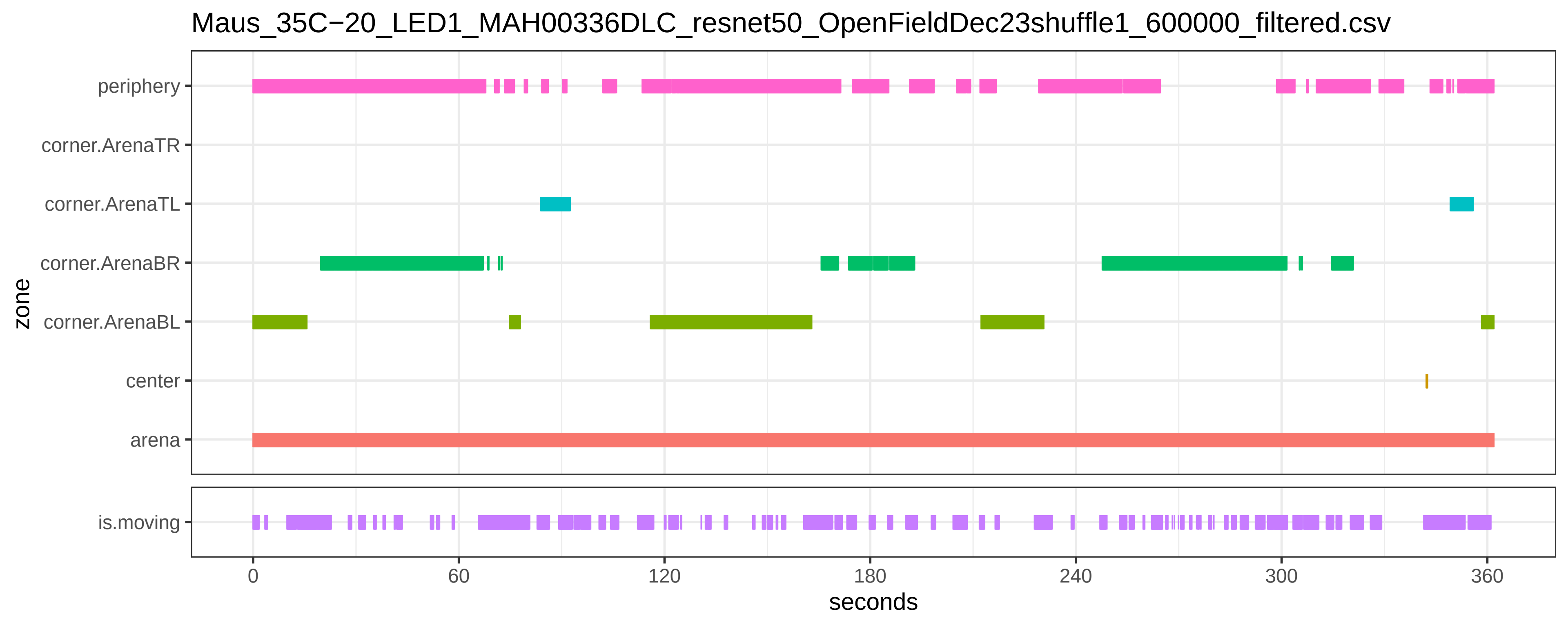
Maus_35C−20_LED1_MAH00336DLC_resnet50_OpenFieldDec23shuffle1_600000_filtered.csv
periphery
corner.ArenaTR
corner.ArenaTL
corner.ArenaBR
corner.ArenaBL
center
arena
zone
is.moving
0
60
120
180
240
300
360
seconds

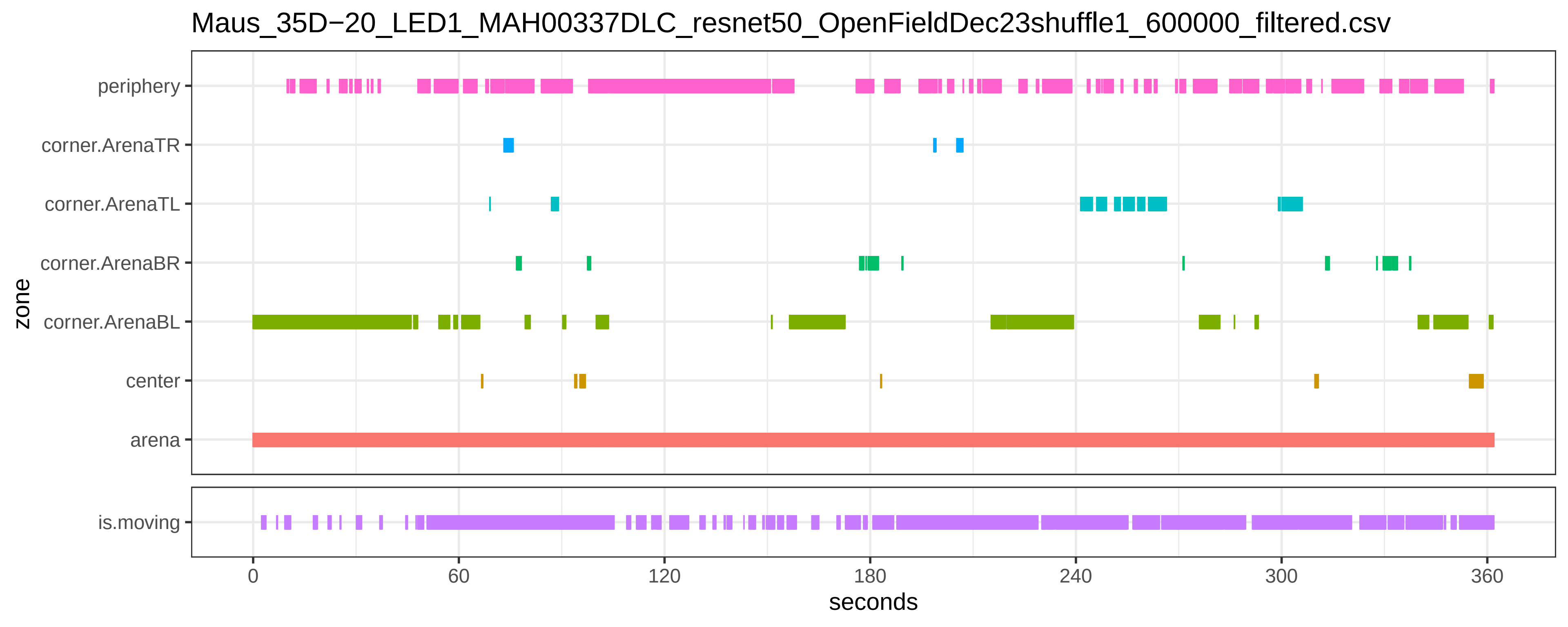
Maus_35D−20_LED1_MAH00337DLC_resnet50_OpenFieldDec23shuffle1_600000_filtered.csv
periphery
corner.ArenaTR
corner.ArenaTL
corner.ArenaBR
corner.ArenaBL
center
arena
zone
is.moving
0
60
120
180
240
300
360
seconds

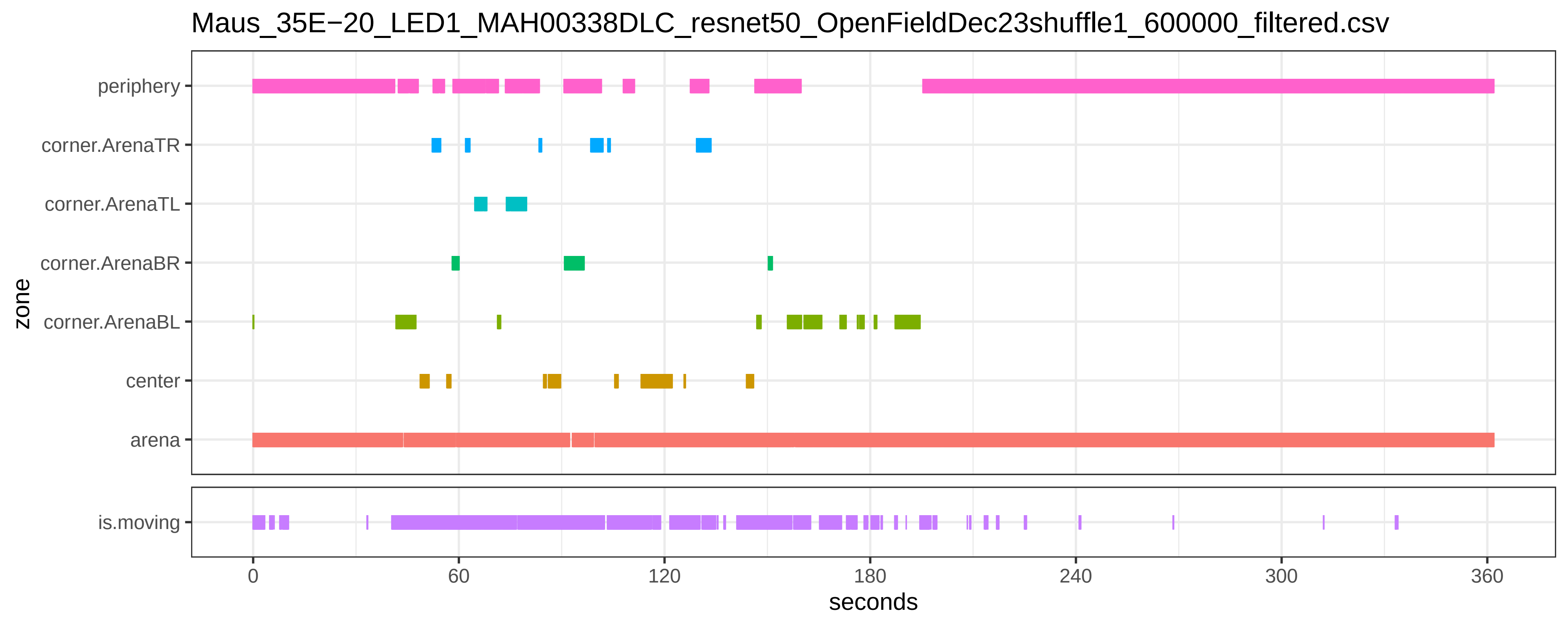
Maus_35E−20_LED1_MAH00338DLC_resnet50_OpenFieldDec23shuffle1_600000_filtered.csv
periphery
corner.ArenaTR
corner.ArenaTL
corner.ArenaBR
corner.ArenaBL
center
arena
zone
is.moving
0
60
120
180
240
300
360
seconds

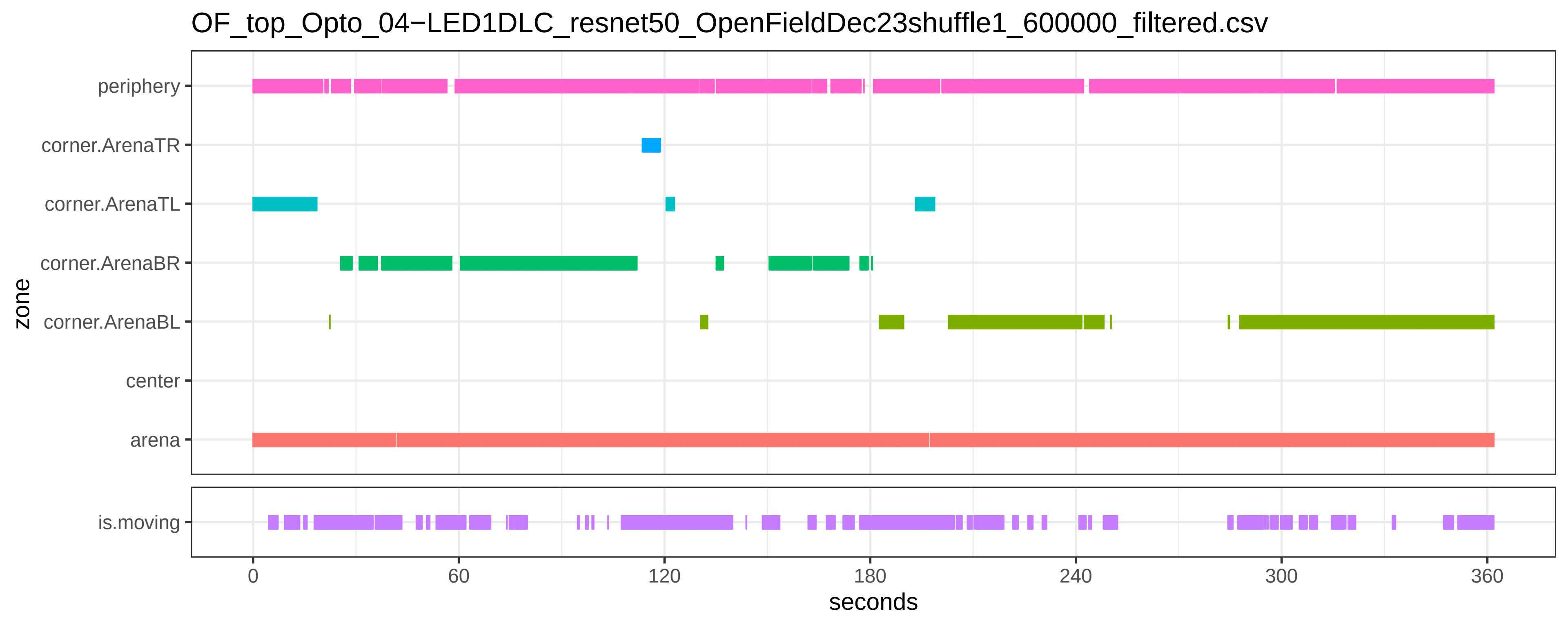
OF_top_Opto_04−LED1DLC_resnet50_OpenFieldDec23shuffle1_600000_filtered.csv
periphery
corner.ArenaTR
corner.ArenaTL
corner.ArenaBR
corner.ArenaBL
center
arena
zone
is.moving
0
60
120
180
240
300
360
seconds

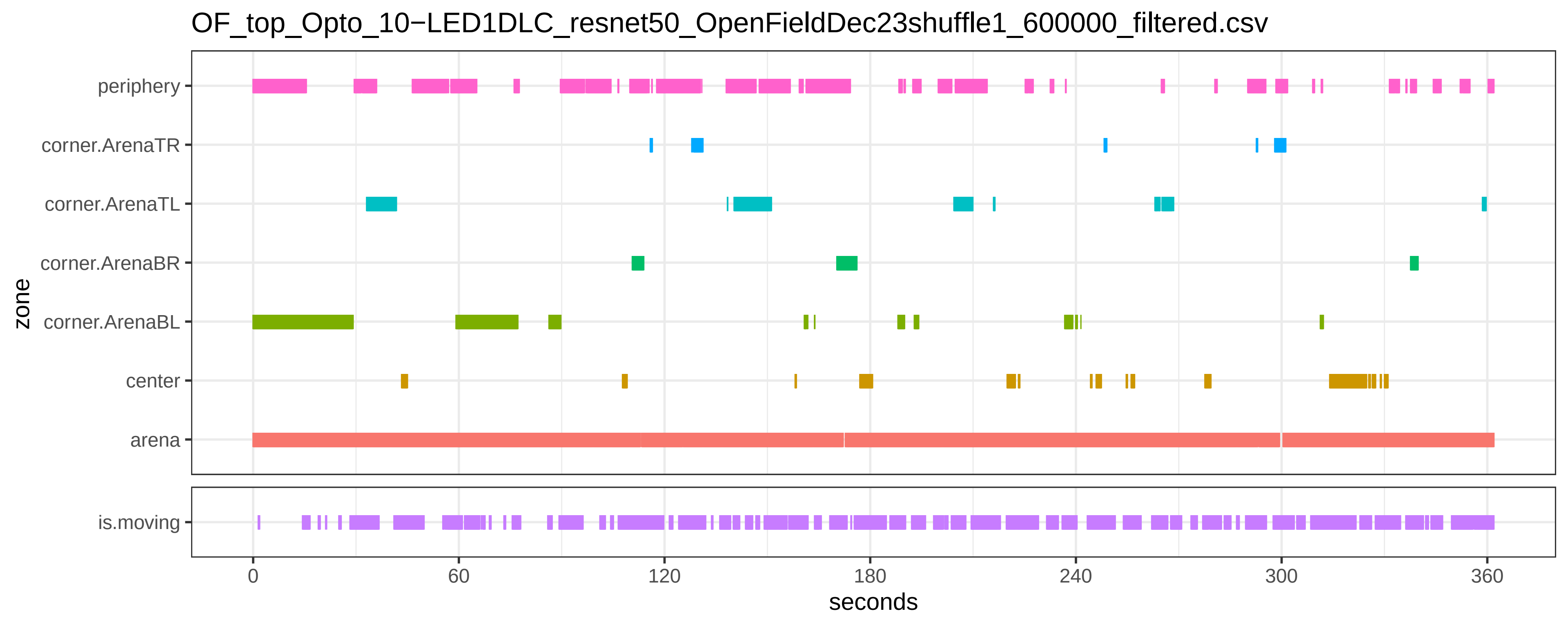

OF_top_Opto_10−LED1DLC_resnet50_OpenFieldDec23shuffle1_600000_filtered.csv
periphery
corner.ArenaTR
corner.ArenaTL
corner.ArenaBR
corner.ArenaBL
center
arena
zone
is.moving
0
60
120
180
240
300
360
seconds

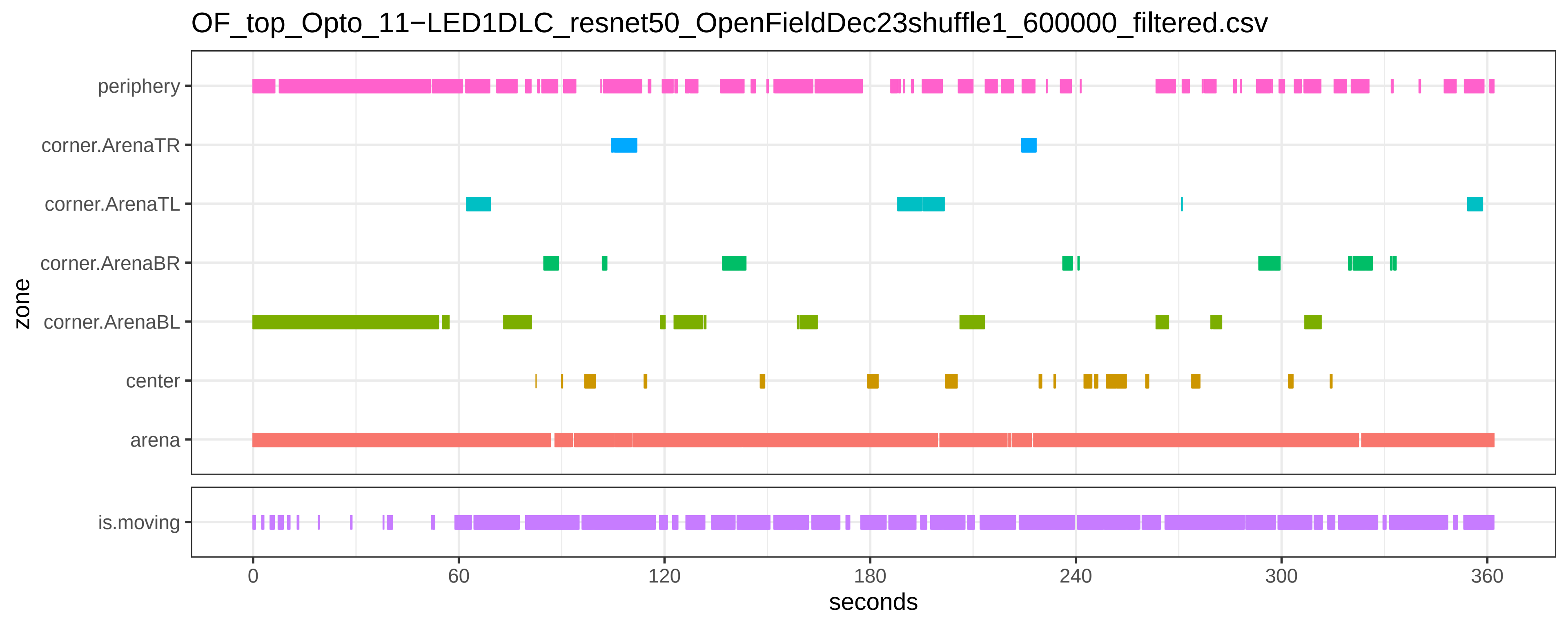

OF_top_Opto_11−LED1DLC_resnet50_OpenFieldDec23shuffle1_600000_filtered.csv
periphery
corner.ArenaTR
corner.ArenaTL
corner.ArenaBR
corner.ArenaBL
center
arena
zone
is.moving
0
60
120
180
240
300
360
seconds

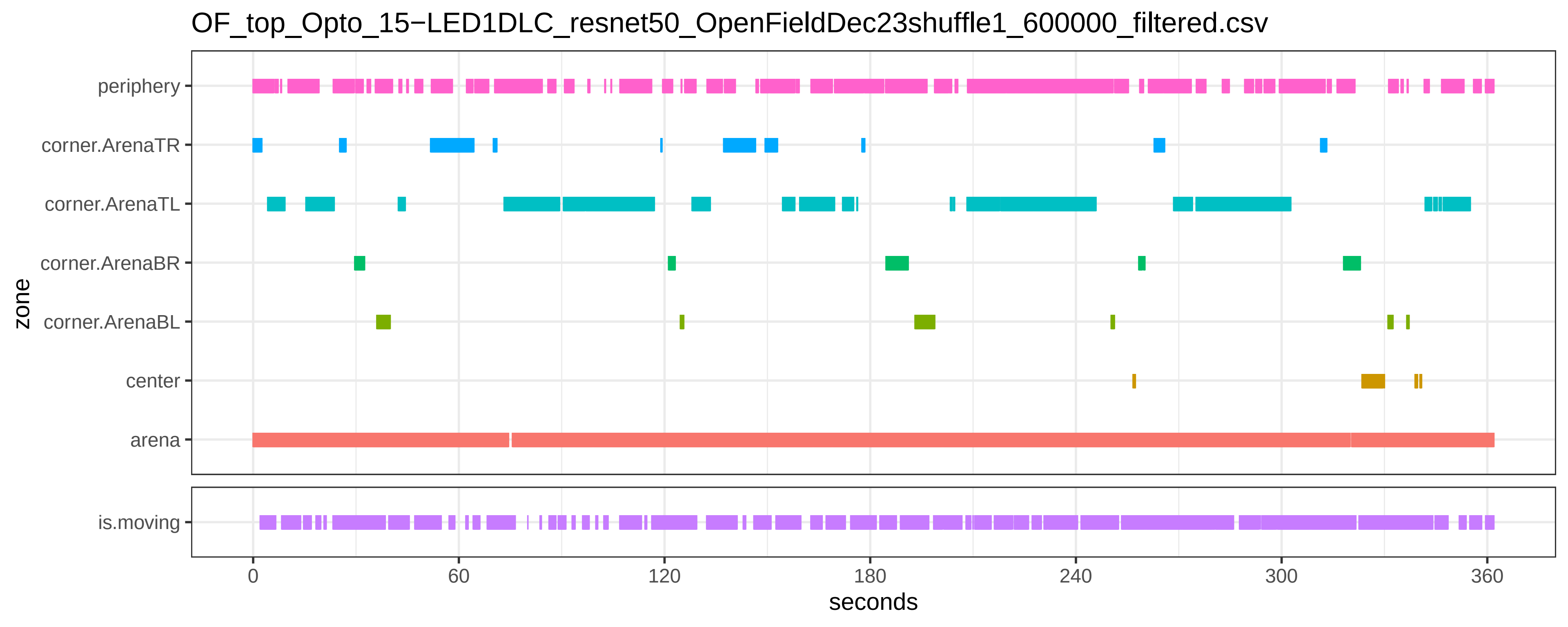

OF_top_Opto_15−LED1DLC_resnet50_OpenFieldDec23shuffle1_600000_filtered.csv
periphery
corner.ArenaTR
corner.ArenaTL
corner.ArenaBR
corner.ArenaBL
center
arena
zone
is.moving
0
60
120
180
240
300
360
seconds

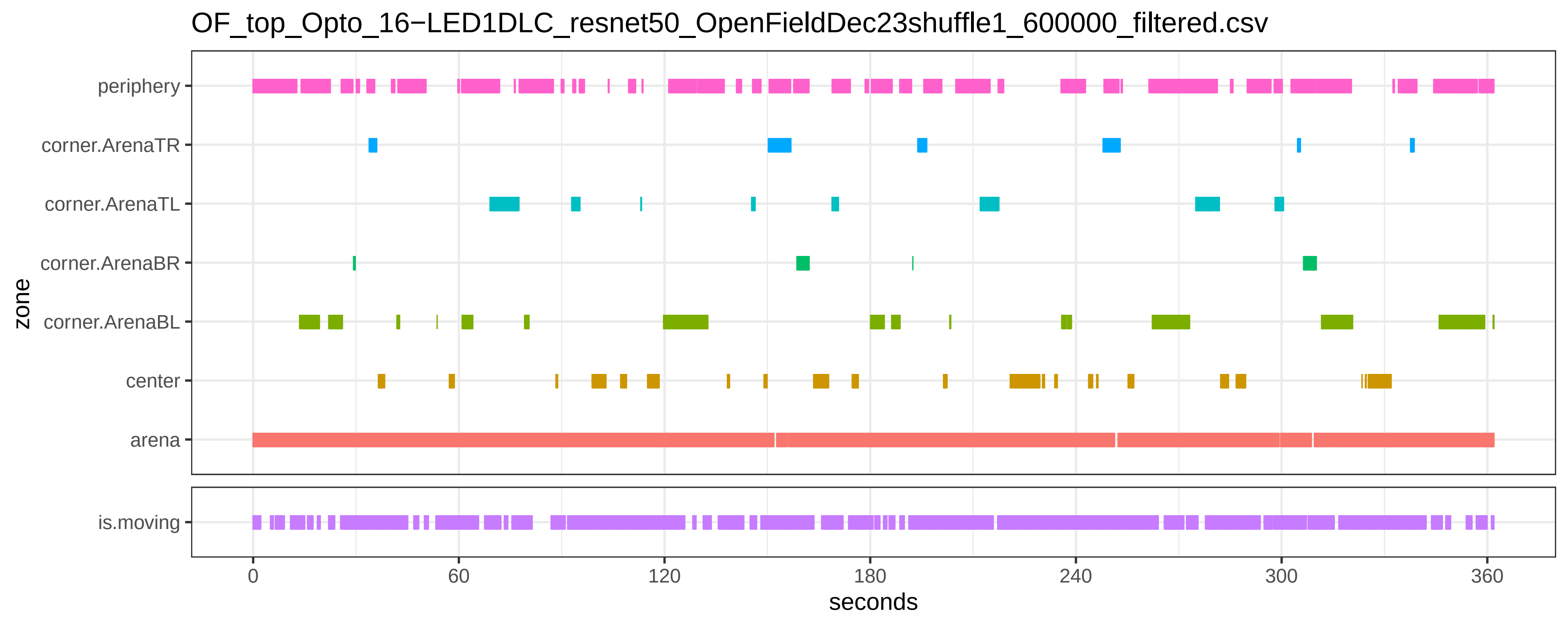

OF_top_Opto_16−LED1DLC_resnet50_OpenFieldDec23shuffle1_600000_filtered.csv
periphery
corner.ArenaTR
corner.ArenaTL
corner.ArenaBR
corner.ArenaBL
center
arena
zone
is.moving
0
60
120
180
240
300
360
seconds

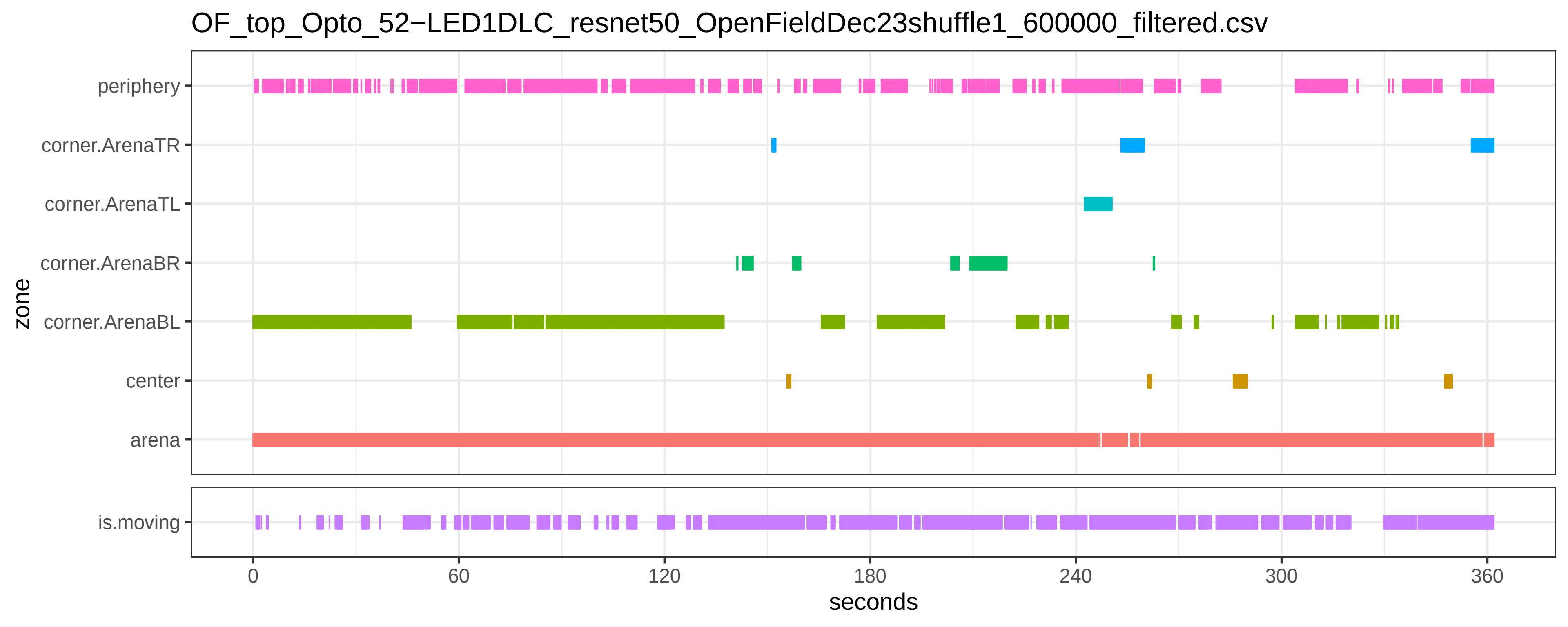

OF_top_Opto_52−LED1DLC_resnet50_OpenFieldDec23shuffle1_600000_filtered.csv
periphery
corner.ArenaTR
corner.ArenaTL
corner.ArenaBR
corner.ArenaBL
center
arena
zone
is.moving
0
60
120
180
240
300
360
seconds

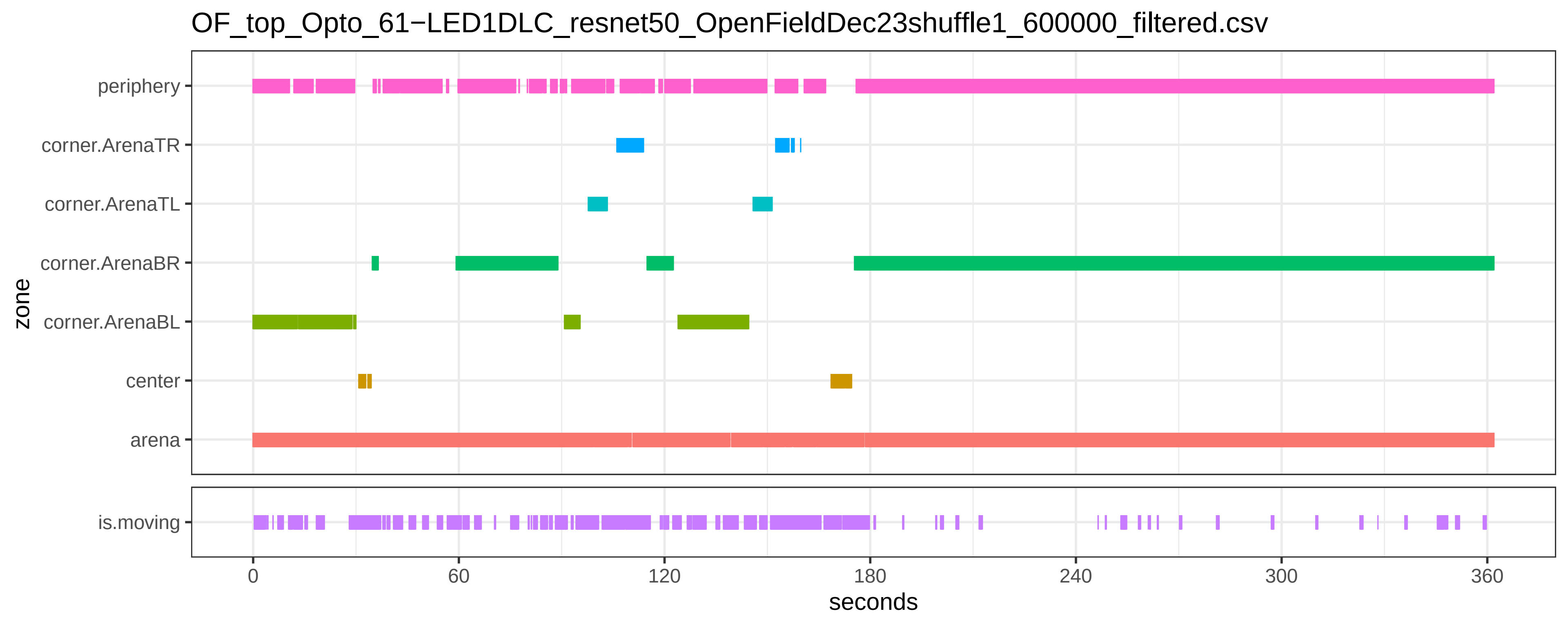

OF_top_Opto_61−LED1DLC_resnet50_OpenFieldDec23shuffle1_600000_filtered.csv
periphery
corner.ArenaTR
corner.ArenaTL
corner.ArenaBR
corner.ArenaBL
center
arena
zone
is.moving
0
60
120
180
240
300
360
seconds

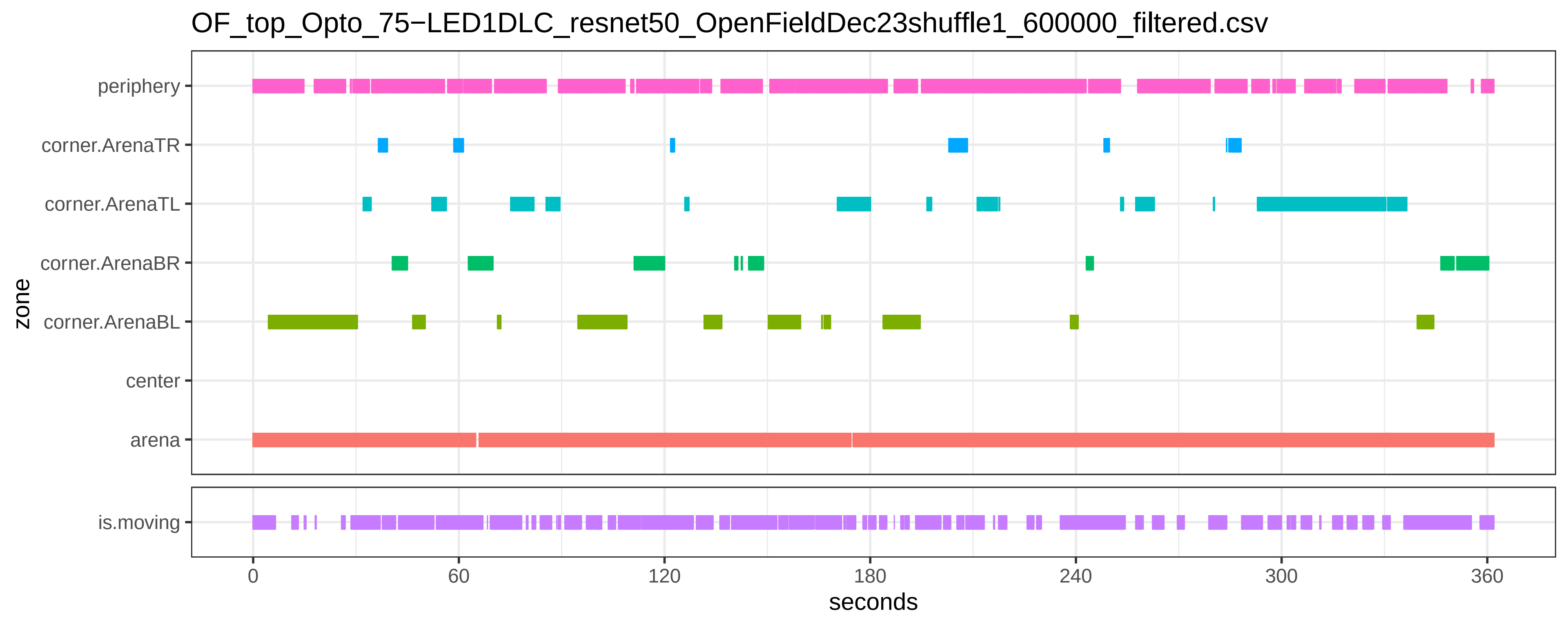

OF_top_Opto_75−LED1DLC_resnet50_OpenFieldDec23shuffle1_600000_filtered.csv
periphery
corner.ArenaTR
corner.ArenaTL
corner.ArenaBR
corner.ArenaBL
center
arena
is.moving
zone
0
60
120
180
240
300
360
seconds

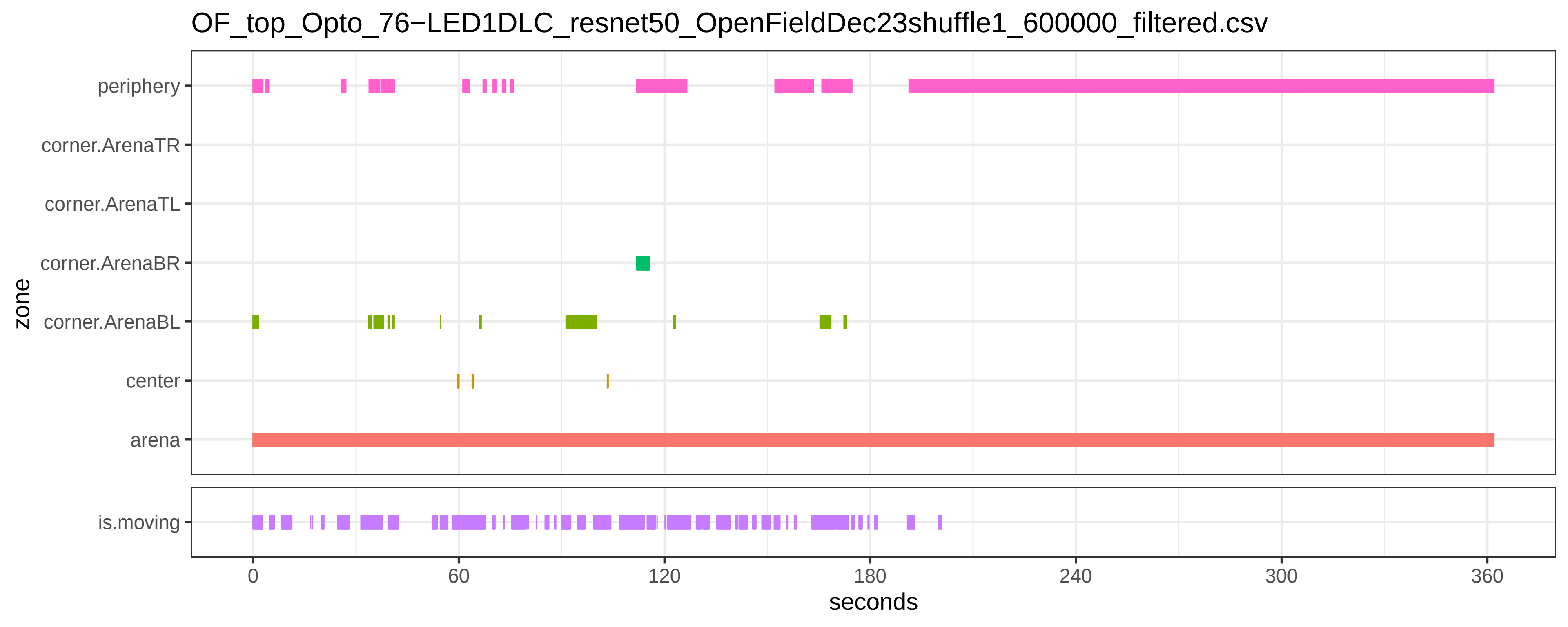
OF_top_Opto_76−LED1DLC_resnet50_OpenFieldDec23shuffle1_600000_filtered.csv
periphery
corner.ArenaTR
corner.ArenaTL
corner.ArenaBR
corner.ArenaBL
center
arena
zone
is.moving
0
60
120
180
240
300
360
seconds

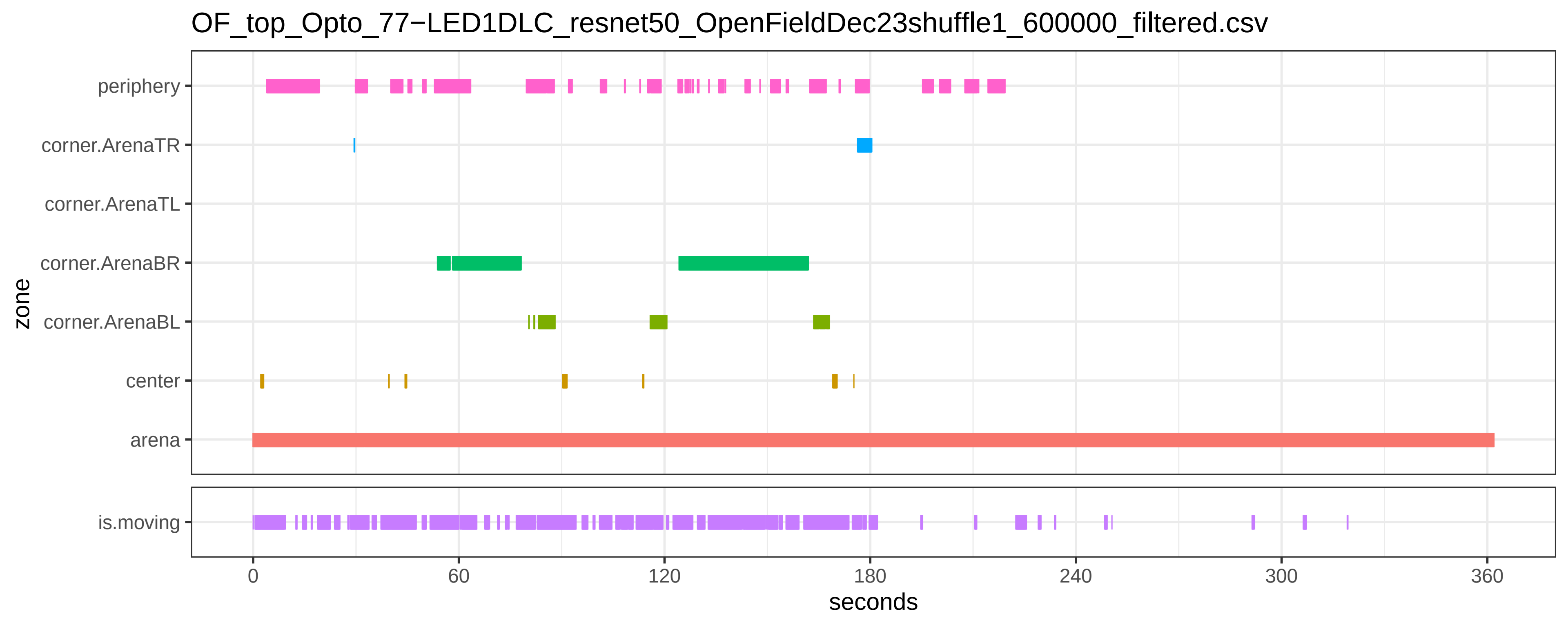

OF_top_Opto_77−LED1DLC_resnet50_OpenFieldDec23shuffle1_600000_filtered.csv
periphery
corner.ArenaTR
corner.ArenaTL
corner.ArenaBR
corner.ArenaBL
center
arena
zone
is.moving
0
60
120
180
240
300
360
seconds
