## SupplementaryFile_S6 for "Chemo- and optogenetic activation of hypothalamic Foxb1-expressing neurons and their terminal endings in the rostral-dorsolateral PAG leads to tachypnea, bradycardia, and immobility"

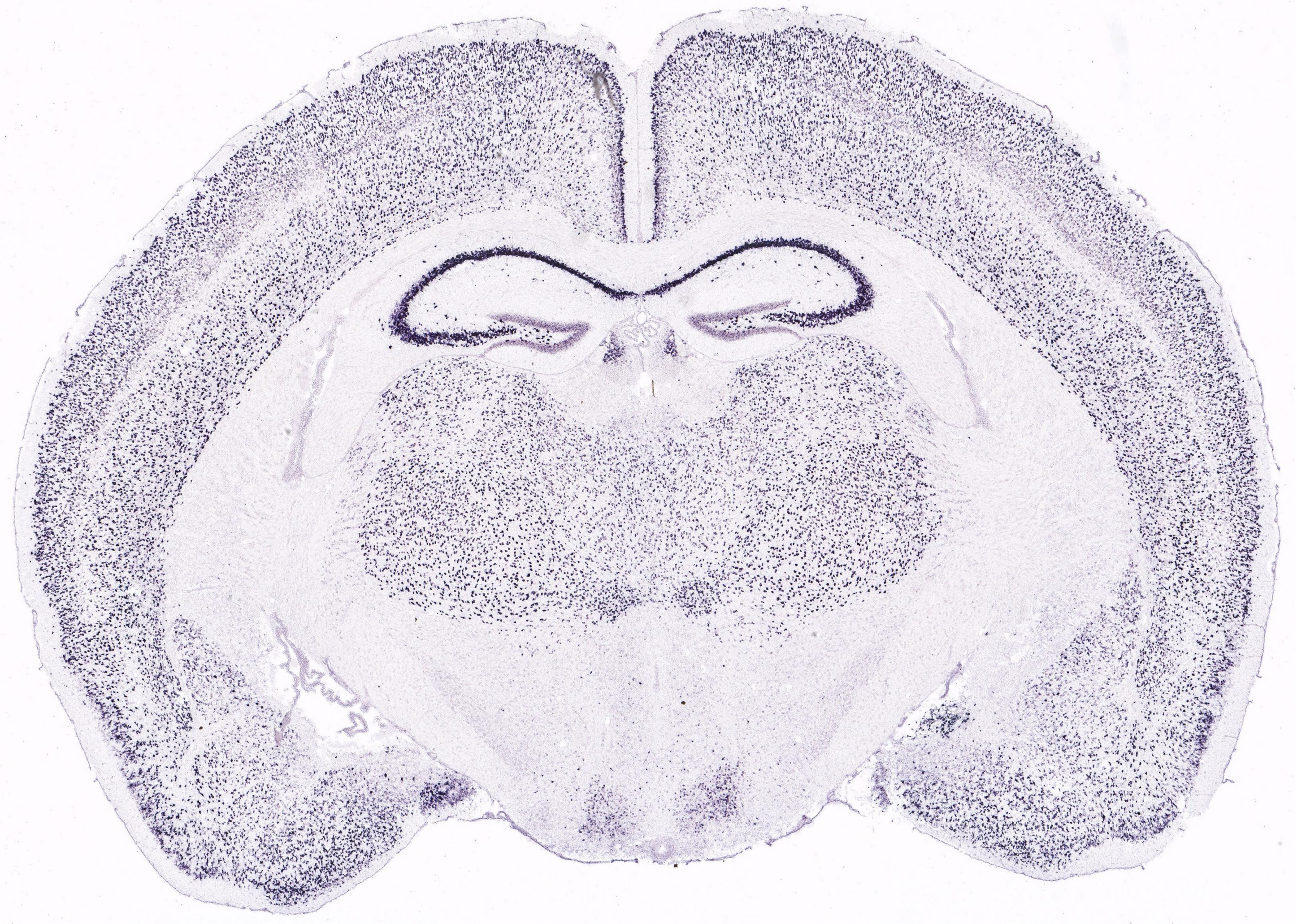
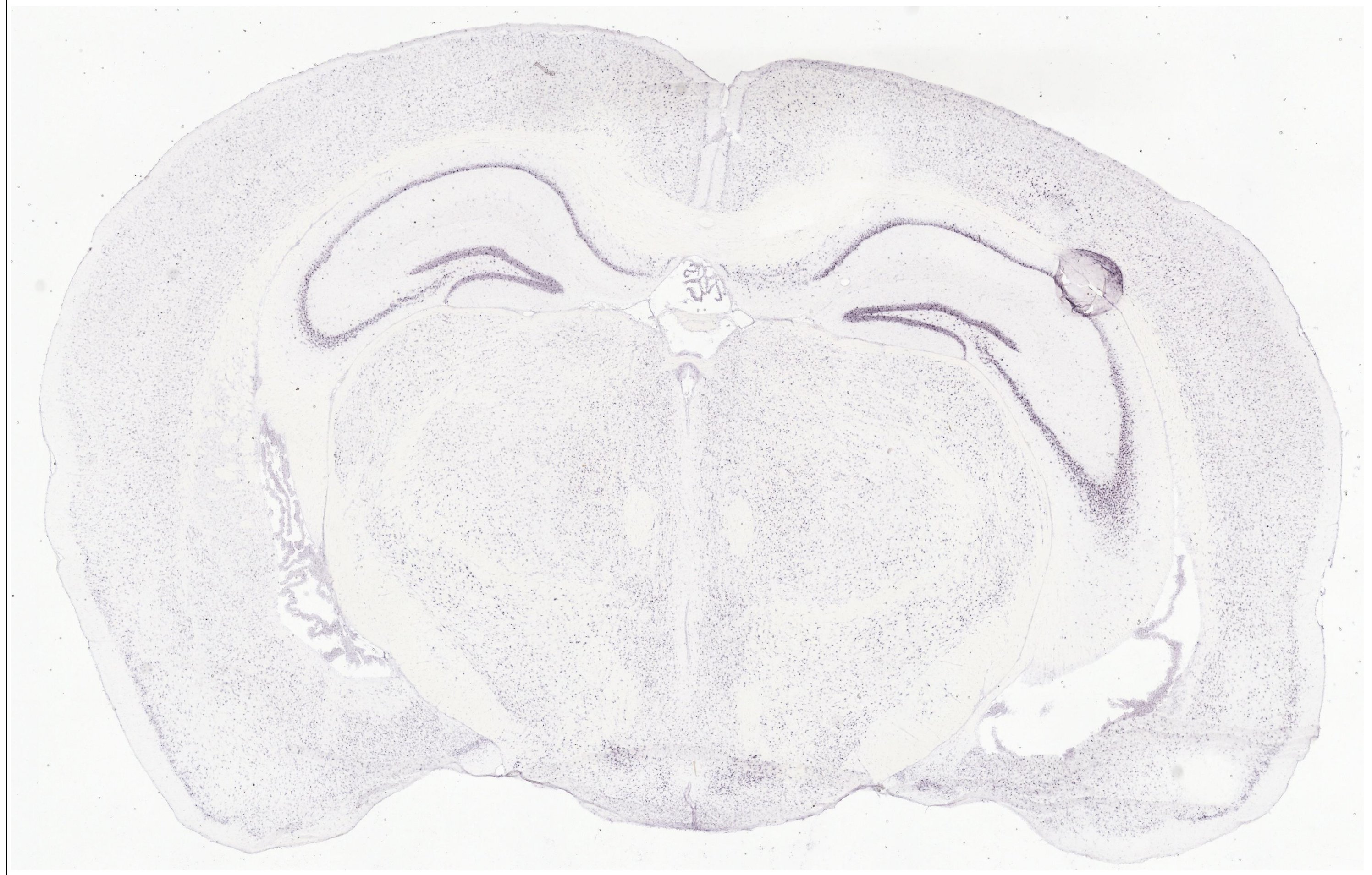
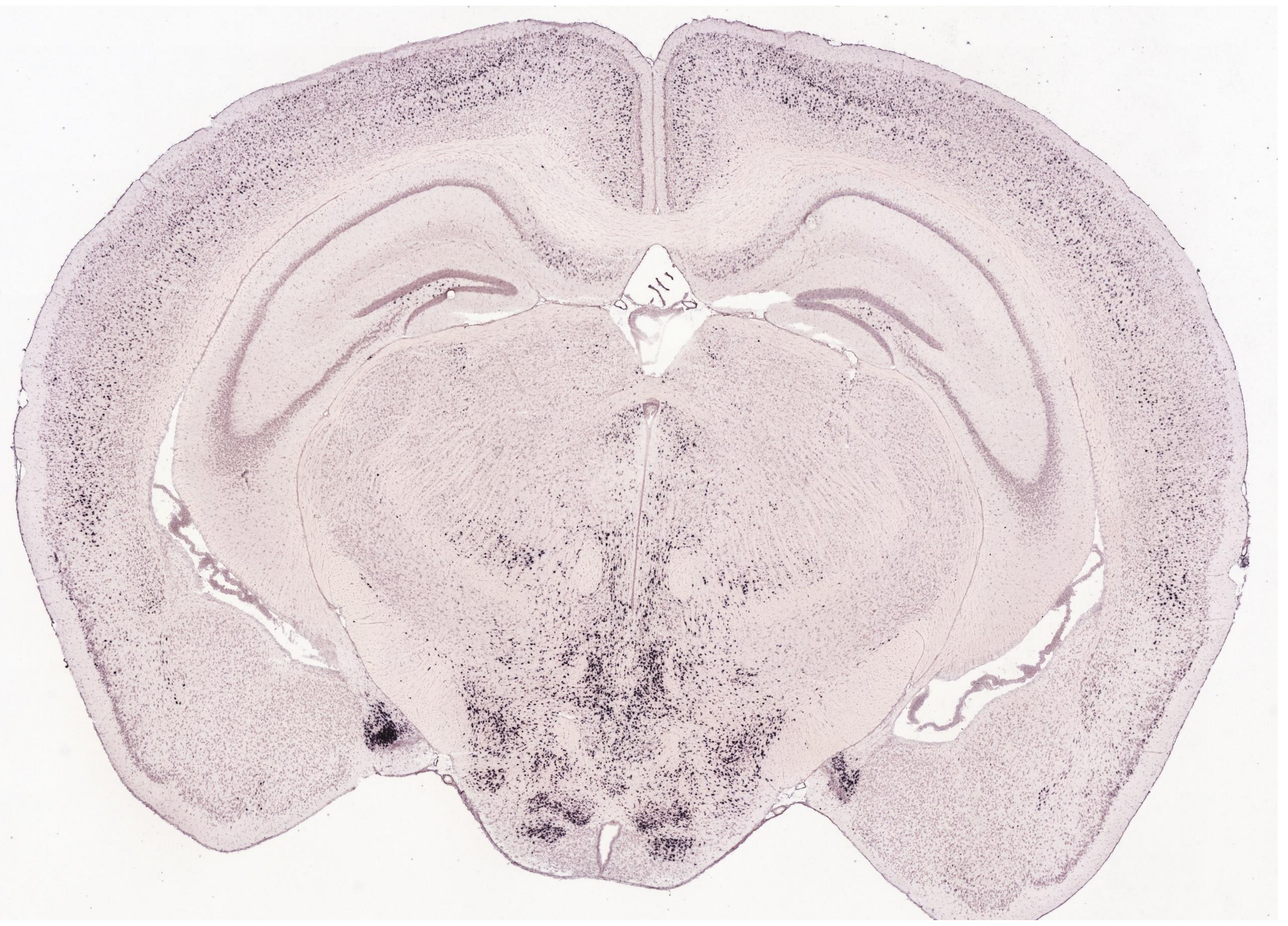
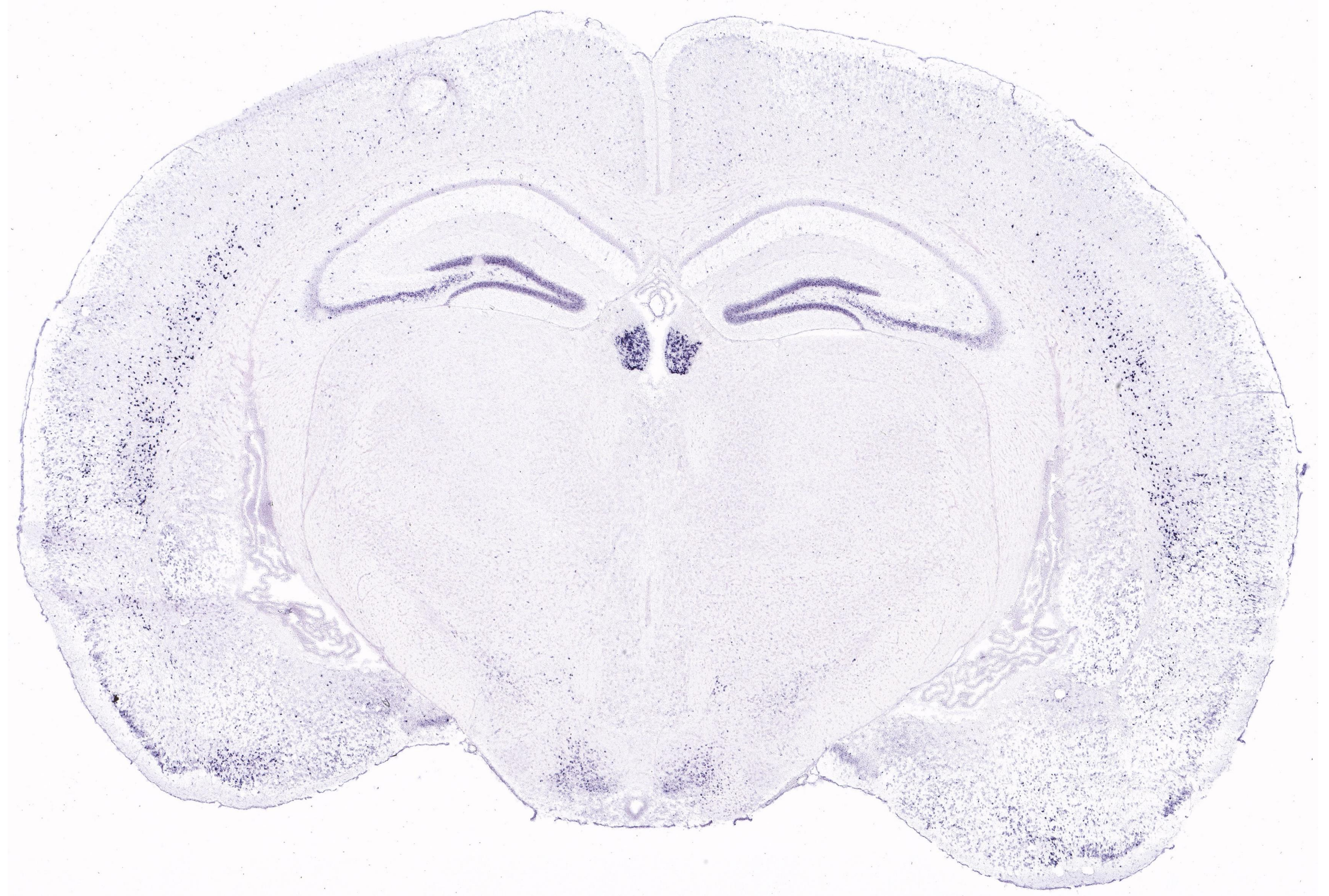
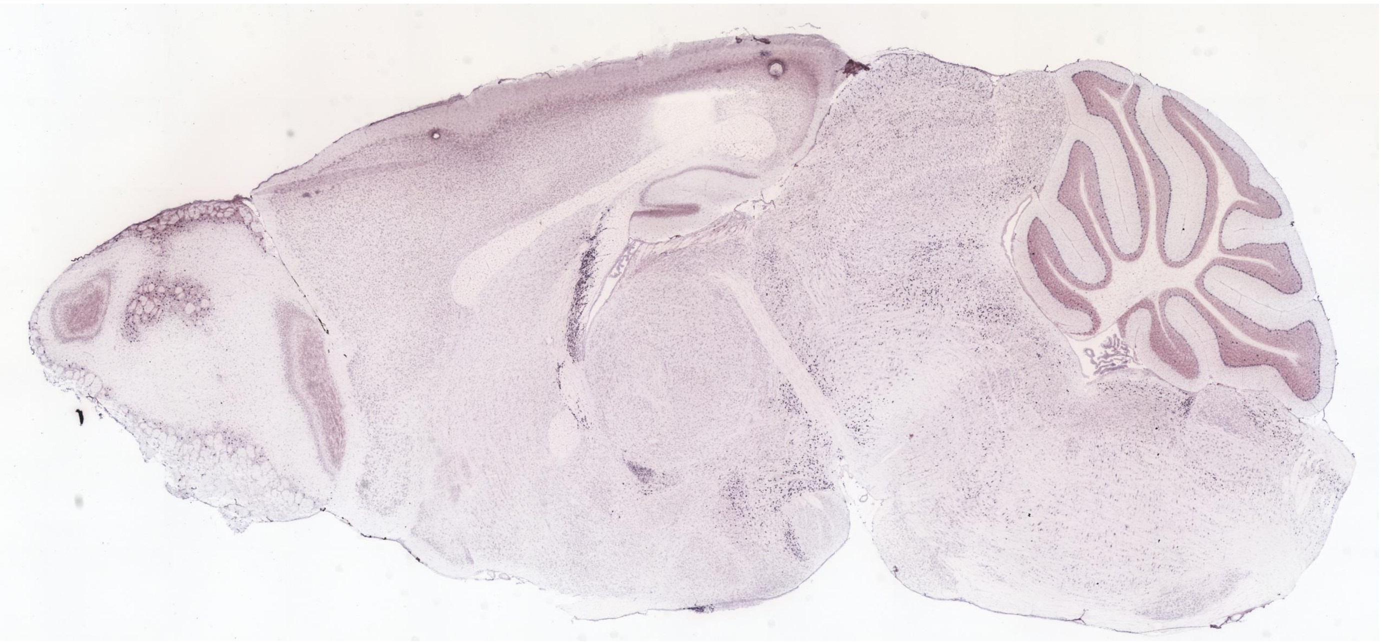
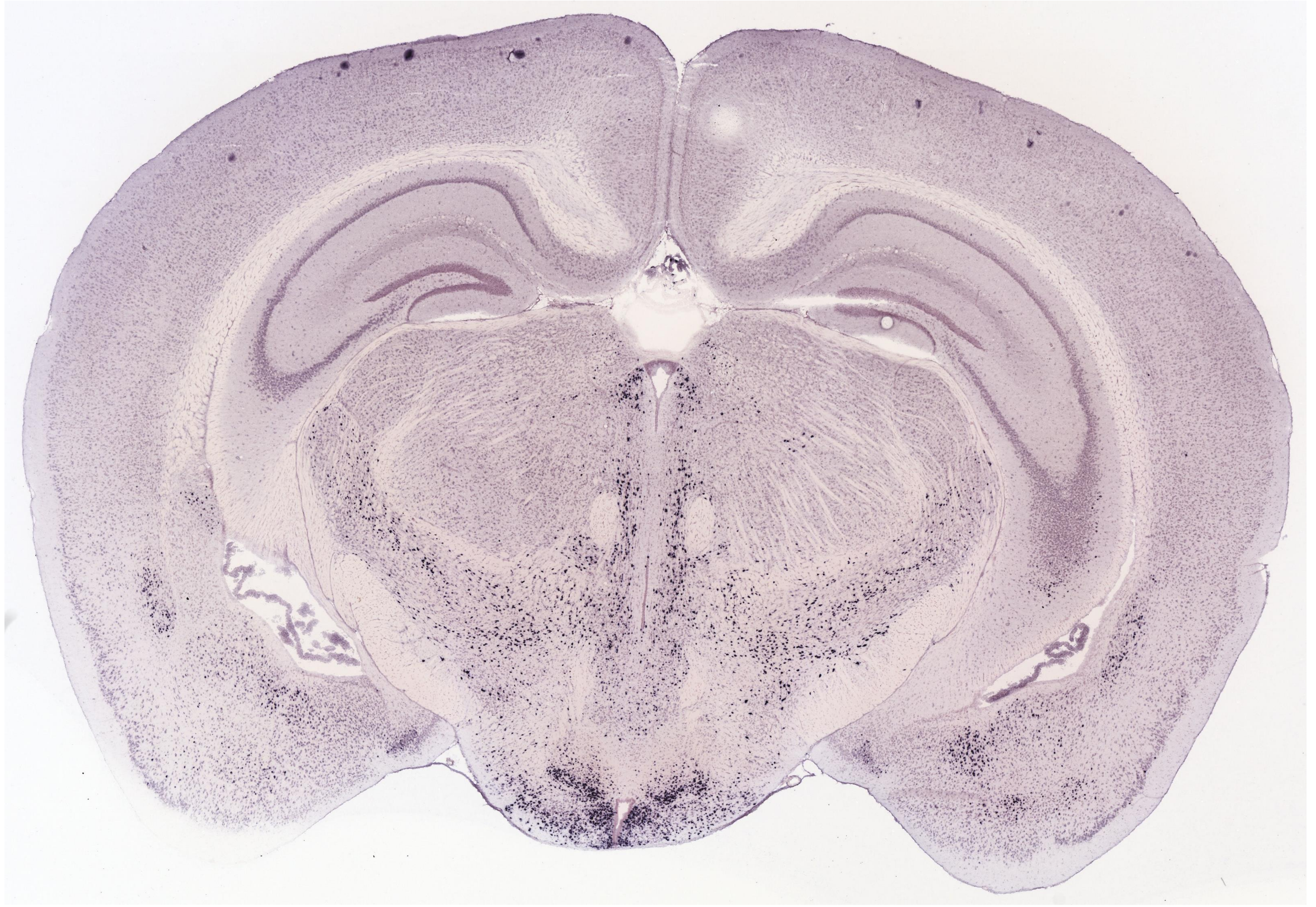
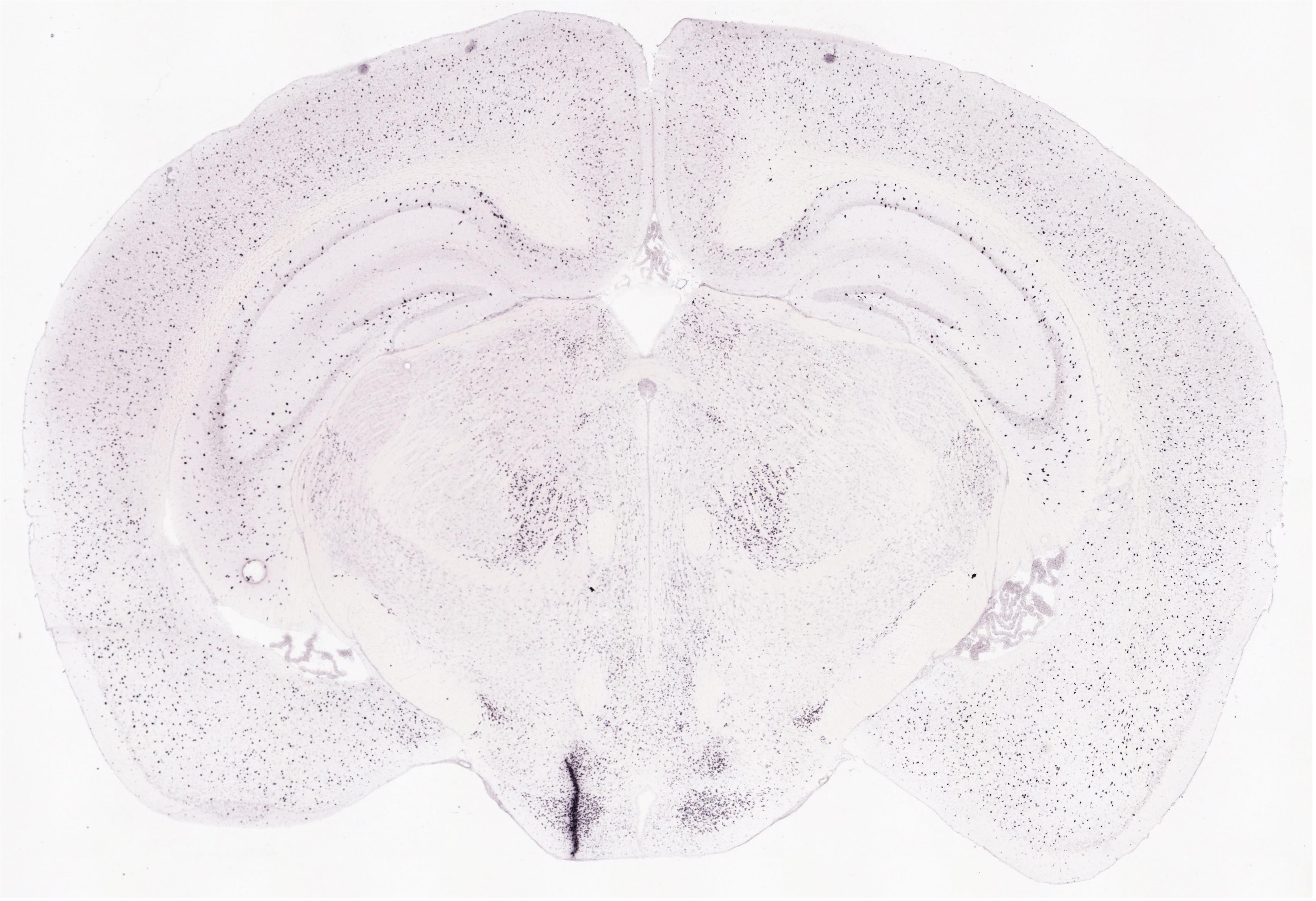
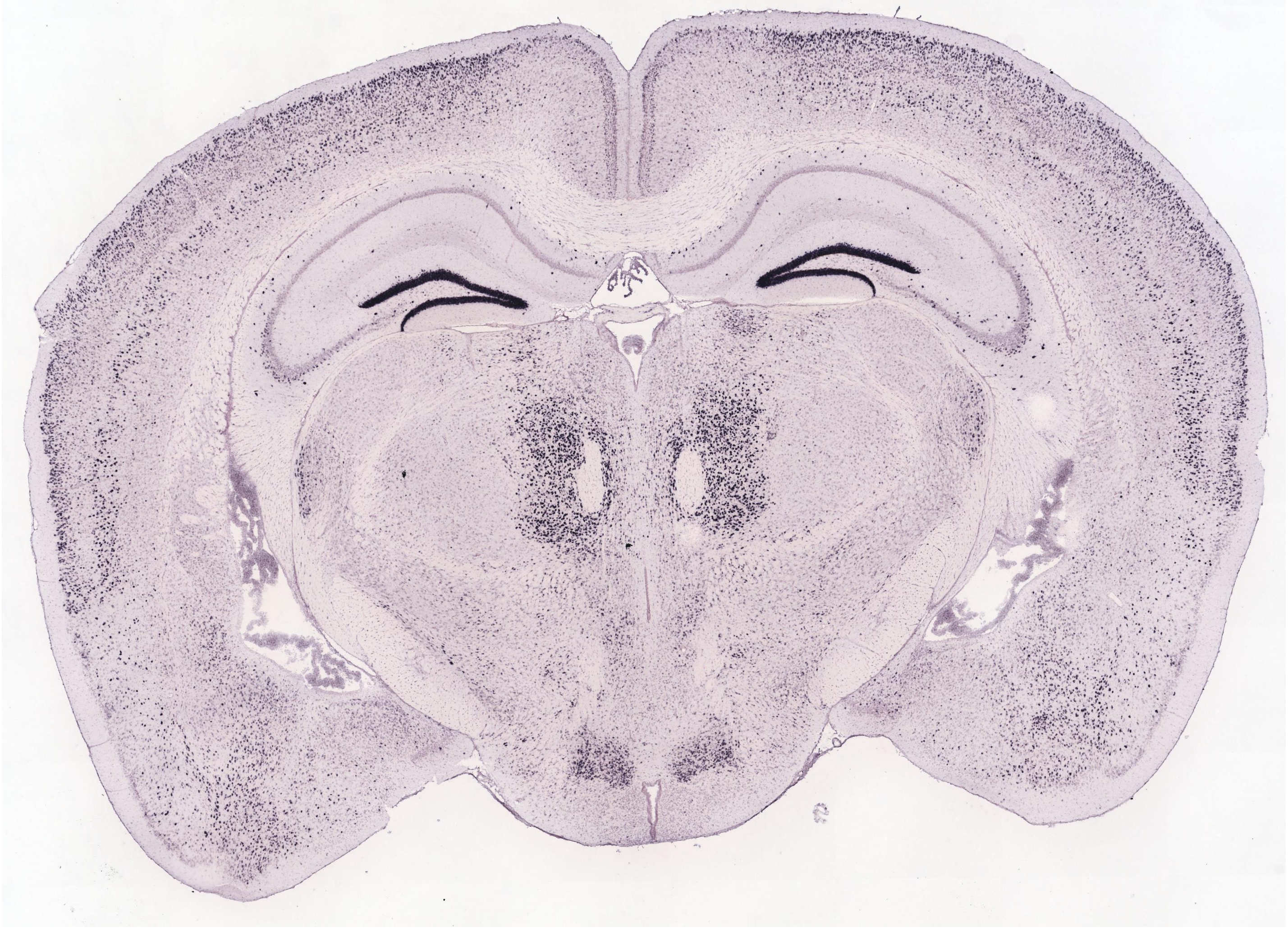
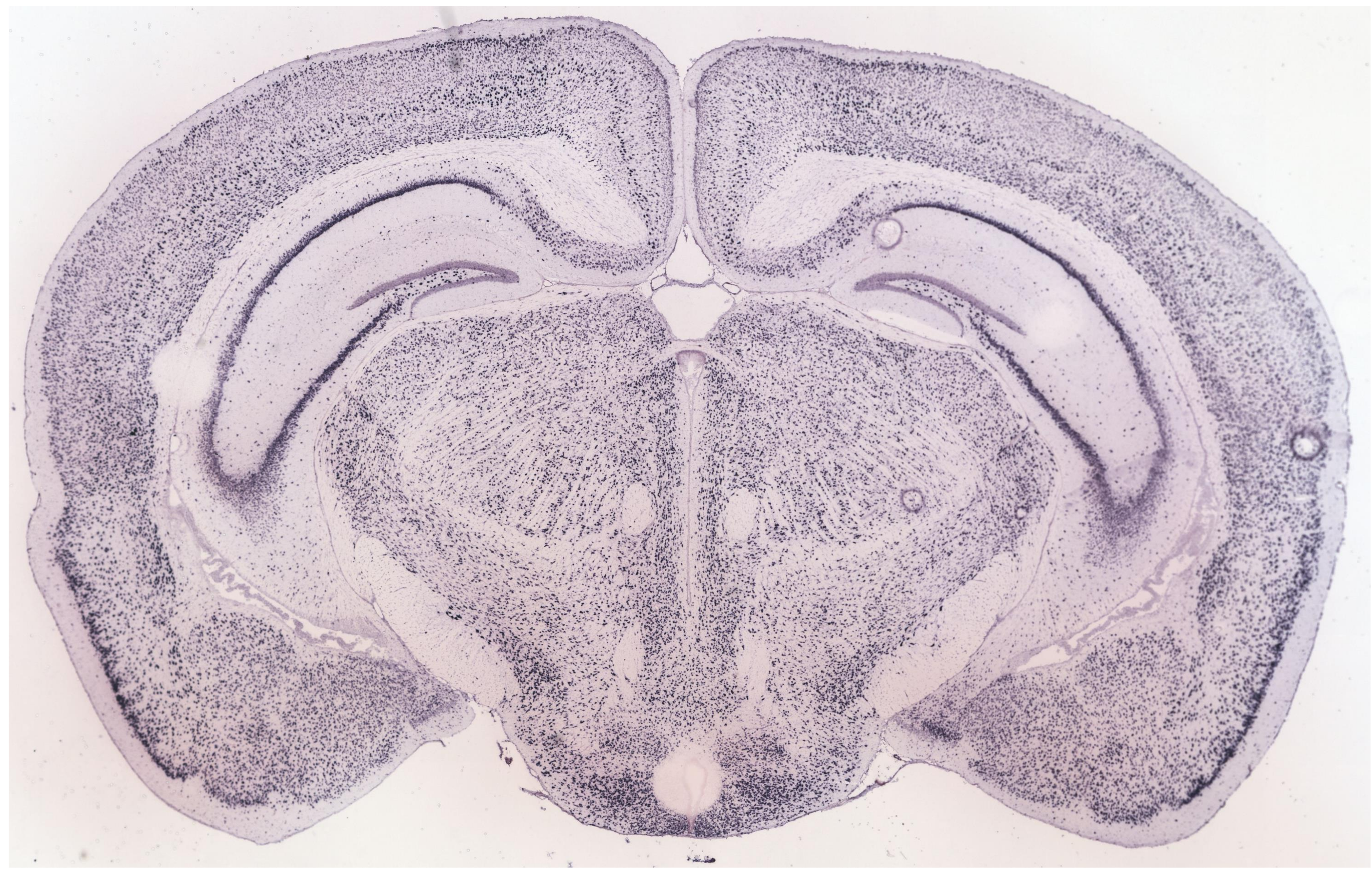
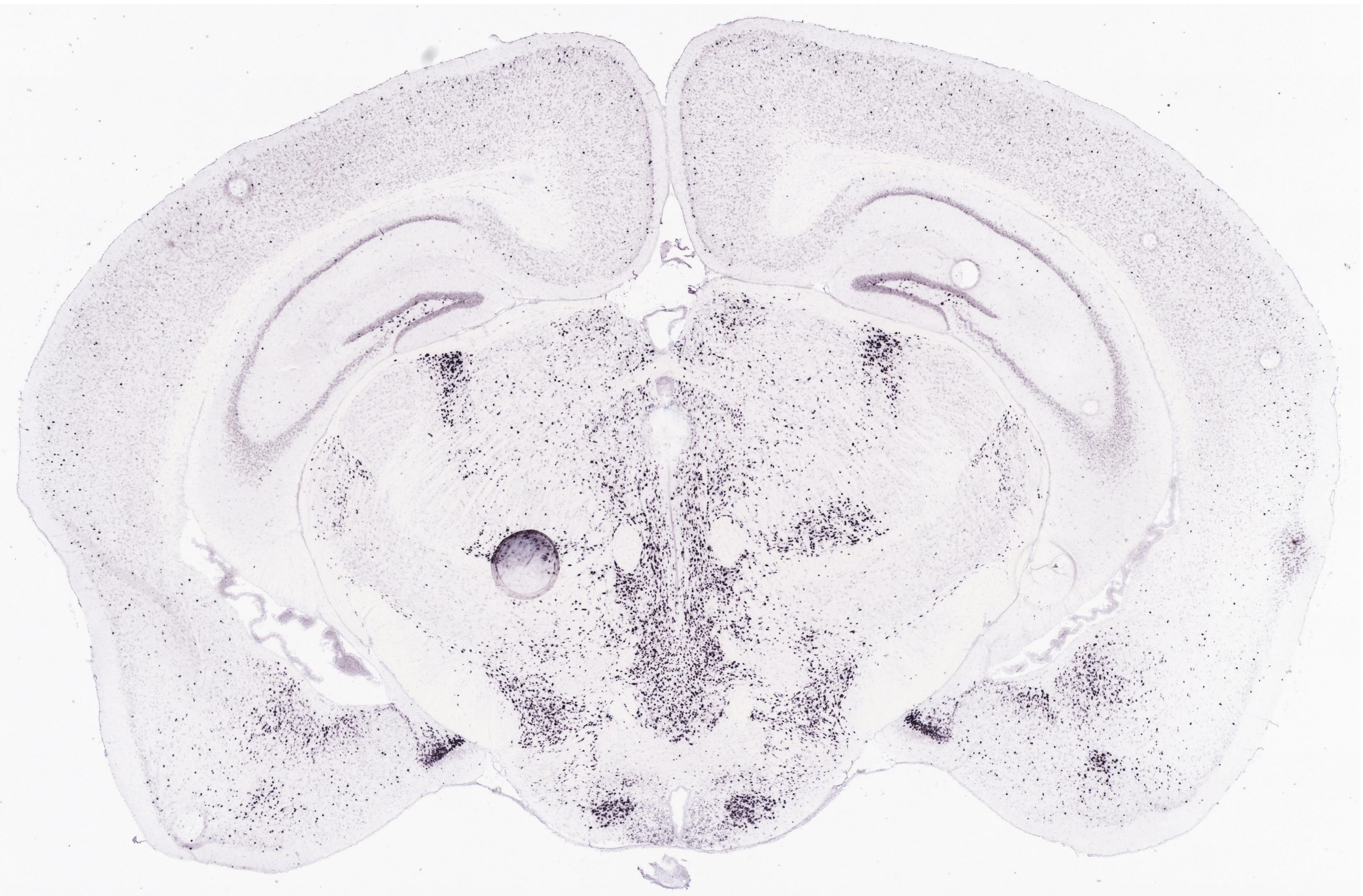
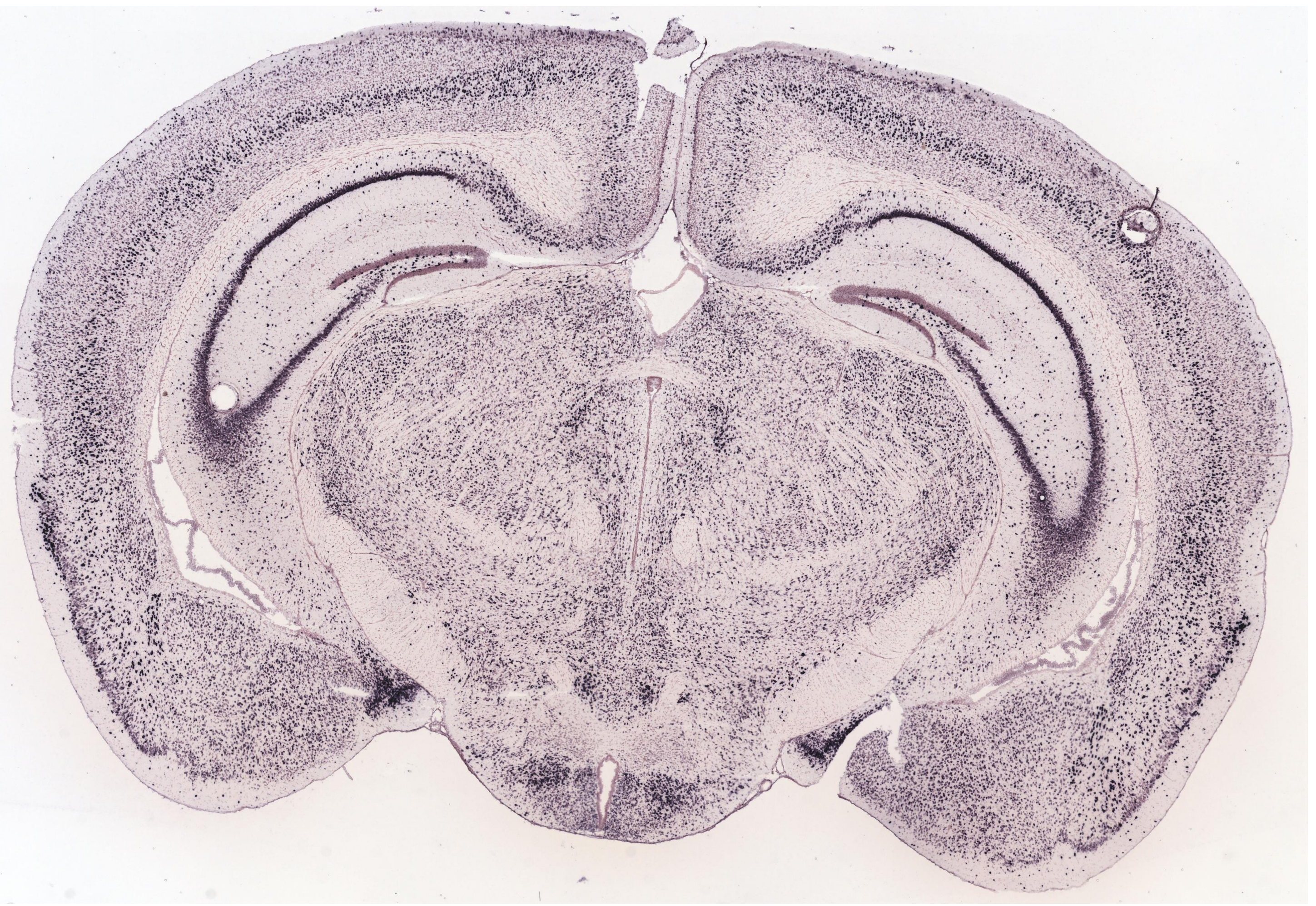

ISH data from Allen Brain Atlas (level of premammillary nuclei).

From top left to bottom right: CCK, Foxb1, Adcyap1, Synpr, Ebf3, Dlk1, Nxph1, Stxbp6, Gap43, Calb2, Rab3c

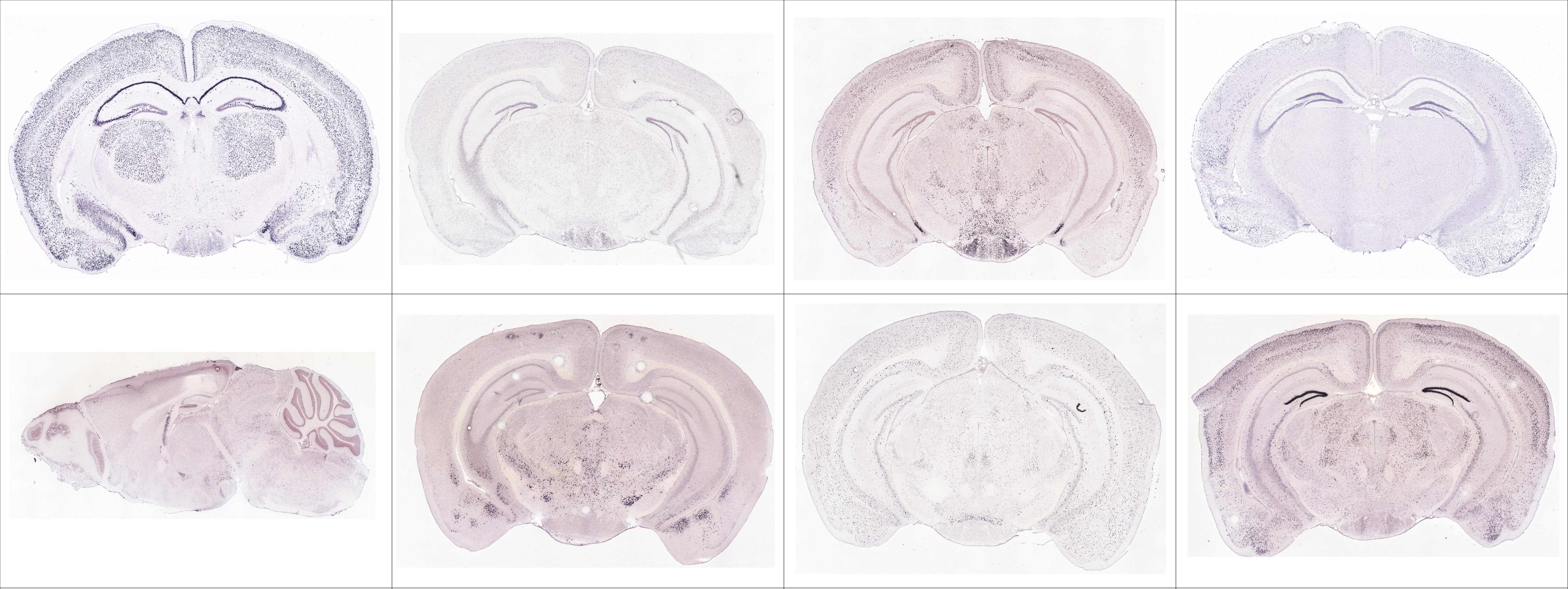

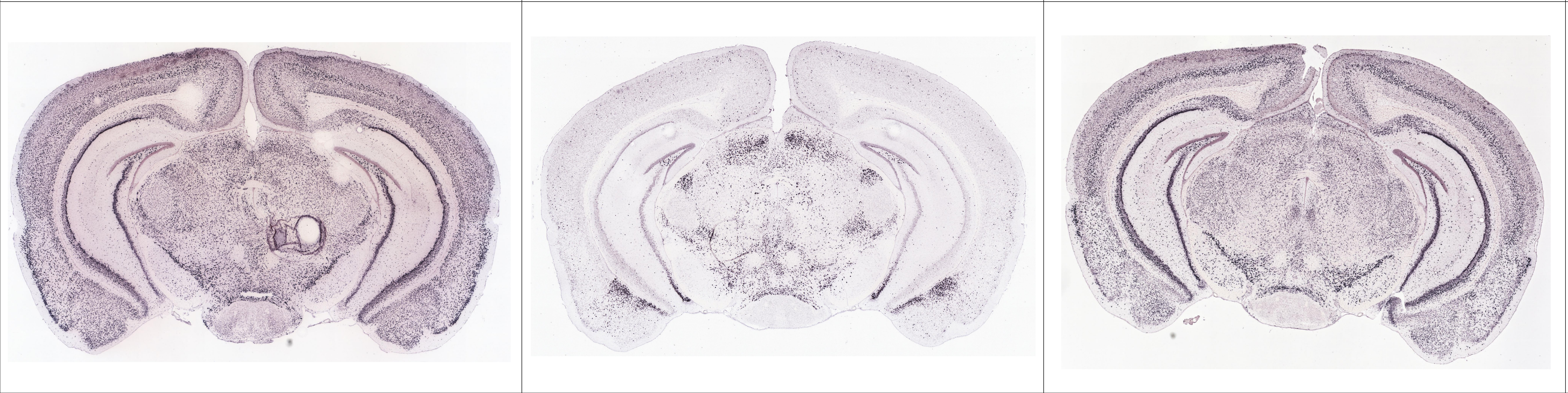

ISH data from Allen Brain Atlas (level of mammillary body).

From top left to bottom right: CCK, Foxb1, Adcyap1, Synpr, Ebf3, Dlk1, Nxph1, Stxbp6, Gap43, Calb2, Rab3c
