## Supplementary material for "Chemo- and optogenetic activation of hypothalamic Foxb1-expressing neurons and their terminal endings in the rostral-dorsolateral PAG leads to tachypnea, bradycardia, and immobility": SuplementaryFile_S7

### Antibodies and fluorophores used for histological analysis

- Rabbit polyclonal to mCherry
  Abcam plc. (product code: ab167453)
  1:1000 in TBS 0.1M, incubation for 2 days

- Mouse monoclonal [2H2] to c-Fos
  Abcam plc. (product code: ab208942)
  1:2000 in TBS 0.1M, incubation for 2 days

- Chicken polyclonal to GFP
  Aves Labs, Inc. (product code: GFP-1020)
  1:200 in TBS 0.1M, incubation for 2 days

- Biotinylated Horse anti-Mouse IgG
  Vector Laboratories, Inc. (product code: BA-2000)
  1:200 in TBS 0.1M, incubation for 2 hours

- Alexa Fluor 647-conjugated Streptavidin
  Jackson ImmunoResearch (product code: 016-600-084)
  1:200 in Tris pH 8.2, incubation for 2 hours

- Cy2-conjugated AffiniPure Donkey anti-Chicken IgG
  Jackson ImmunoResearch (product code: 703-225-155)
  1:200 in Tris pH 8.2, incubation for 2 hours

- Cy3-conjugated AffiniPure Donkey anti-Rabbit IgG
  Jackson ImmunoResearch (product code: 711-165-152)
  1:200 in Tris pH 8.2, incubation for 2 hours

- DAPI
  Life Technologies Corporation (product code: D1306)
  1:5000 in TBS 0.1M, incubation for 5 minutes
